# Circular van Krevelen diagram for visualizing metabolic pathways

**DOI:** 10.1101/2025.05.31.657198

**Authors:** Shuzhao Li

## Abstract

Emerging biochemical data require effective pathway visualization. However, traditional metabolic maps were based on manual layouts, falling behind new scientific discoveries. Automated visualization of metabolic pathways has been investigated heavily, but resulted in few consensuses. We report here a new approach based on circularized van Krevelen diagram. This method is based on chemical principles, providing the consistency critical for community exchanges and collaborations. Example applications include metabolomics data interpretation and a new set of metabolic maps.

## Introduction

The metabolic chart of pathways is a quintessential presence in biochemistry education and research, first drawn by Donald Nicholson in 1955 (Nicholson, 2001). In the genomics era, KEGG (Kyoto Encyclopedia of Genes and Genomes, Kanehisa et al, 2025) metabolic maps have been a constant fixture in visualizing metabolic pathways. The KEGG maps were also laid out manually, then computerized using fixed coordinates. It is laborious to generate pathway maps manually. More importantly, these manual maps become restrictive in the age of large-scale molecular discoveries and computational intelligence. For example, modern metabolomics and exposomics deal with a large number of new compounds that are not in classical pathways (Mitchell et al, 2024, Chi et al, 2025, David et al, 2021), demanding computable solutions that support continuous expansion of reactions and pathways. Automated metabolic pathway visualization has been investigated extensively (Kutmon et al, 2015, Pon et al, 2015, Sidiropoulos et al, 2017, Paley et al, 2021, Agrawal et al, 2024). However, no widely accepted solution has emerged. Because unlike empirical gene networks, metabolic pathway visualization needs to connect with the underlying chemical principles. As the past efforts were optimized for print media, a new design is required to aid computing biochemistry in the big data era.

Biochemistry on earth is based on carbon and involves only a limited number of key elements. Dirk Willem van Krevelen popularized the plots of H:C ratio versus O:C ratio in industrial chemistry since 1950s (Wu et al, 2004, Kim et al, 2003). As metabolomics has now emerged as high-throughput measurements of small molecules in biological systems (Li, ed., 2020), van Krevelen diagrams have been applied to visualizing metabolomics data (Tziotis et al, 2011, Kew et al, 2017, Brockman et al, 2018, Laszakovits et al, 2021). Major classes of metabolites can be separated to some extent on a classical van Krevelen diagram (Brockman et al, 2018), indicating potentials to support automated layout of metabolic pathways. We postulate here that a coordinate system based on elemental ratios can support metabolic pathway visualization at the interface between humans and computers. A new approach based on circularized van Krevelen diagram is developed and its application to human metabolic maps is presented in this paper as proof of principle. This method is principled, consistent and applicable to broad metabolomics data.

## Results

### Emerging biochemical data motivate chemistry based pathway visualization

The traditional manual layout of a global metabolic map has reached its limit. Typical human genome scale metabolic models (GSMMs, Brunk et al, 2018, Robinson et al, 2020) contain about 4,000 compounds, while the Human Metabolome Database (HMDB, Wishart et al, 2022) catalogs over 200,000 compounds (**Figure 1a**). We use the term compounds here, as some of them do not fall into the definition of biological metabolites. This scale of data necessitates automated layout for visualization and computer aided navigation. We propose in this paper that modified van Krevelen plots on a polar coordinate can serve as a foundation of layout algorithms (examples in **Figure 1b, c**).

**Figure 1:**
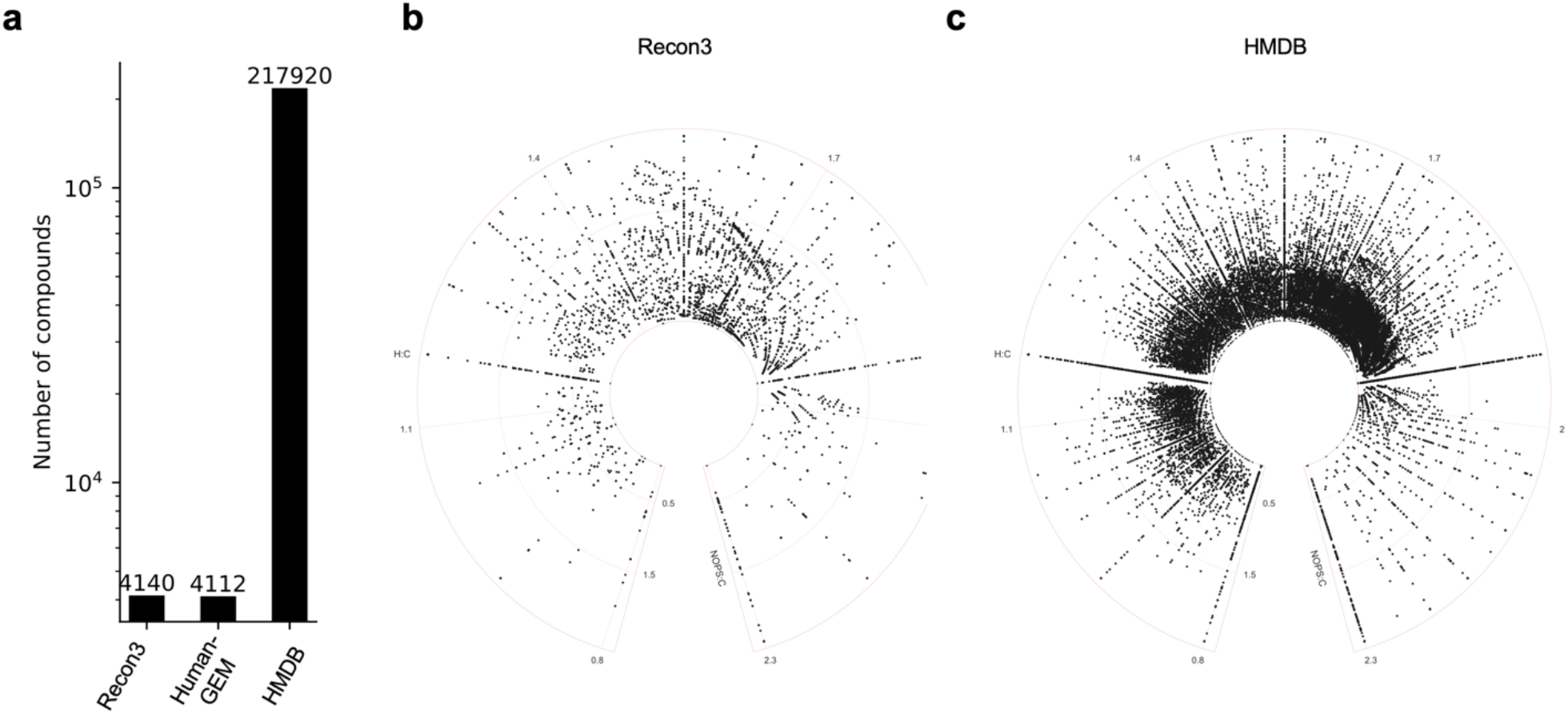
Numbers of compounds in human biochemistry. **a**. numbers of compounds in genome-scale metabolic models (RECON3, human-GEM) and a leading metabolite database (HMDB). **b.** Circular van Krevelen plots for compounds in RECON3 and HMDB, respectively.

The original van Krevelen diagram uses O:C ratio versus H:C ratio. For example, in a molecule with formula C_26_H_43_NO_5_, the H:C ratio is 43/26 = 1.654, and O:C ratio 5/26 = 0.192. Such ranges are limited for biological molecules. The distributions of H:C and O:C ratios in all compounds in a GSMM (Recon3) and HMDB are shown in **Figure 2**. In applications to biological molecules, nitrogen and other elements are often important. One can add the N:C ratio as another dimension (Wu et al, 2004). Here, we introduce a new NOPS:C ratio, weighted by atomic numbers. For the above molecule, its NOPS:C ratio is calculated as (1*14 + 5*16 + 0*31 + 0*32)/(26 * 12) = 0.301. The NOPS:C ratios have a broader distribution that those of O:C (**Figure 2**, right), which benefits space utilization in layout algorithms. Because isotopes have different masses, this weighted metric can be easily extended to isotope analysis and tracing experiments.

**Figure 2.**
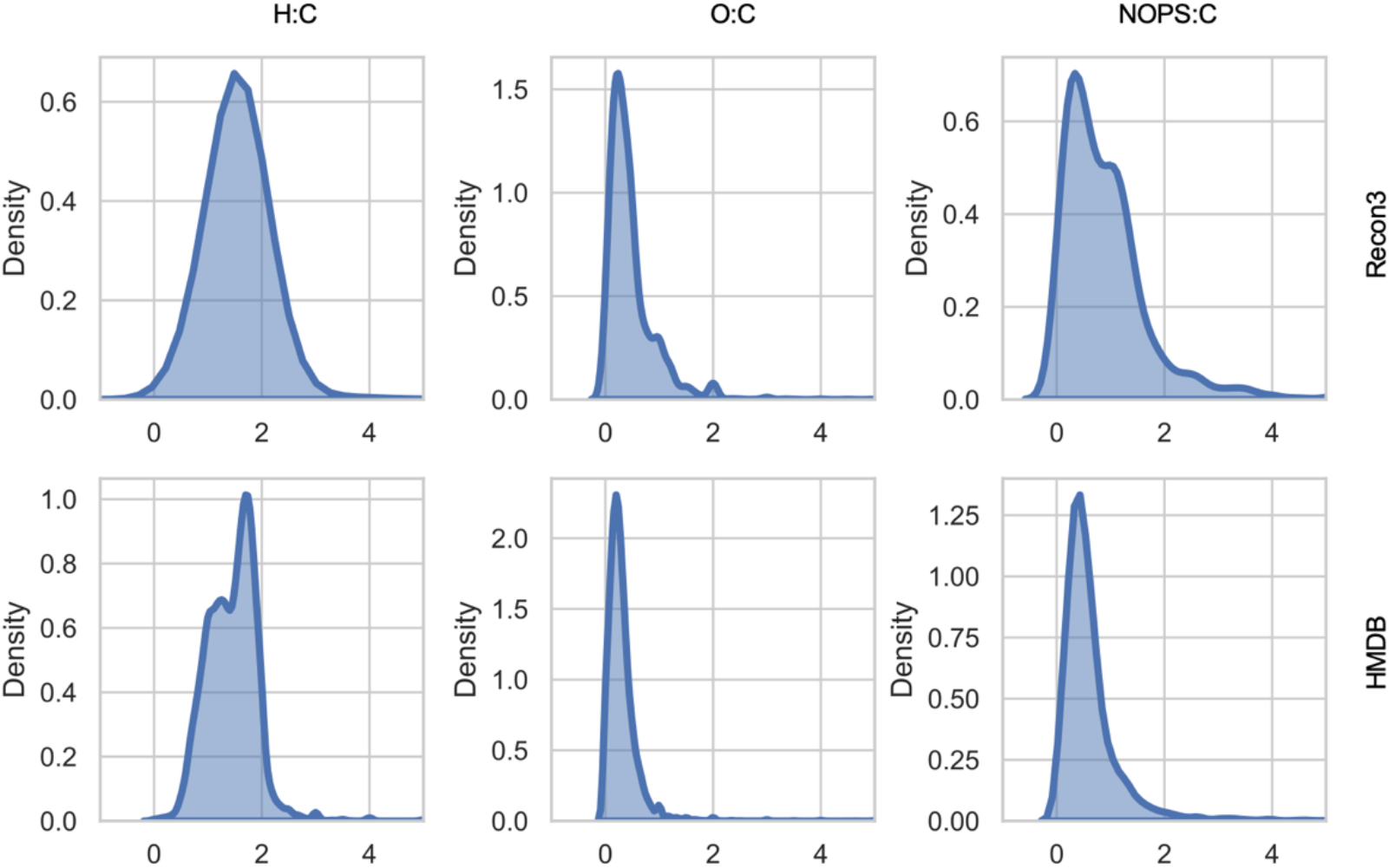
Distribution of elemental ratios in Recon3 (top) and HMDB (bottom).

Using one example pathway (aminosugar metabolism), the difference between O:C and NOPS:C ratios as y-axis is illustrated in **Figure 3a, b**. The metabolic reactions in this pathway have now been added to the plots as directed edges, indicating reasonable usability for pathway visualization. When more compounds are added, three concerns related to scalability arise: a) poor space utilization with some very high-density and very low-density areas; b) vaguely defined boundaries, lack of reference landmarks for visual navigation; and c) difficult navigation of the reaction edges (connectors). Converting the Cartesian coordinates to polar coordinates brings clear improvements. Here, H:C ratio is used as angular coordinate and NOPS:C ratio for radial distance when data are projected onto a polar coordinate system (global examples in examples in **Figure 1b, c**; smaller examples in **Figure 3d, e**). The circular plots have less waste and cluttering, and reduce the space used by connectors, The outer boundary defines a constant space for the chemicals, easier to locate a compound than on an open plain. The most important is that each compound has a fixed coordinate on these plots, a consistency factor critical for community exchanges and collaborations, akin to geological maps and astronomical systems.

**Figure 3:**
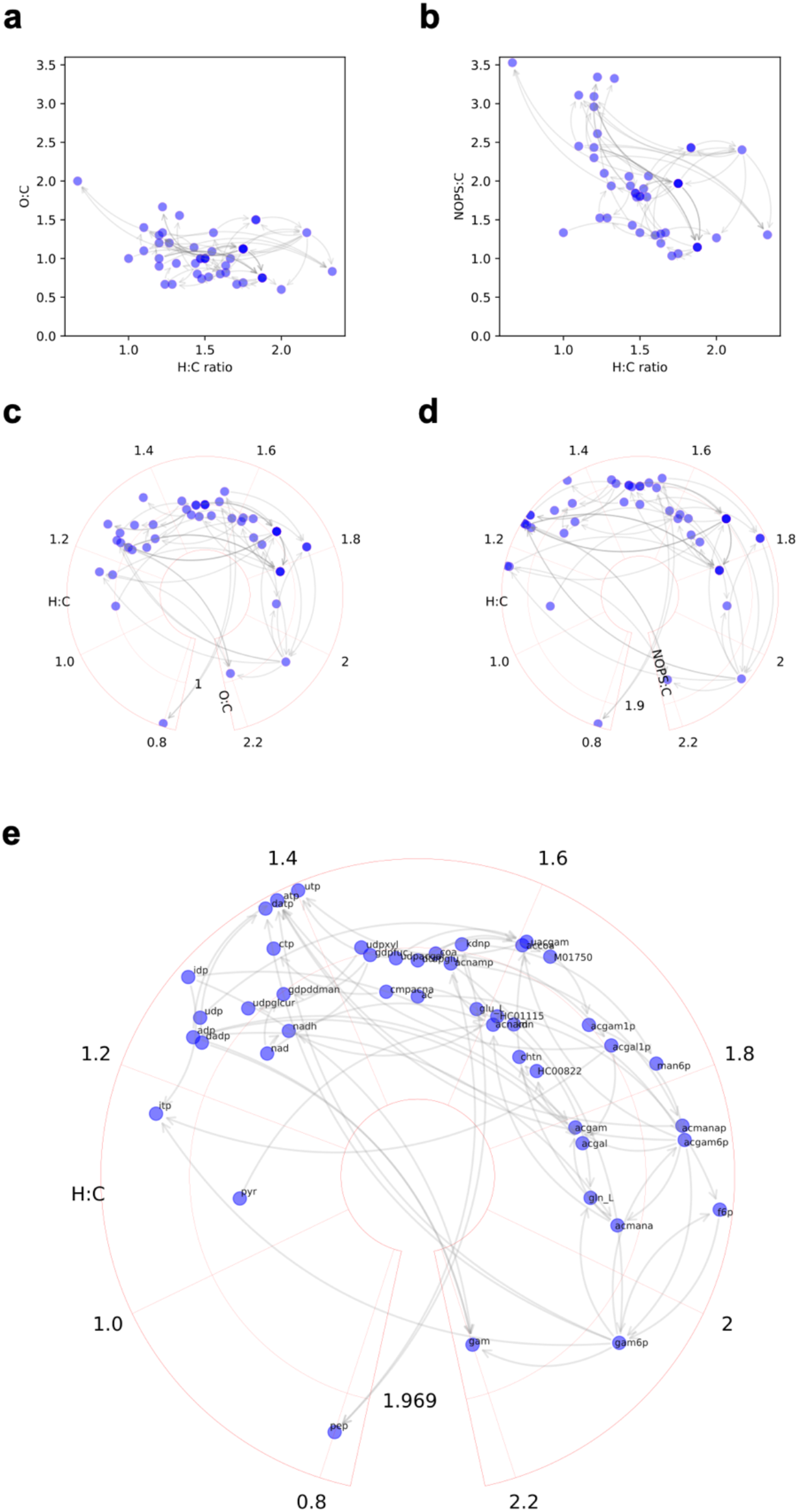
Modifications of van Krevelen plots by weighted element ratios and polar coordinates. Aminosugar metabolism pathway is used as examples. Each node a metabolite, each edge a reaction step. Conventional van Krevelen plot on Cartesian coordinates using O:C ratios or weighted NOPS:C ratios as y-axis (**a, b**). Circular van Krevelen plot on polar coordinates using O:C ratios or weighted NOPS:C ratios as y-axis (**c, d**). **e**. Circular van Krevelen plot with compound labels.

### Circular van Krevelen plots applied to biological data

The layout of compounds on a circular van Krevelen plot is based on their chemical formulas. This is illustrated on human caffeine metabolism in **Figure 4**. As caffeine is degraded to smaller molecules, distinct groups of compounds are shown by similar structures, and decreasing order of H:C ratio. Because the reactions remove methyl groups that have more hydrogen atoms than carbons in the purine rings. As discussed above, new data drive expansion of pathways, which in turn demands automated and logical accommodation in pathway visualization. A new caffeine metabolite, 1,3,7-Trimethyldihydrourate, has been recently identified in humans (Liu et al, 2021). Adding this new metabolite is trivial, as its coordinates on circular van Krevelen plot are calculated directly from its formula (blue in **Figure 4**).

**Figure 4:**
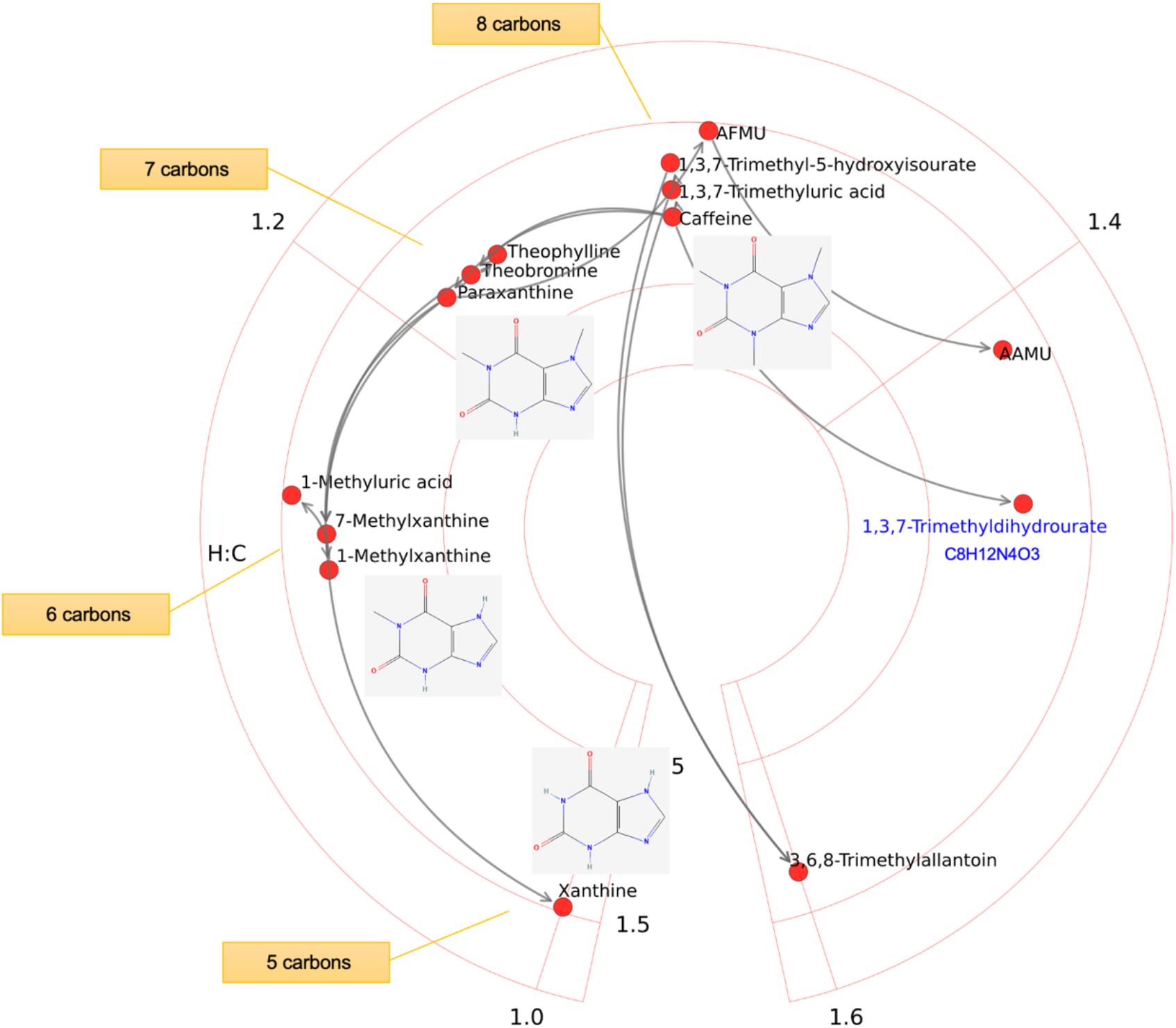
Caffeine metabolism in humans. A recently identified caffeine metabolite in humans is colored in blue.

Using circular van Krevelen plots, the full set of metabolic pathway maps in Recon3 are generated as proof of principle (**Suppl File 1**). Three lipid pathways are shown in **Figure 5a**, to exemplify their visual patterns, due to many molecular species varied by regular chain units.

**Figure 5:**
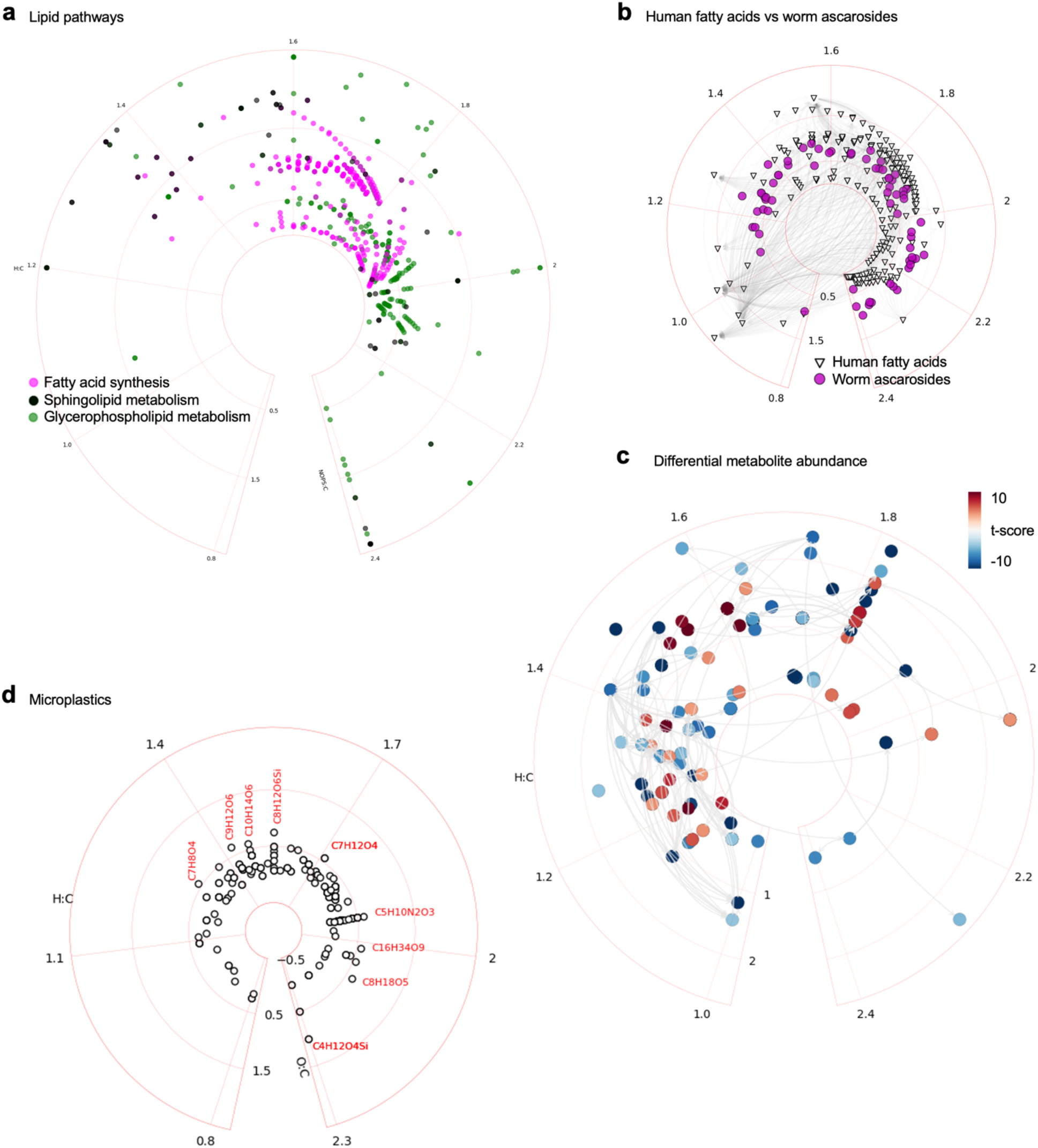
Example applications of circular van Krevelen plots. a) Three pathways from the Recon3 model. b) Comparative metabolomics on human and worm pathways (Artyukhin et al, 2018). Differential metabolite abundance on a metabolic network in immune cell activation (Li et al, 2013). Plotting micro- and nano-plastics found in human samples (Dusza et al, 2022).

Such patterns can be used to identify compounds, and they are easily visible in the global maps (**Figure 1b, c**). Since many lipids in the same subclass are catalyzed by the same enzyme, such reaction edges can be optimized in the future by abstraction. Interesting, nematodes use ascarosides for their biology. These ascarosides are different from human fatty acids but also have side chains of varying length. Artyukhin et al. (2018) employed metabolomics analysis to reveal that beta oxidation impacts large number of these ascarosides. The result from Artyukhin et al. (2018) is overlaid here on human pathway of fatty acids, enabling visual comparison (**Figure 5b**). These pathway plots can be customized in many ways to suit the types of data and data analysis. In typical studies of comparing metabolite abundance, difference can be coded by colors in heat maps. Similarly, differential metabolite abundance can be mapped to a circular van Krevelen plot of metabolic network. **Figure 5c** shows a reproduction of the data from Li et al (2013) using this style. The plots are flexible enough to be applied to micro- and nano-plastics found in human samples (**Figure 5d**, Dusza et al, 2022).

### Implementation of circular van Krevelen plot in the lcvk Python package

We have implemented this approach in a software package based on the widely used Matplotlib library in Python, where the polar projection serves as a basis for circular van Krevelen plots. Chemical formula manipulation is using mass2chem. The main wrapper function in ‘lcvk.polarPlot.cplot_LCVK_pathway’ provides arguments to control many features directly. For example, changing a few parameters can optimize the plots significantly (**Figure S1**). A set of Jupyter notebook examples and templates are provided as part of the software repository.

A direct limitation of these van Krevelen plots is that the coordinates are based on chemical formulas. By default, compounds with the same chemical formula have the same coordinates. This is undesired in many situations. This can be overcome by additional methods, e.g. replacing the formula node by a pie chart representing cohabitant molecules. A simple optimization method is included in the software package to reduce overlap nodes (e.g. **Figure S1d**). Once the data size is big enough, the density problem is unavoidable on a 2-D format. Interactive environments, including common web based JavaScript plots, will be needed to navigate multiple layers of data.

## Discussion

New data from metabolomics and exposomics have exceed the scale of GSMMs greatly. This demands new visualization tools that are built on consistent and extensible principles. We have reported here a new approach based on circularized van Krevelen diagram to generate metabolic maps. This approach leads to a fixed coordinate for each metabolite, which is readily calculated by its chemical formula. This enables reliable exchange of data and models, and offers the flexibility to visualize any chemical network. In global metabolic maps (Figure 1b, c), the distribution of molecules follows their chemical properties and molecules of the same H:C ratios become natural landmarks. Therefore, the method is well suited for -omics scale analysis, and future development for interactive navigations and scientific discoveries (Lozano et al, 2021).

## Data and Code Availability

Source code, example data and notebook templates are freely available at https://github.com/shuzhao-li-lab/Li_CVK_diagram.

## Author Contributions

Project design, Software development, Data analysis and writing: SL.

## Acknowledgements

This work was in part supported by NIH grant R01 AI149746 (NIAID) and ARPA-H award D24AC00345.

## Supplemental Information

**Supplemental File S1**. Full list of pathway figures in Recon3 by default lcvk plot function.

**Supplemental Figure S1:**
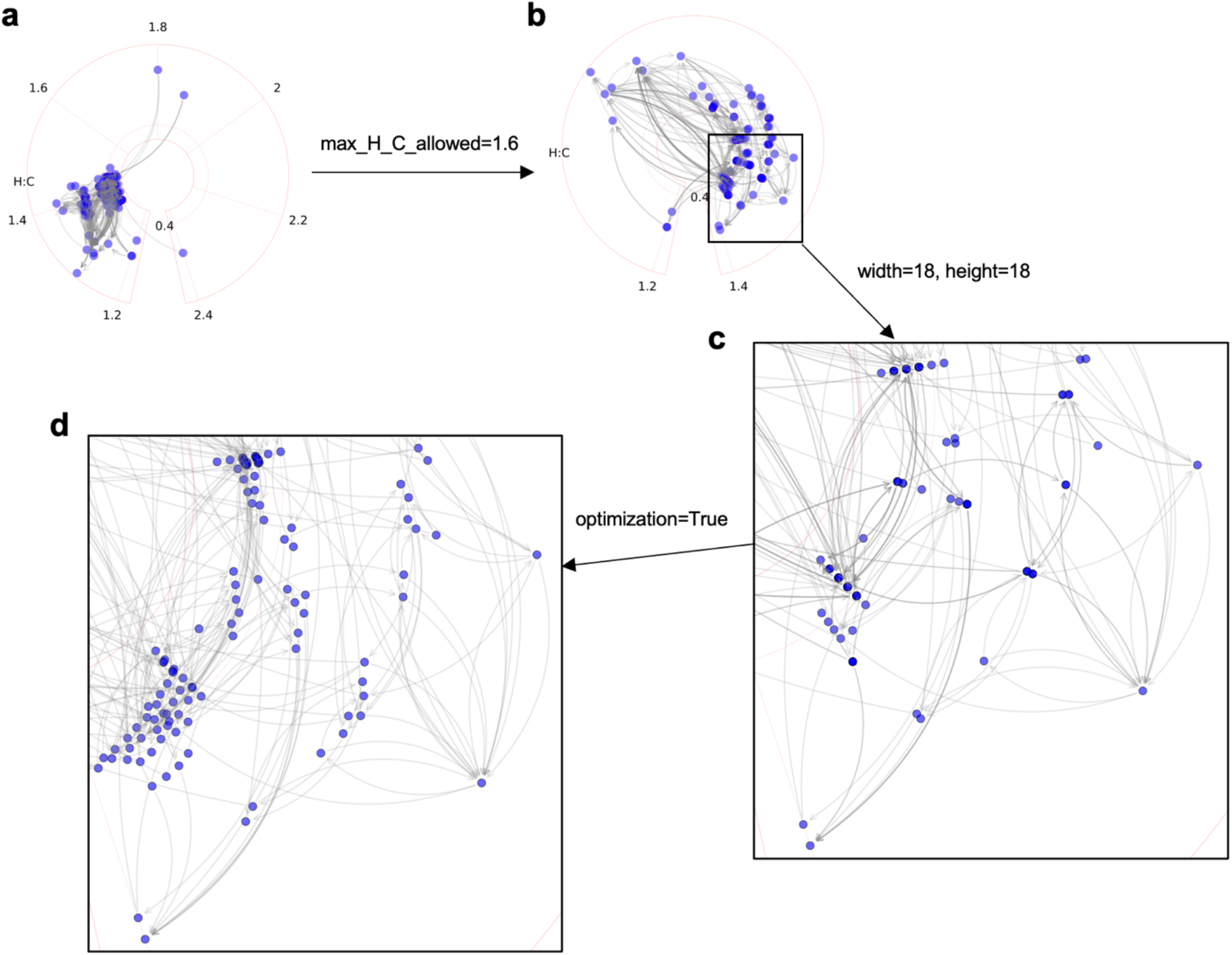
Scalable and zoomable pathway and network visualization. Optimization of this plot (**a**) by rescaling H:C range (**b**), by changing figure size (**c**), and by an optimization function to avoid overlap nodes (**d**).

