## Supplementary figures and images for "Circular van Krevelen diagram for visualizing metabolic pathways"

### Alanine and aspartate metabolism.pdf

#### Alanine and aspartate metabolism

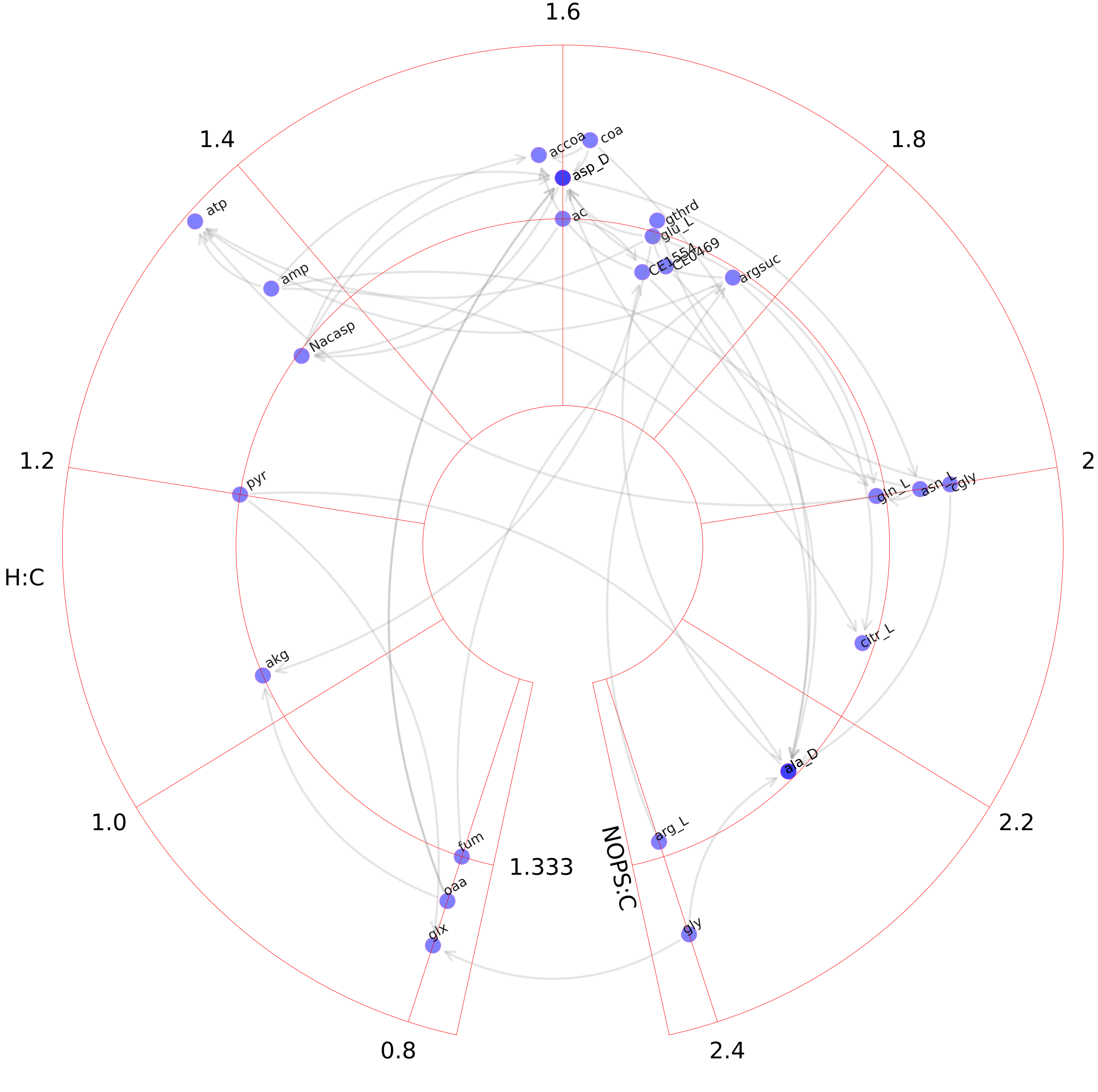

### Alkaloid synthesis.pdf

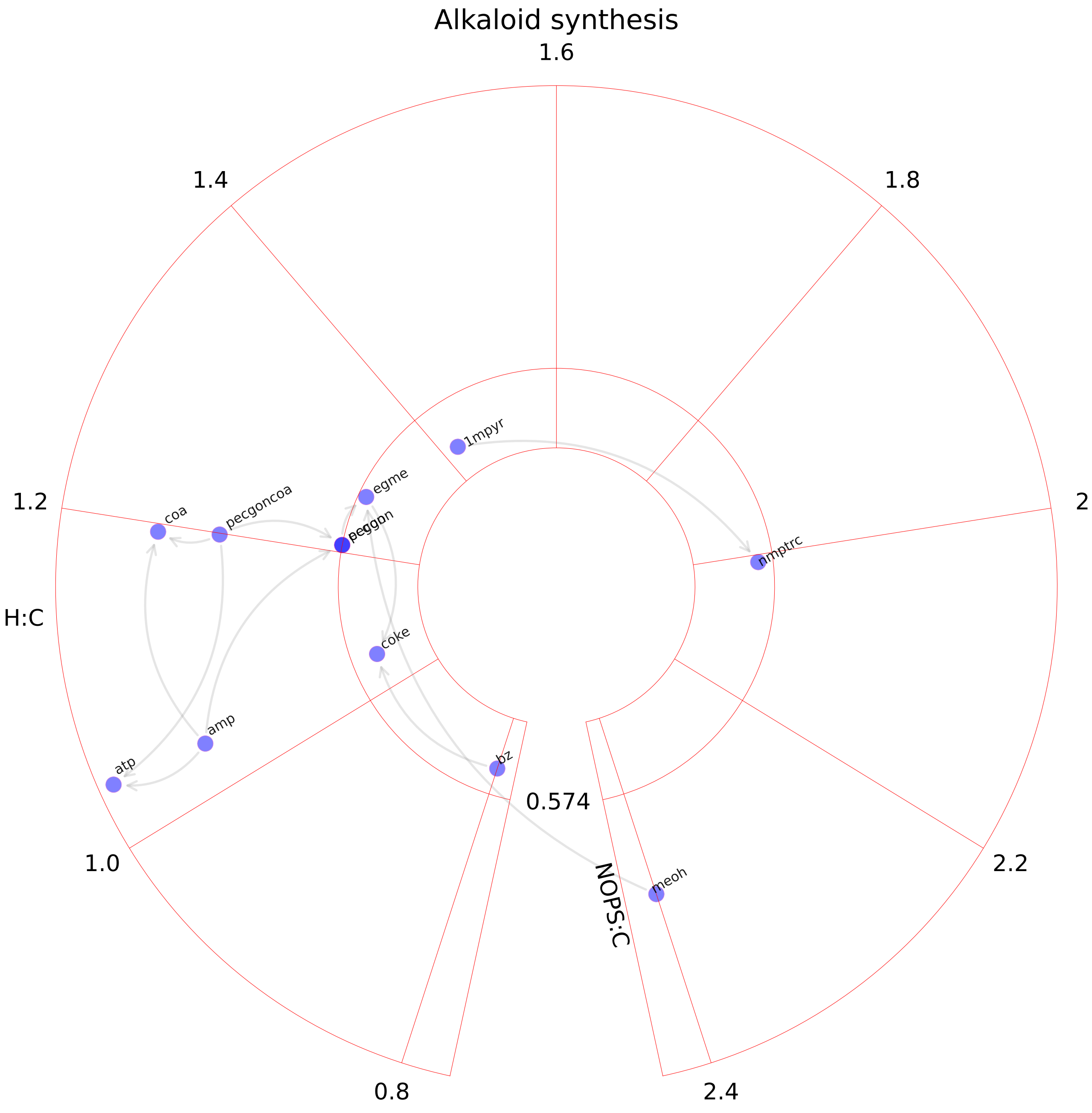

### Aminoacyl-tRNA biosynthesis.pdf

#### Aminoacyl-tRNA biosynthesis

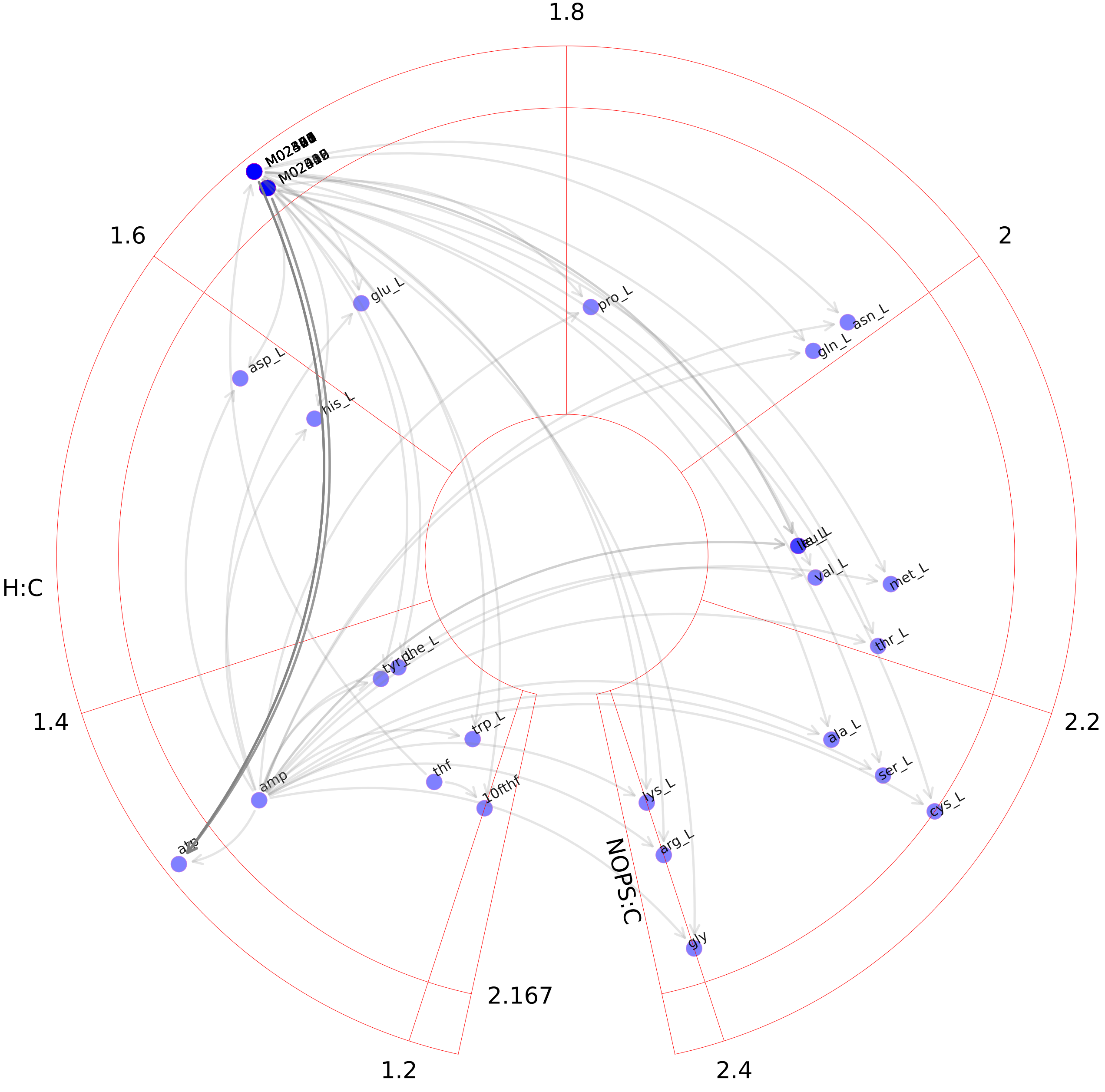

### Aminosugar metabolism.pdf

#### Aminosugar metabolism

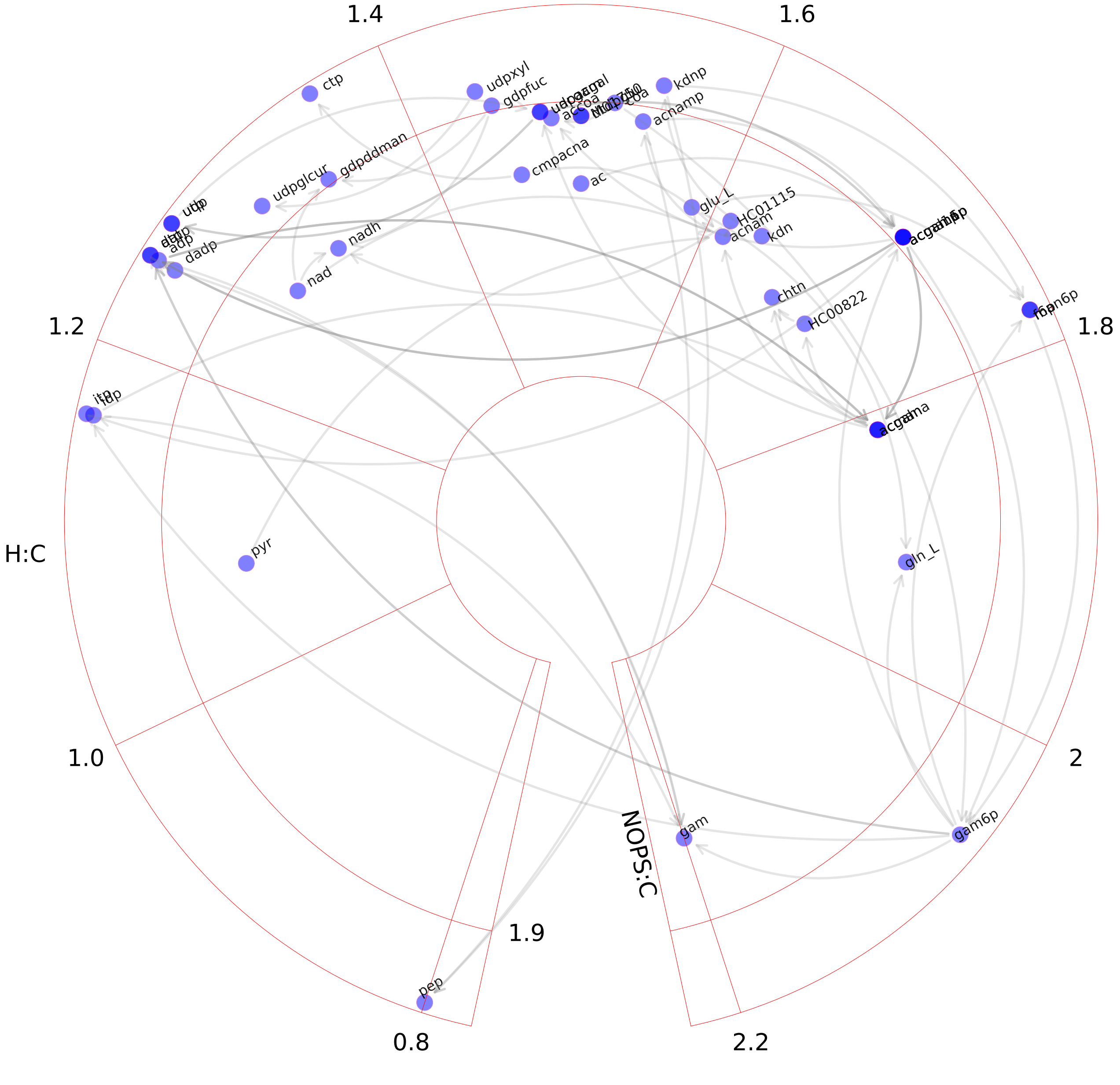

### Androgen and estrogen synthesis and metabolism.pdf

#### Androgen and estrogen synthesis and metabolism

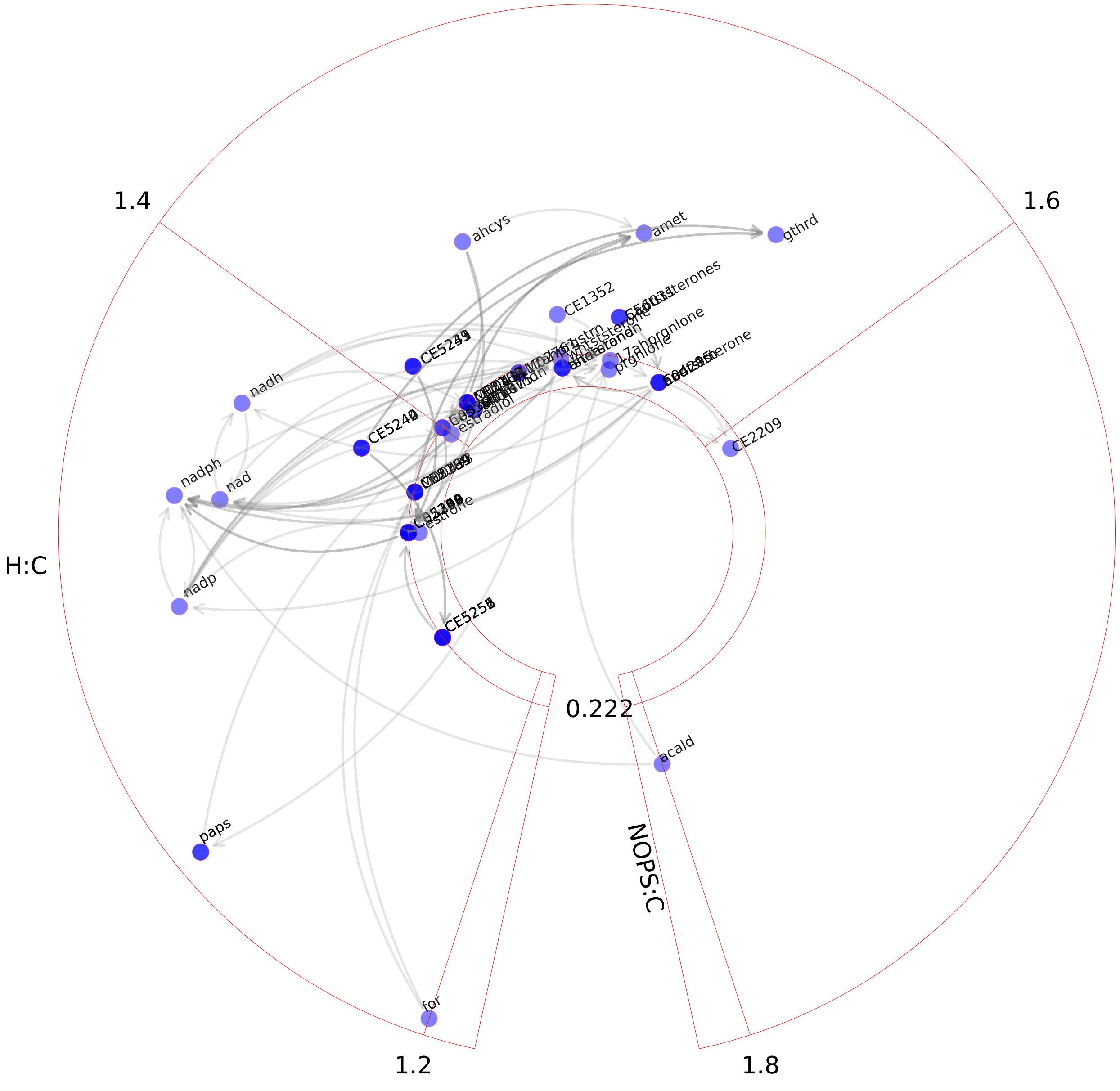

### Arachidonic acid metabolism.pdf

Arachidonic acid metabolism

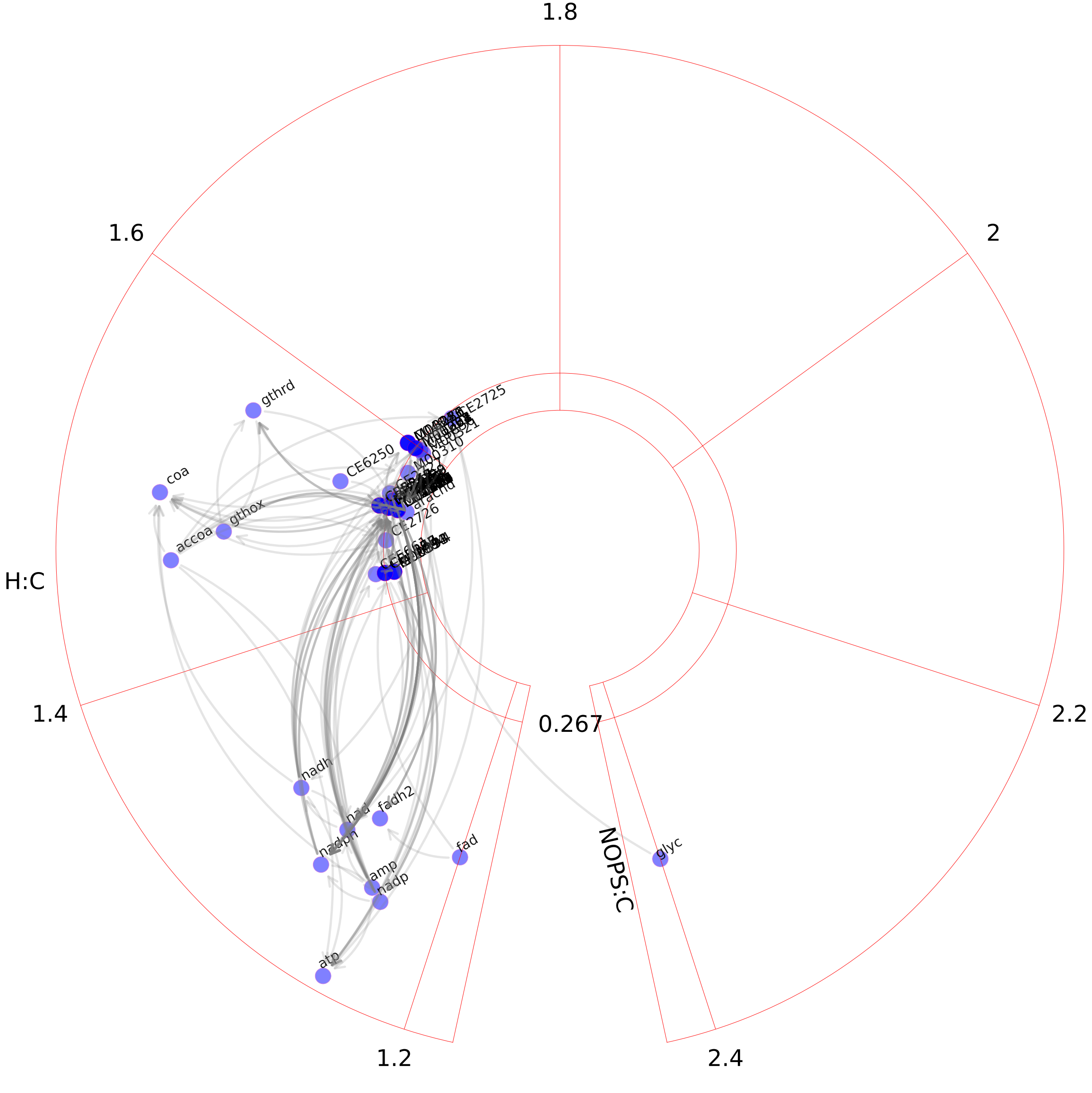

### Arginine and proline metabolism.pdf

#### Arginine and proline metabolism

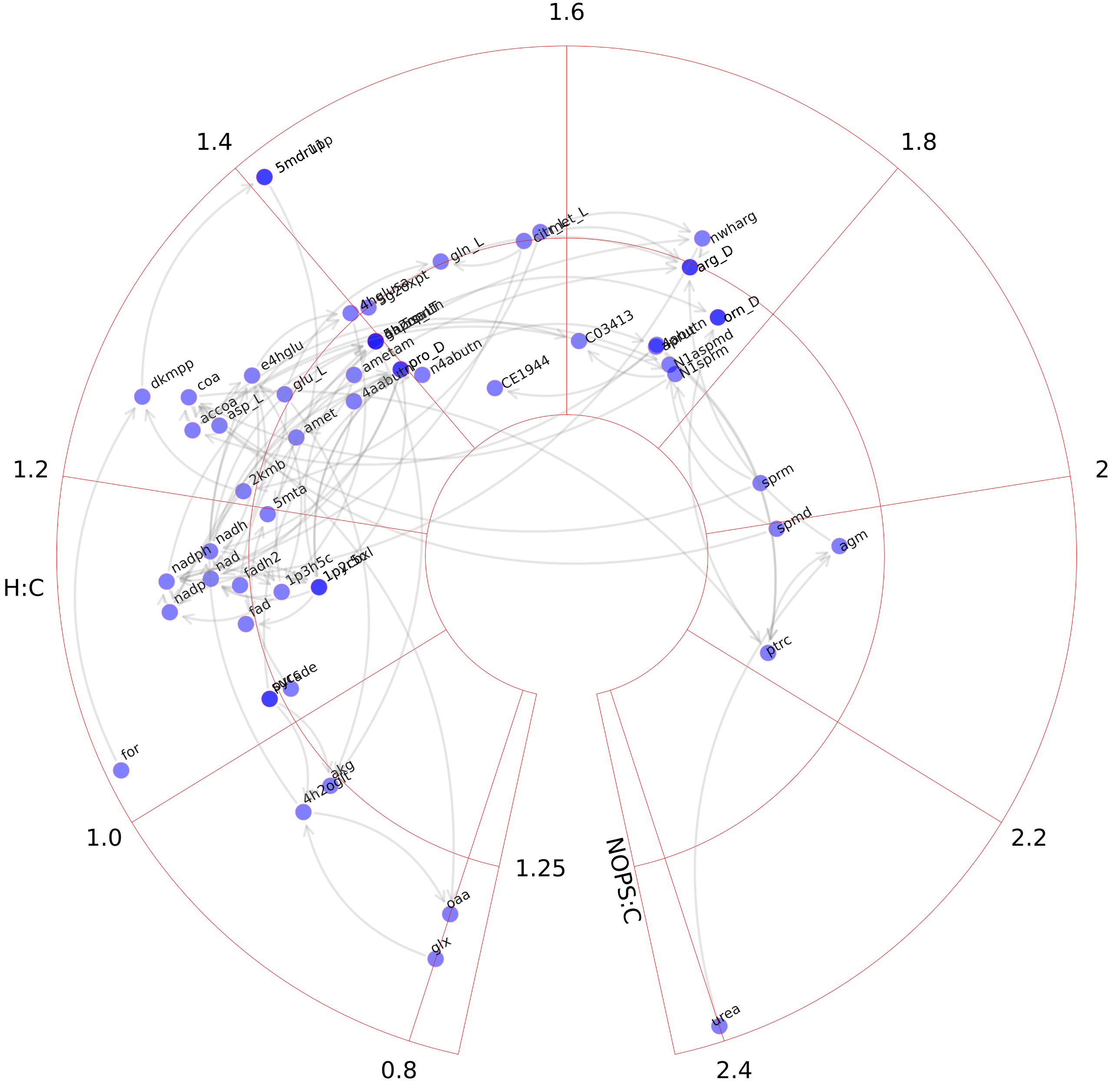

### Beta-Alanine metabolism.pdf

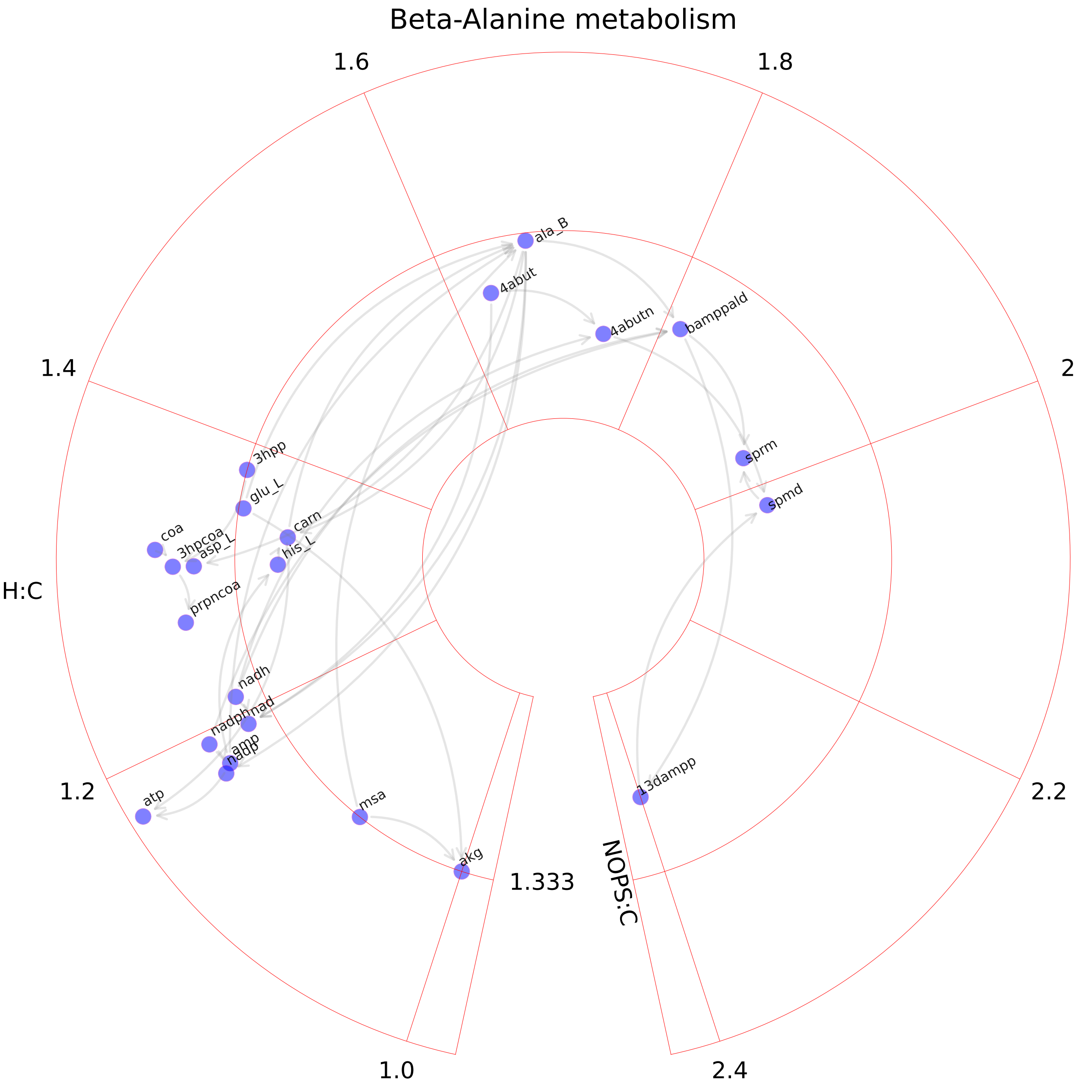

### Bile acid synthesis.pdf

#### Bile acid synthesis

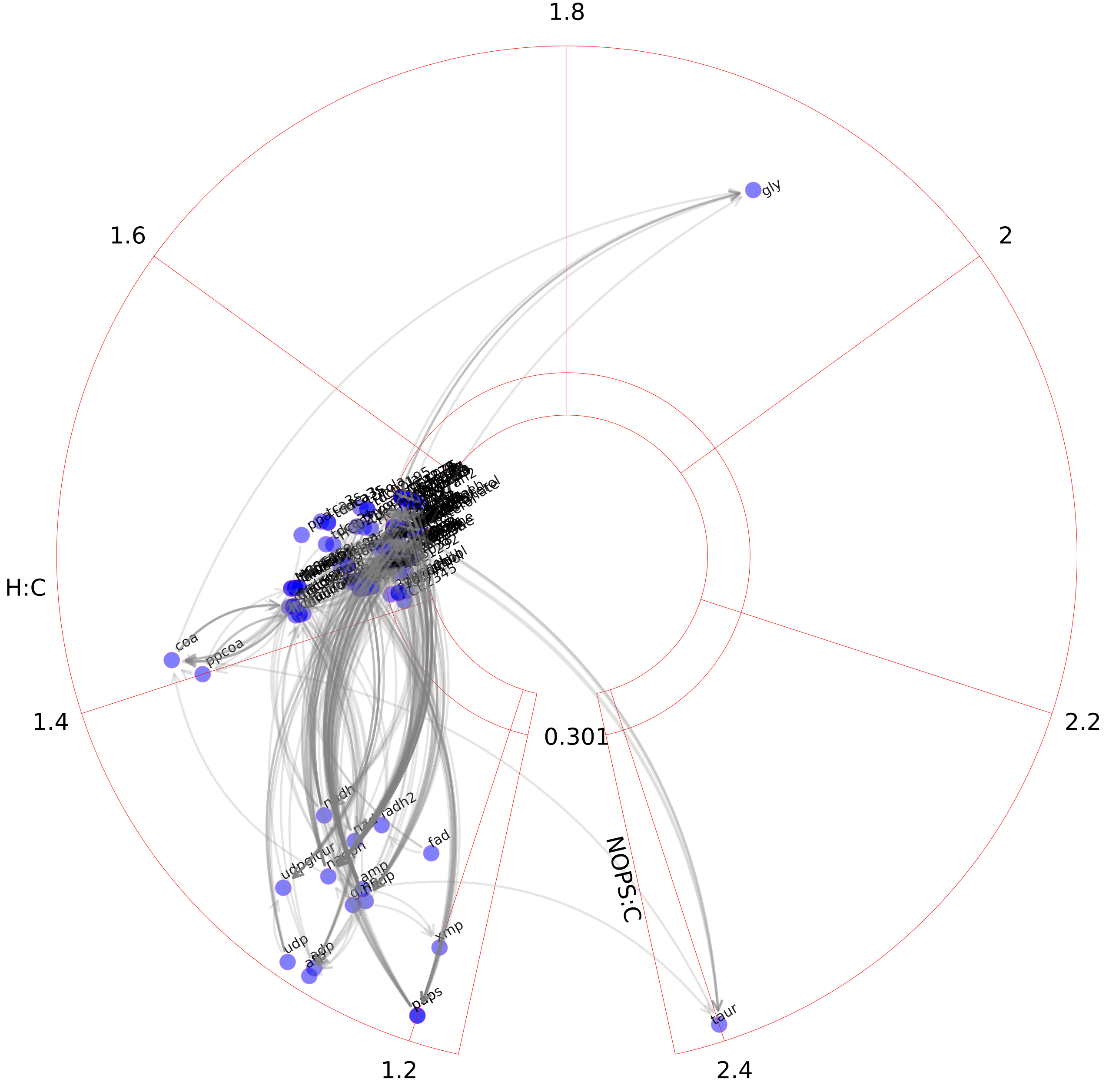

### Biotin metabolism.pdf

#### Biotin metabolism

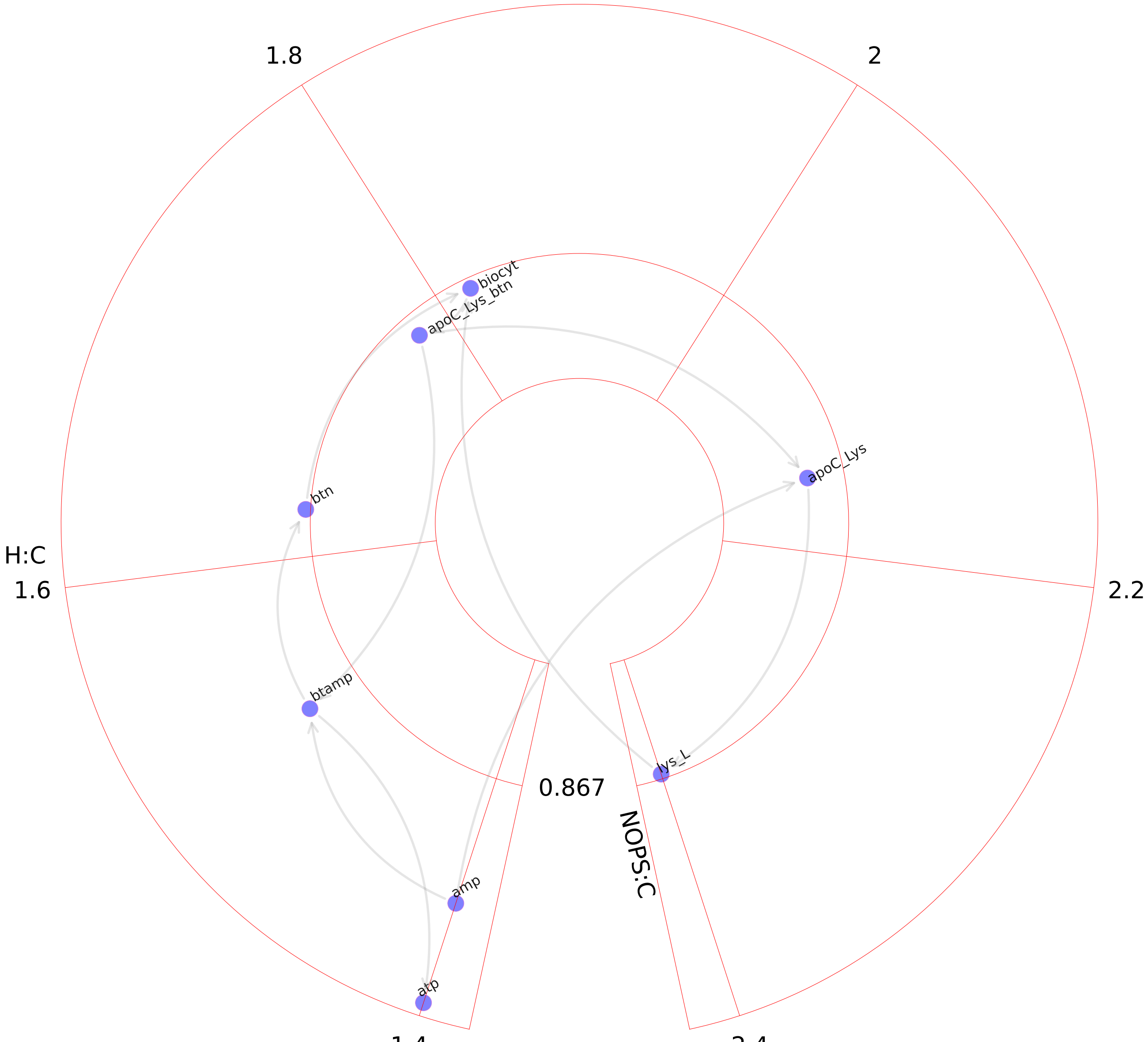

### Blood group synthesis.pdf

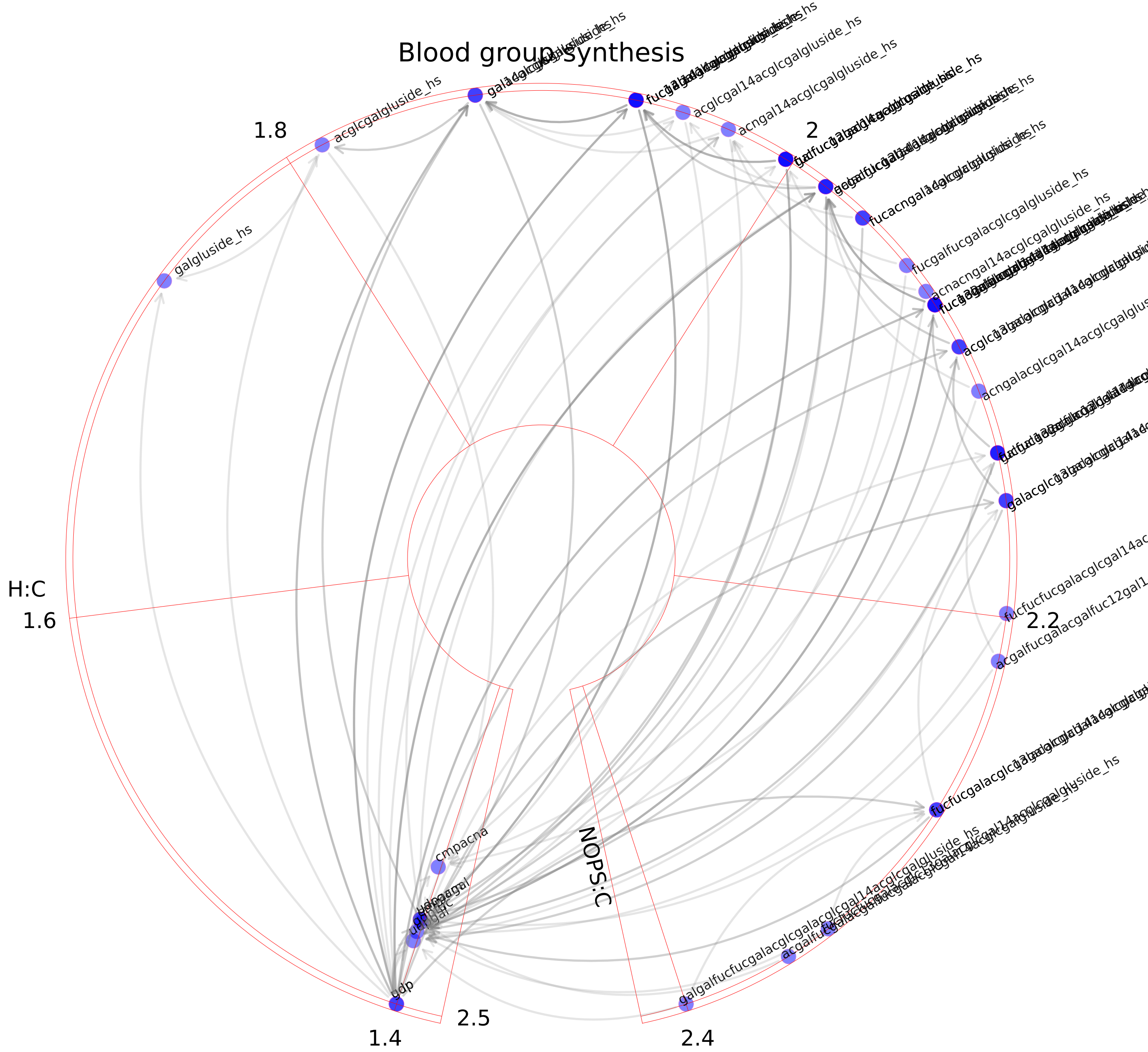

### Butanoate metabolism.pdf

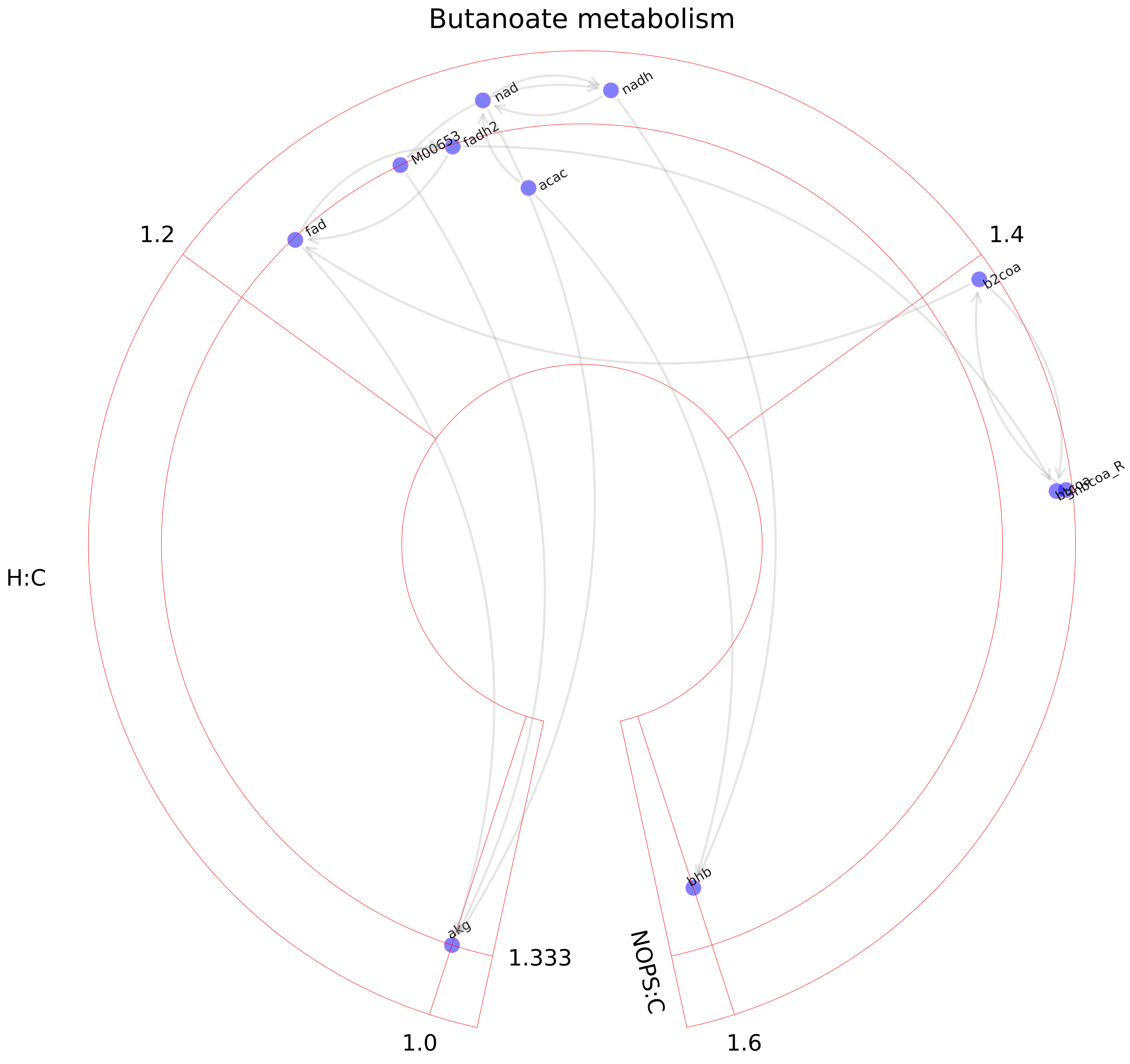

### C5-branched dibasic acid metabolism.pdf

#### C5-branched dibasic acid metabolism

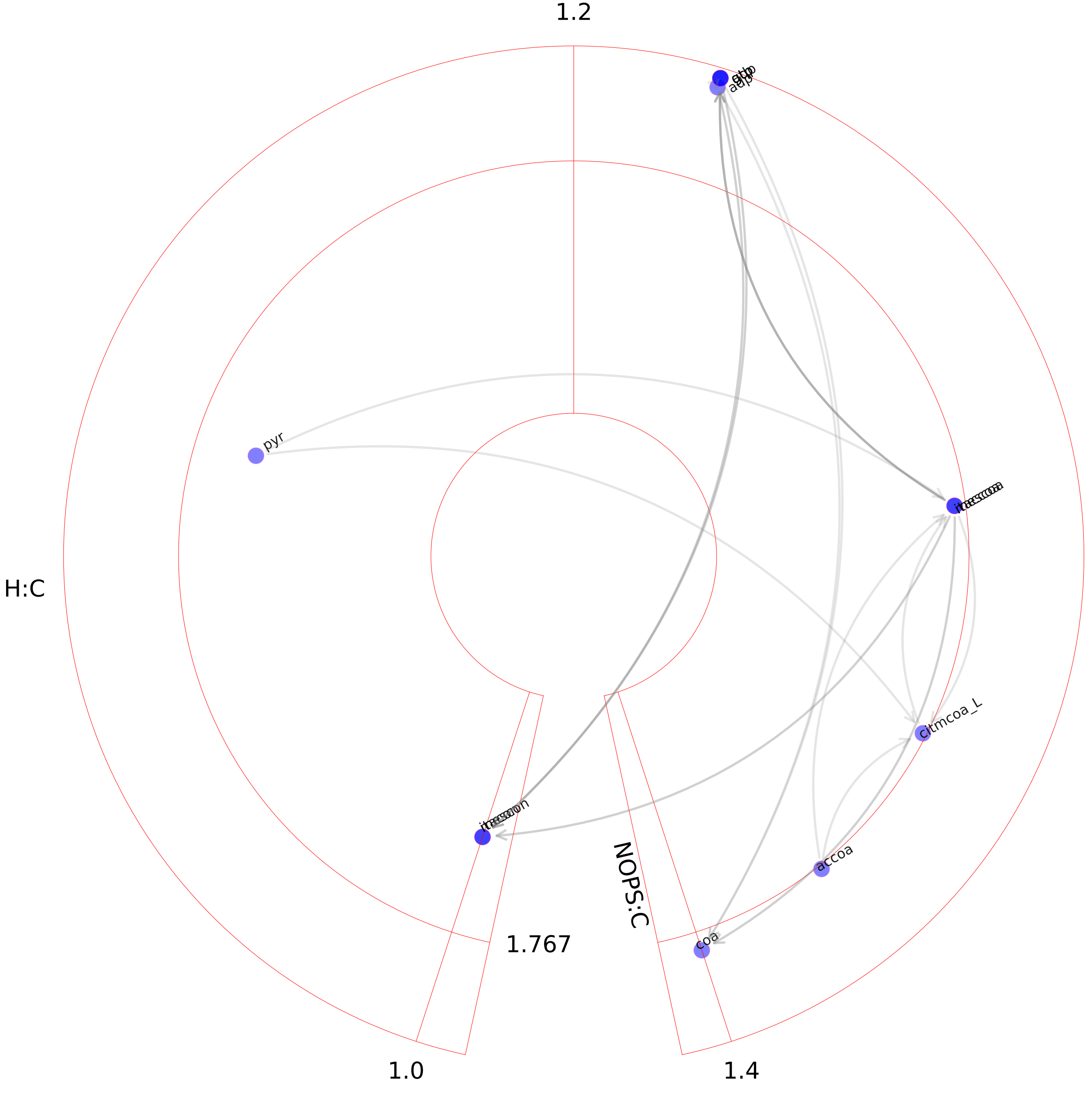

### Cholesterol metabolism.pdf

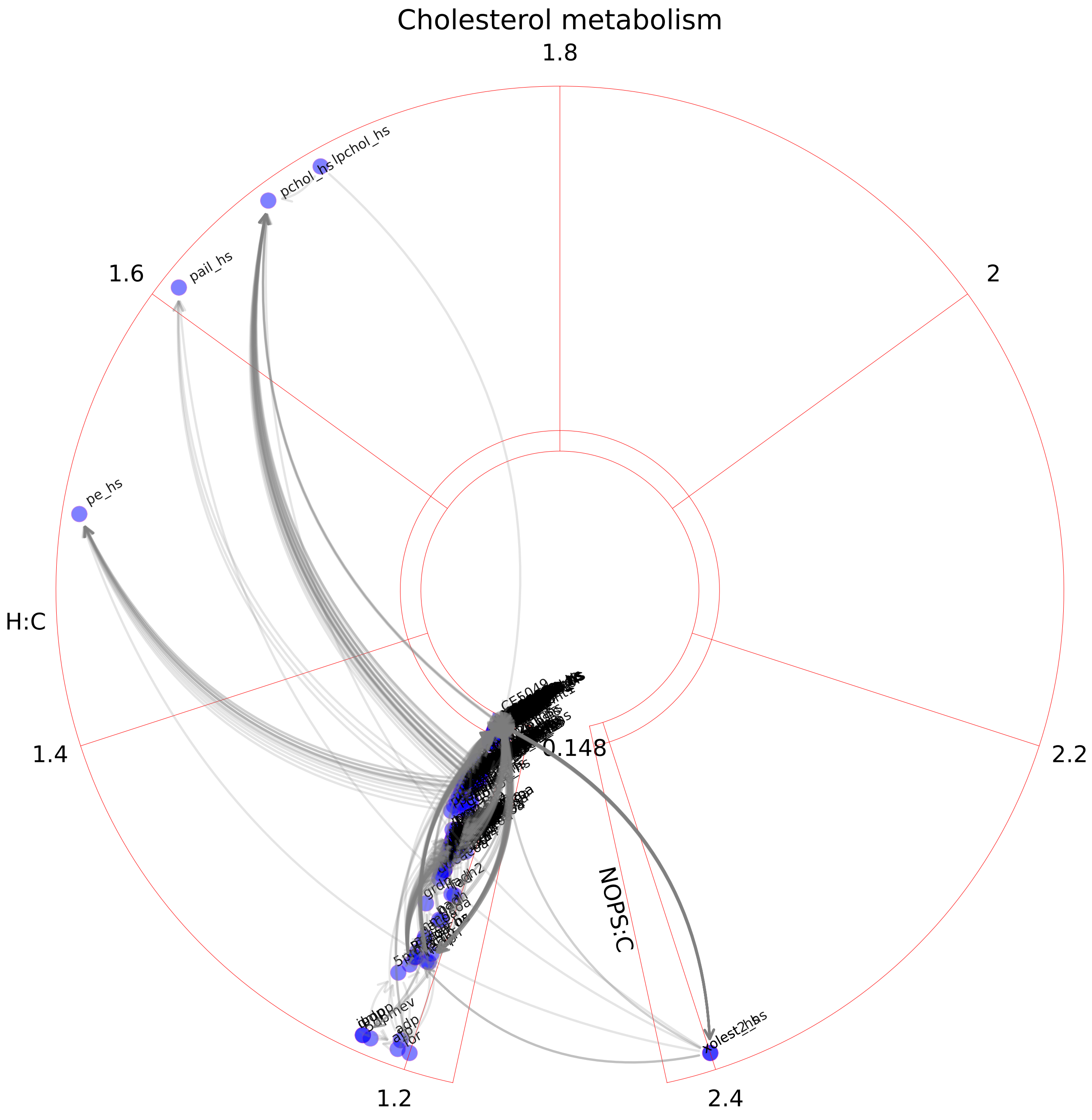

### Chondroitin sulfate degradation.pdf

#### Chondroitin sulfate degradation

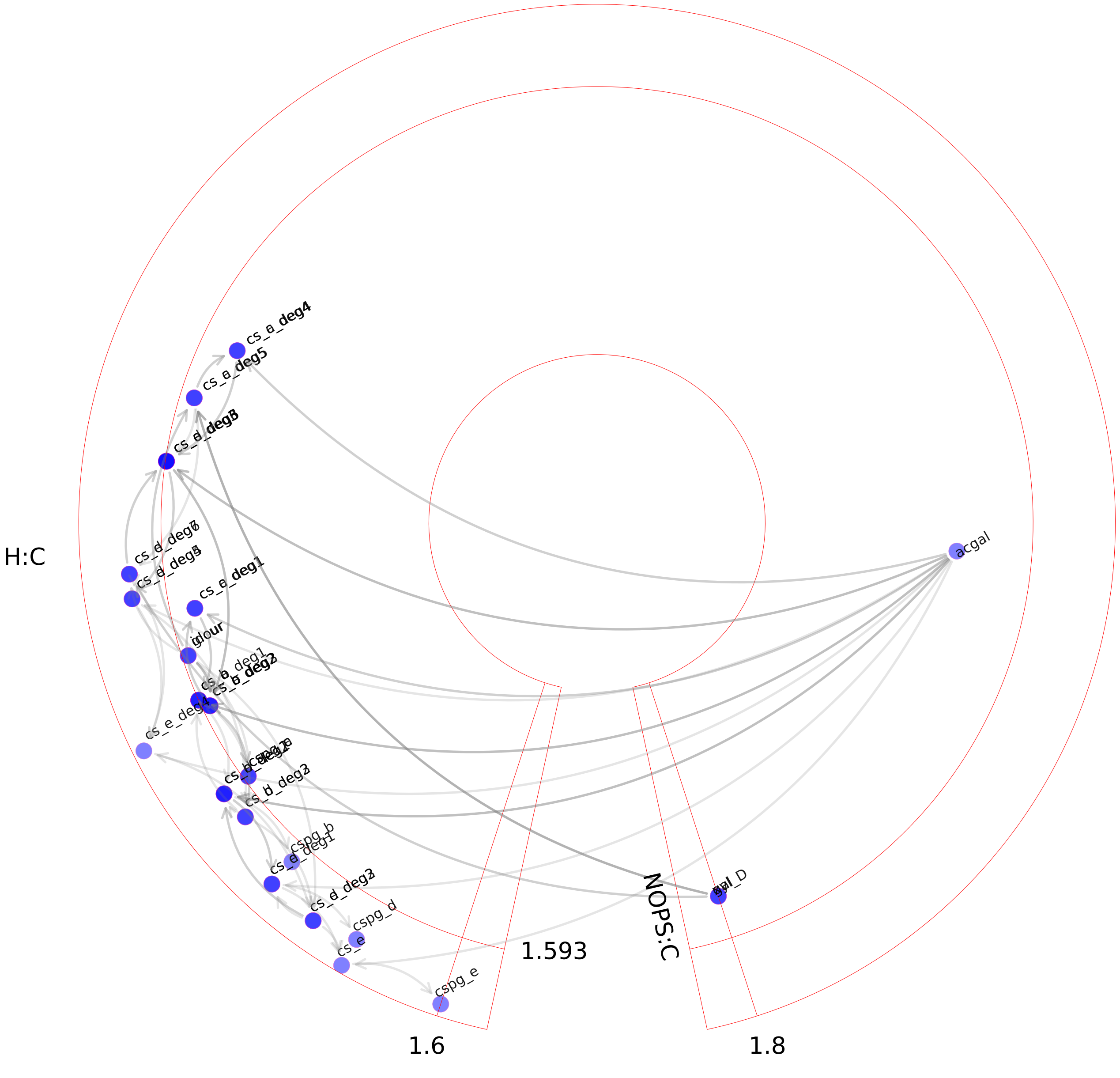

### Chondroitin synthesis.pdf

#### Chondroitin synthesis

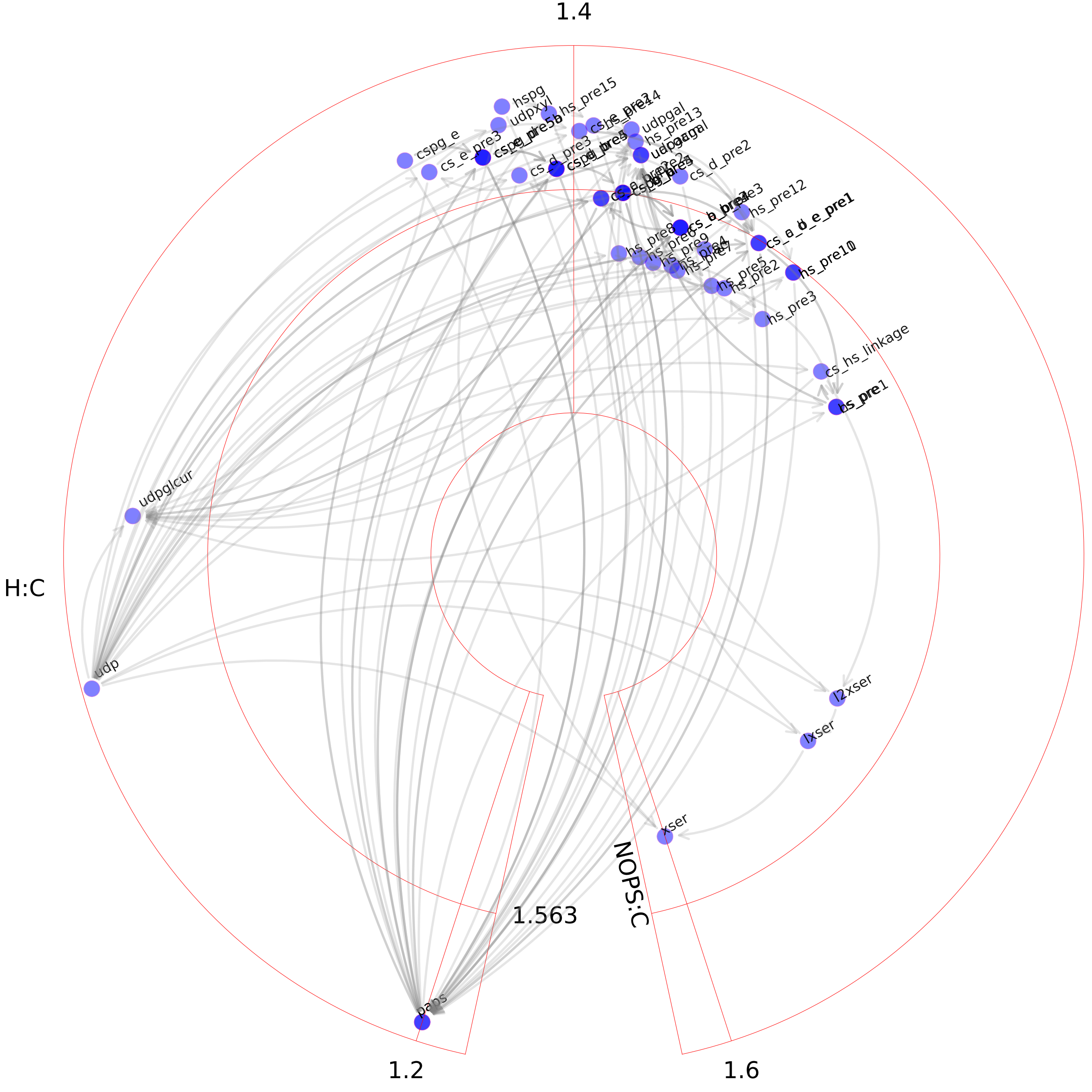

### Citric acid cycle.pdf

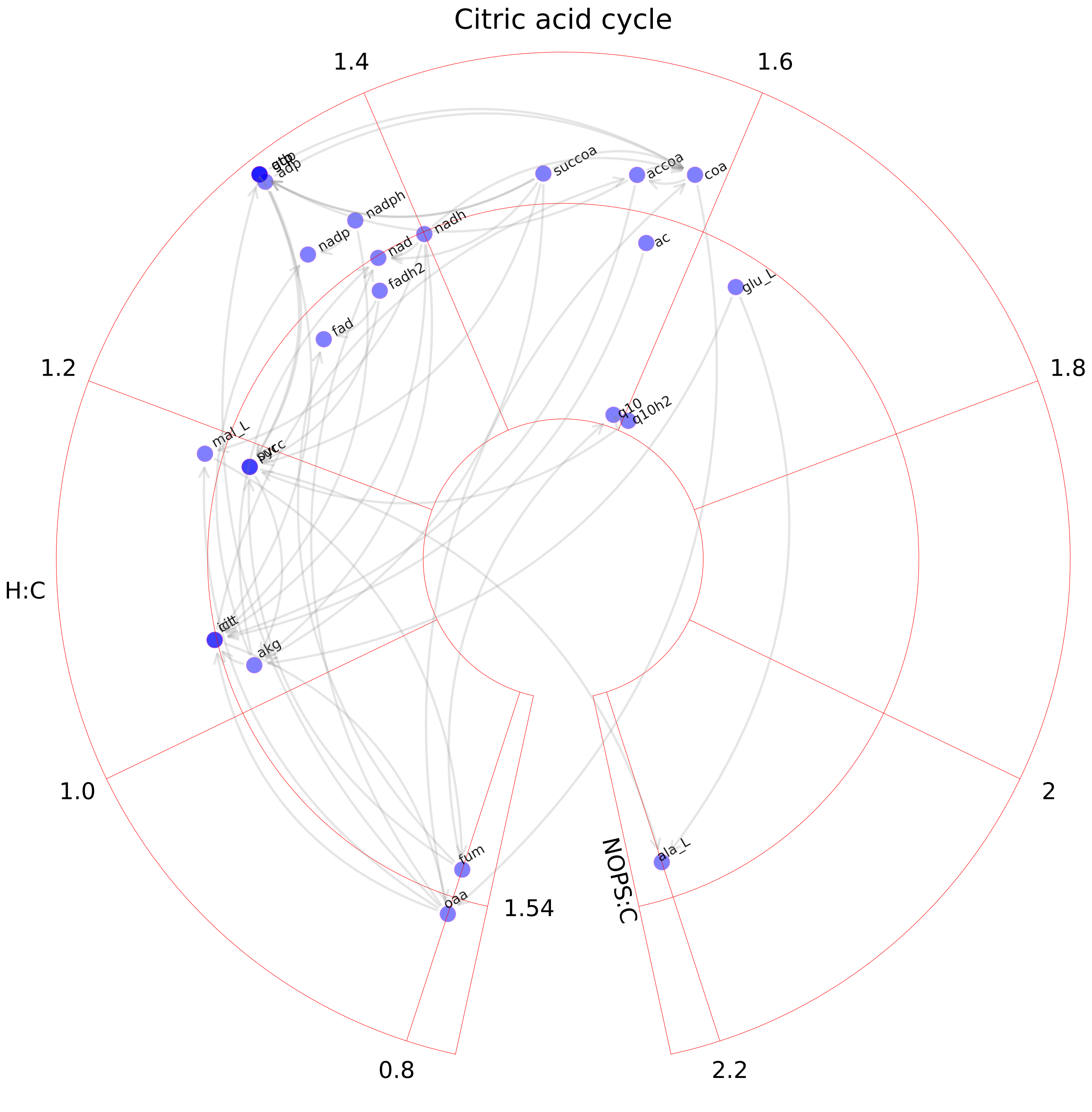

### CoA catabolism.pdf

#### CoA catabolism

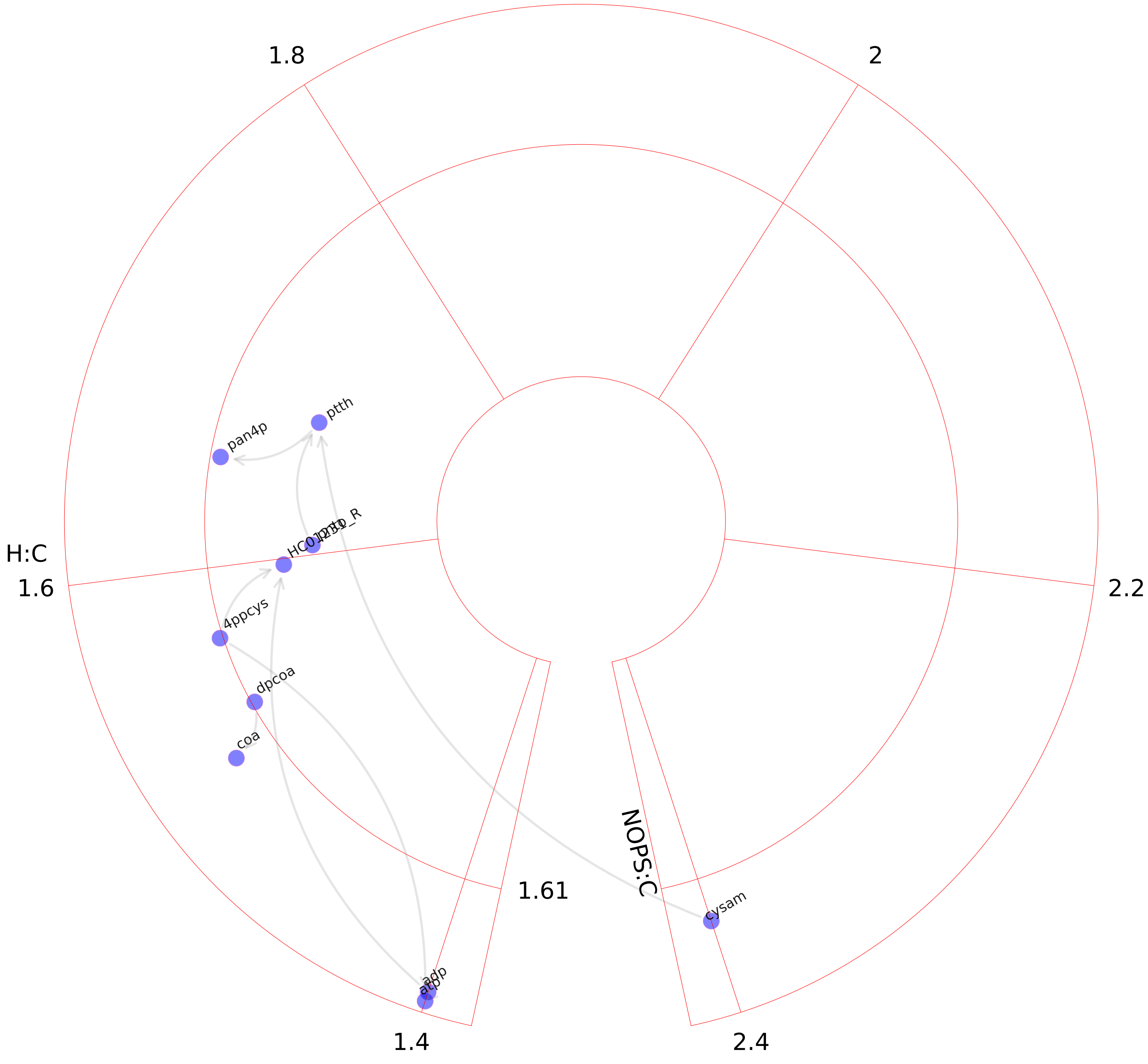

### CoA synthesis.pdf

#### CoA synthesis

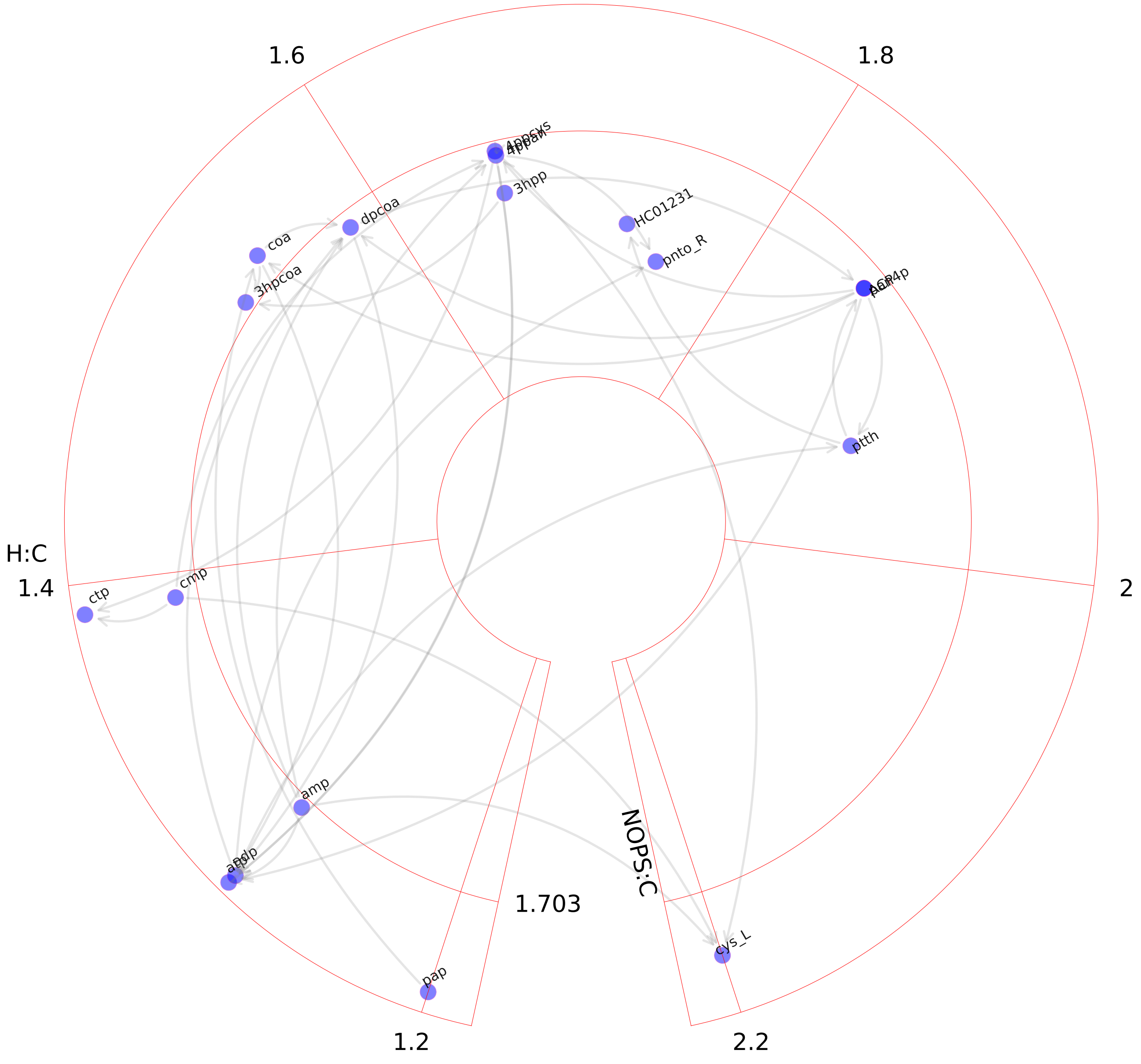

### Cytochrome metabolism.pdf

Cytochrome metabolism

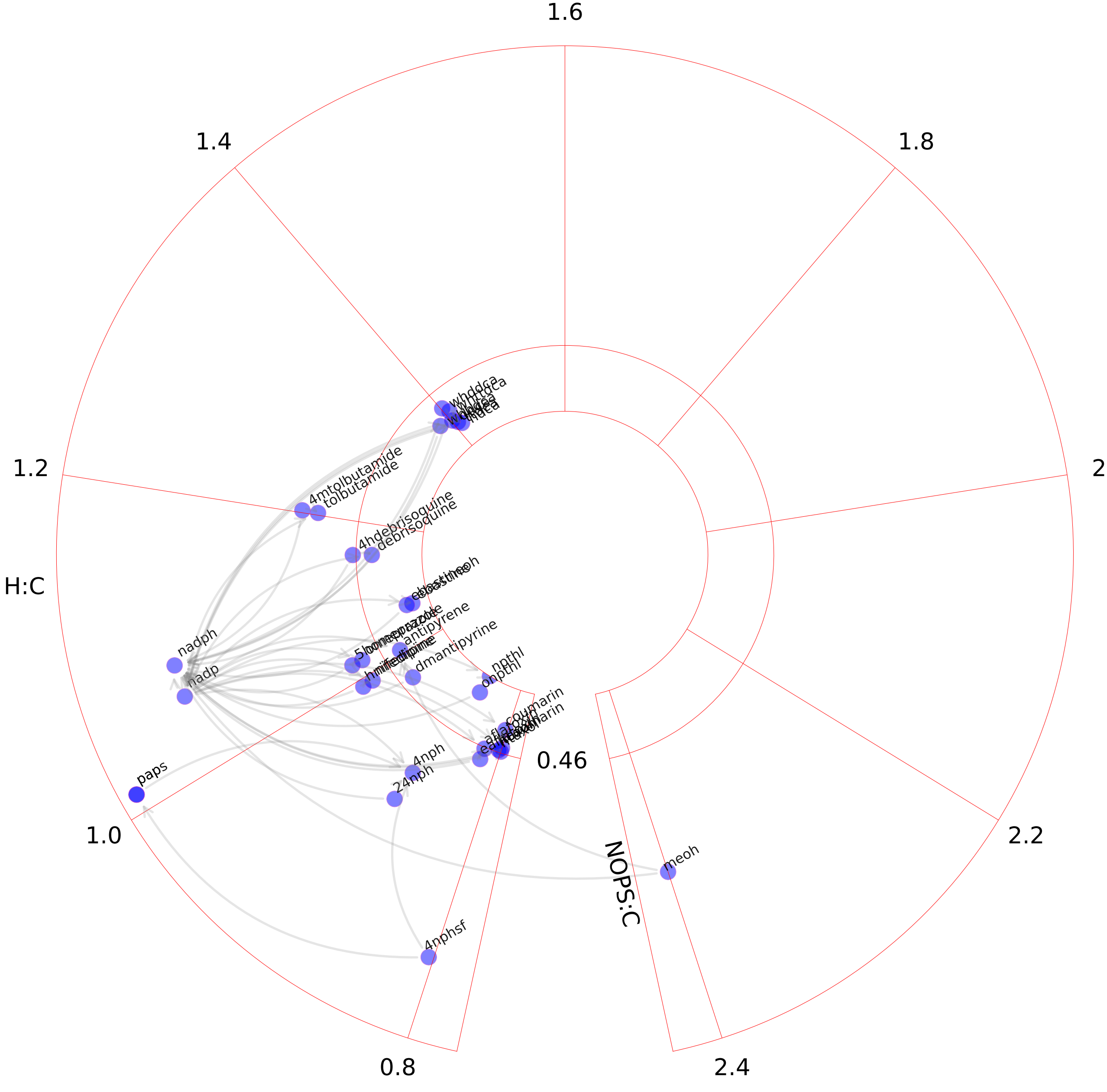

### D-alanine metabolism.pdf

#### D-alanine metabolism

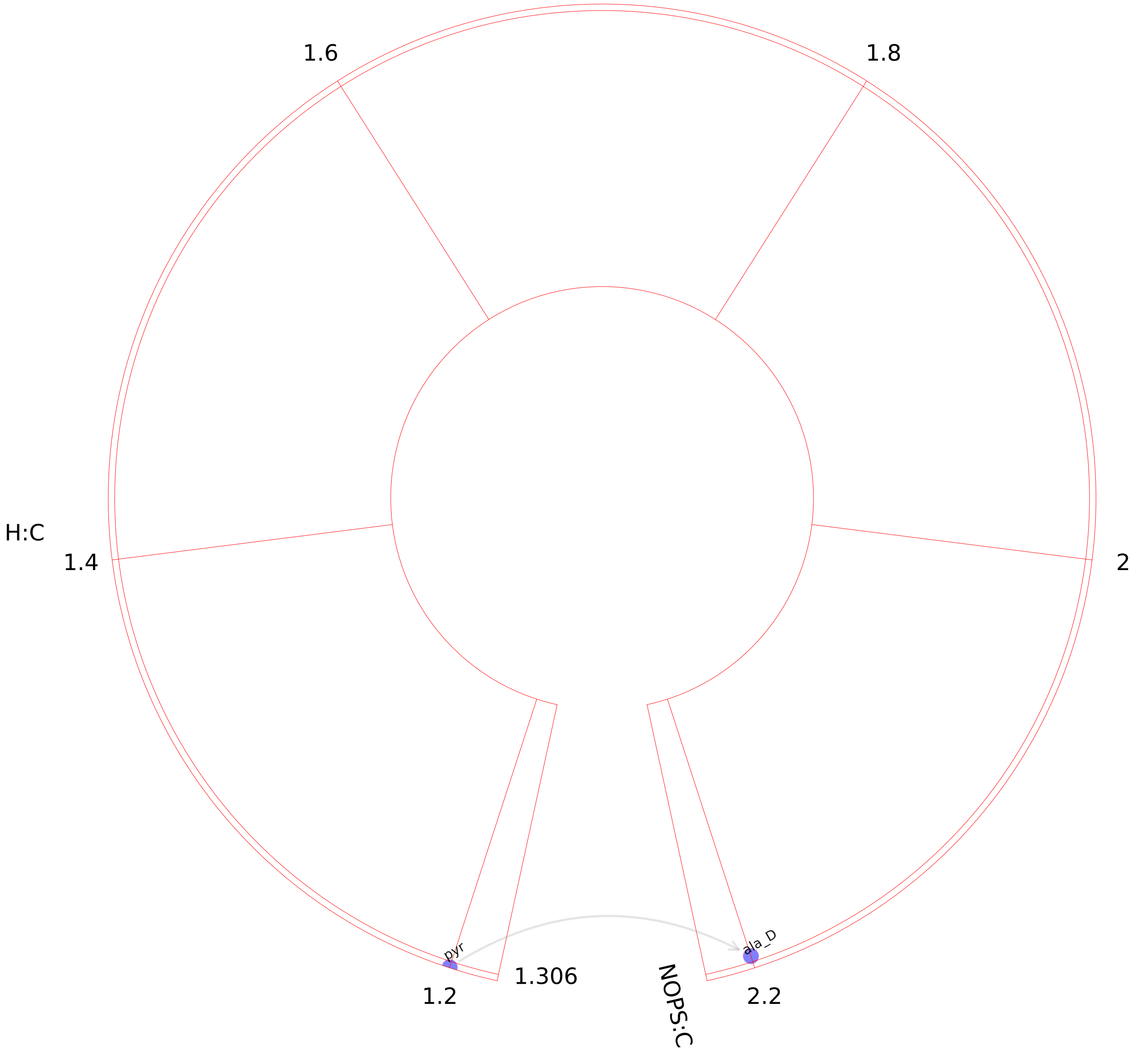

### Dietary fiber binding.pdf

#### Dietary fiber binding

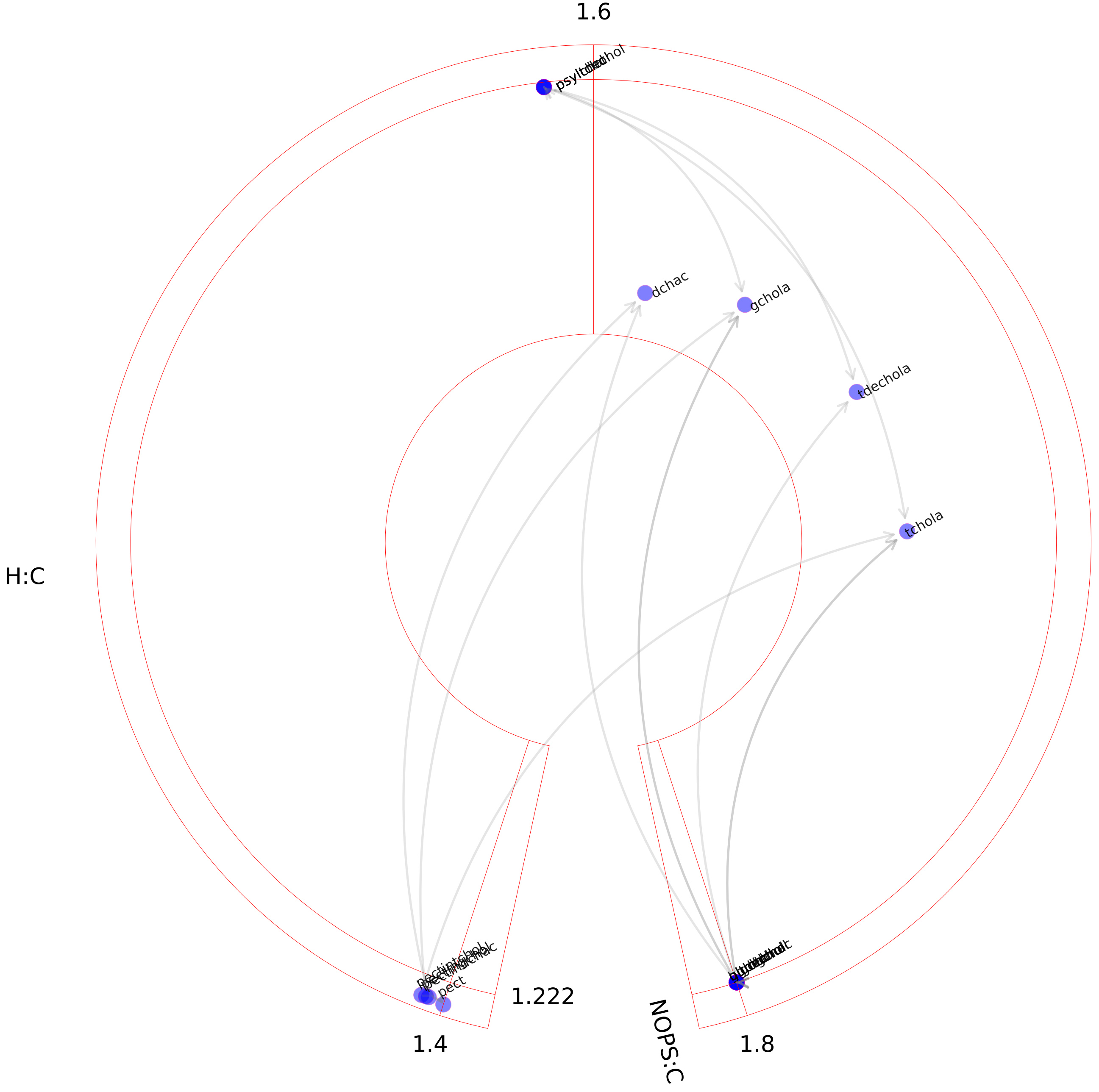

### Drug metabolism.pdf

#### Drug metabolism

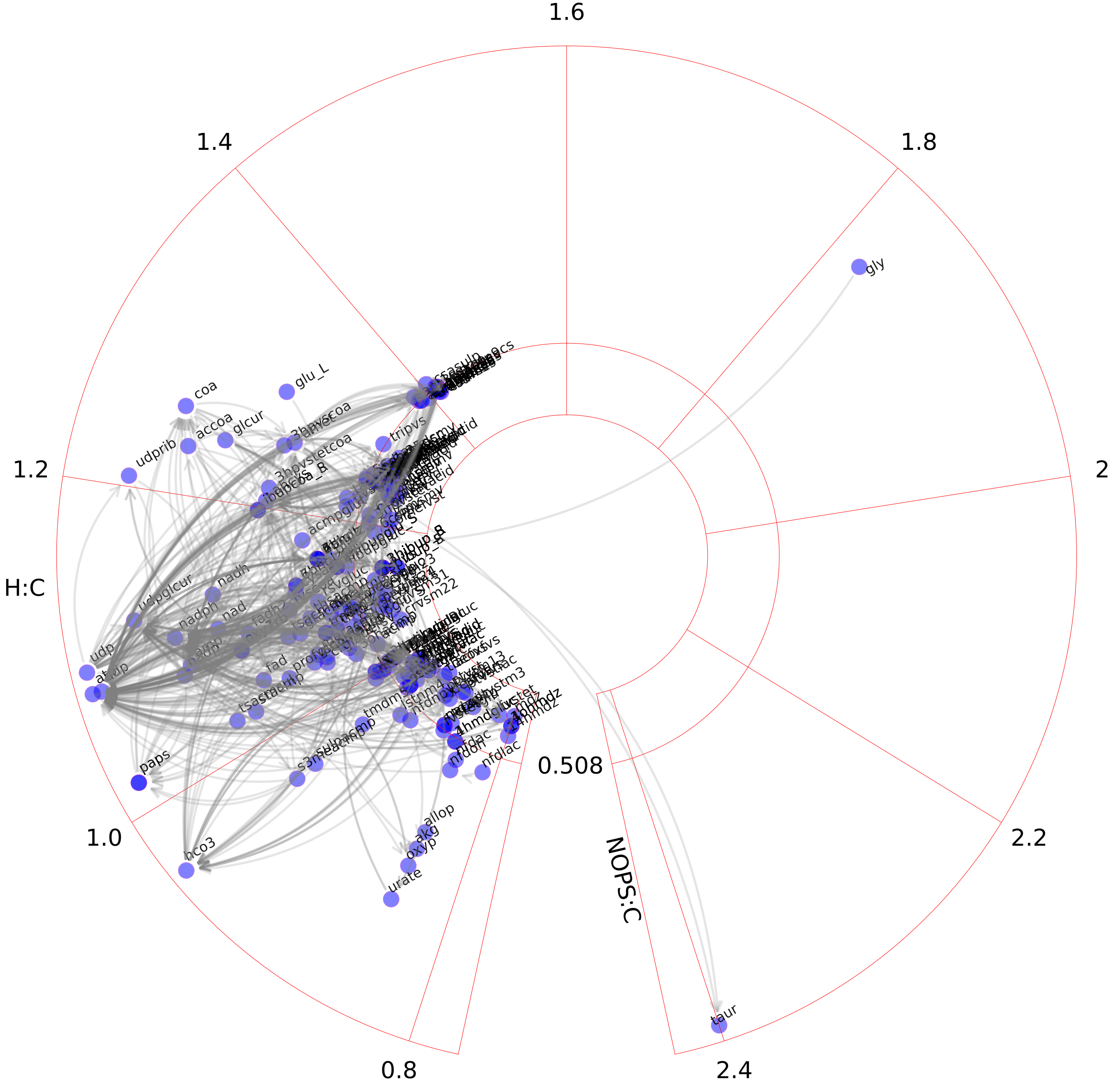

### Eicosanoid metabolism.pdf

Eicosanoid metabolism

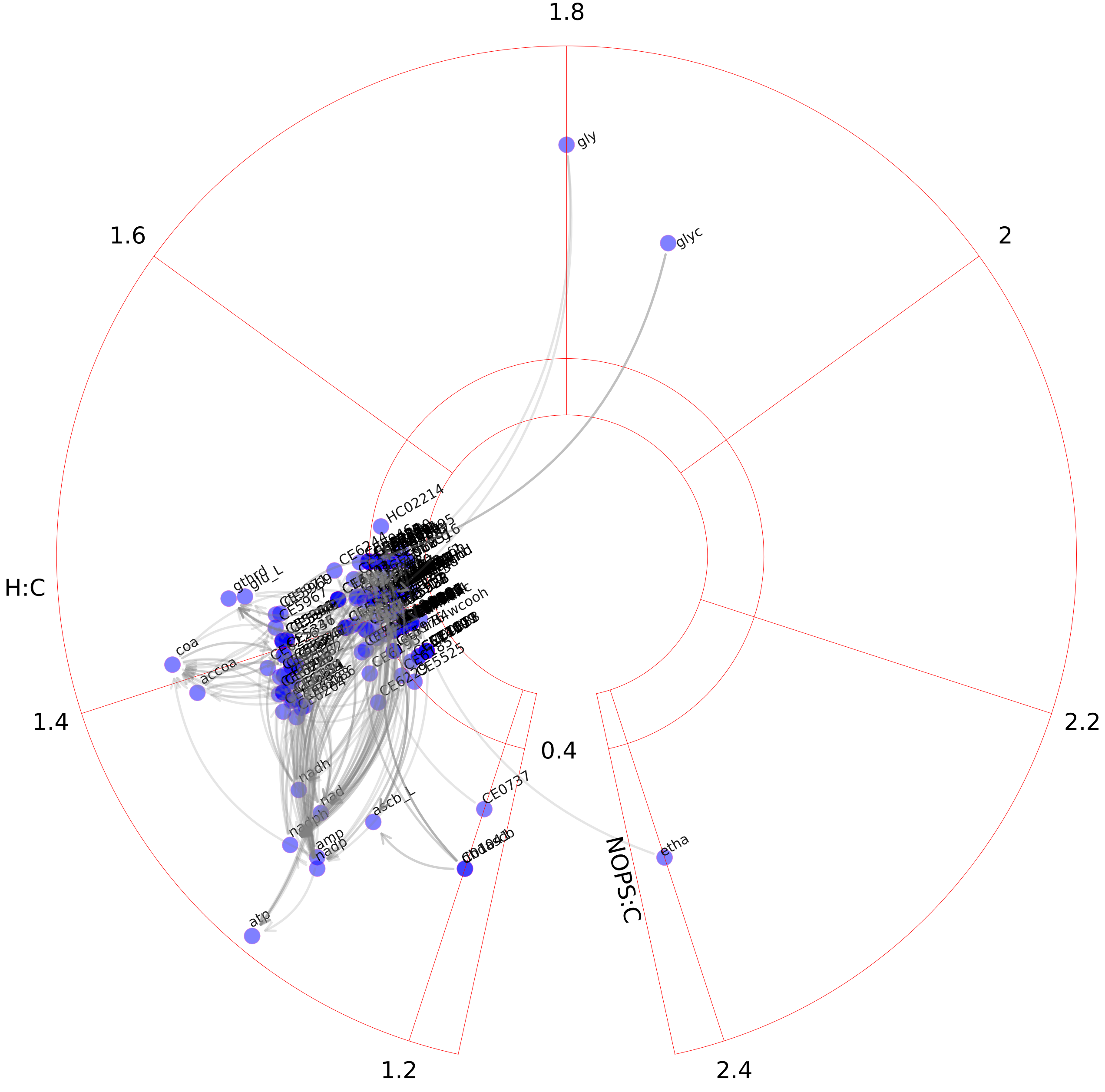

### Fatty acid oxidation.pdf

#### Fatty acid oxidation

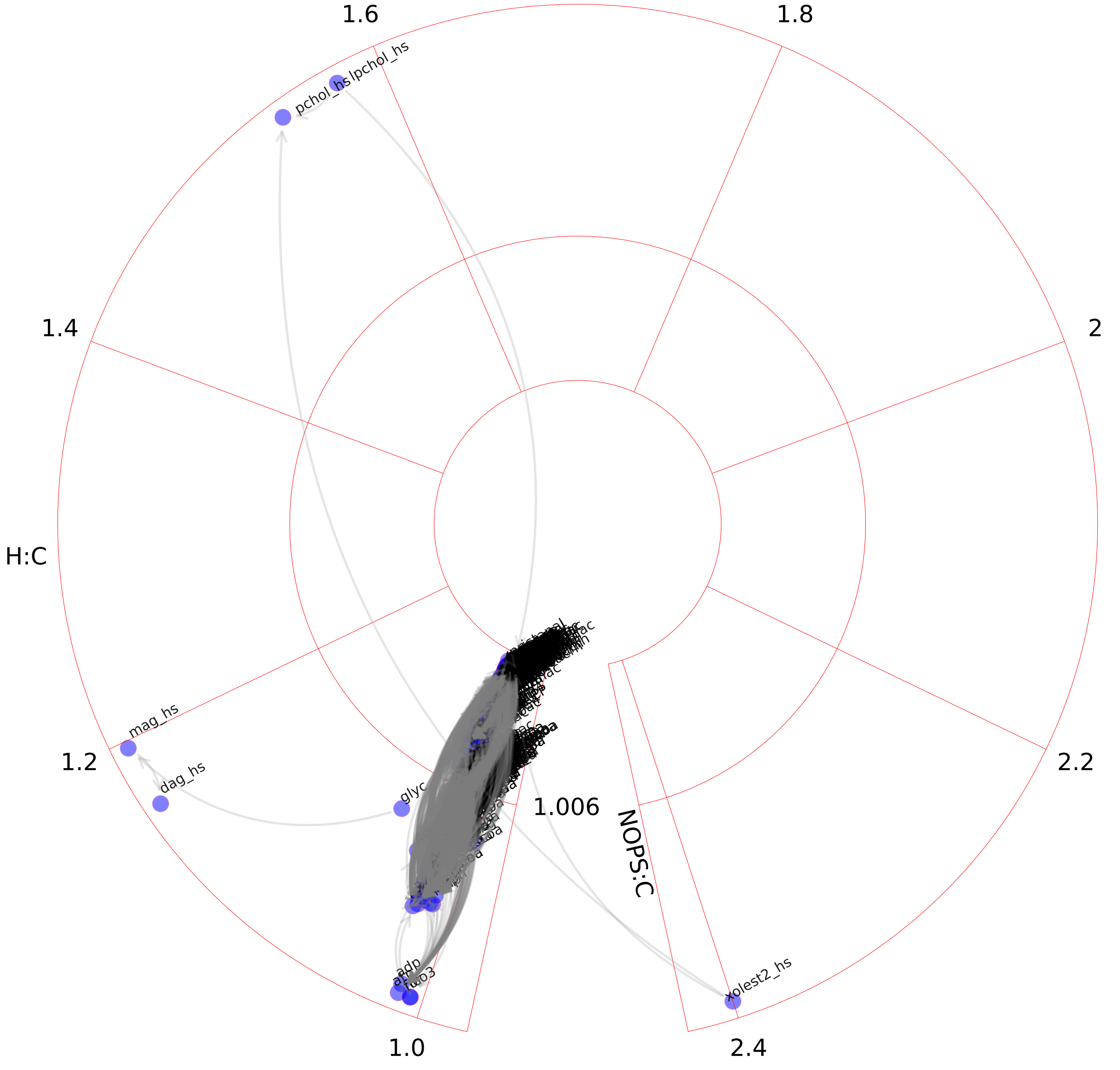

### Fatty acid synthesis.pdf

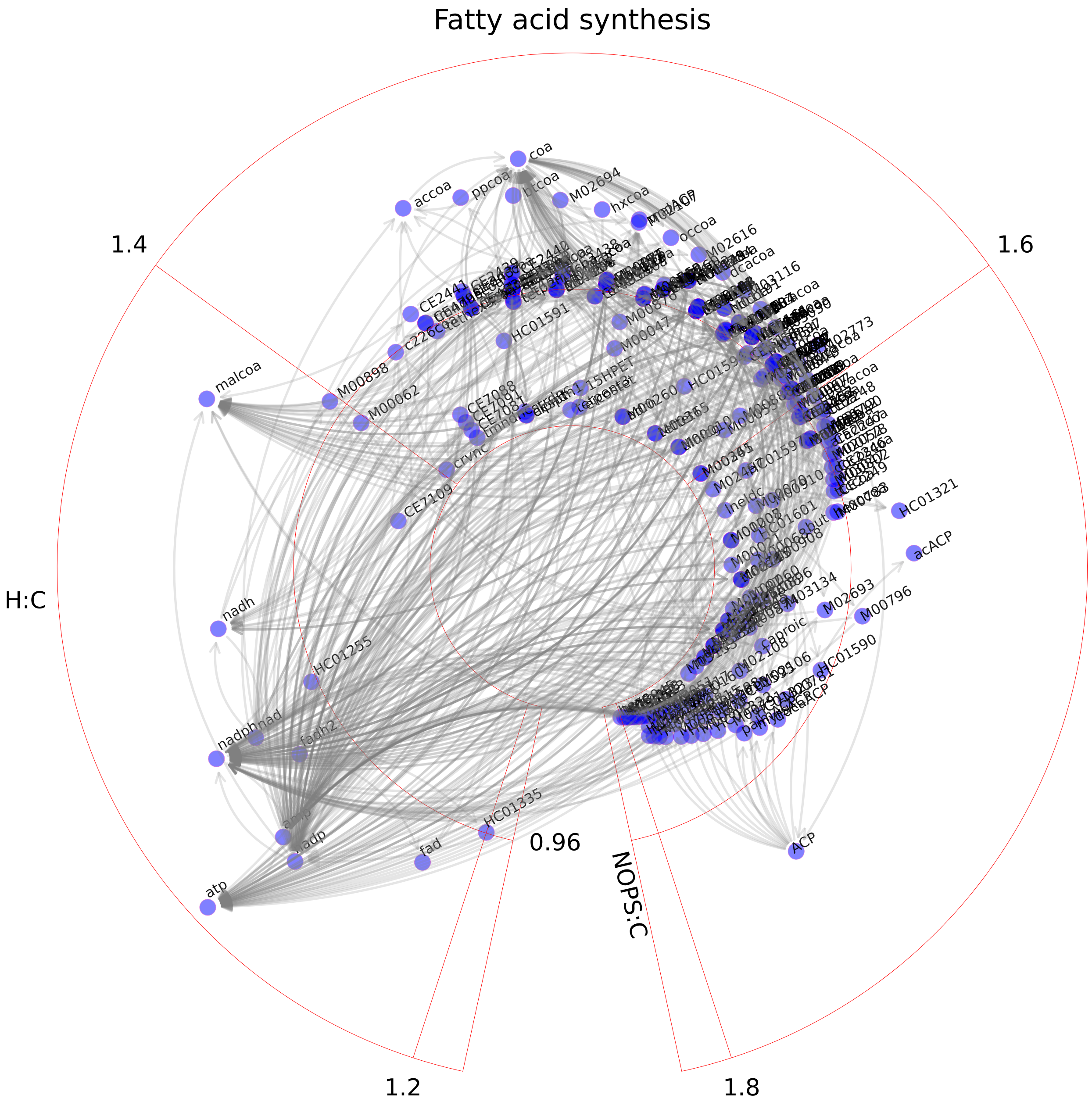

### Folate metabolism.pdf

#### Folate metabolism

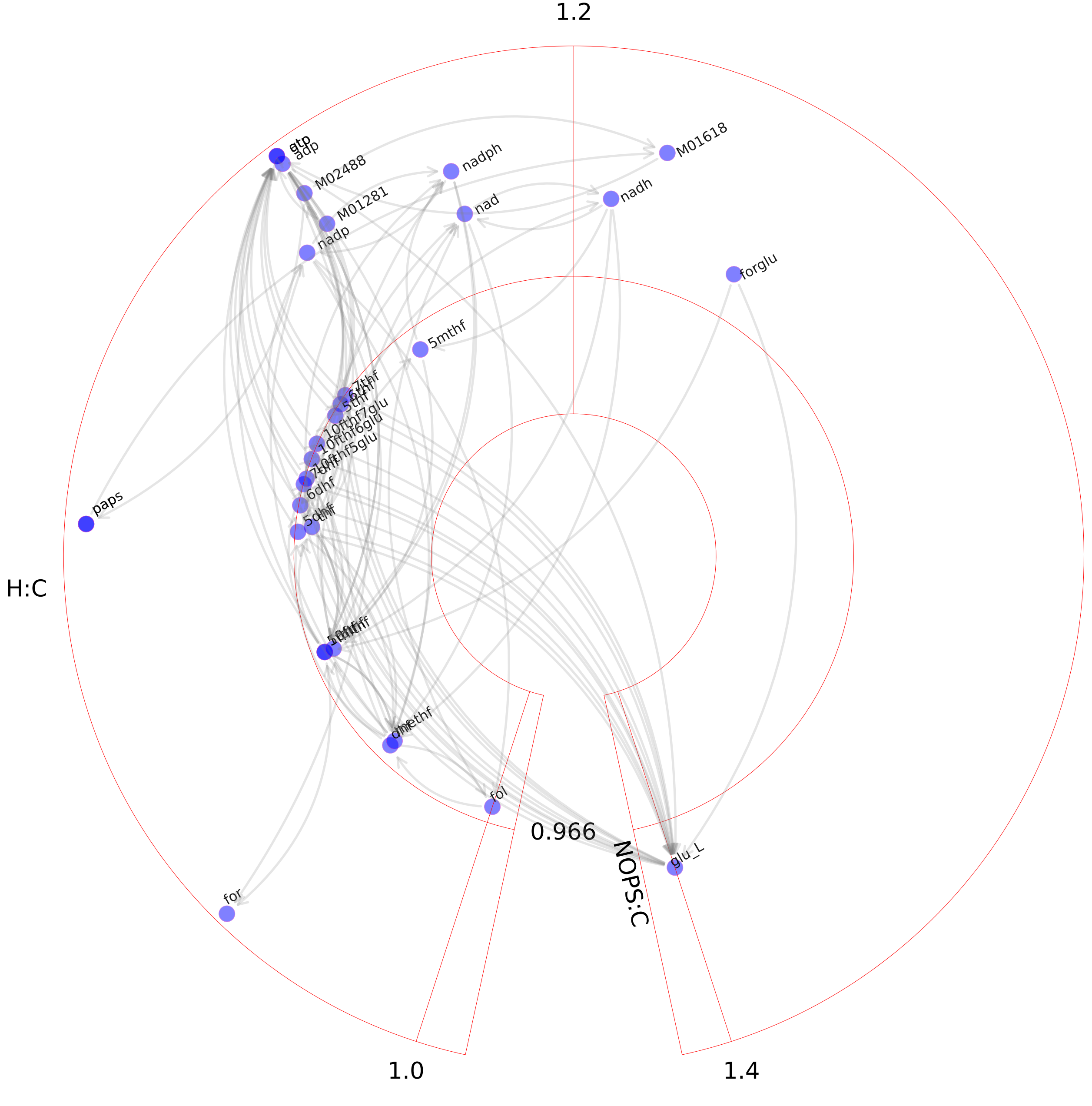

### Fructose and mannose metabolism.pdf

#### Fructose and mannose metabolism

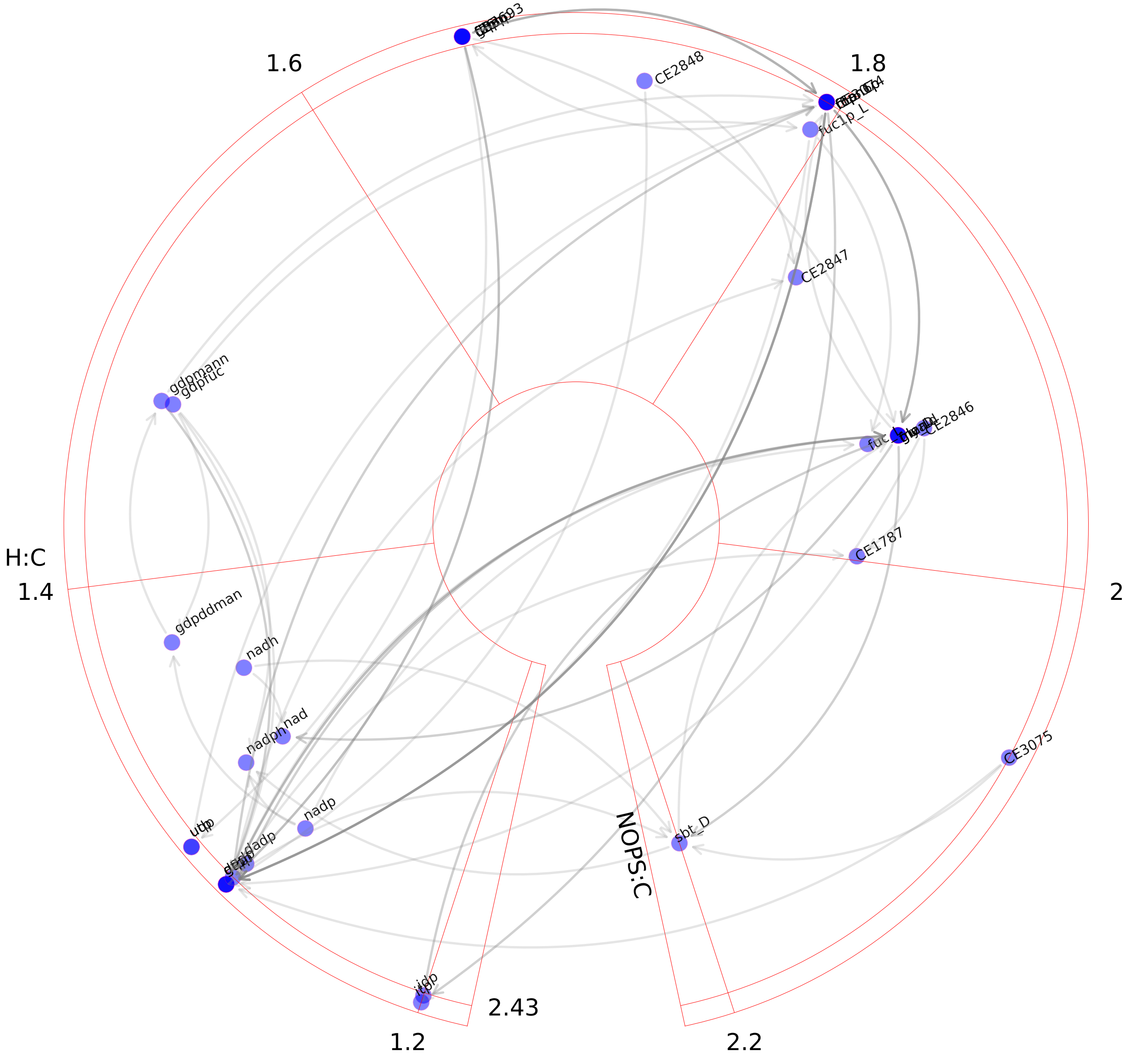

### Galactose metabolism.pdf

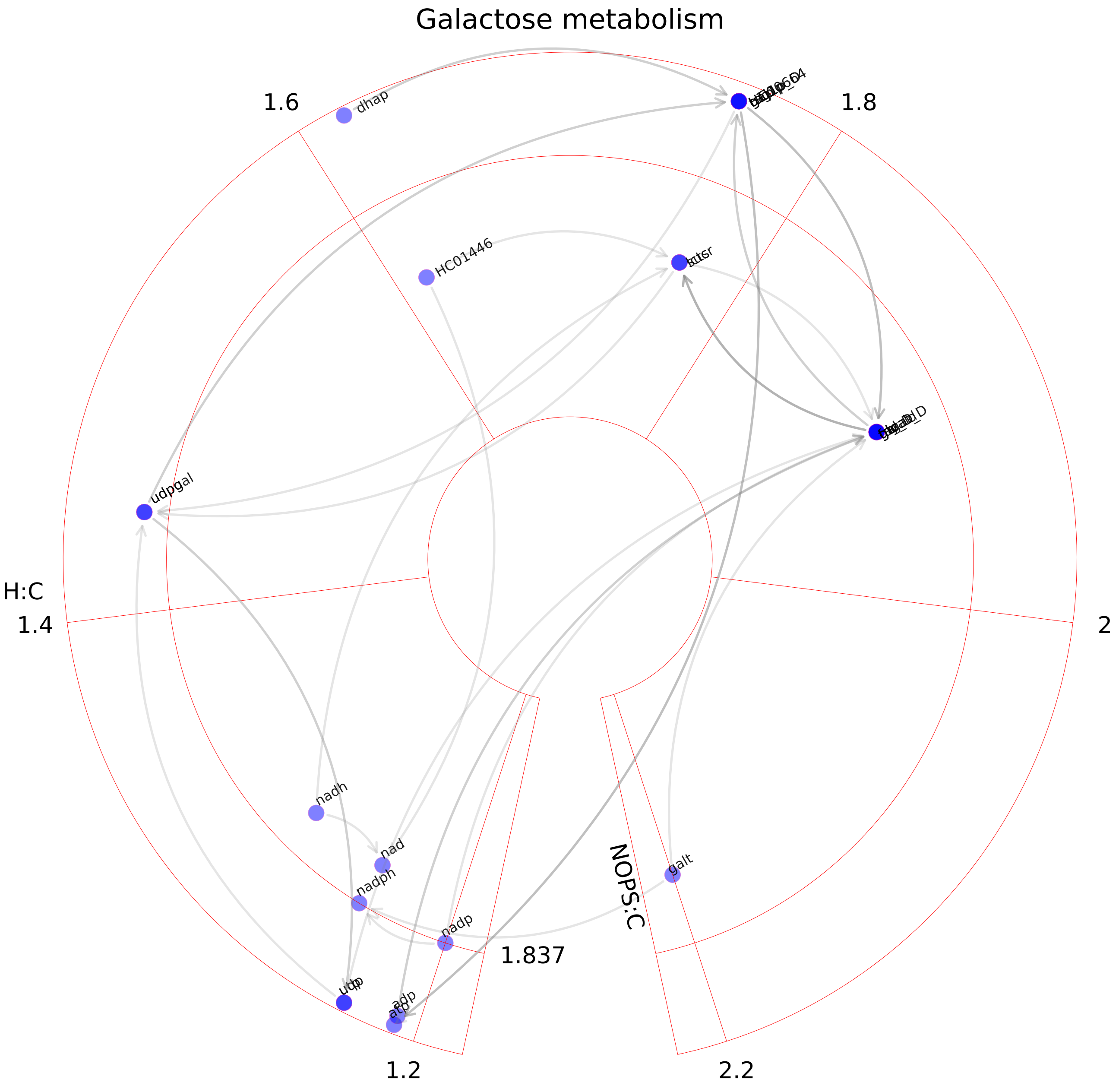

### Glutamate metabolism.pdf

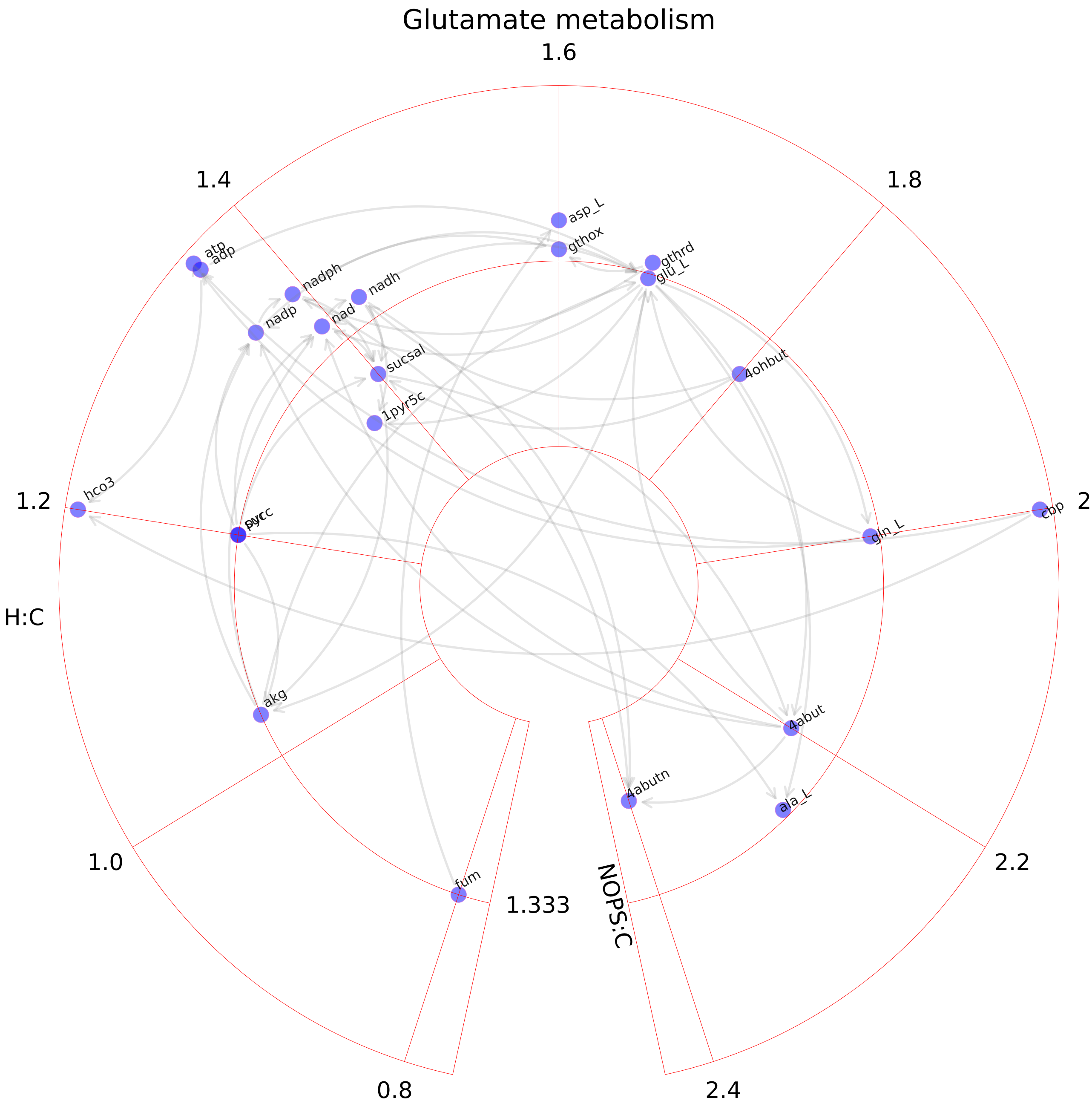

### Glutathione metabolism.pdf

#### Glutathione metabolism

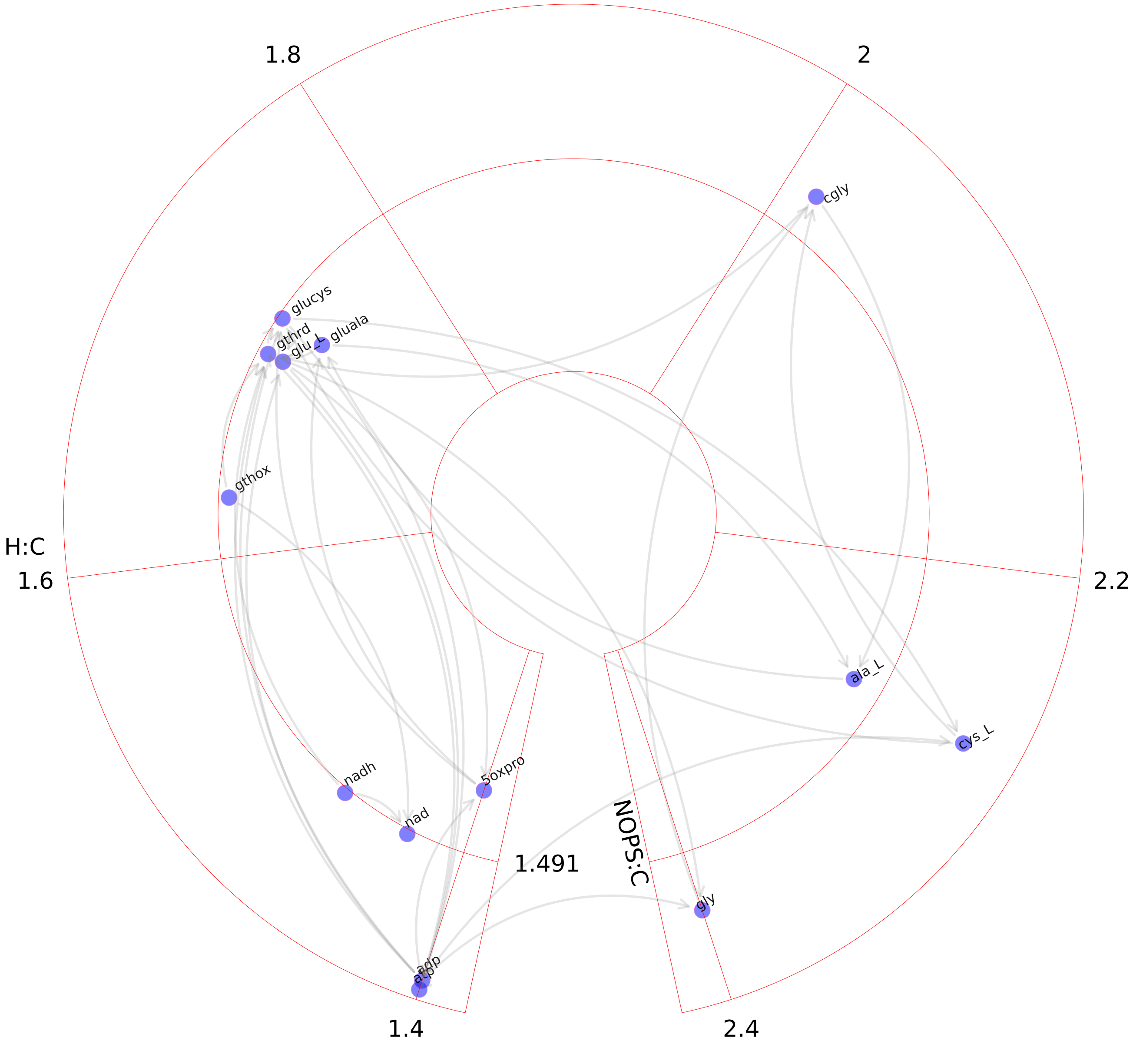

### Glycerophospholipid metabolism.pdf

#### Glycerophospholipid metabolism

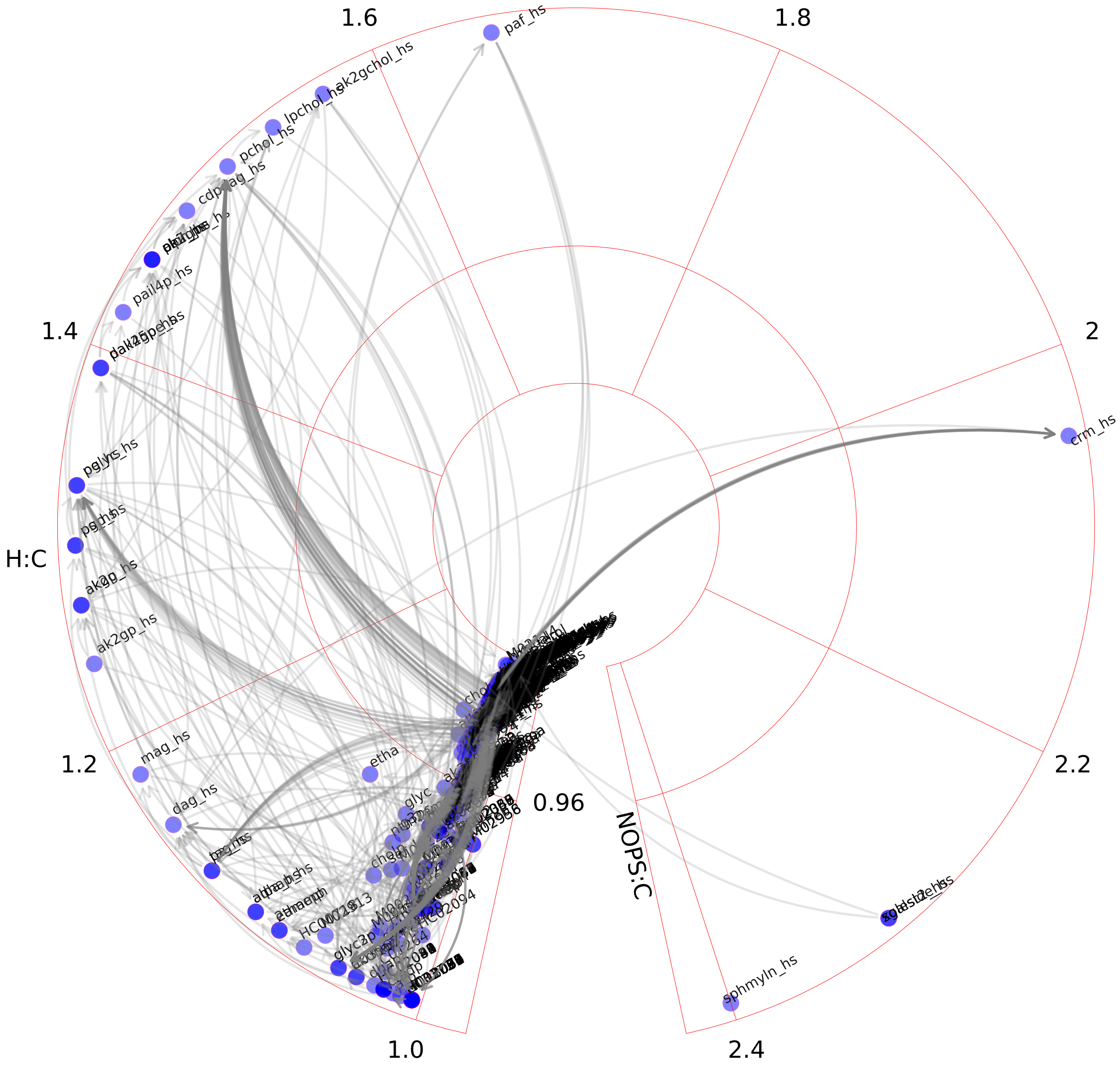

### Glycine, serine, alanine, and threonine metabolism.pdf

Glycine, serine, alanine, and threonine metabolism

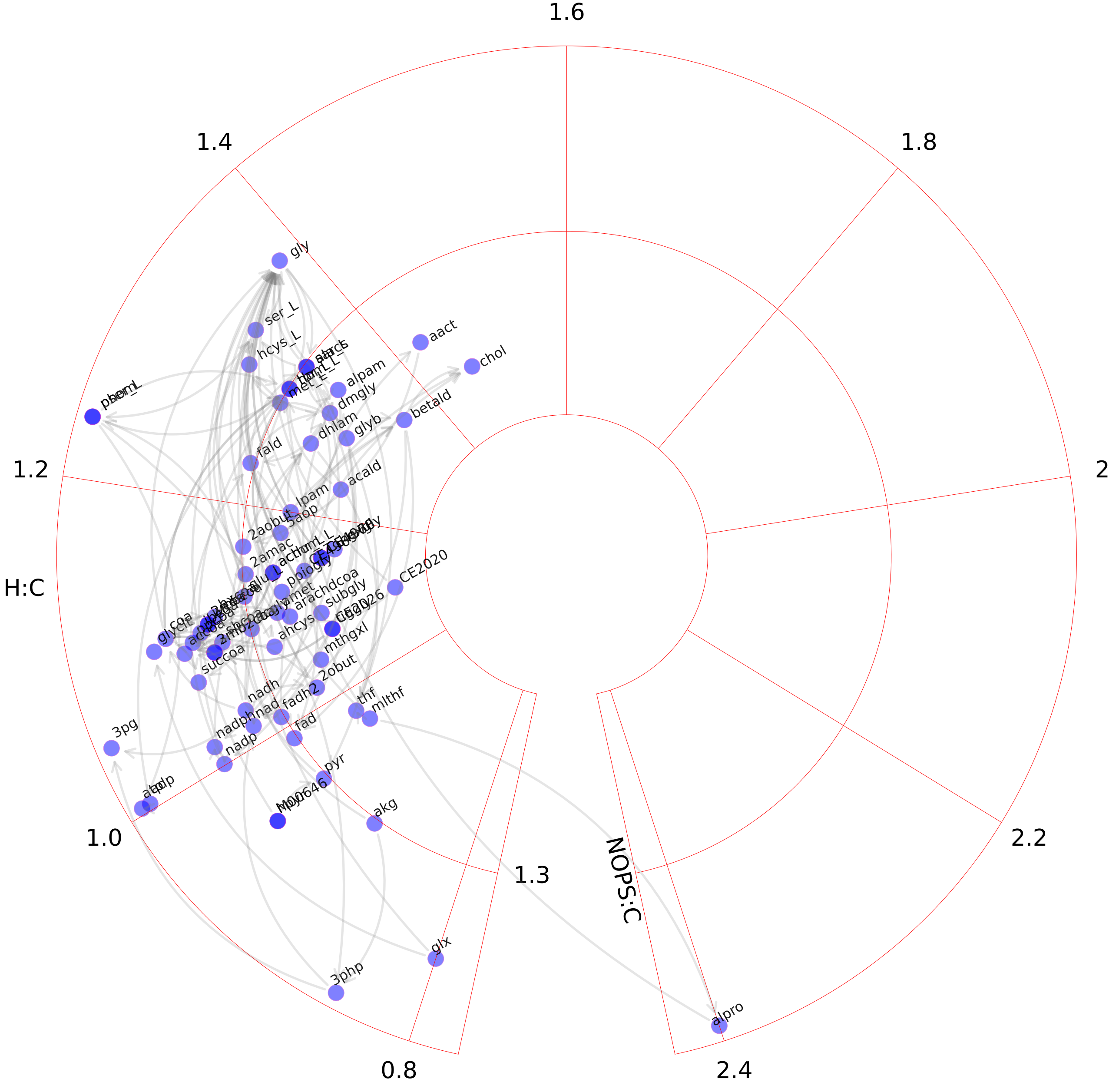

### Glycolysis_gluconeogenesis.pdf

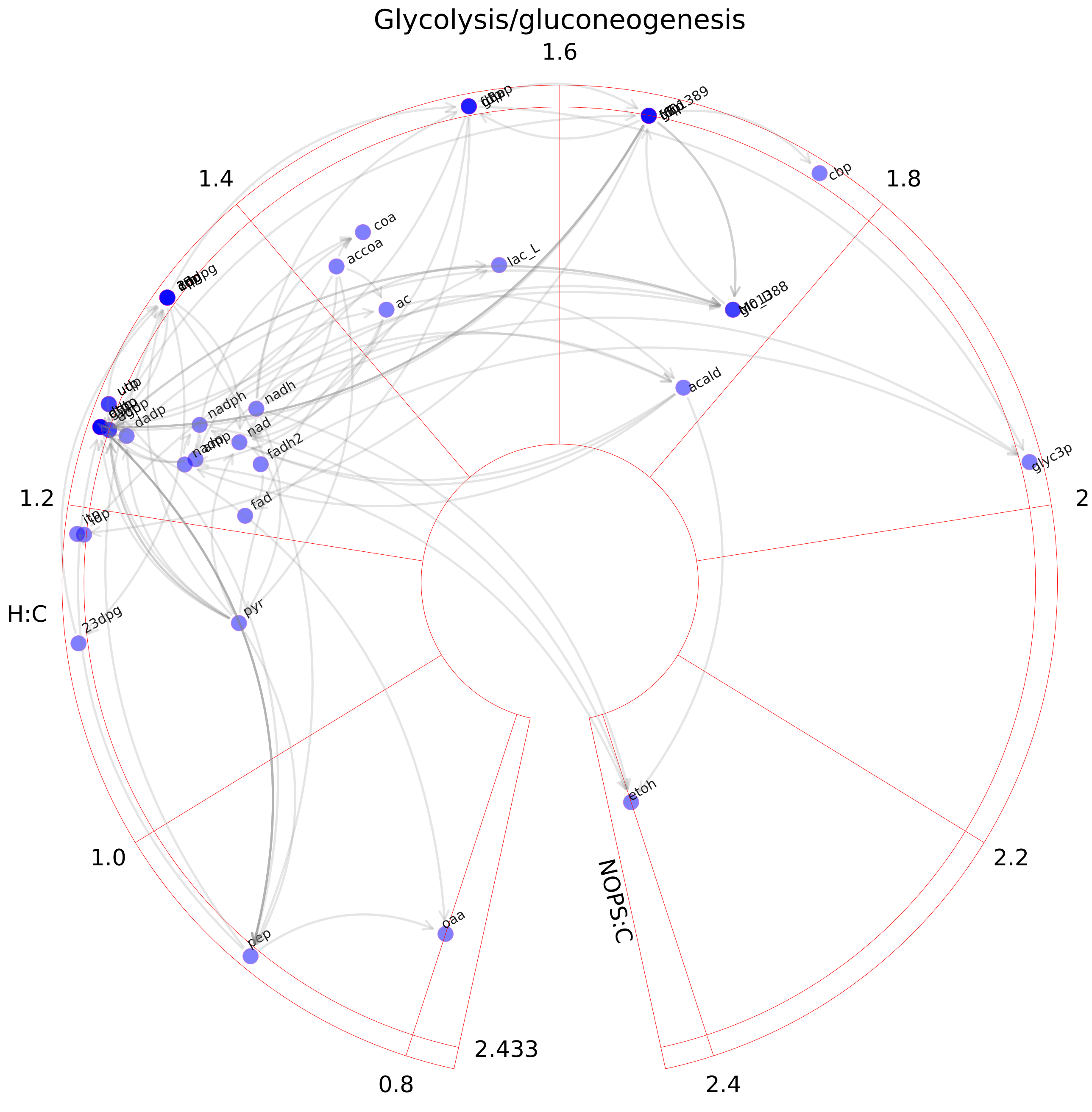

### Glycosphingolipid metabolism.pdf

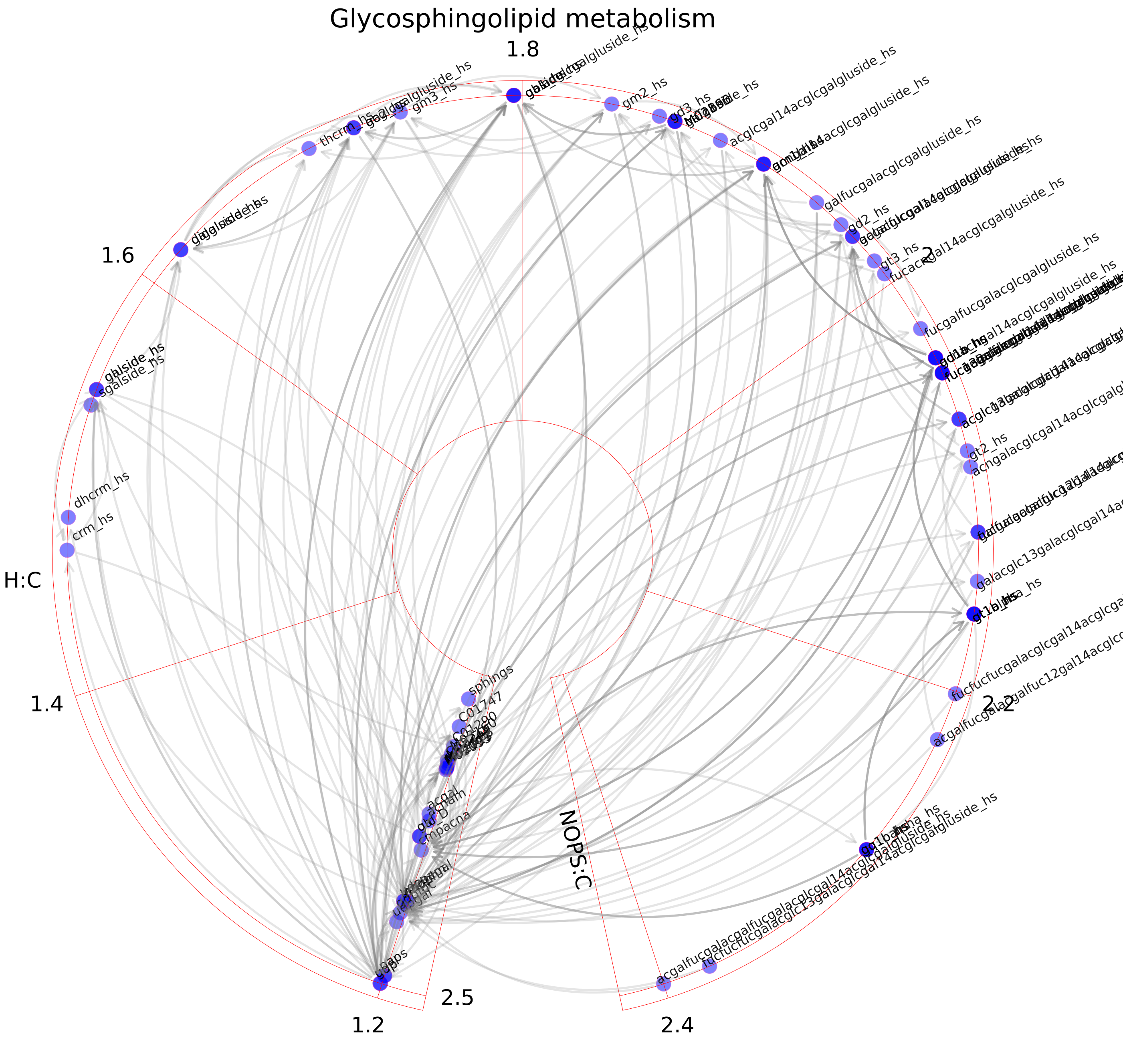

### Glyoxylate and dicarboxylate metabolism.pdf

Glyoxylate and dicarboxylate metabolism

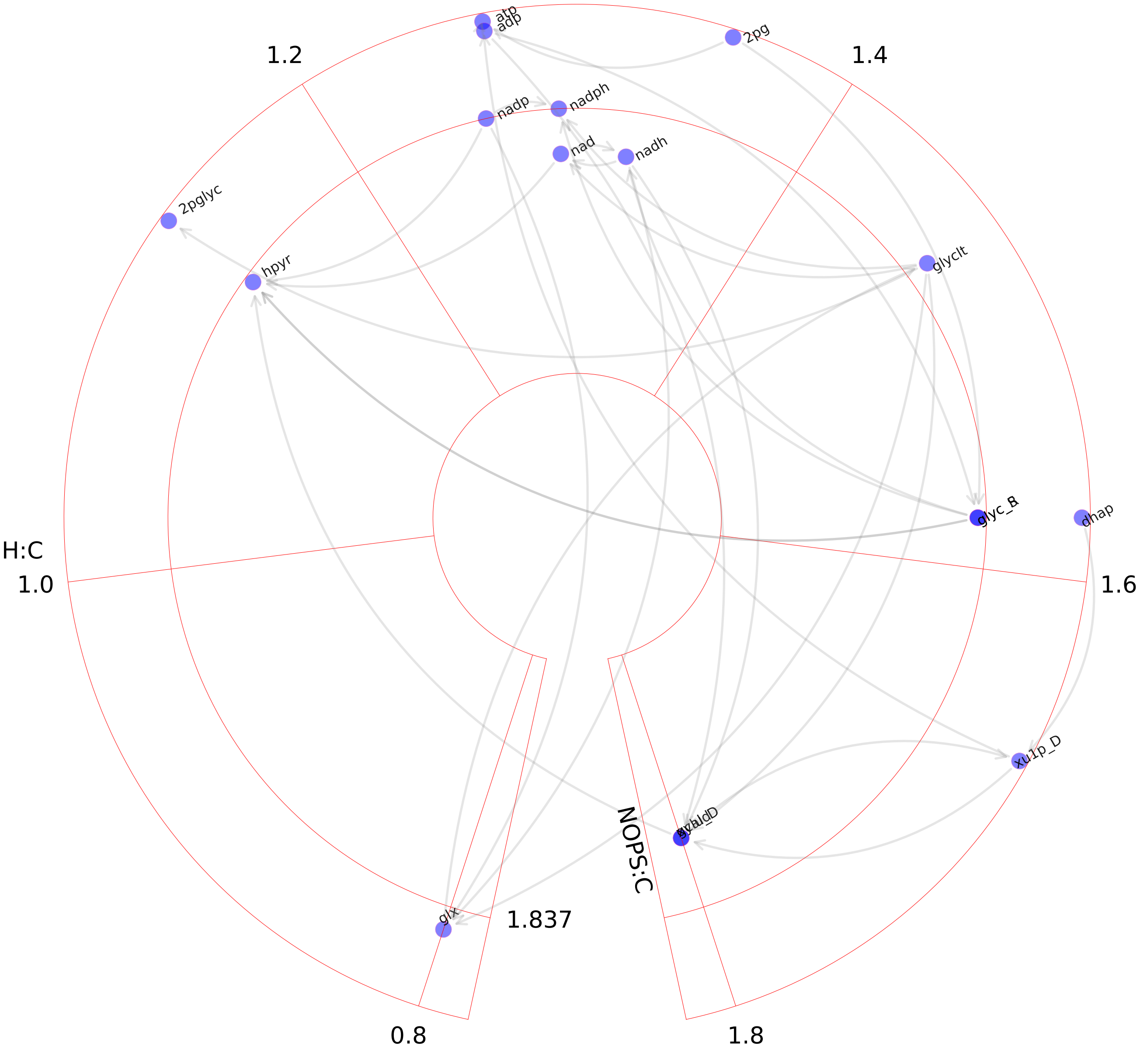

### Heme degradation.pdf

#### Heme degradation

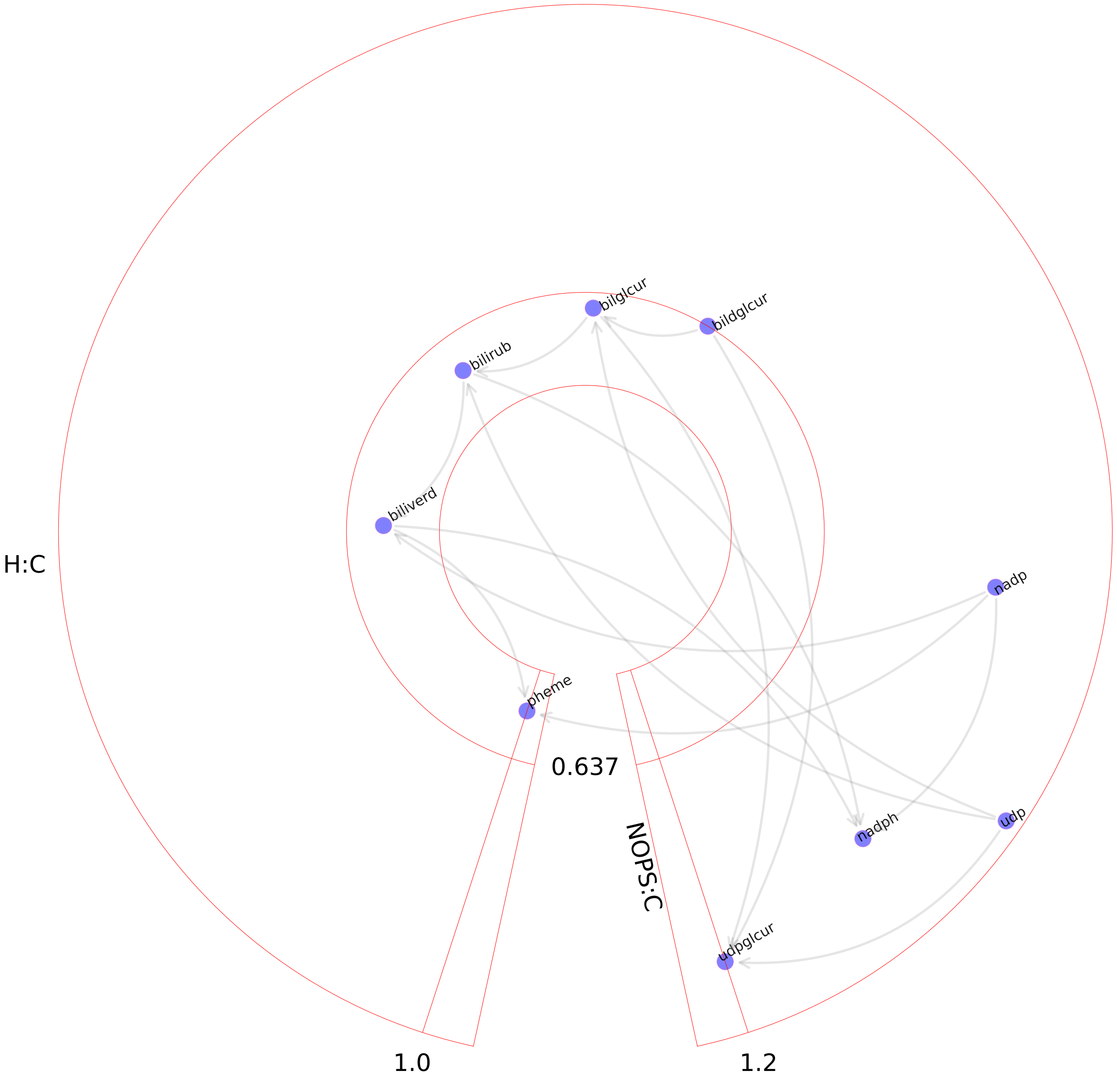

### Heme synthesis.pdf

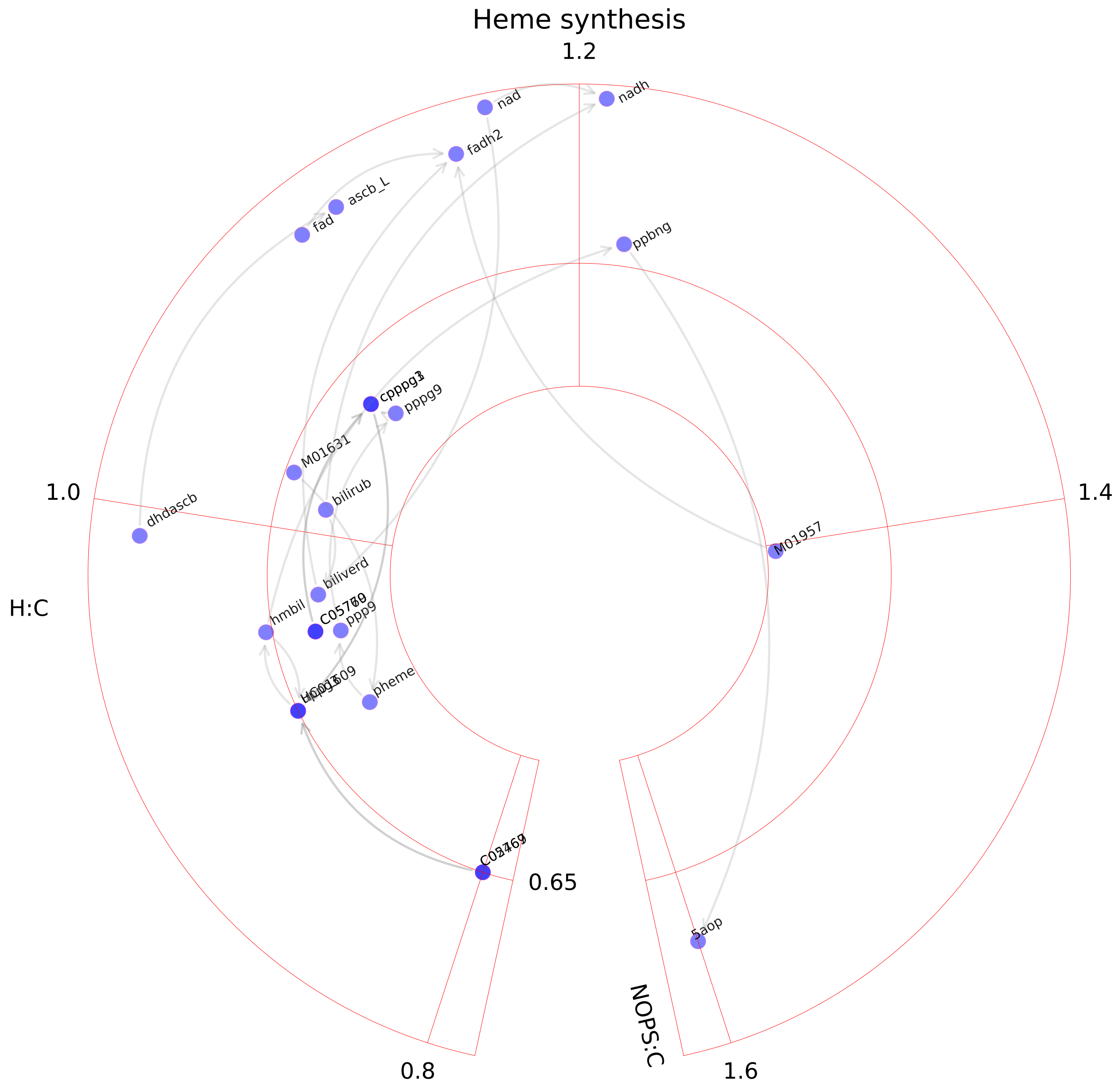

### Heparan sulfate degradation.pdf

Heparan sulfate degradation

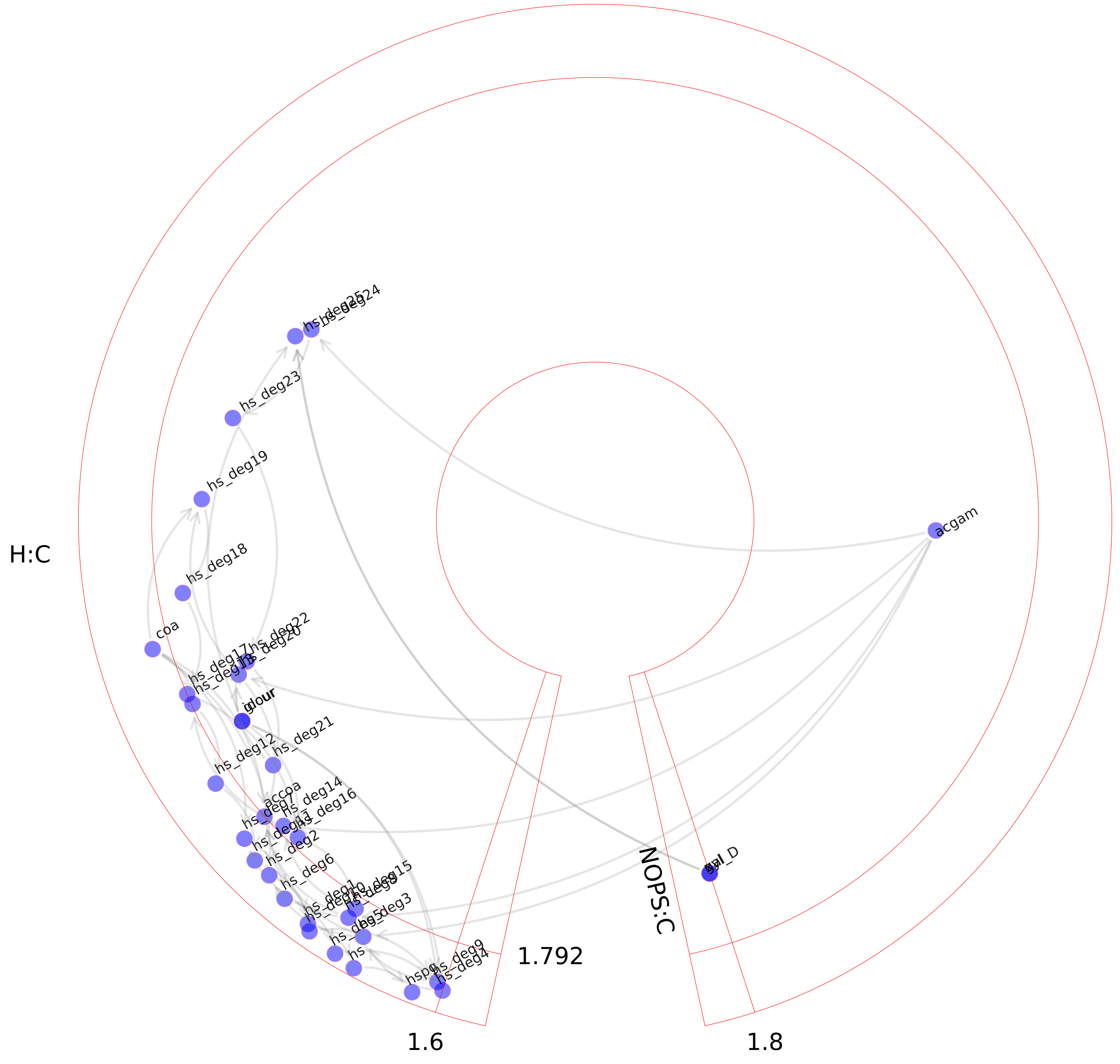

### Hippurate metabolism.pdf

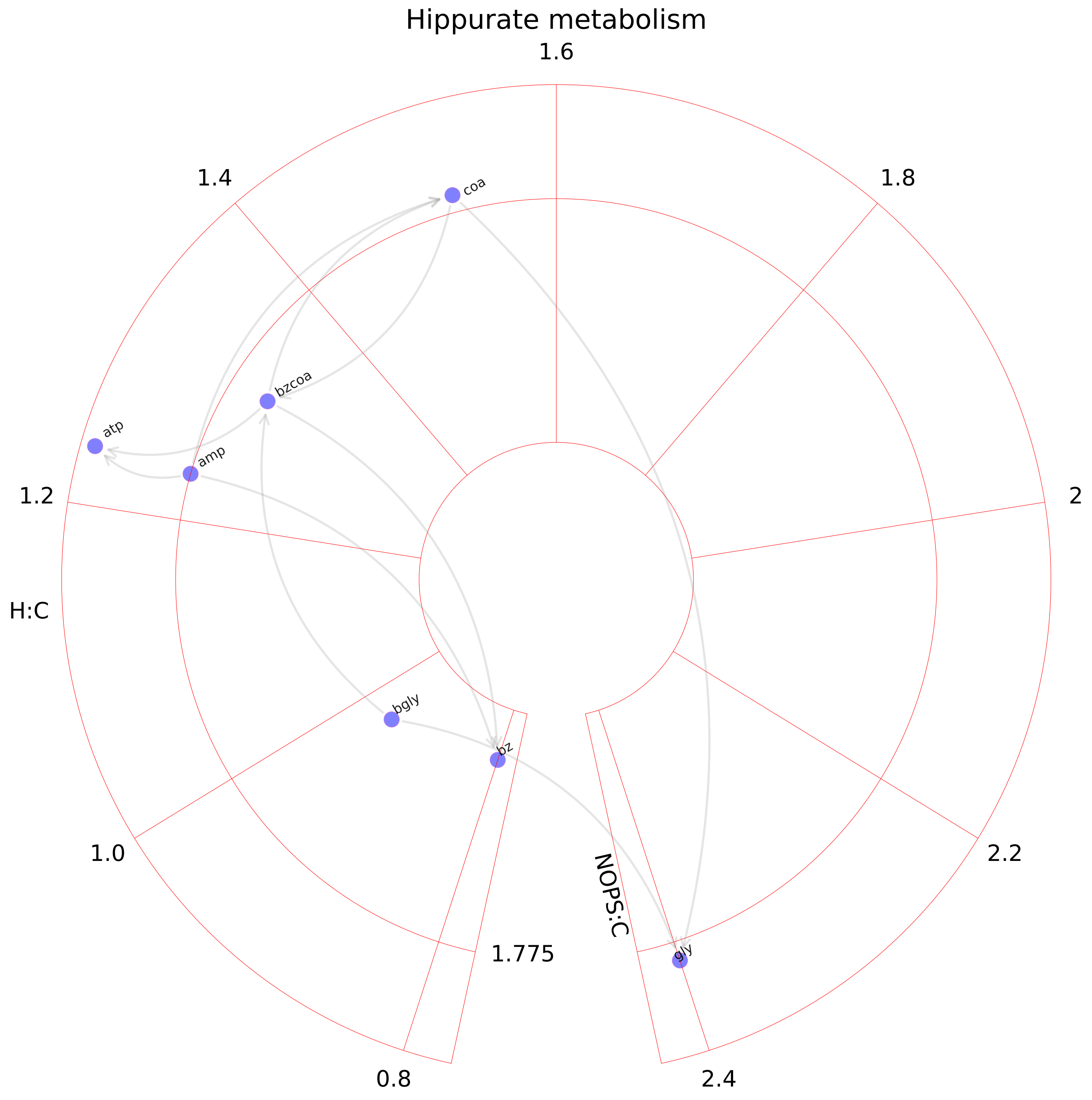

### Histidine metabolism.pdf

#### Histidine metabolism

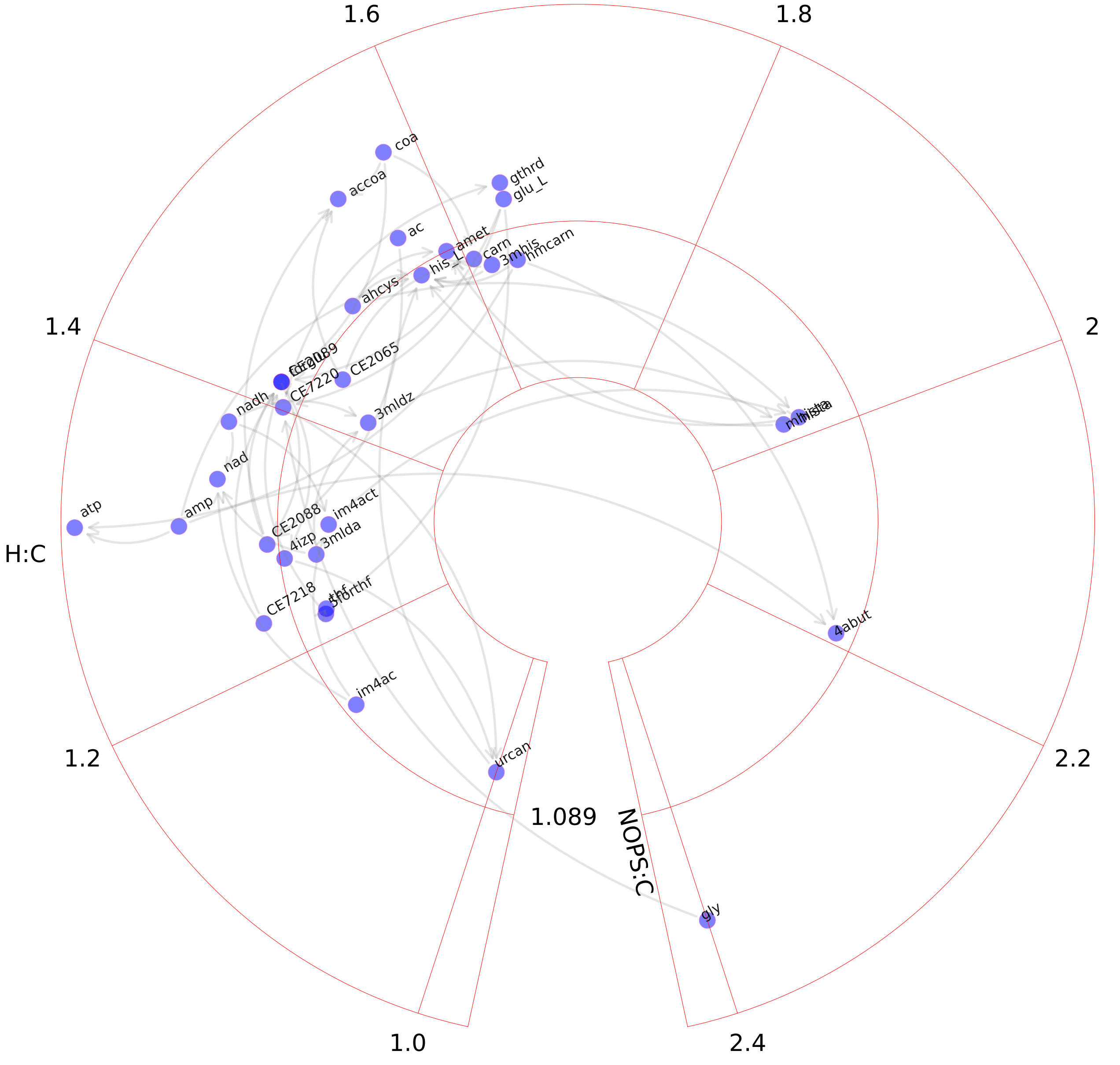

### Hyaluronan metabolism.pdf

#### Hyaluronan metabolism

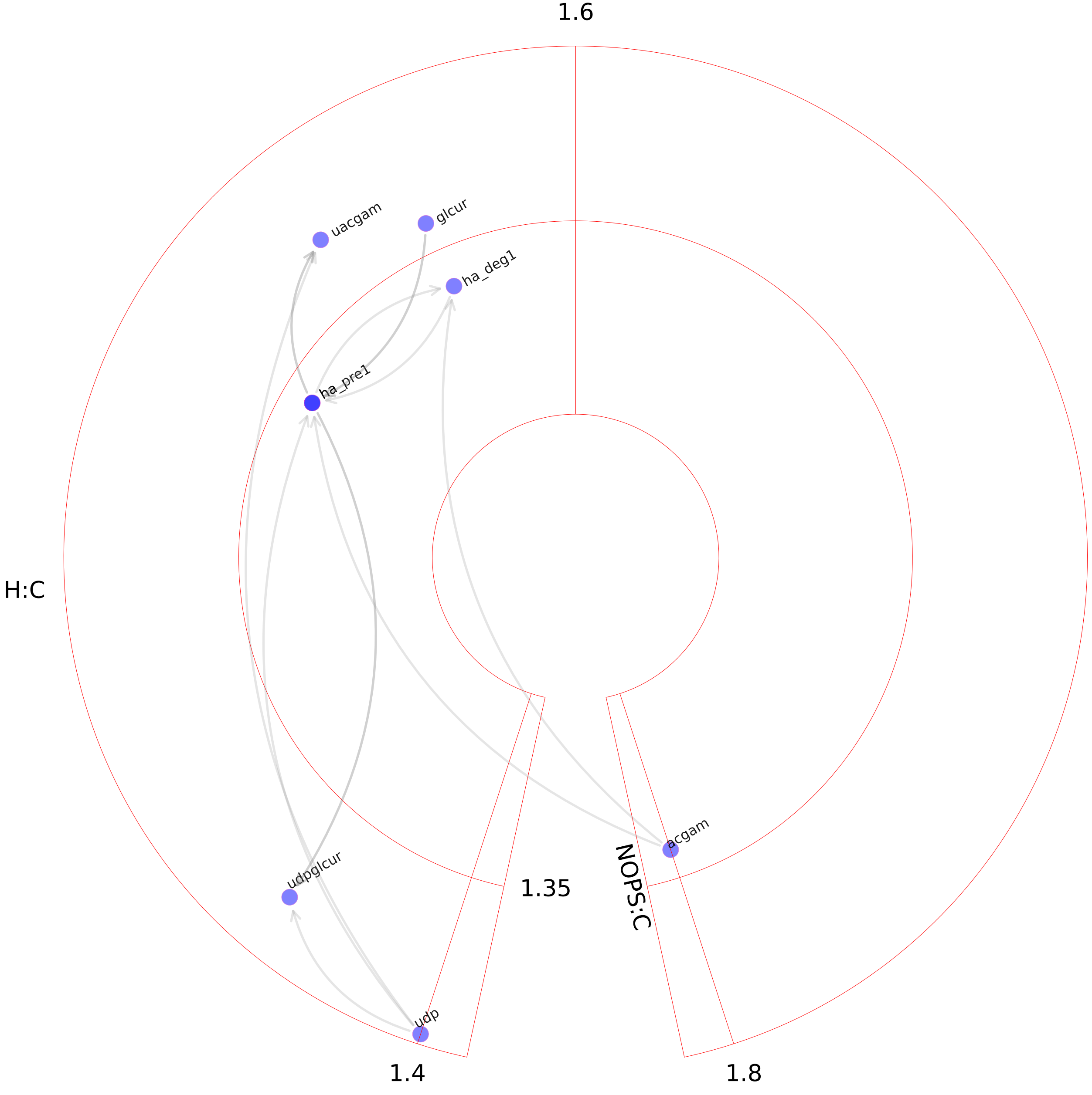

### Inositol phosphate metabolism.pdf

Inositol phosphate metabolism

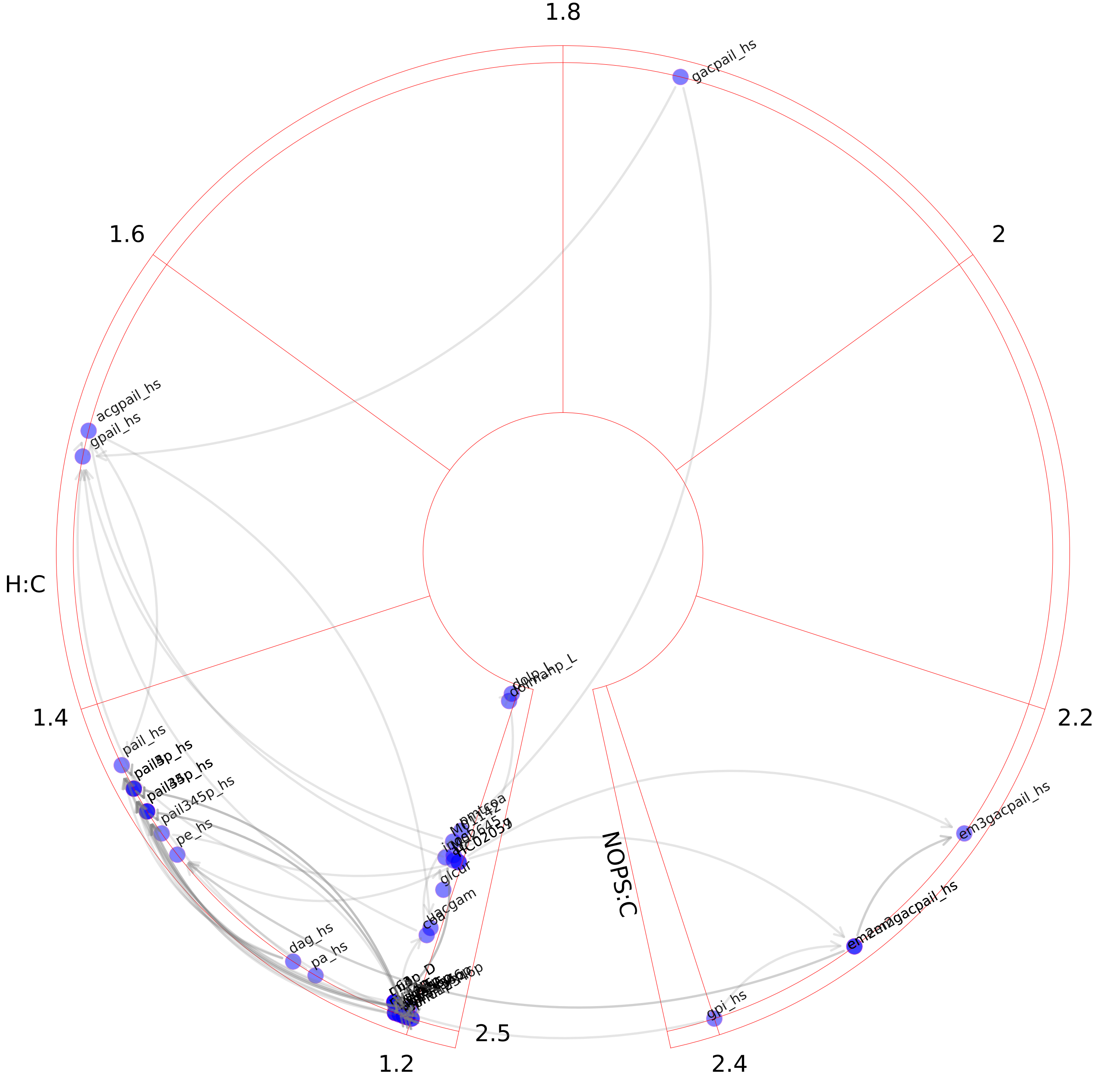

### Keratan sulfate synthesis.pdf

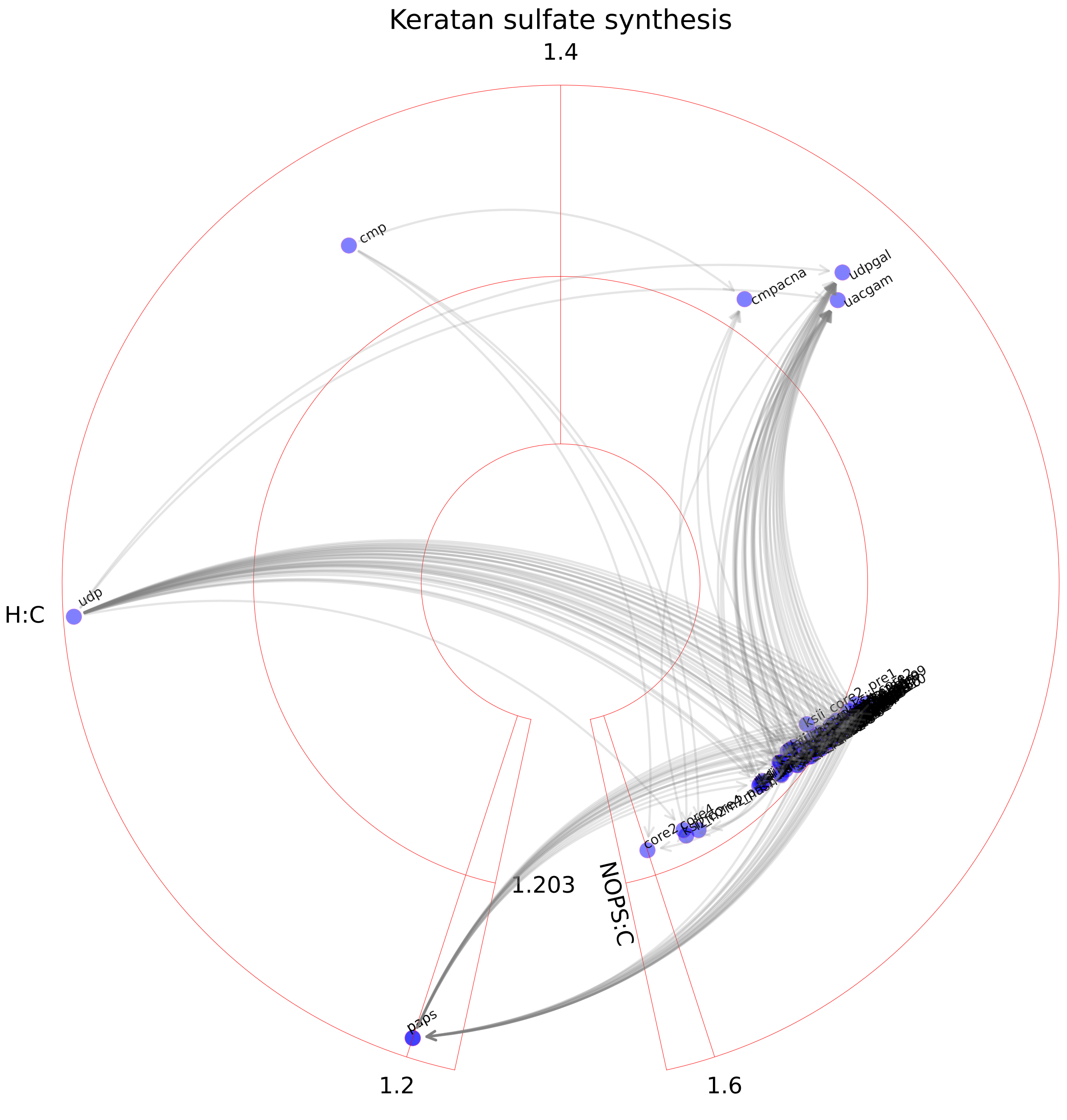

### Leukotriene metabolism.pdf

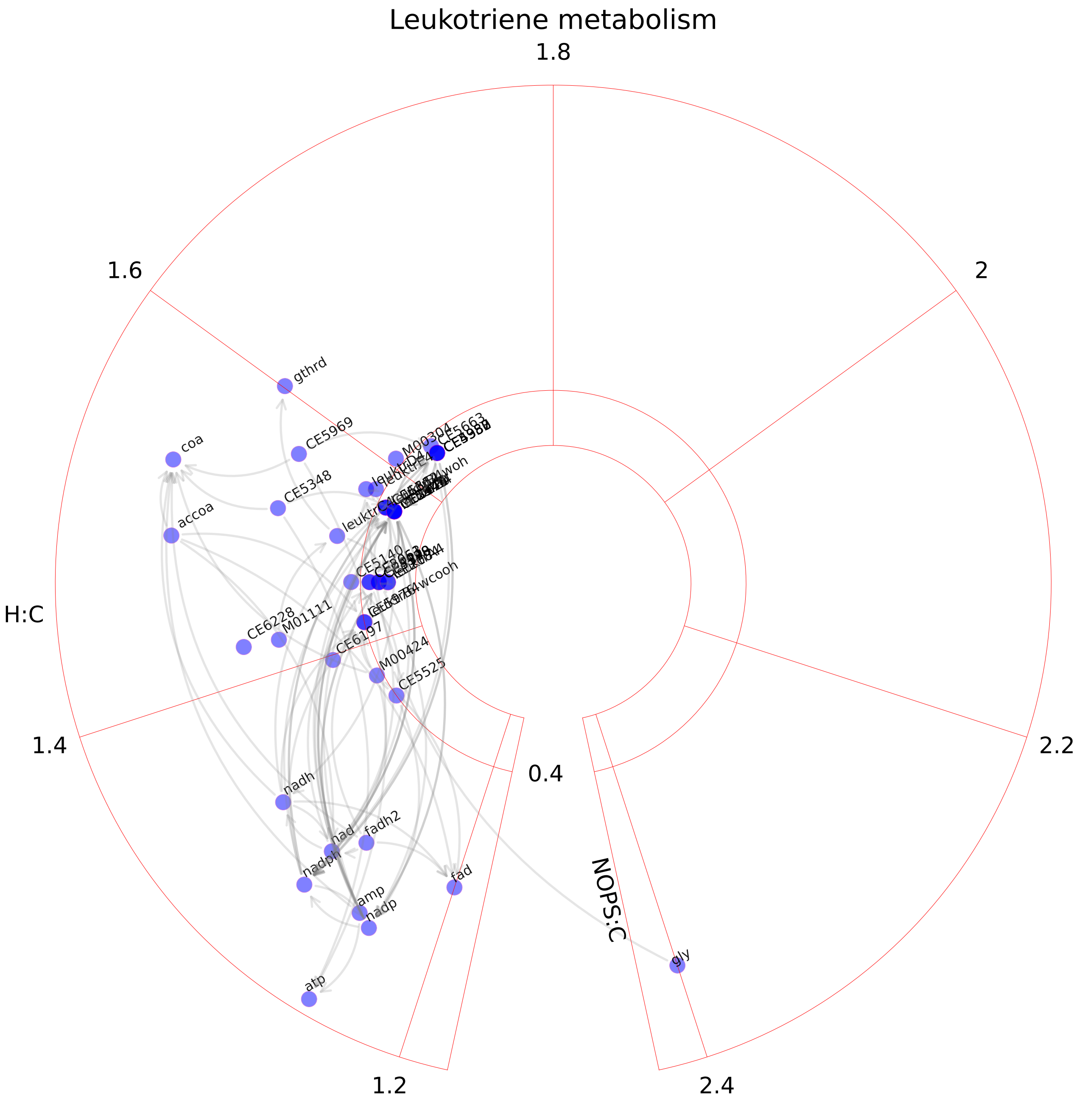

### Limonene and pinene degradation.pdf

##### Limonene and pinene degradation

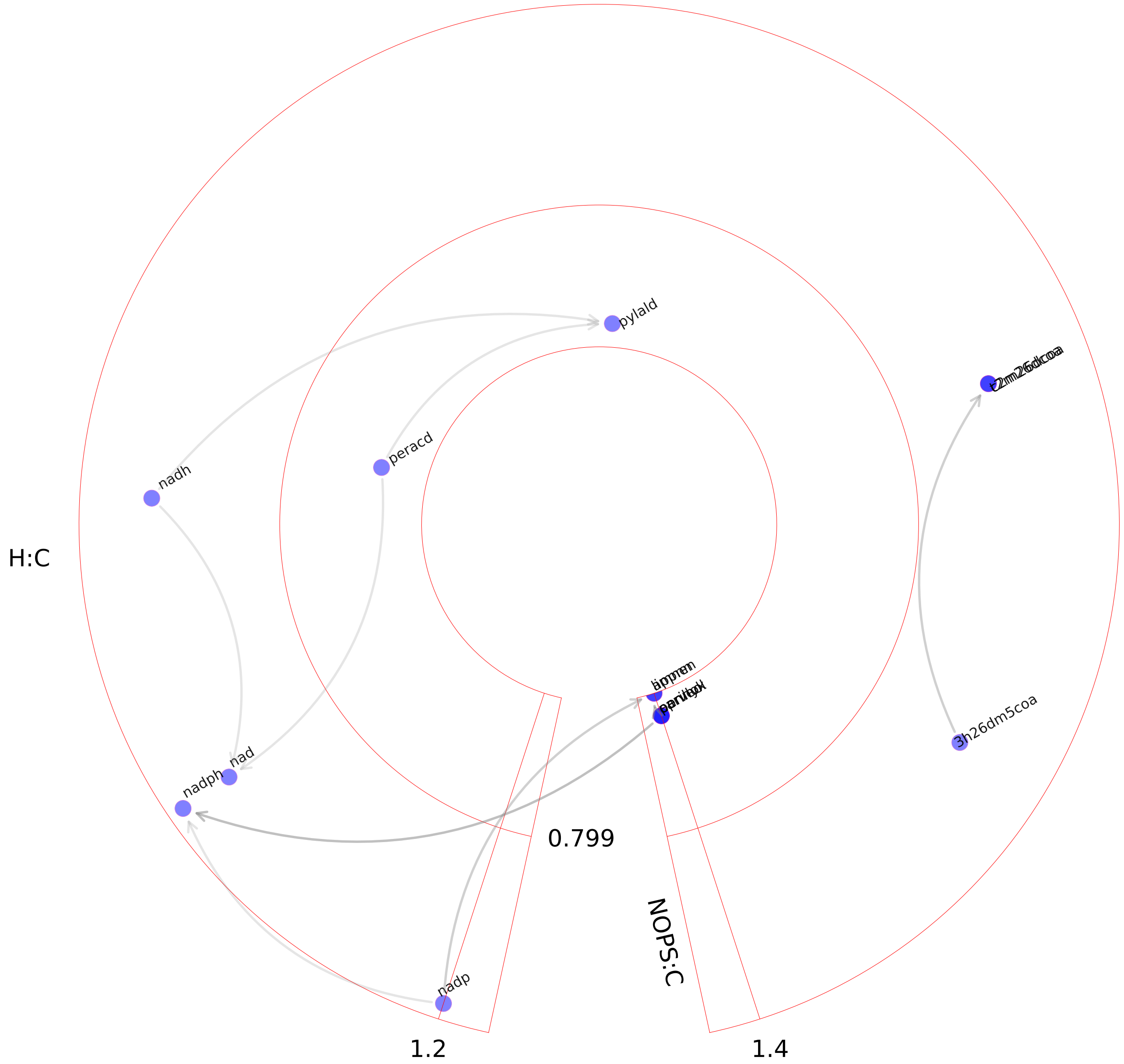

### Linoleate metabolism.pdf

#### Linoleate metabolism

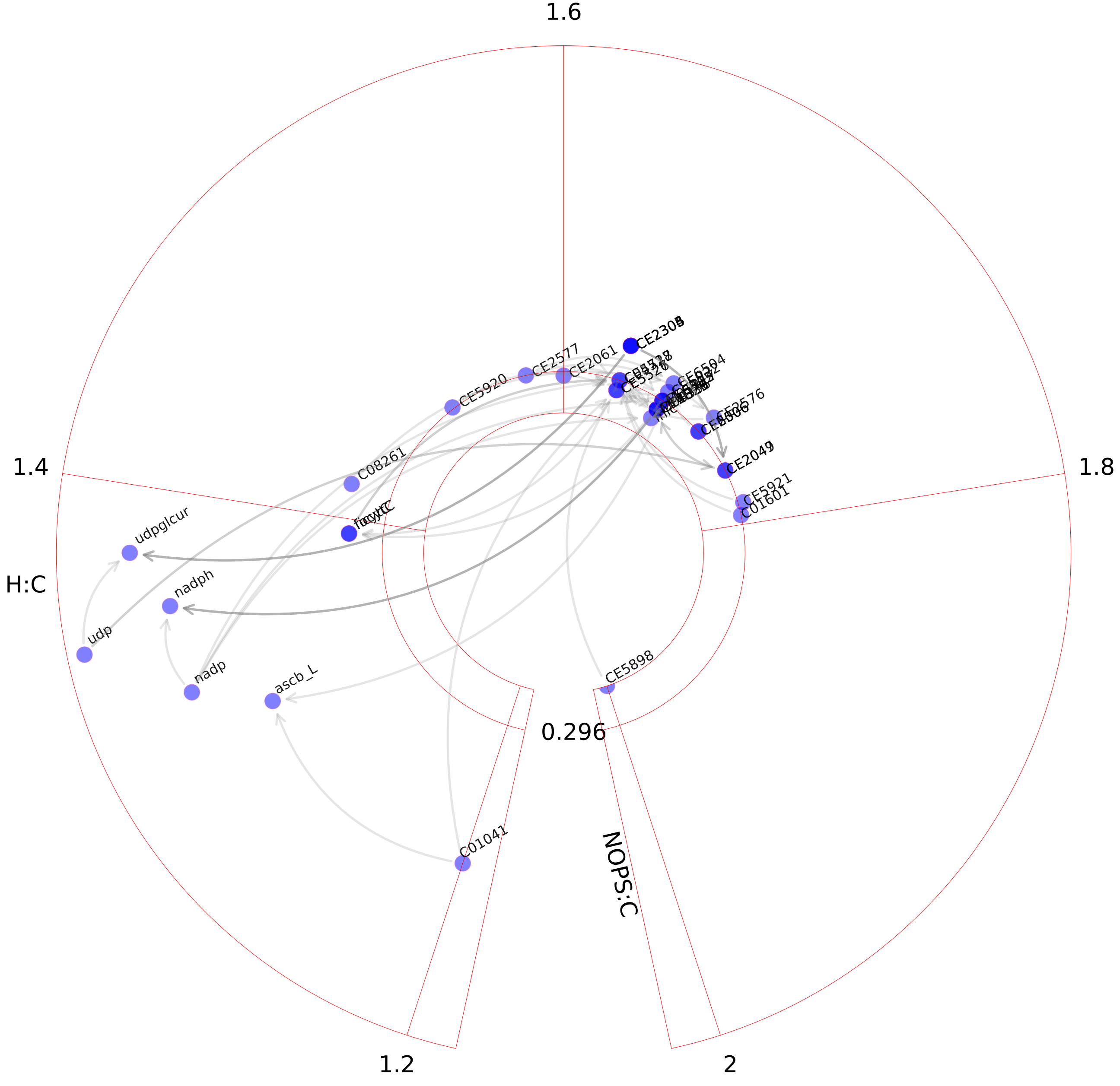

### Lipoate metabolism.pdf

#### Lipoate metabolism

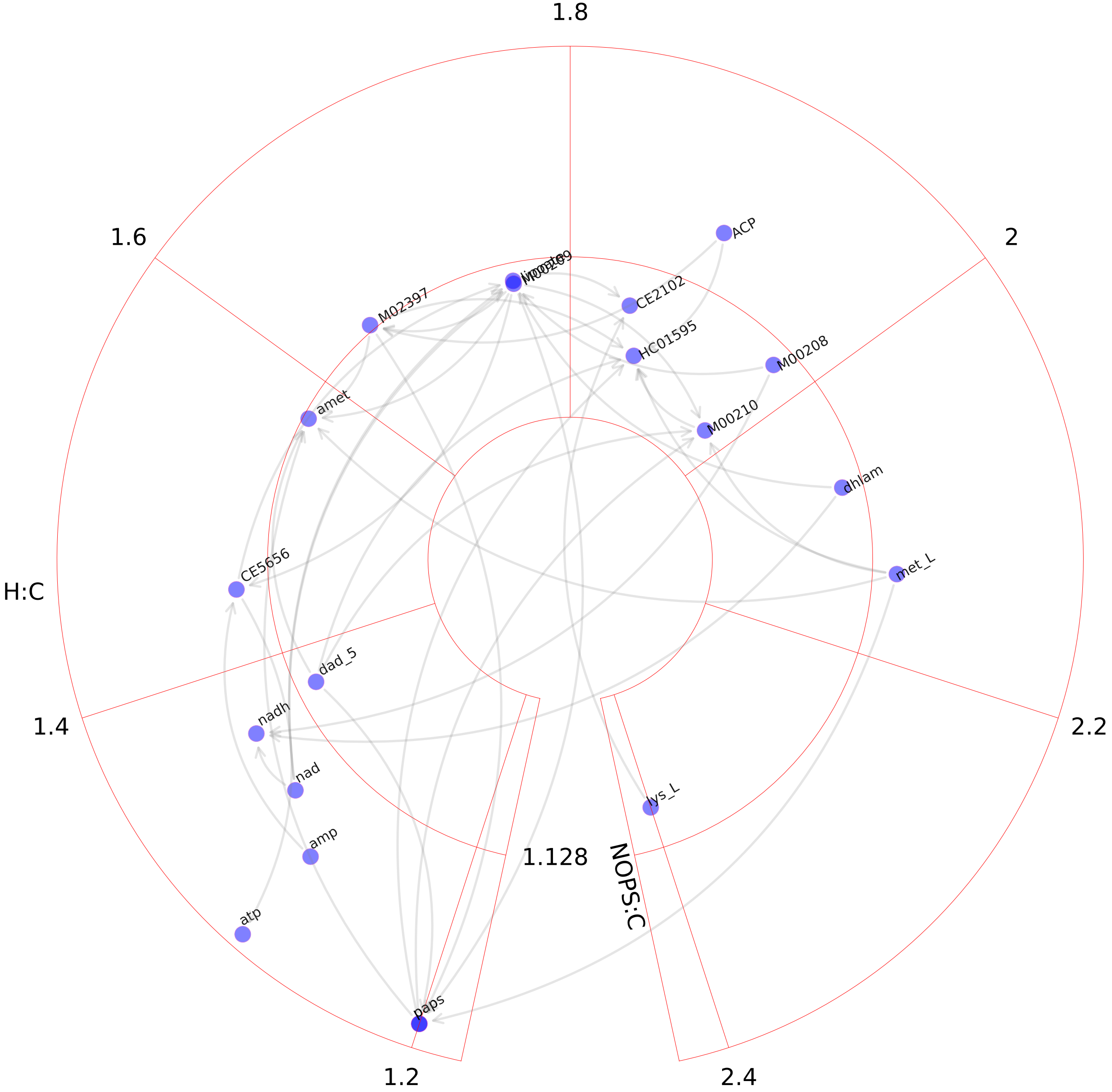

### Lysine metabolism.pdf

#### Lysine metabolism

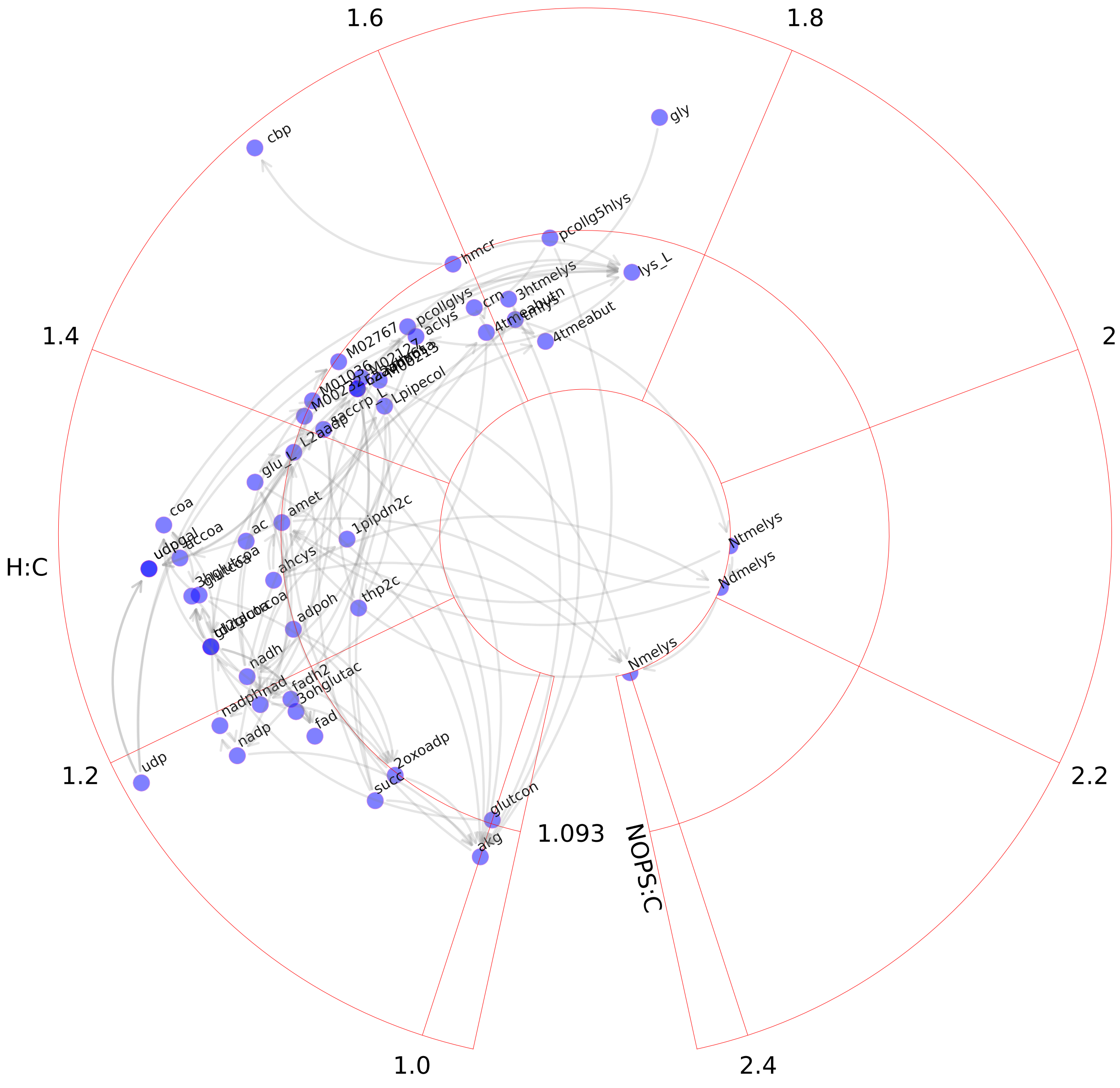

### Methionine and cysteine metabolism.pdf

#### Methionine and cysteine metabolism

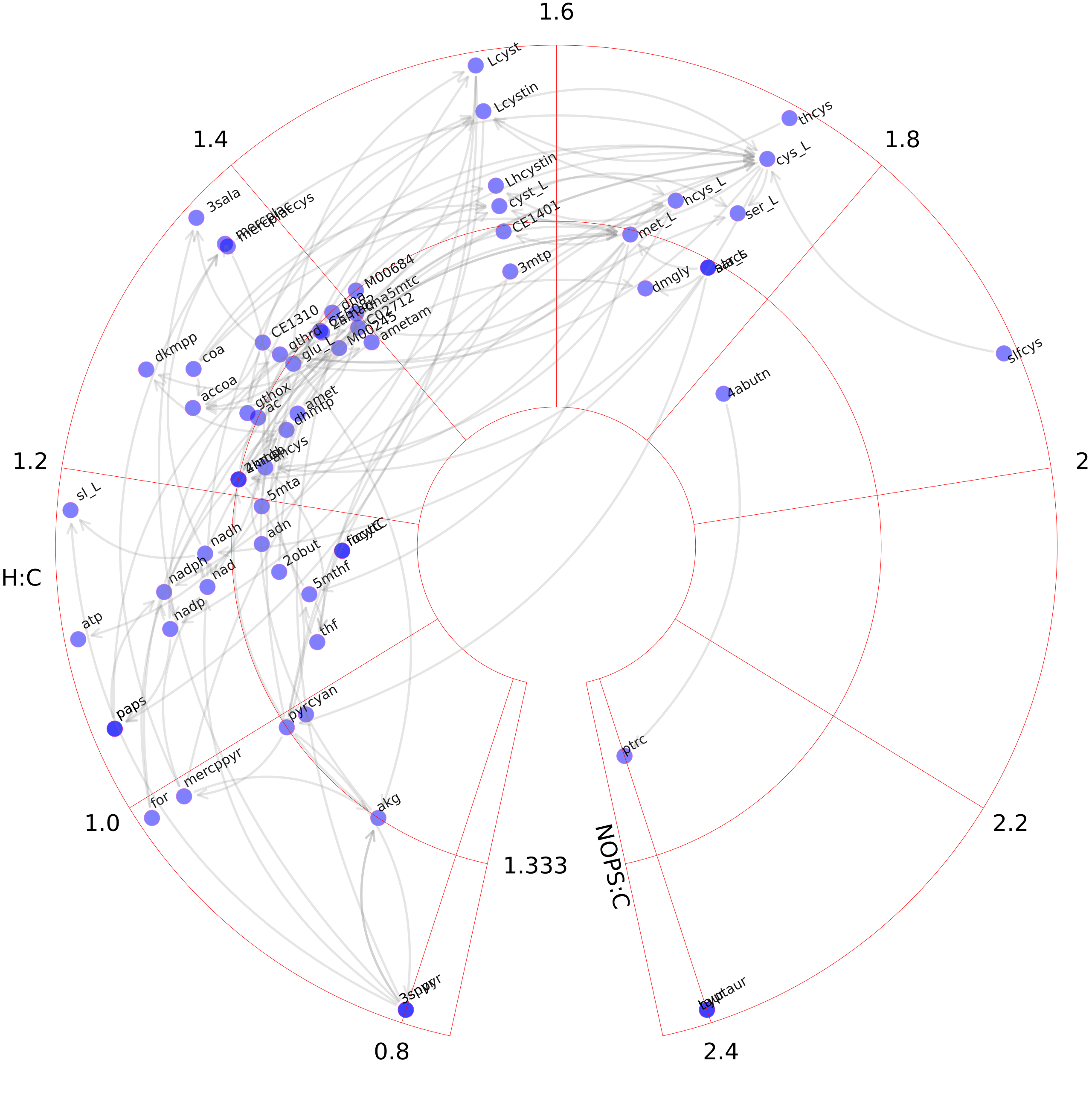

### N-glycan degradation.pdf

#### N-glycan degradation

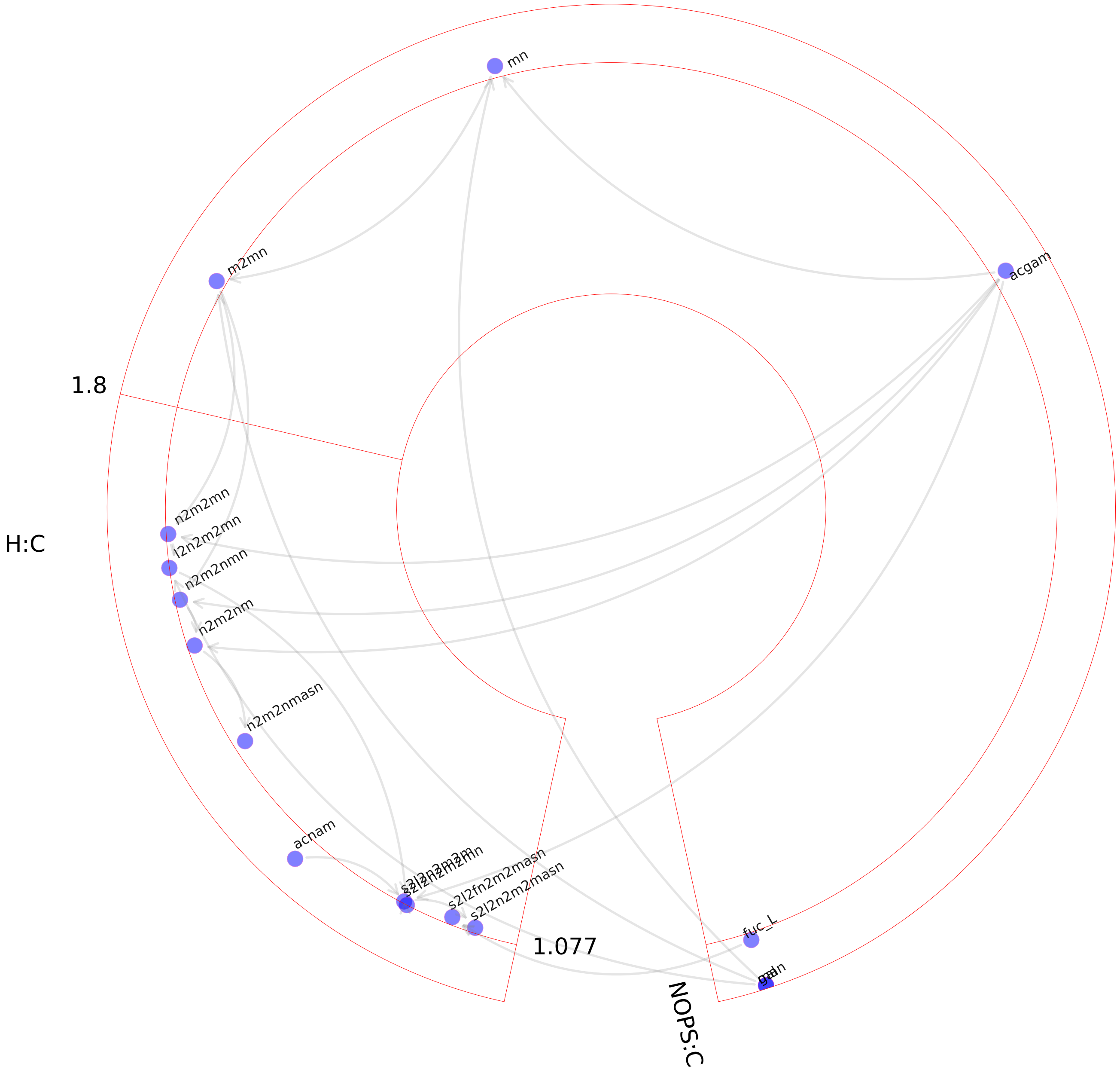

### N-glycan metabolism.pdf

#### N-glycan metabolism

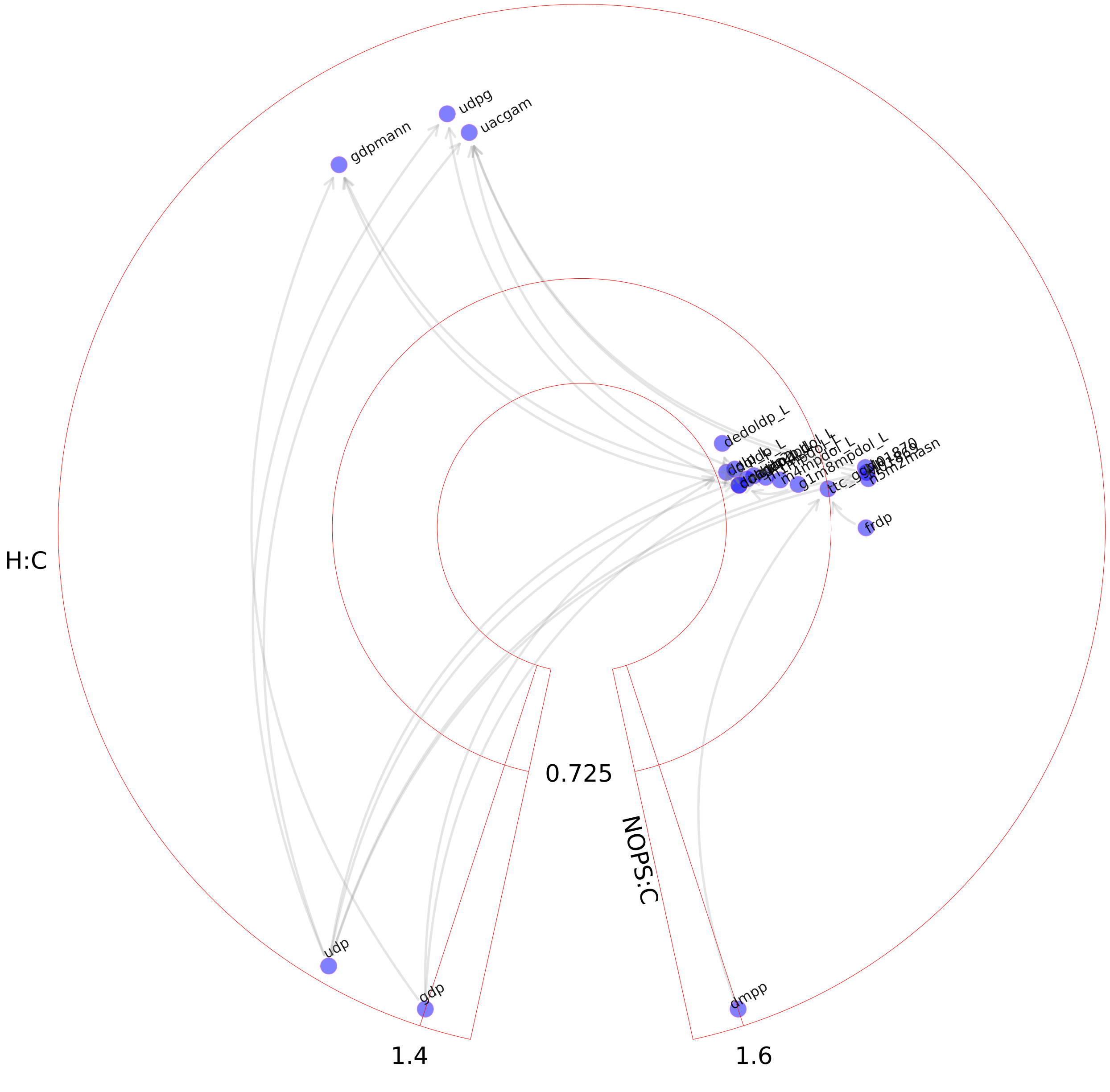

### N-glycan synthesis.pdf

#### N-glycan synthesis

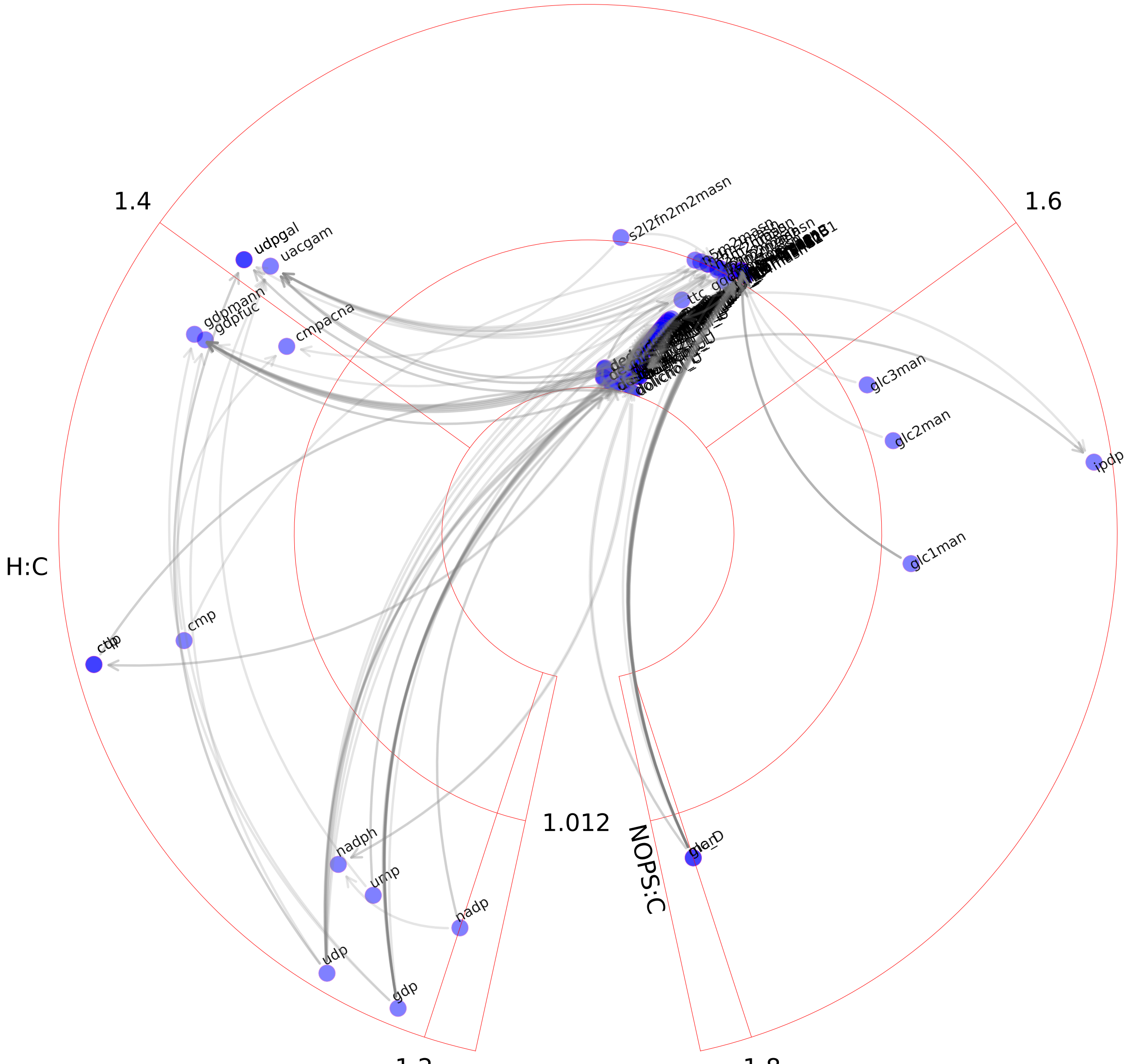

### NAD metabolism.pdf

#### NAD metabolism

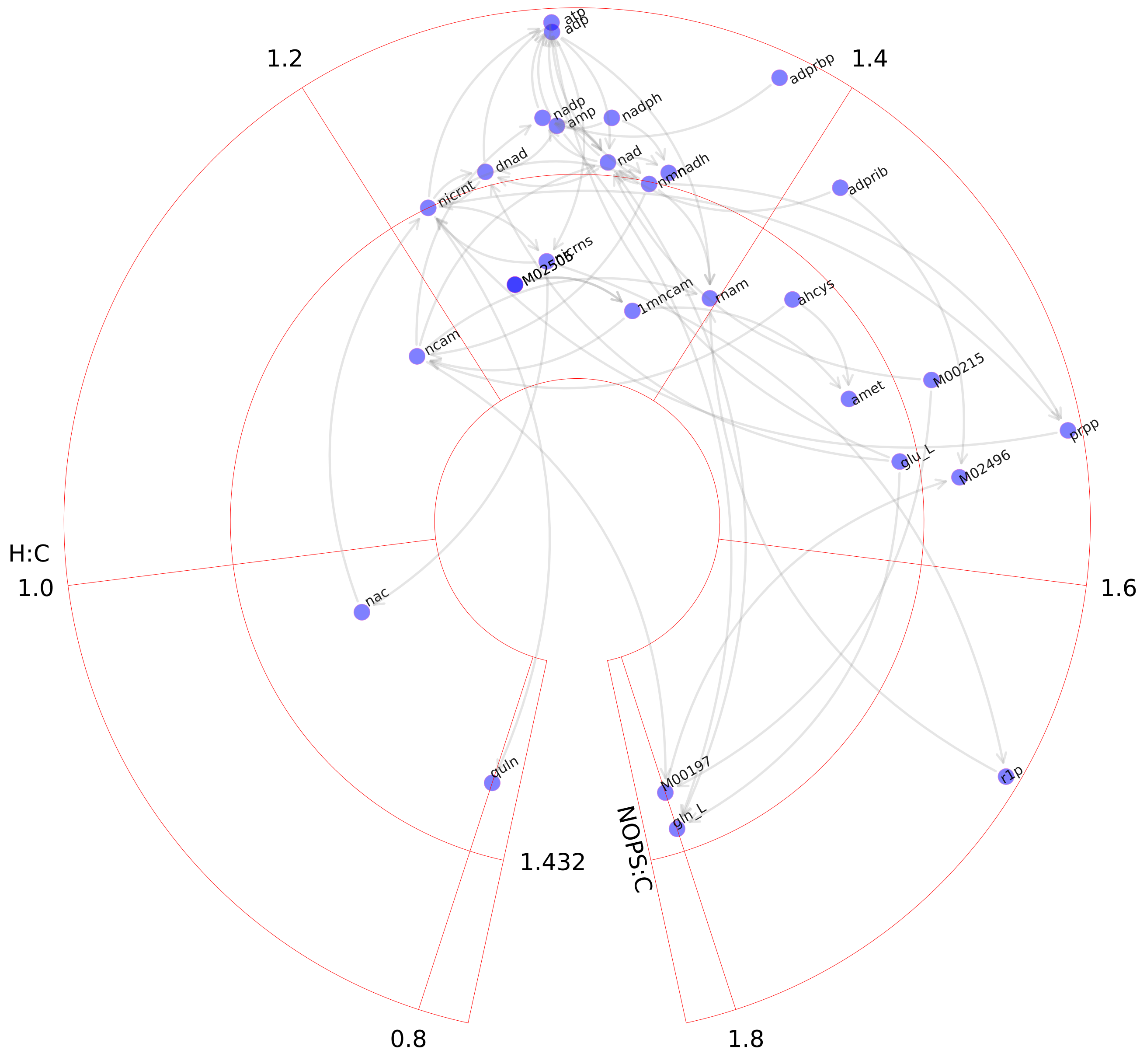

### Nucleotide interconversion.pdf

#### Nucleotide interconversion

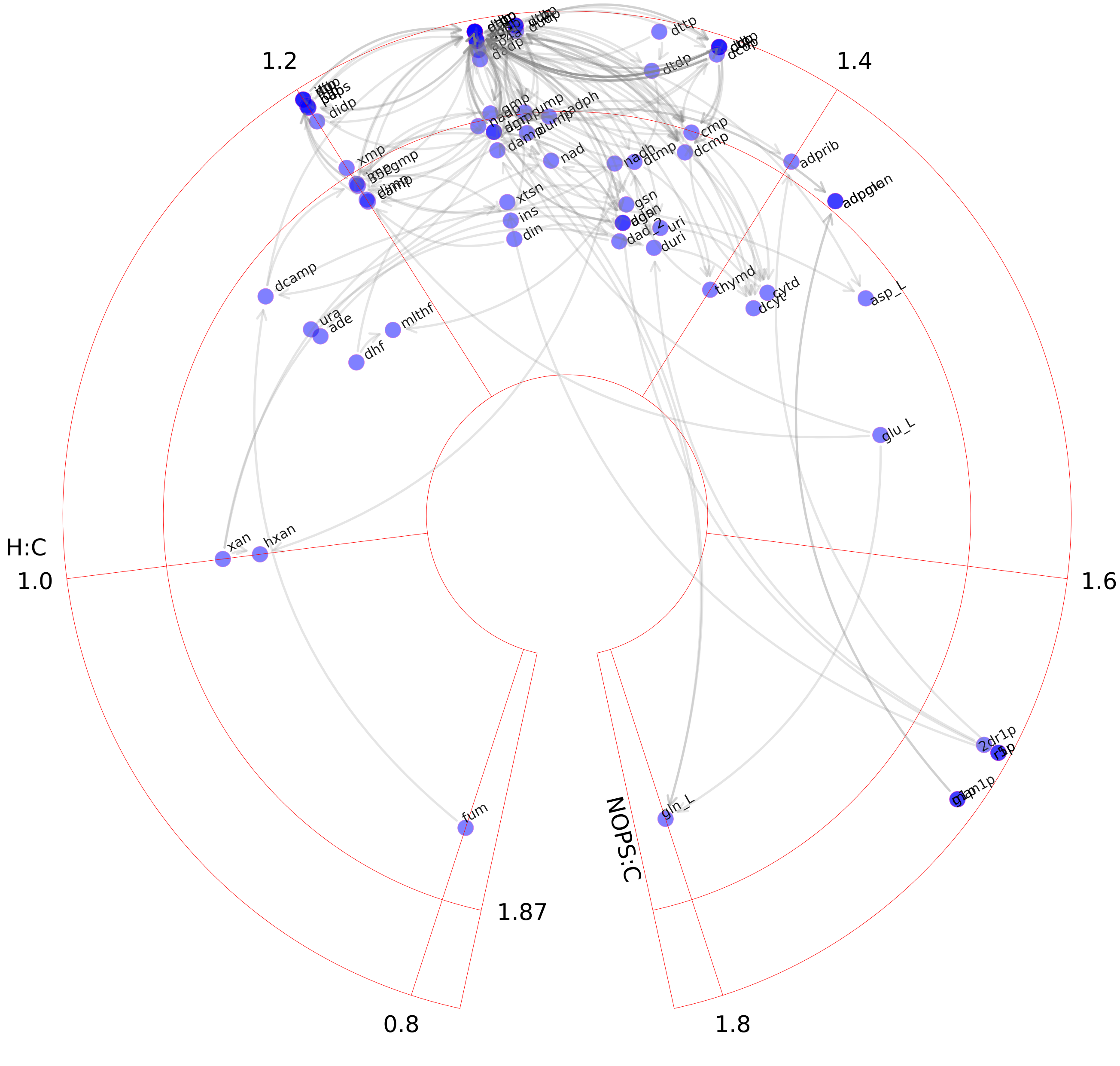

### Nucleotide salvage pathway.pdf

#### Nucleotide salvage pathway

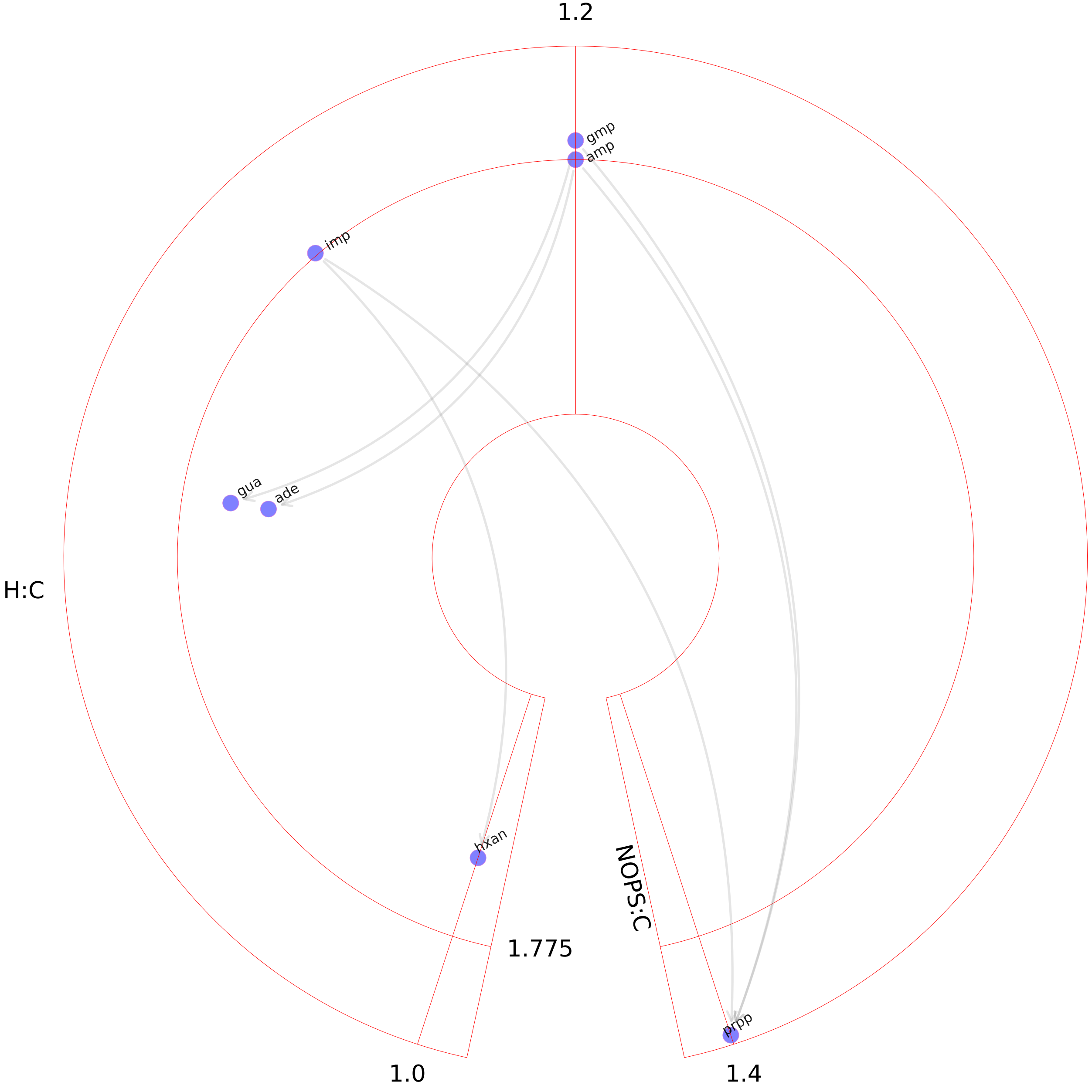

### Nucleotide sugar metabolism.pdf

#### Nucleotide sugar metabolism

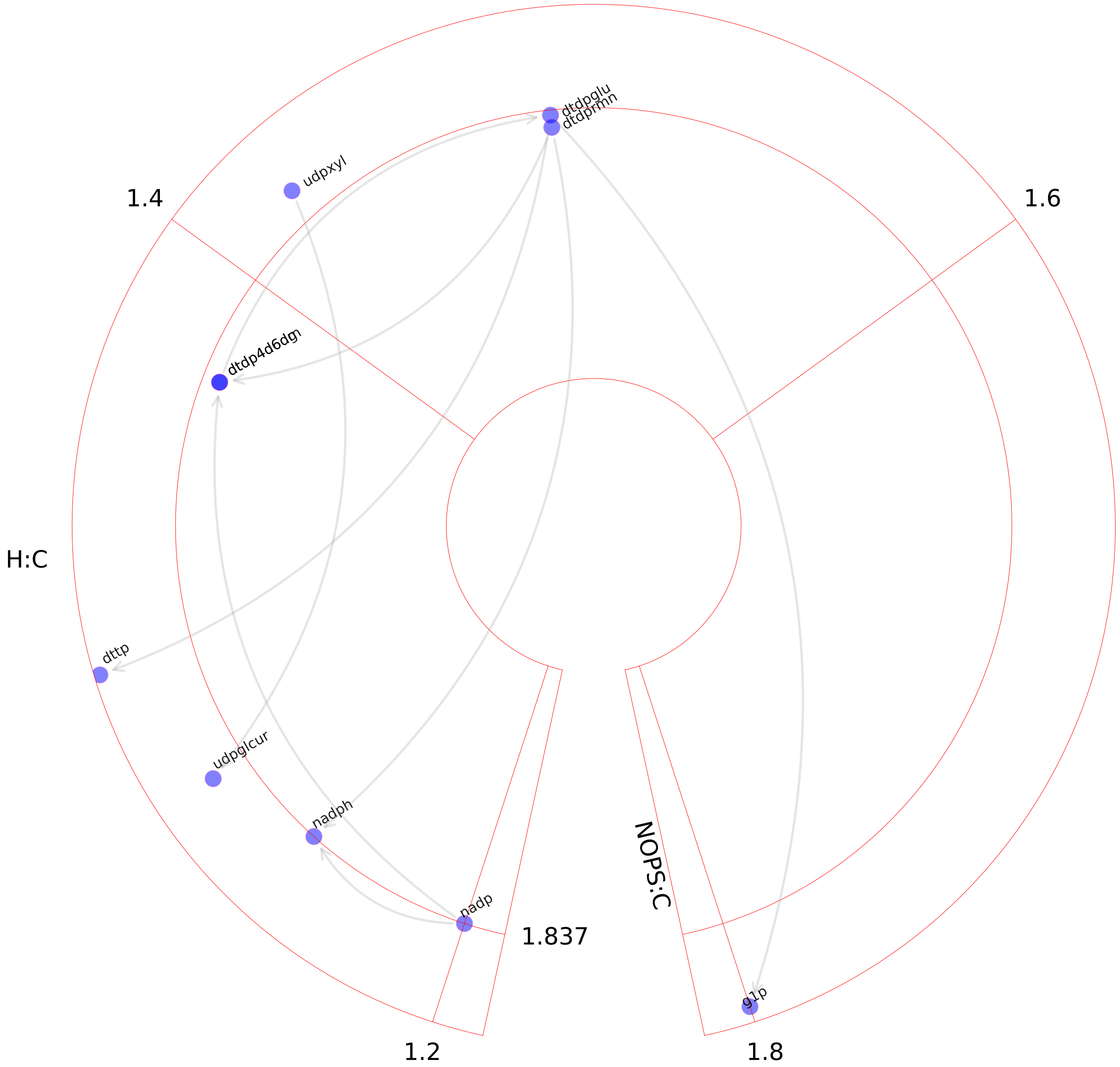

### O-glycan metabolism.pdf

#### O-glycan metabolism

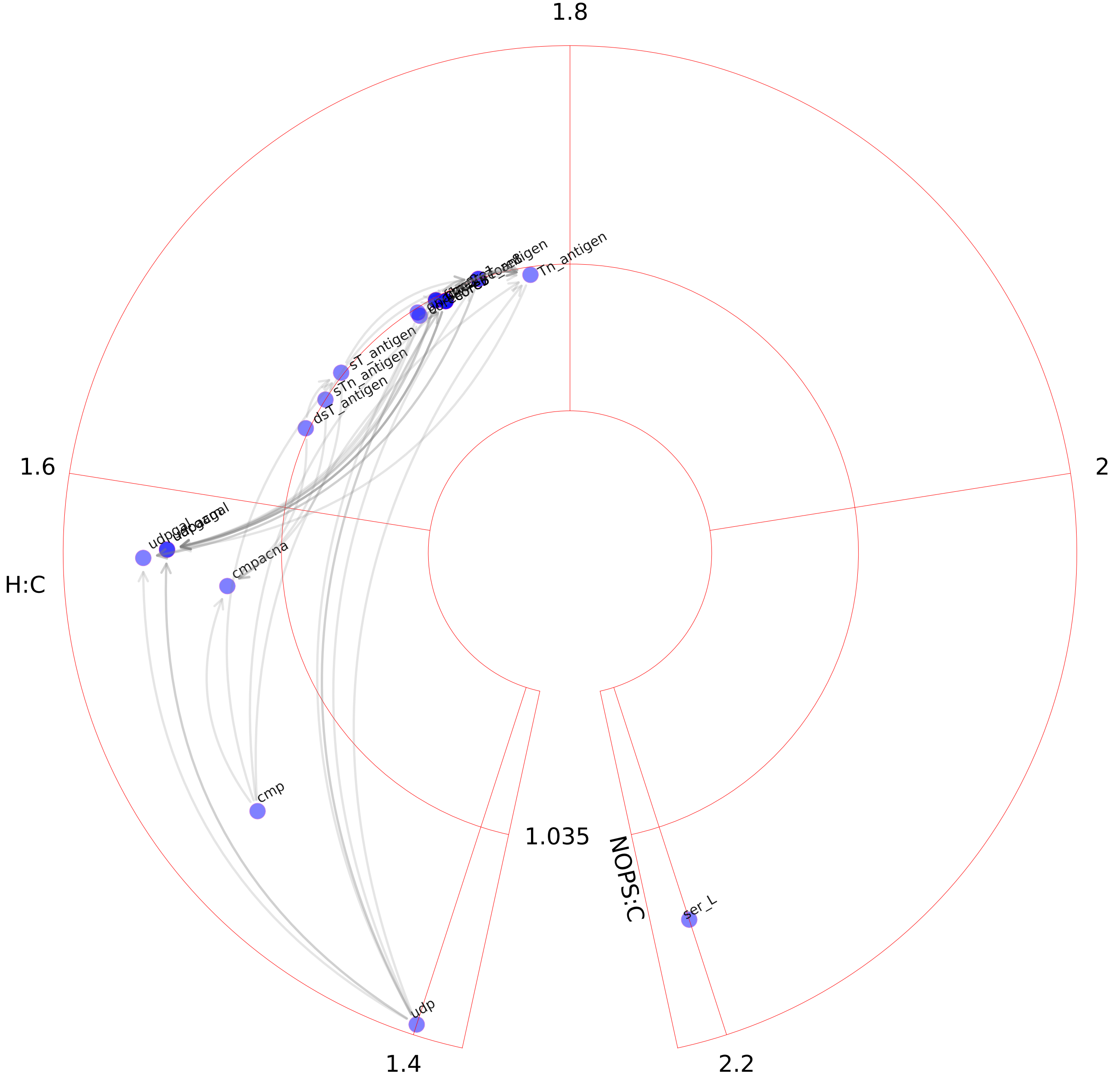

### Oxidative phosphorylation.pdf

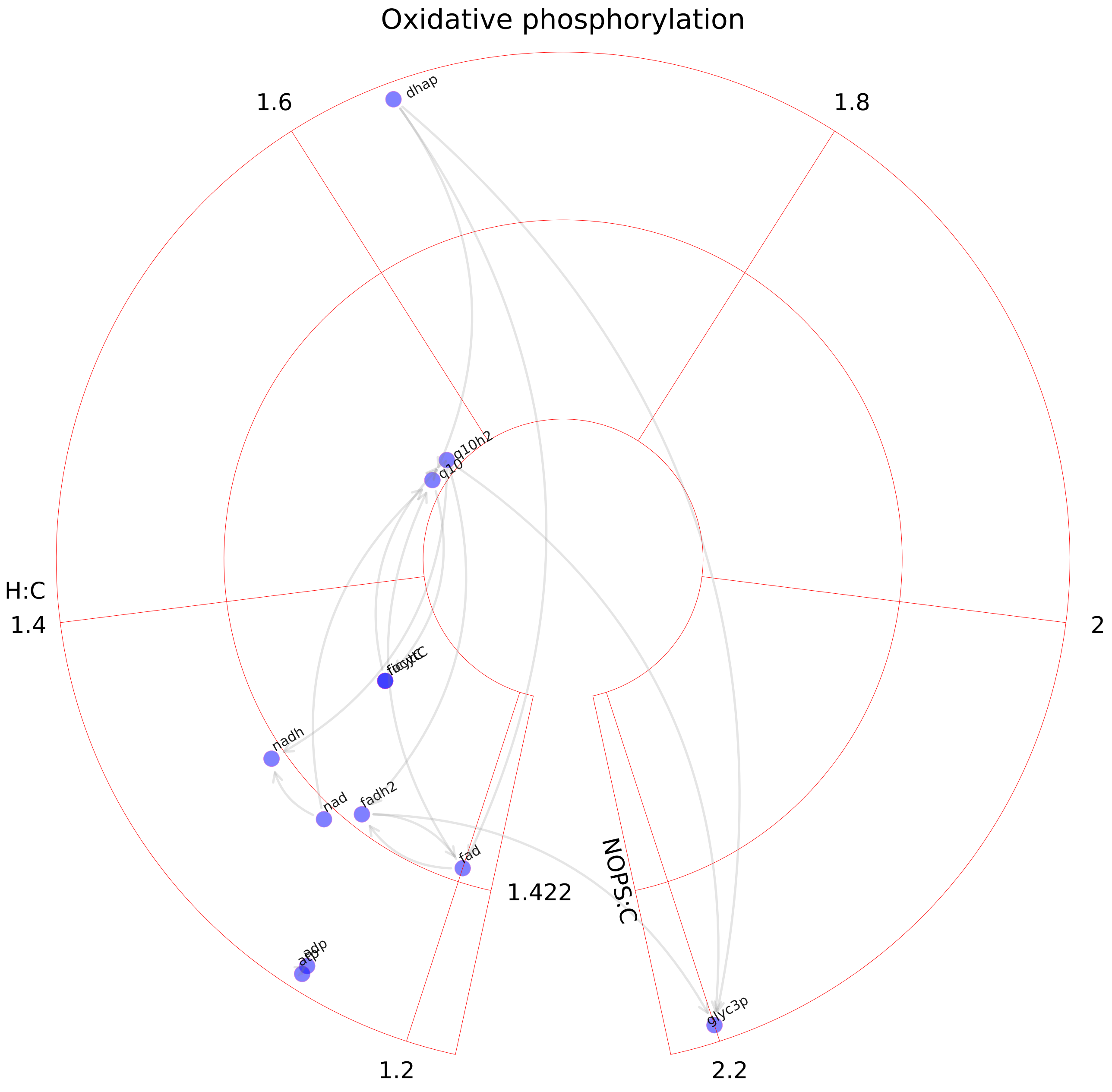

### Pentose phosphate pathway.pdf

#### Pentose phosphate pathway

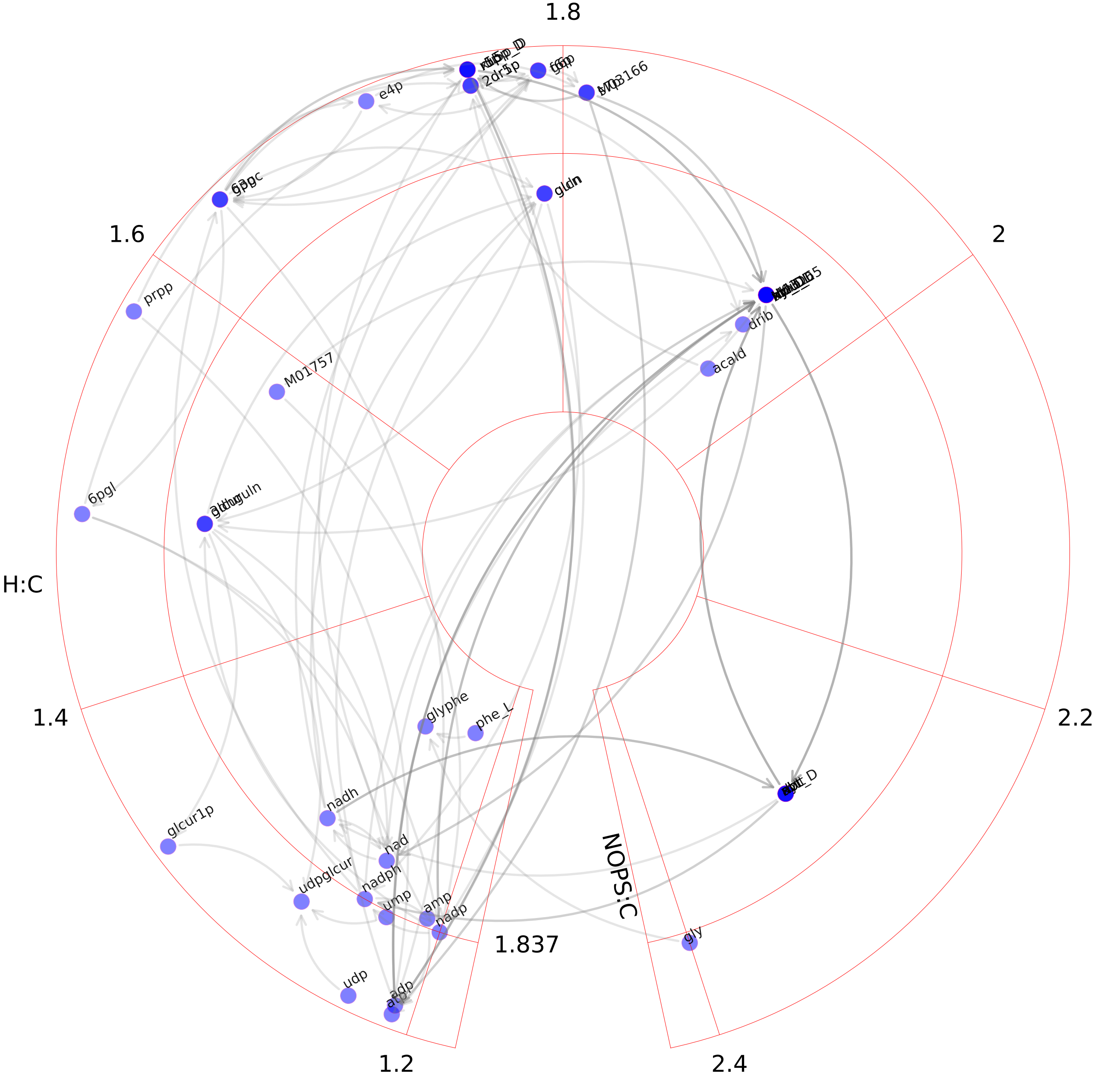

### Peptide metabolism.pdf

#### Peptide metabolism

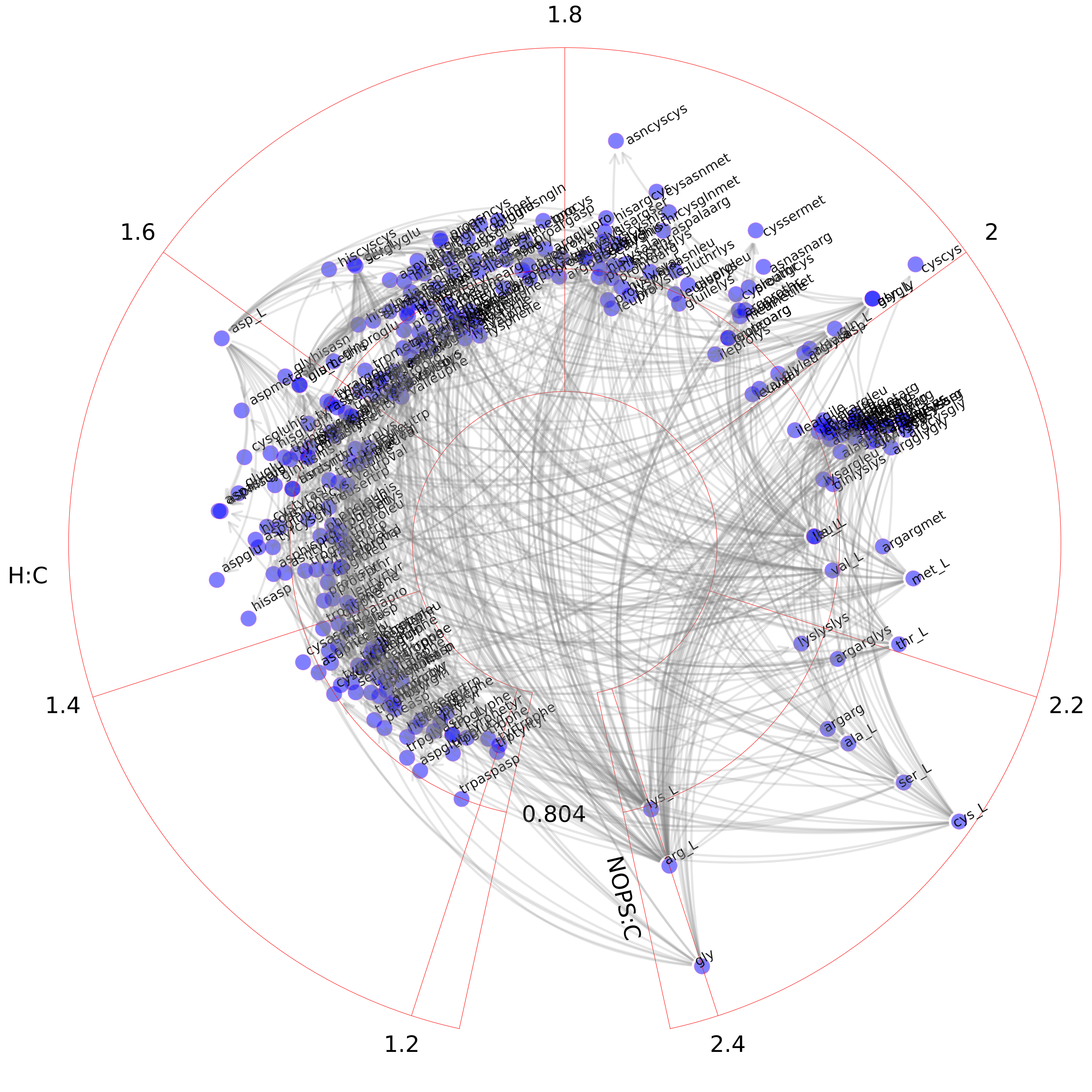

### Phenylalanine metabolism.pdf

Phenylalanine metabolism

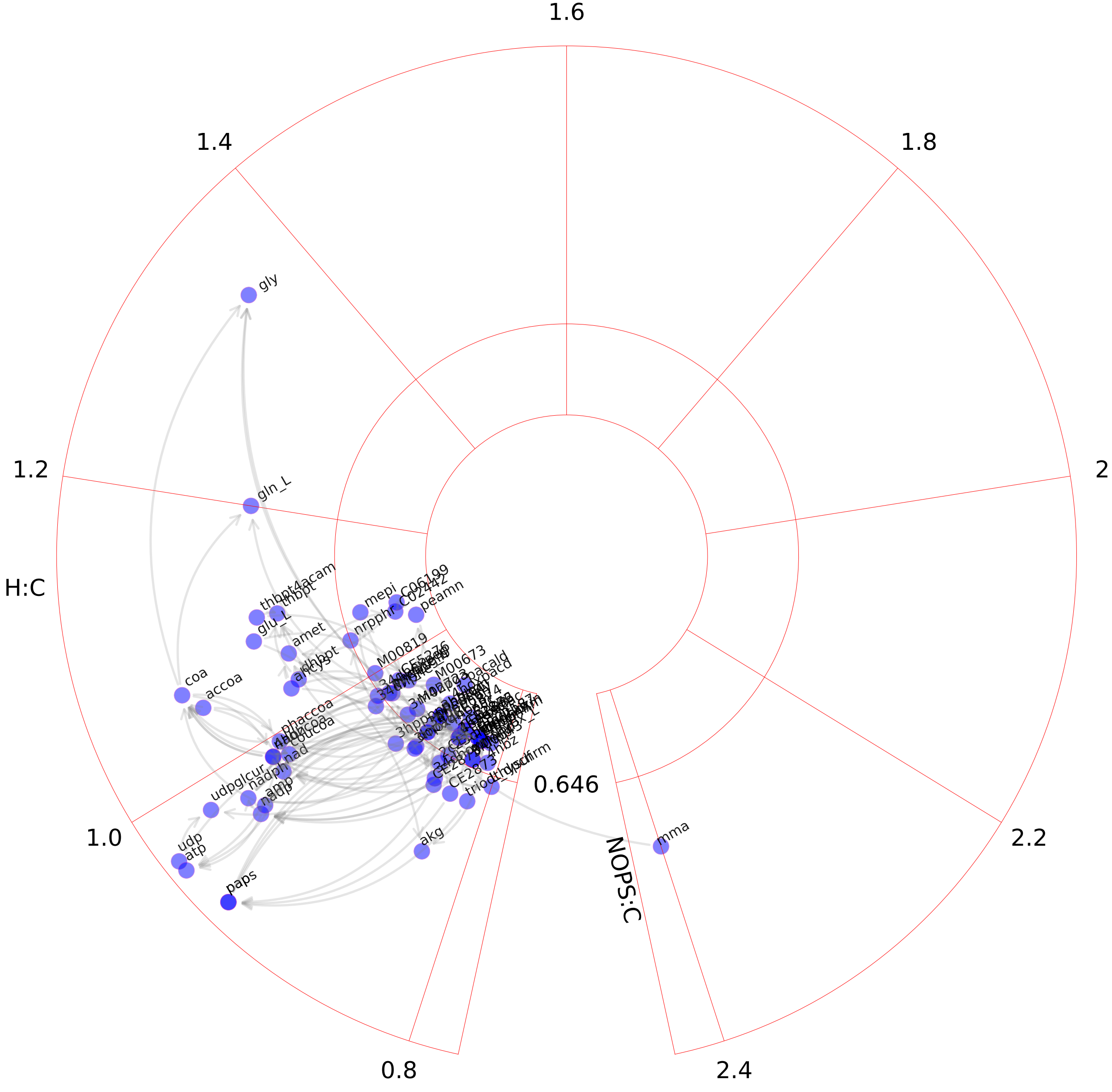

### Phosphatidylinositol phosphate metabolism.pdf

#### Phosphatidylinositol phosphate metabolism

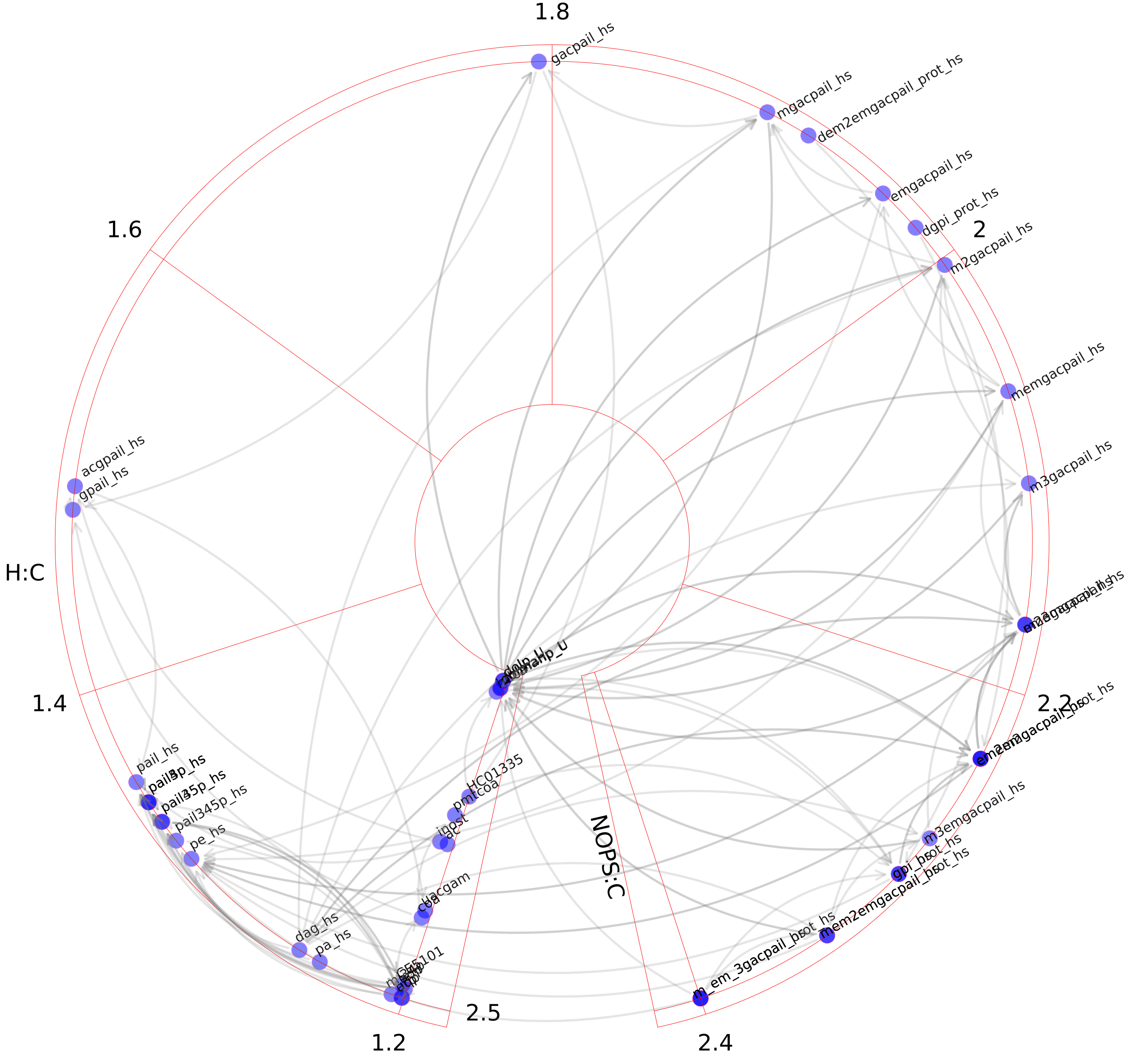

### Propanoate metabolism.pdf

Propanoate metabolism

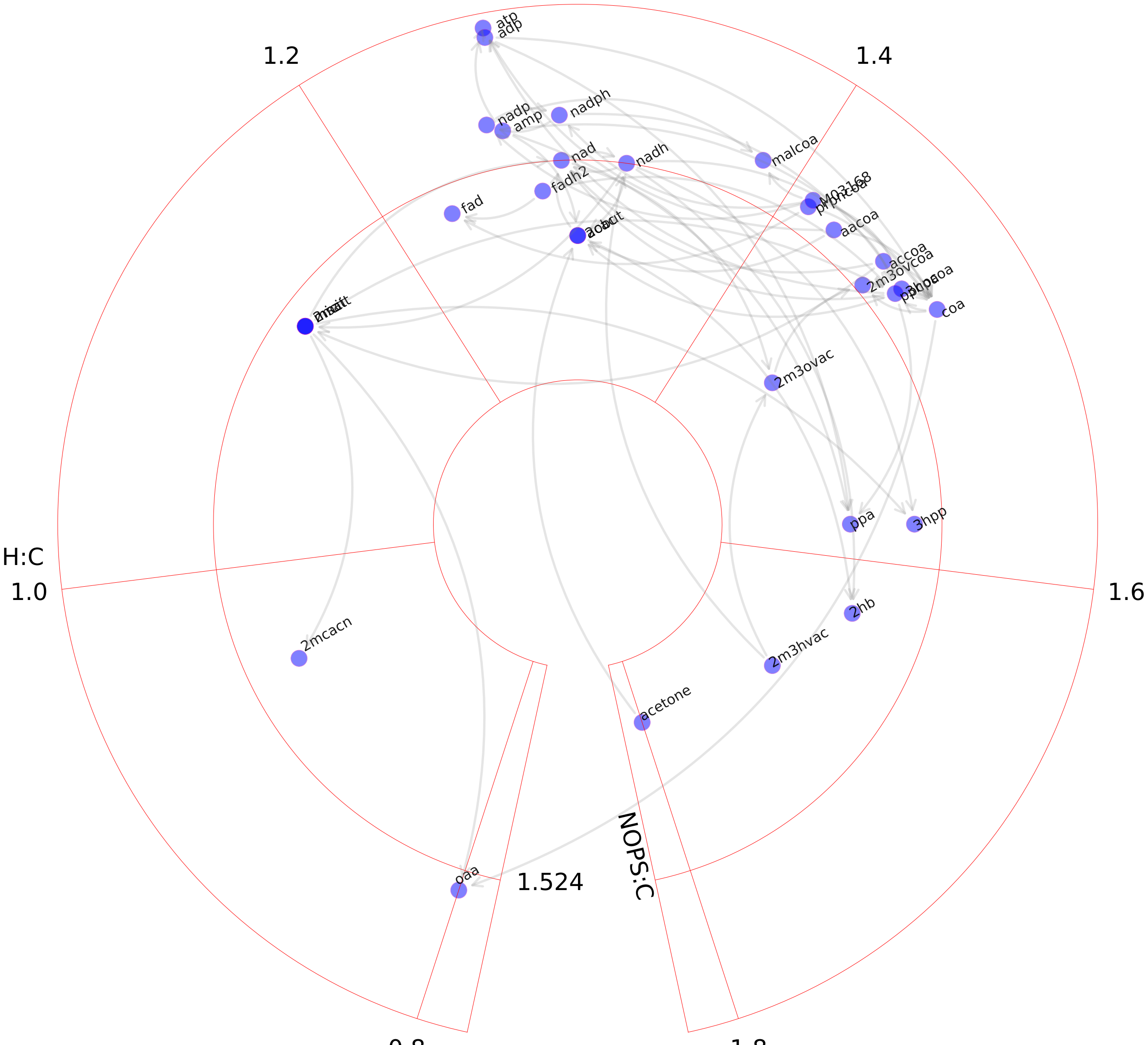

### Purine catabolism.pdf

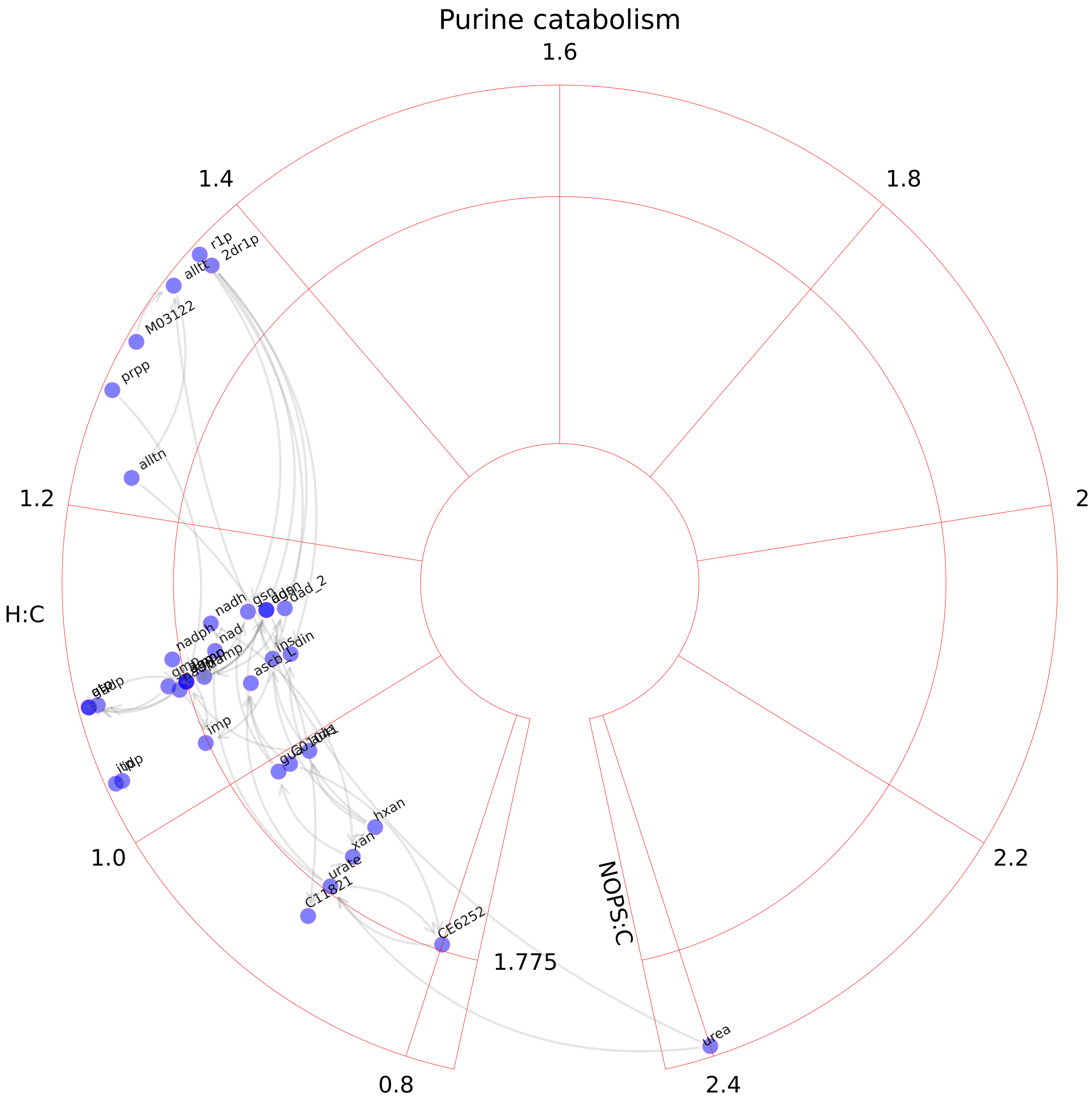

### Purine synthesis.pdf

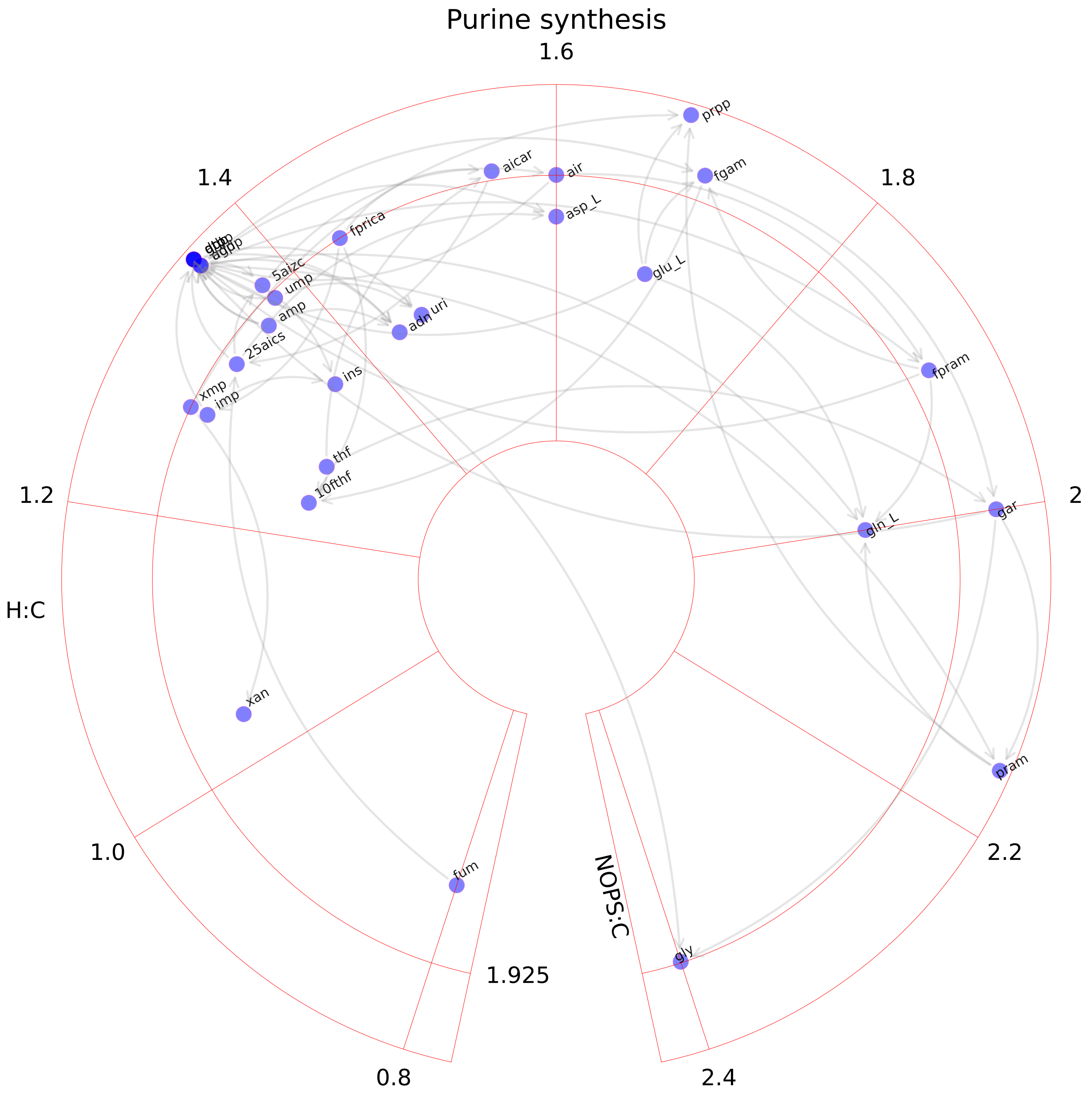

### Pyrimidine catabolism.pdf

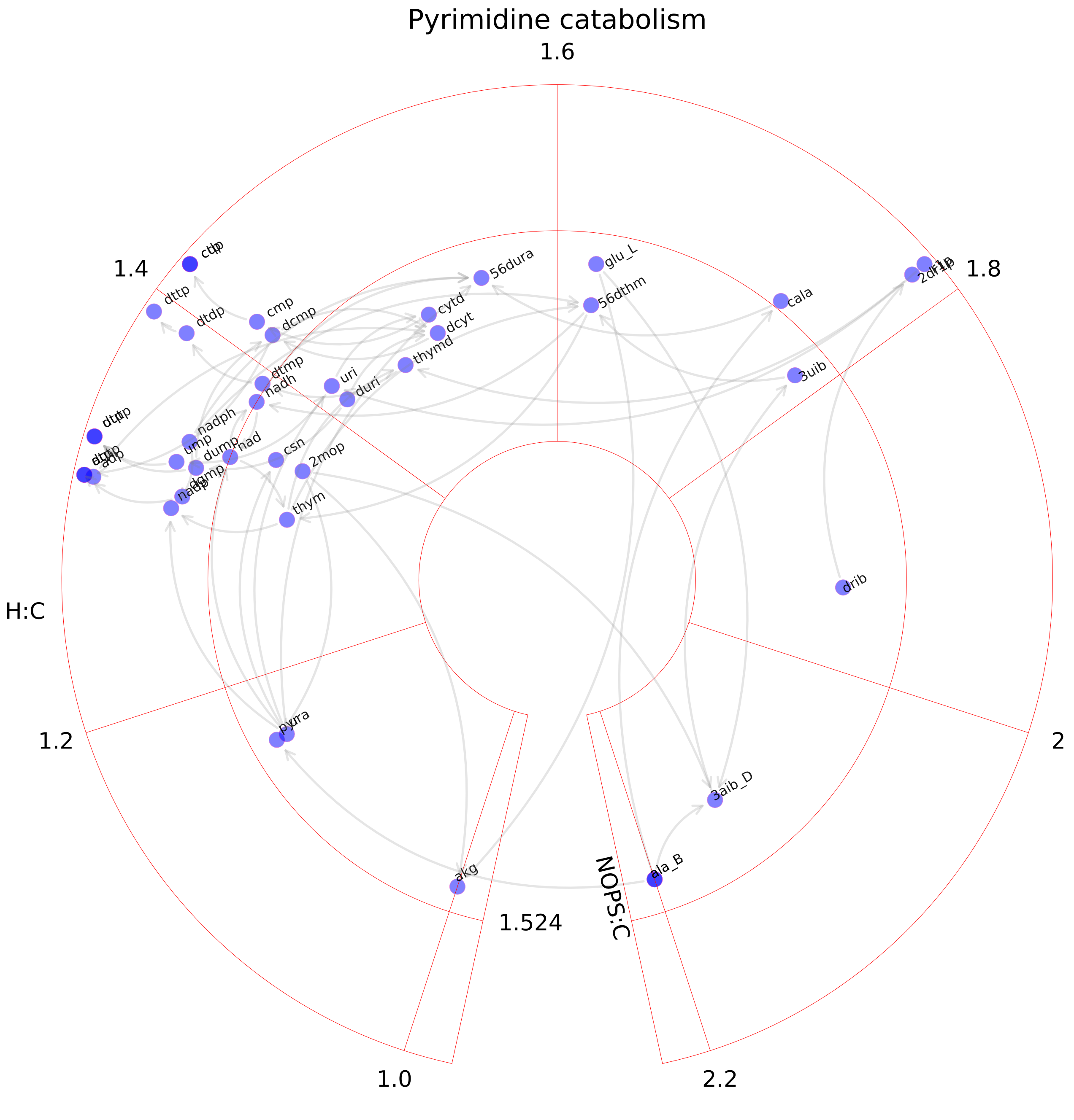

### Pyrimidine synthesis.pdf

Pyrimidine synthesis

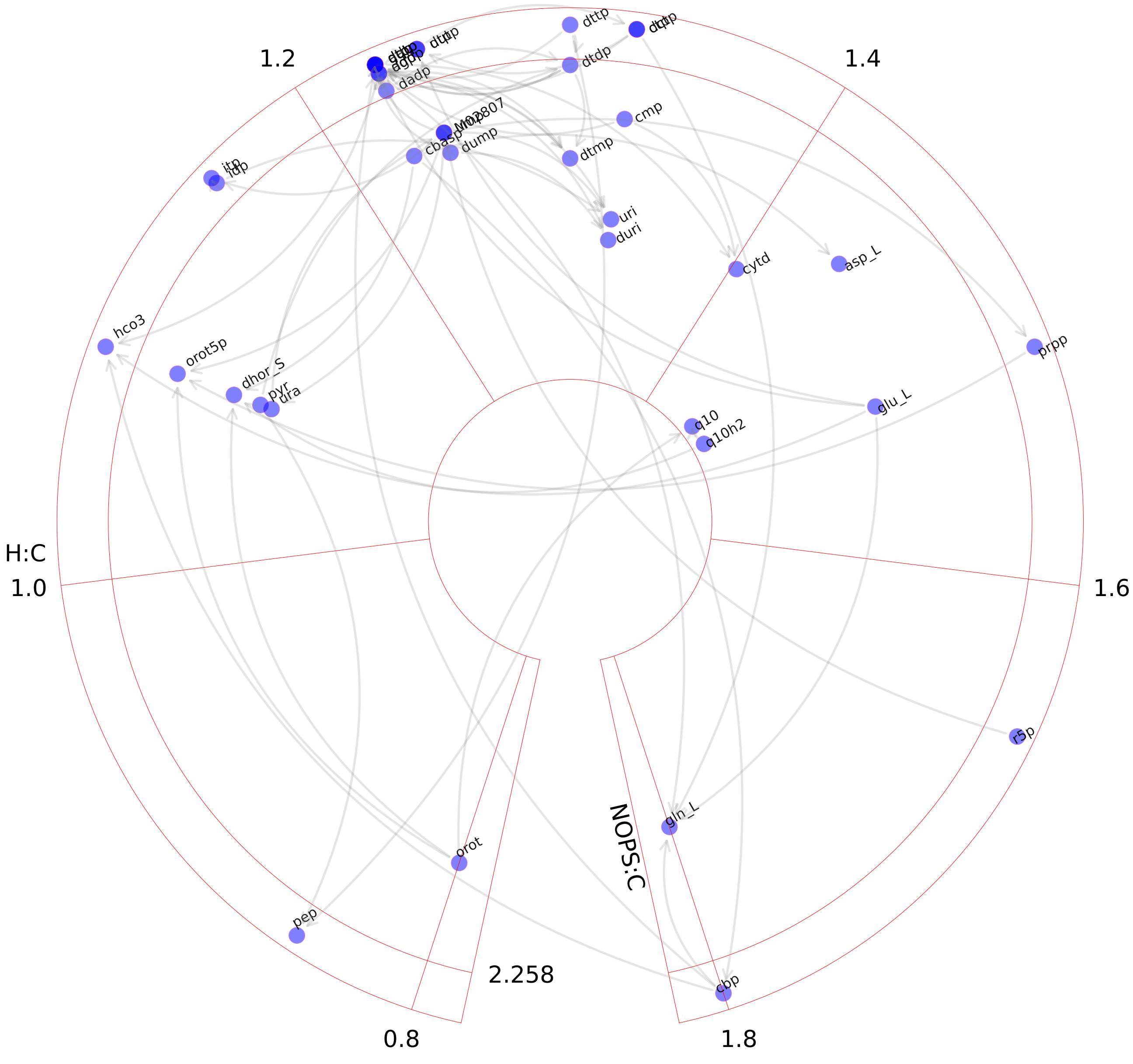

### Pyruvate metabolism.pdf

Pyruvate metabolism

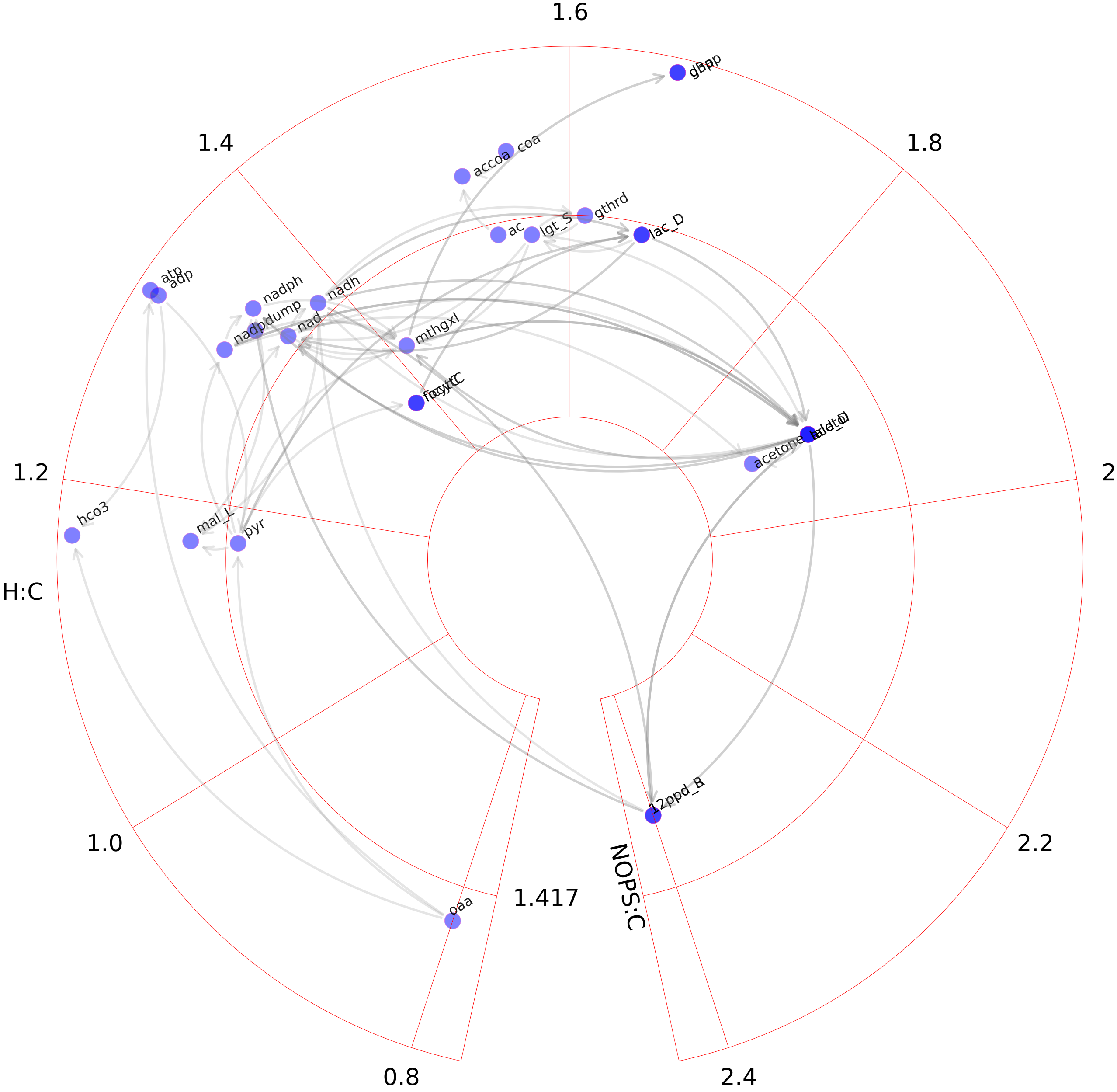

### R group synthesis.pdf

#### R group synthesis

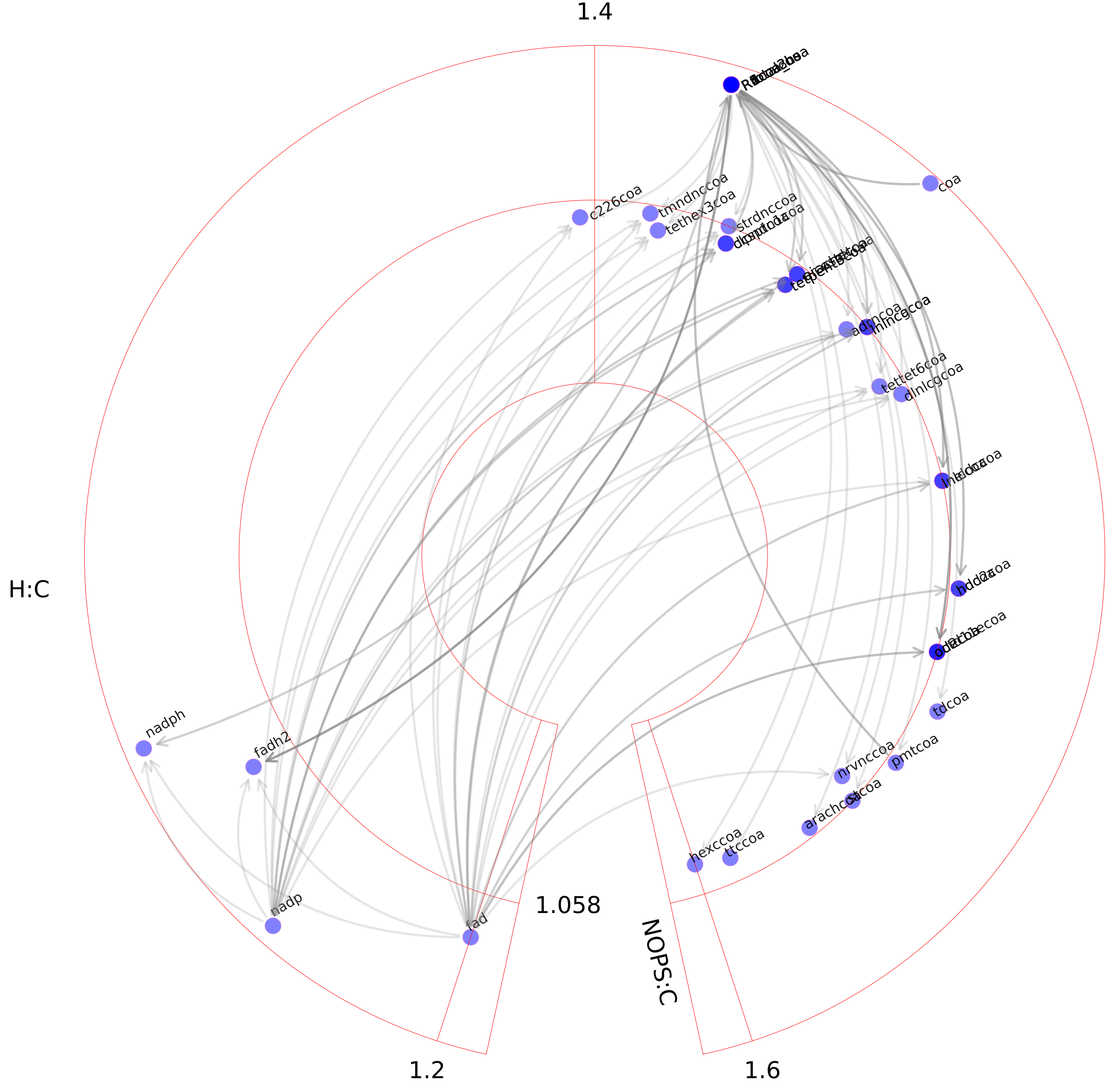

### ROS detoxification.pdf

ROS detoxification

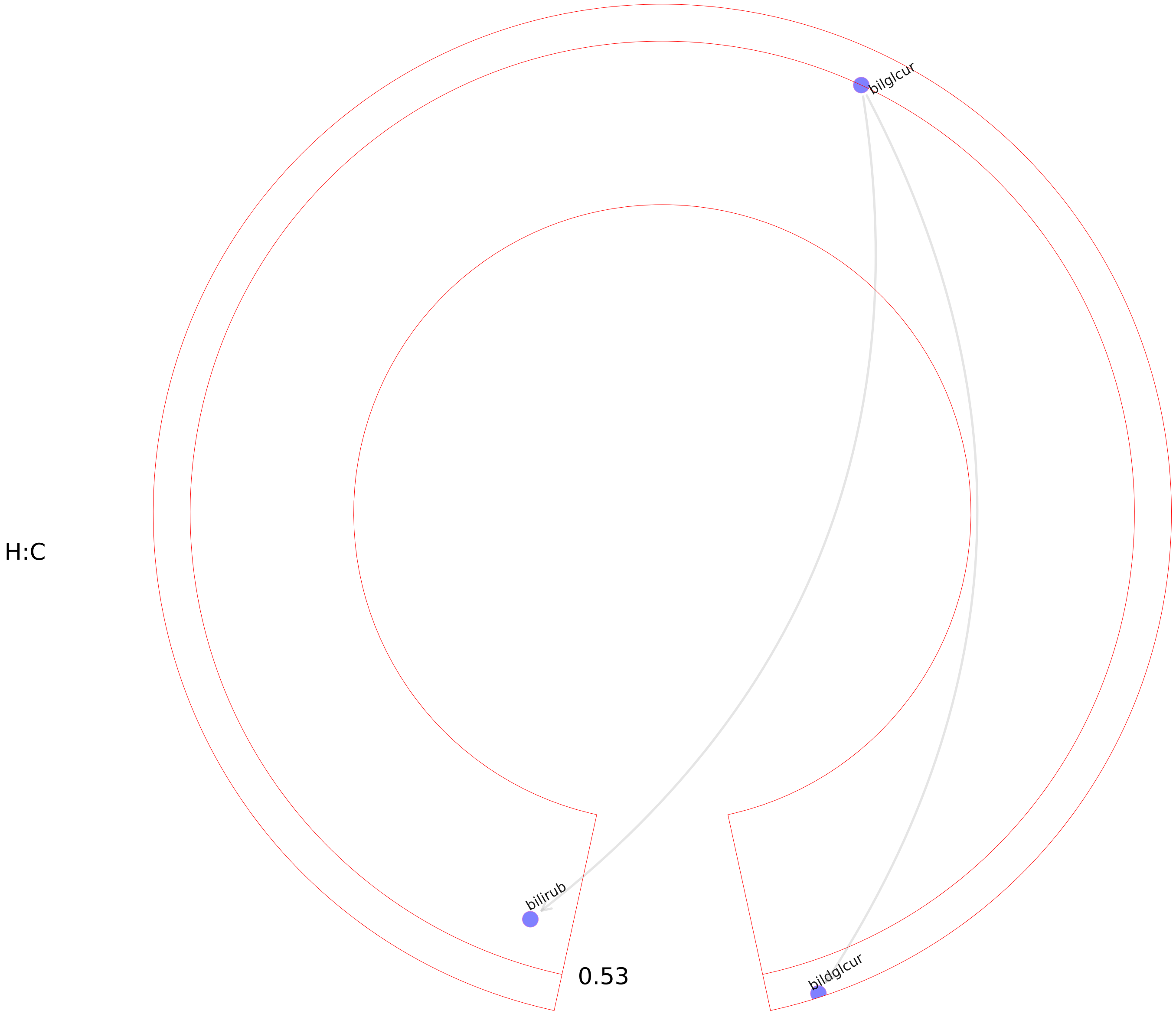

### Selenoamino acid metabolism.pdf

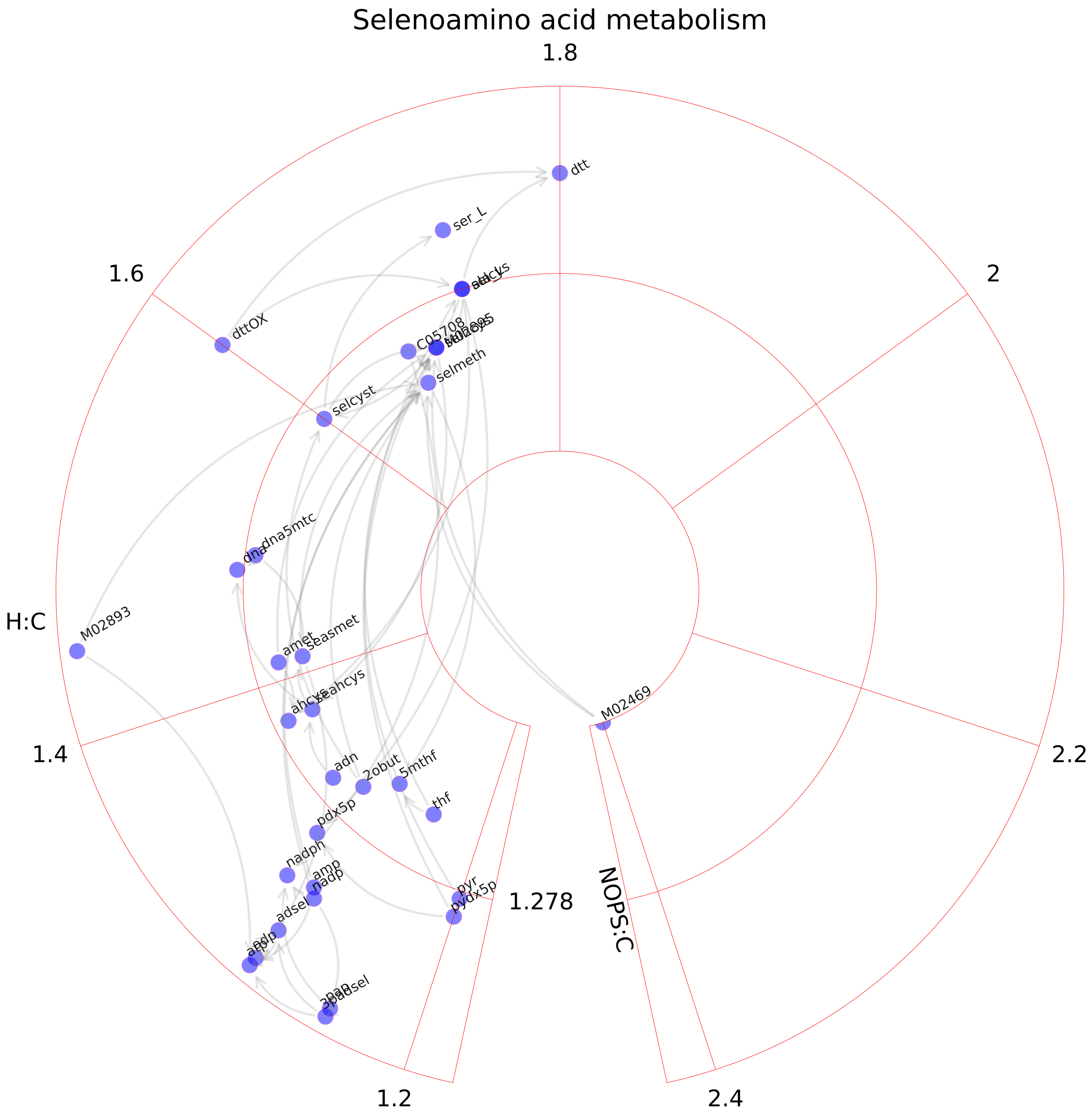

### Sphingolipid metabolism.pdf

#### Sphingolipid metabolism

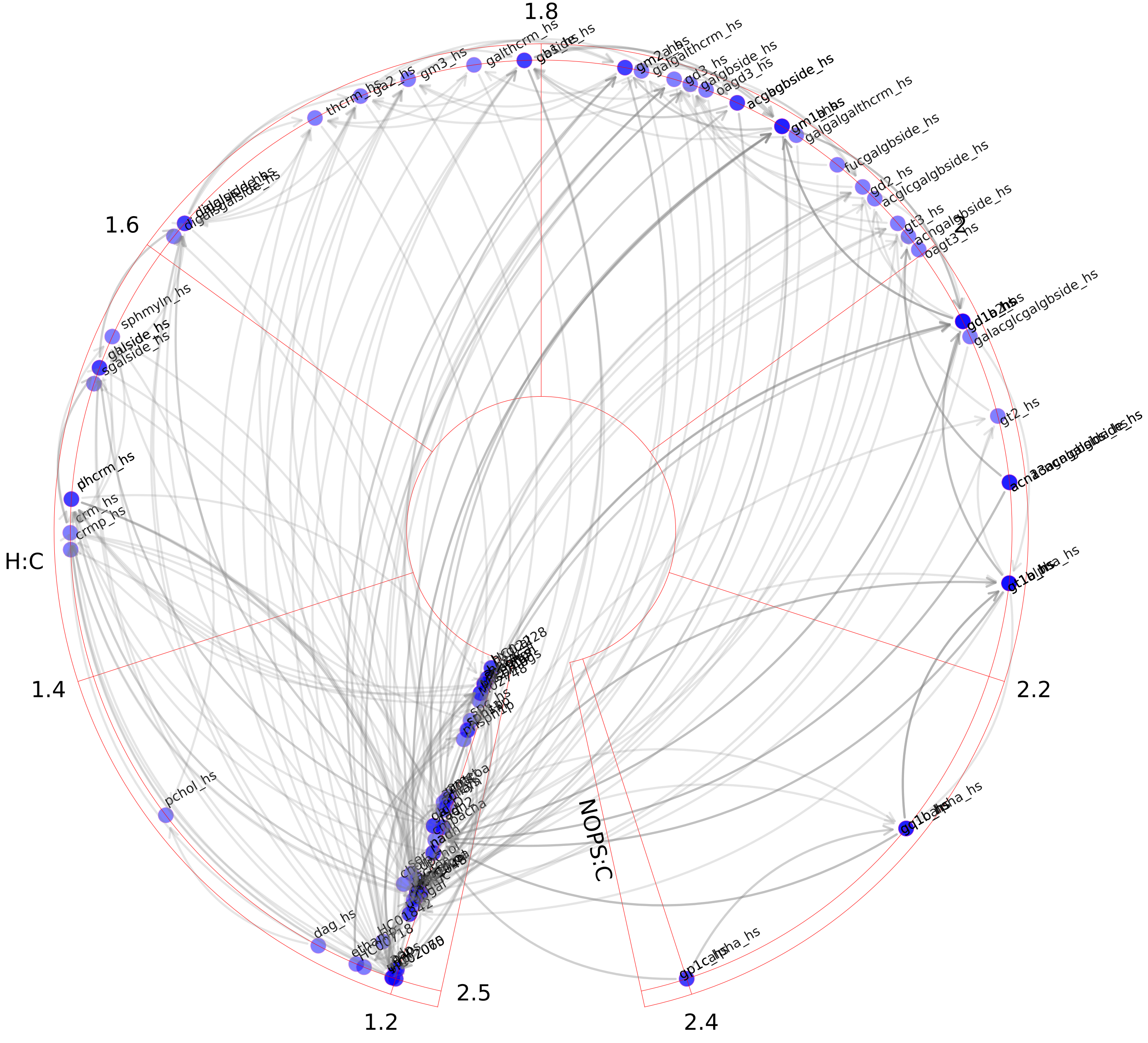

### Squalene and cholesterol synthesis.pdf

#### Squalene and cholesterol synthesis

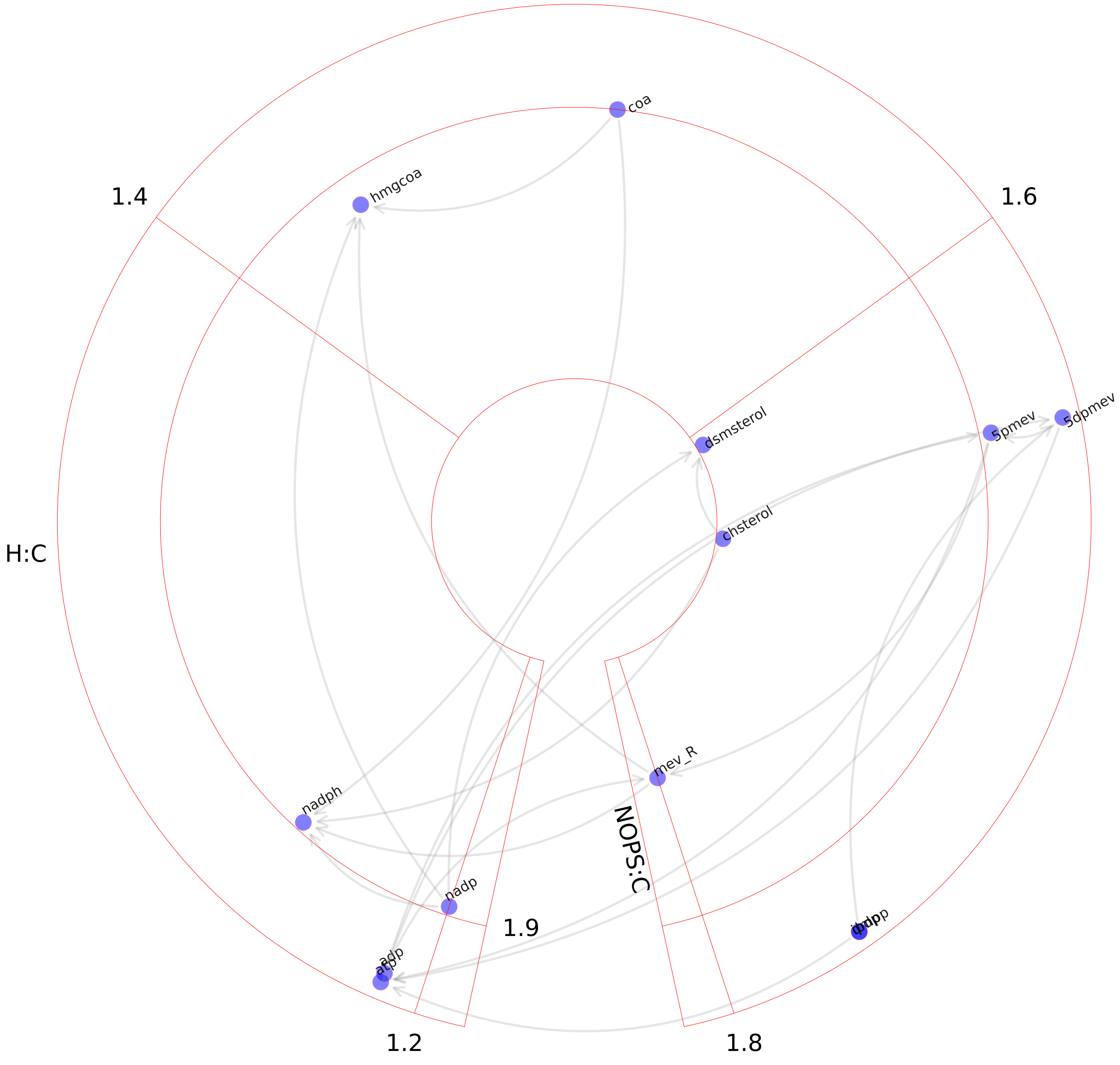

### Starch and sucrose metabolism.pdf

#### Starch and sucrose metabolism

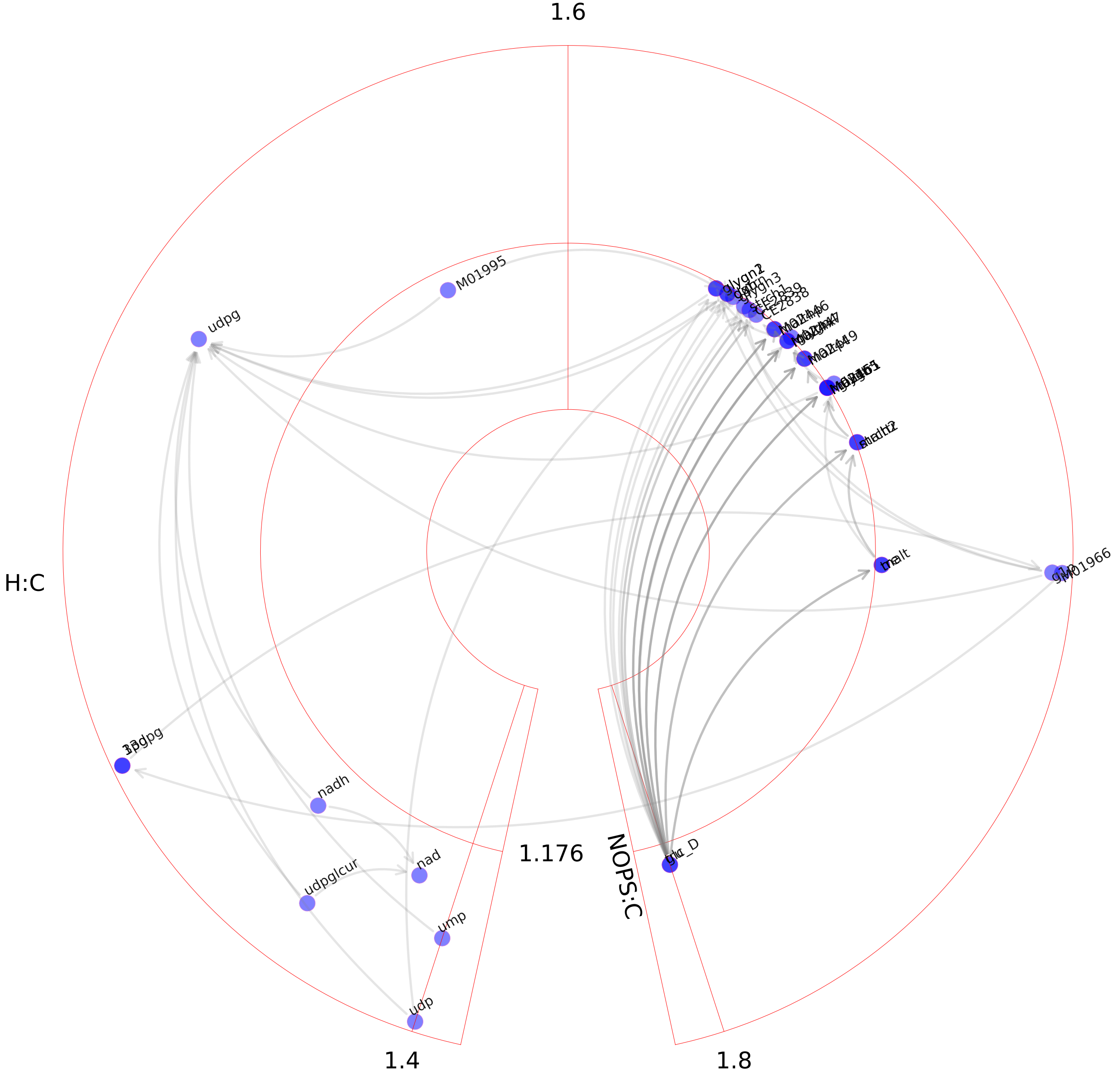

### Steroid metabolism.pdf

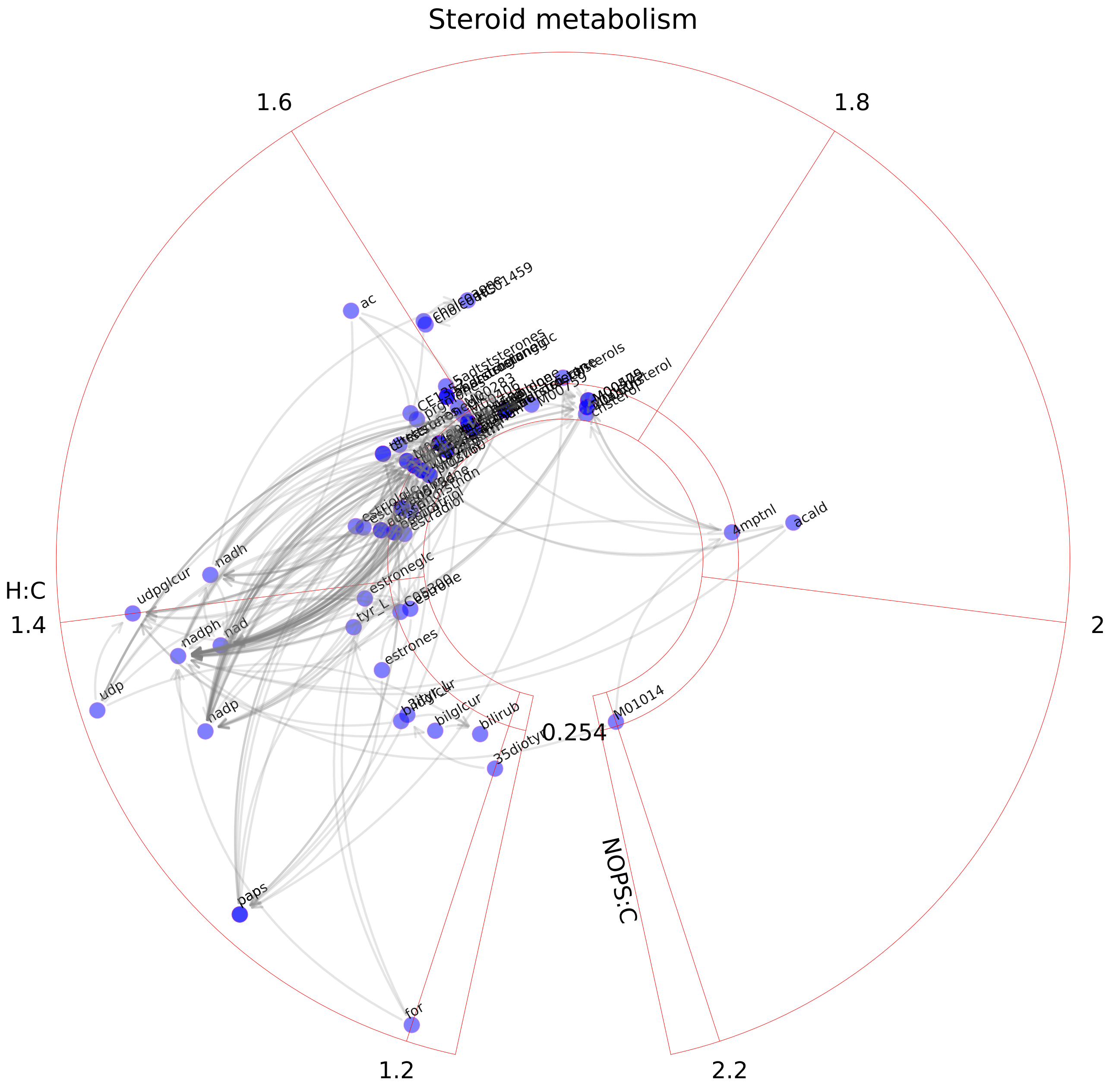

### Stilbene, coumarine and lignin synthesis.pdf

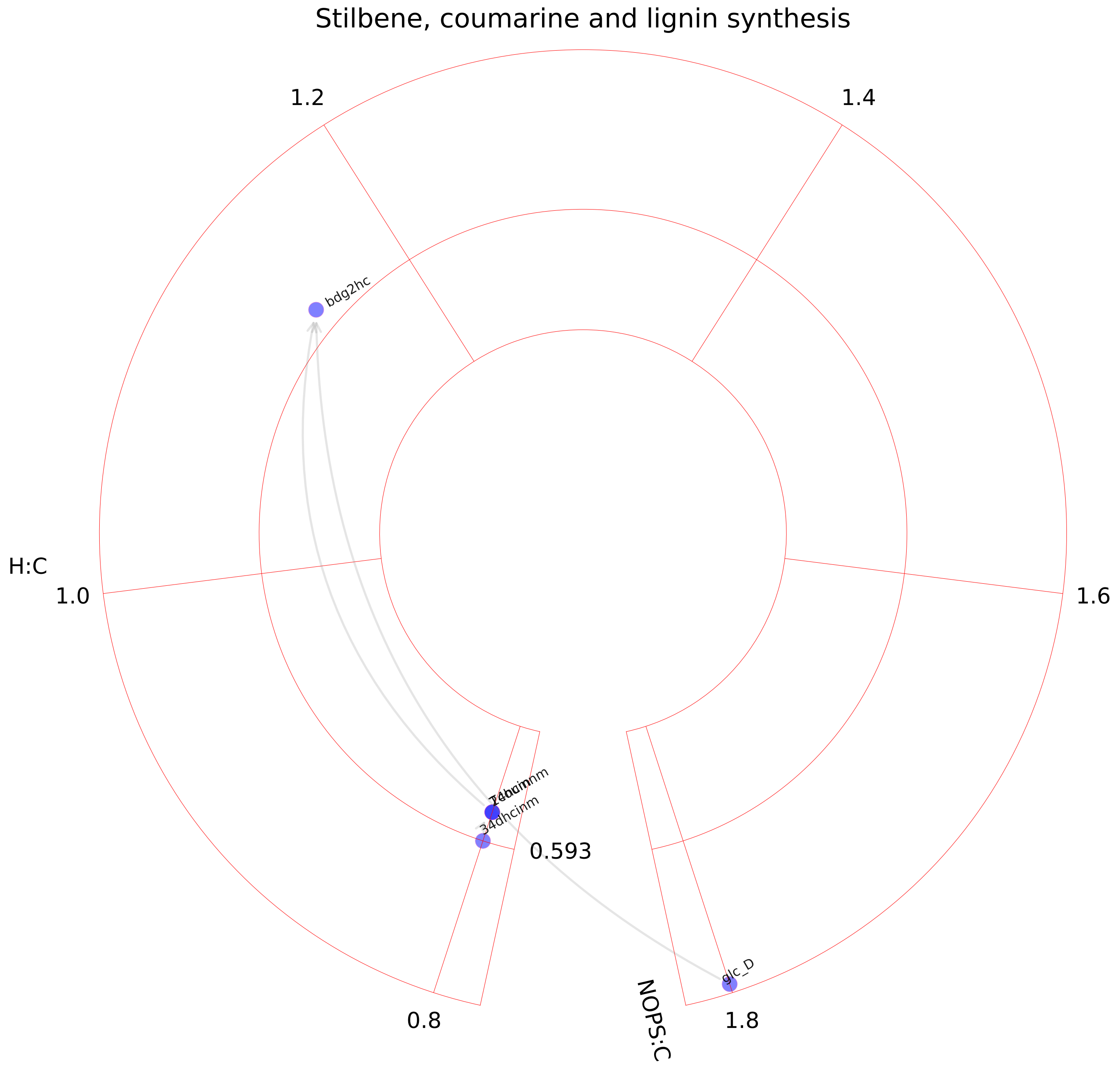

### Taurine and hypotaurine metabolism.pdf

#### Taurine and hypotaurine metabolism

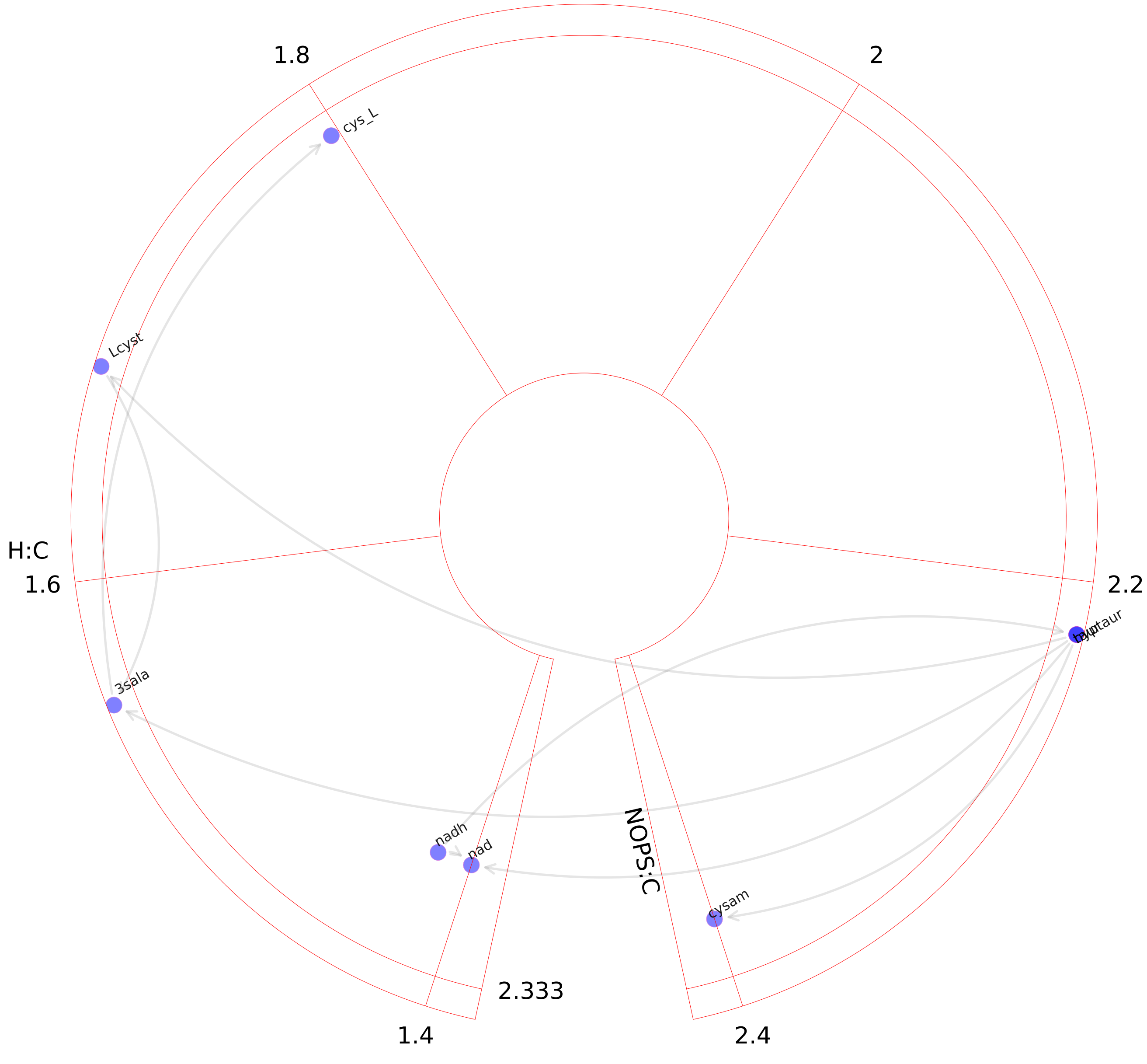

### Tetrahydrobiopterin metabolism.pdf

##### Tetrahydrobiopterin metabolism

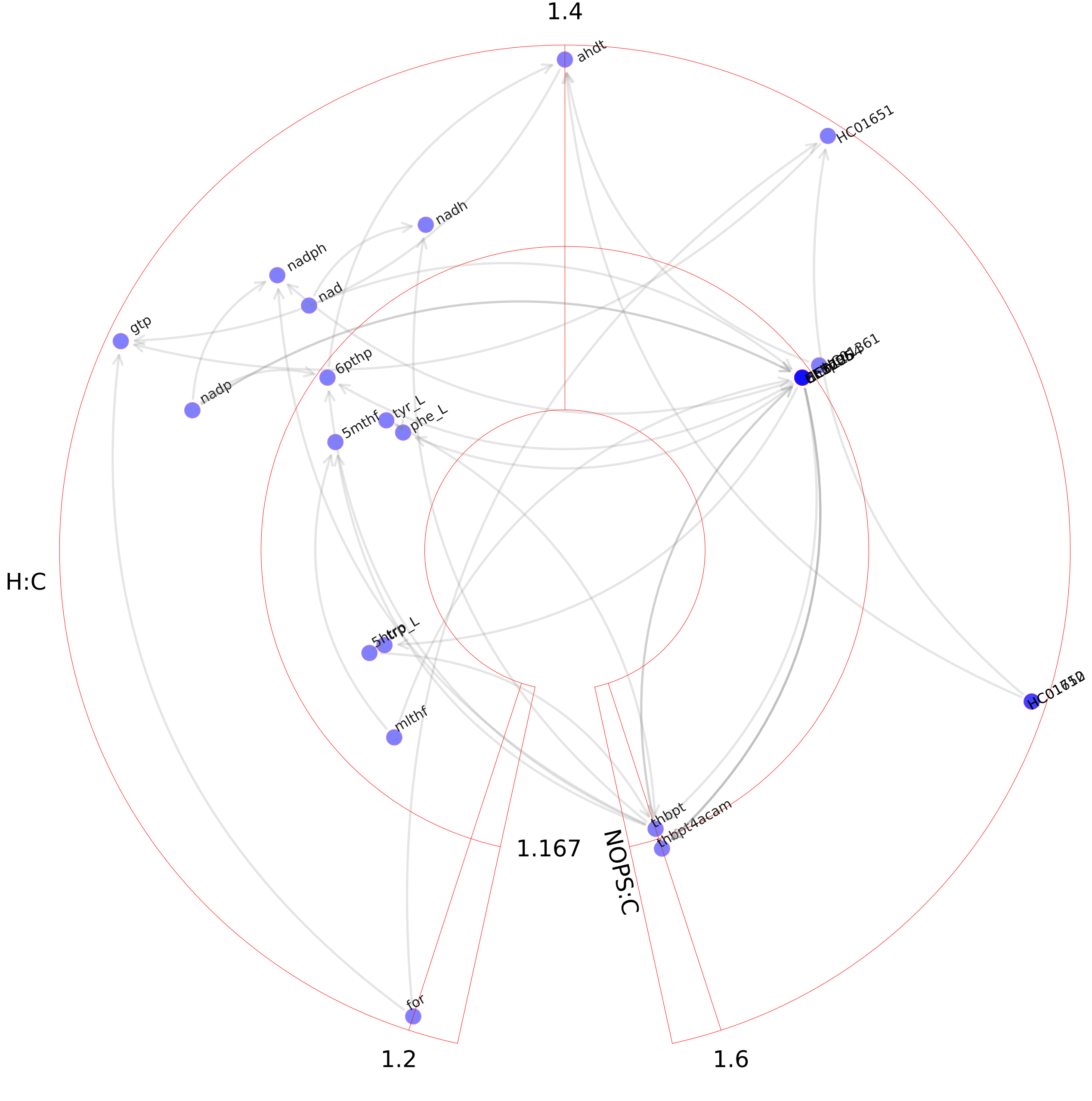

### Thiamine metabolism.pdf

#### Thiamine metabolism

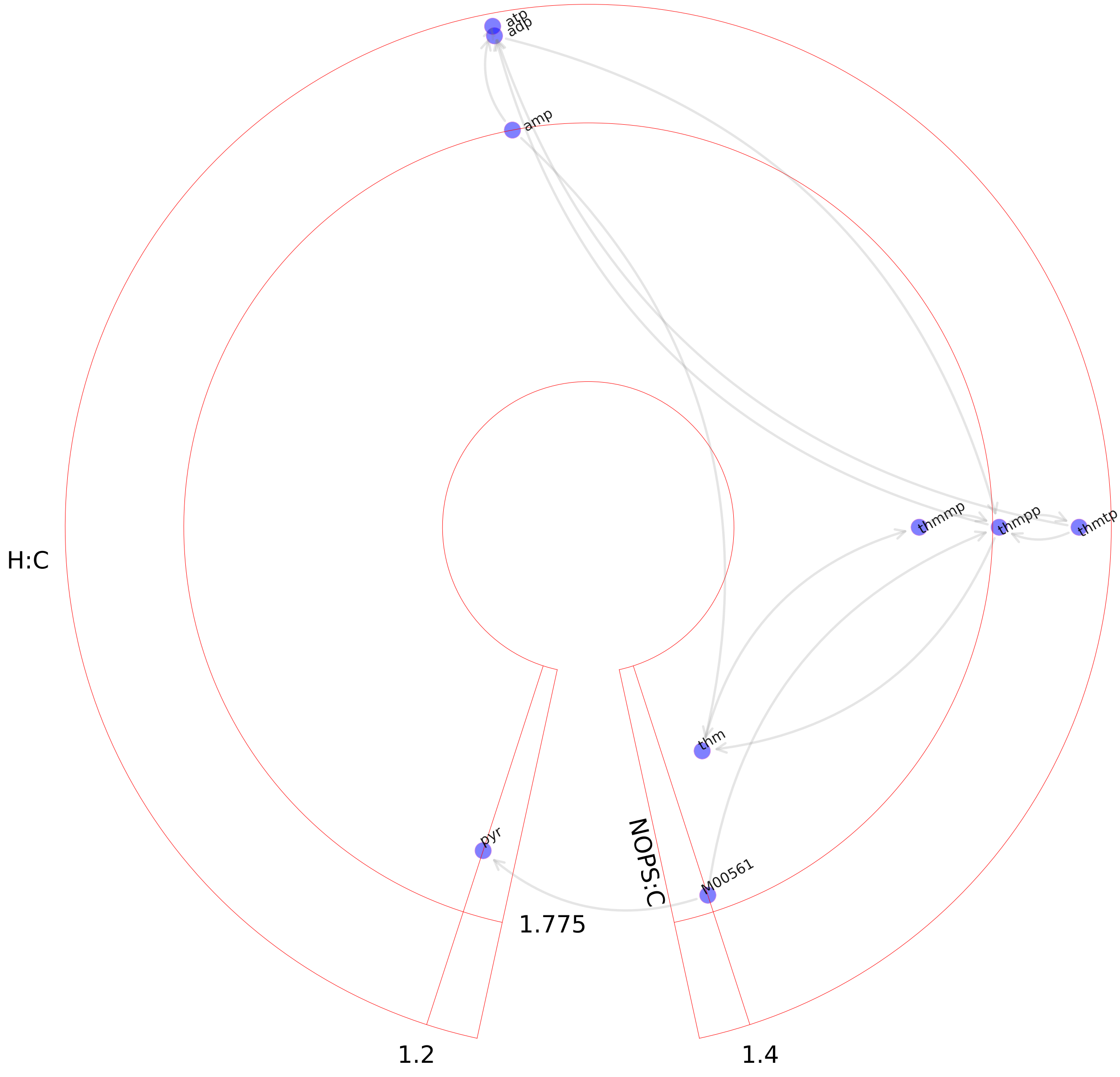

### Triacylglycerol synthesis.pdf

#### Triacylglycerol synthesis

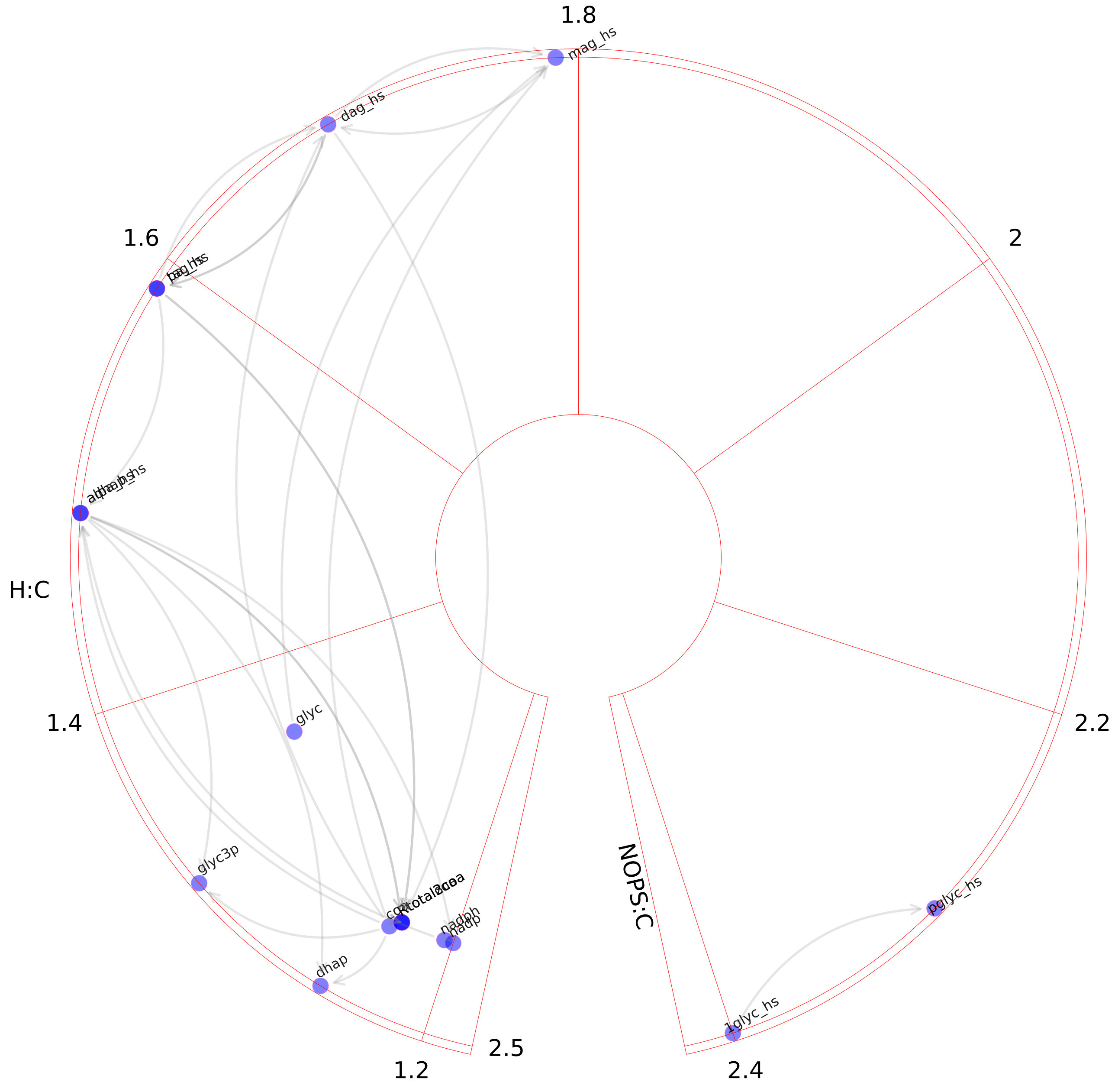

### Triglycerides formation.pdf

#### Triglycerides formation

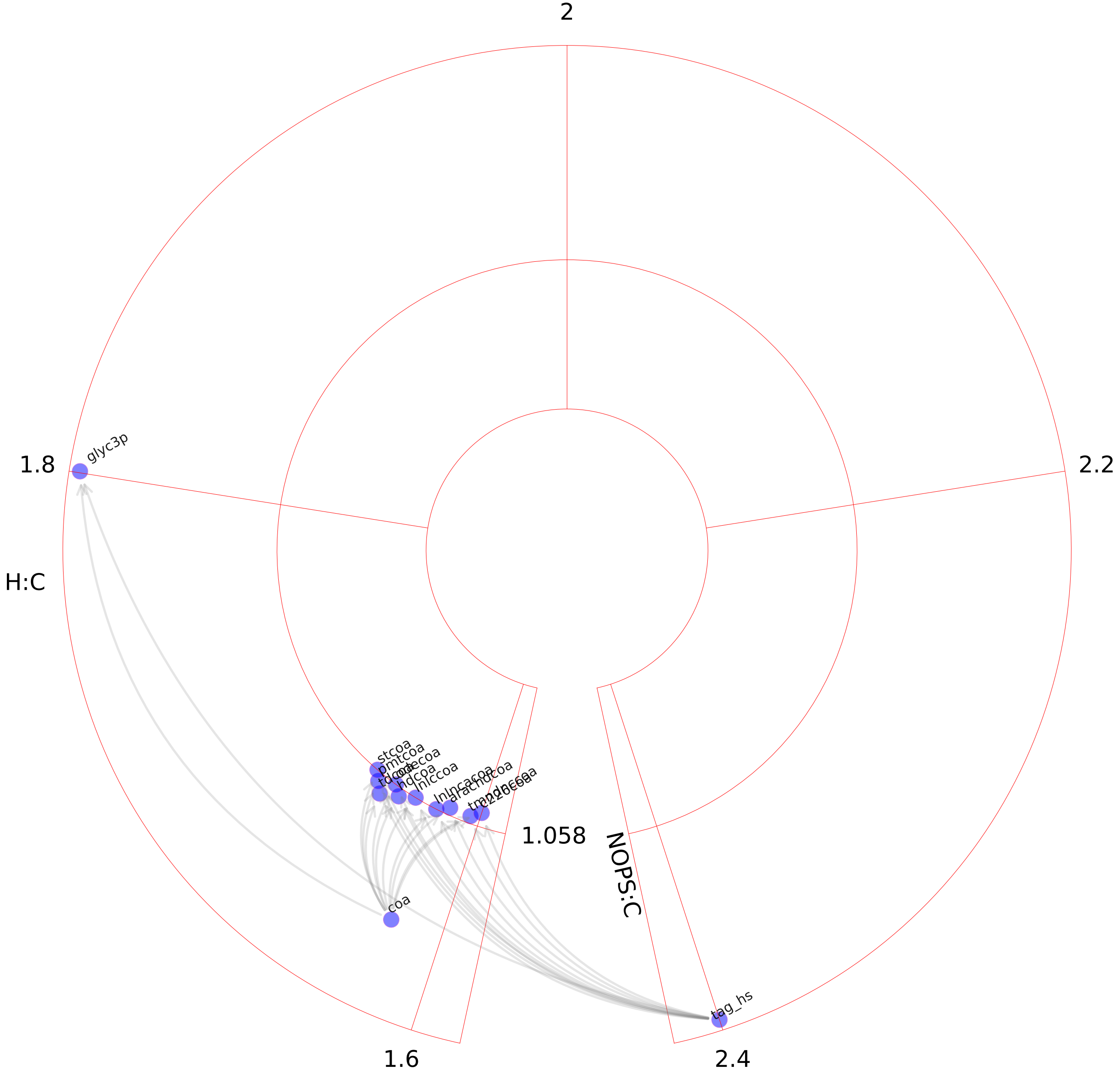

### Tryptophan metabolism.pdf

#### Tryptophan metabolism

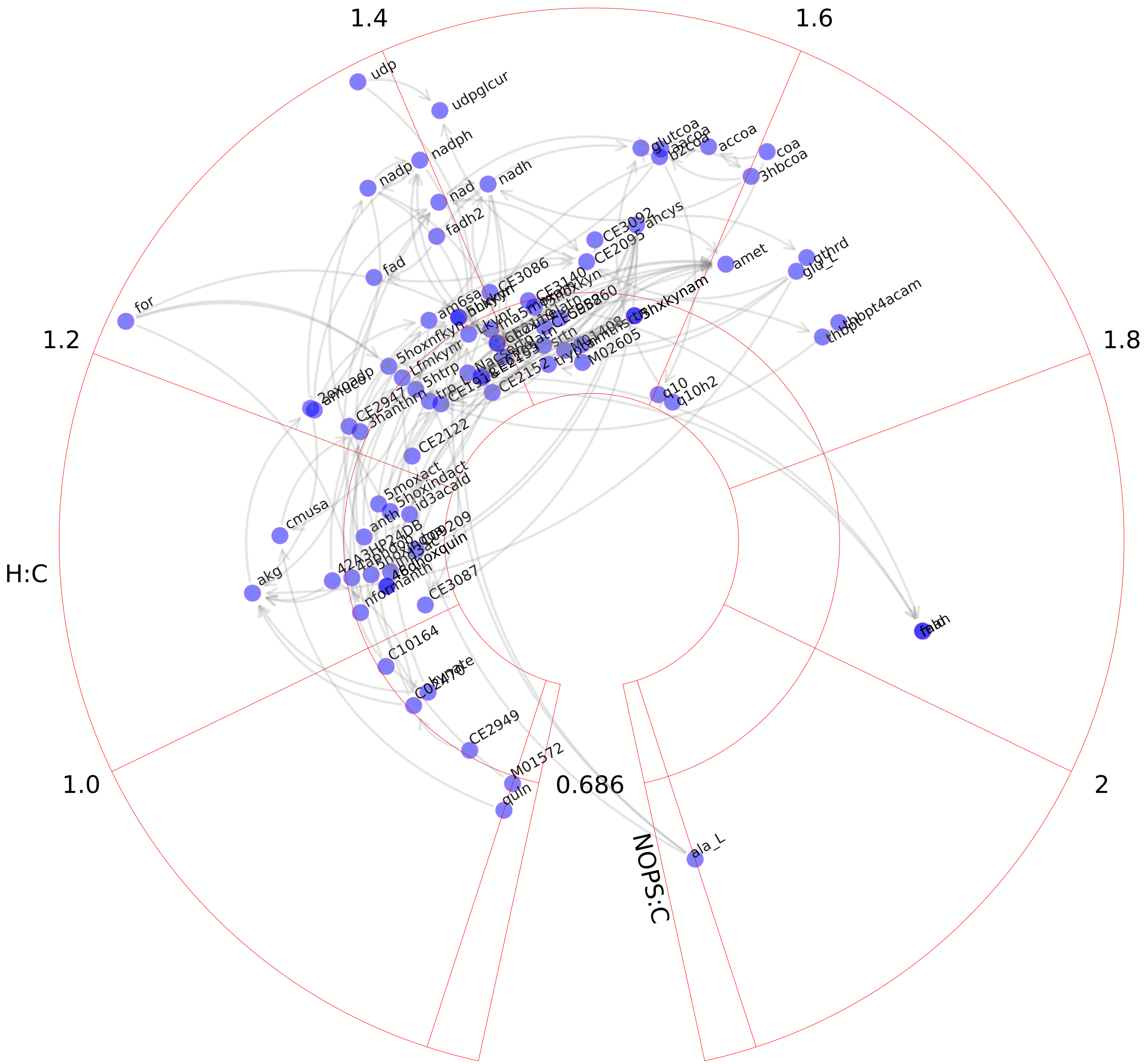

### Tyrosine metabolism.pdf

#### Tyrosine metabolism

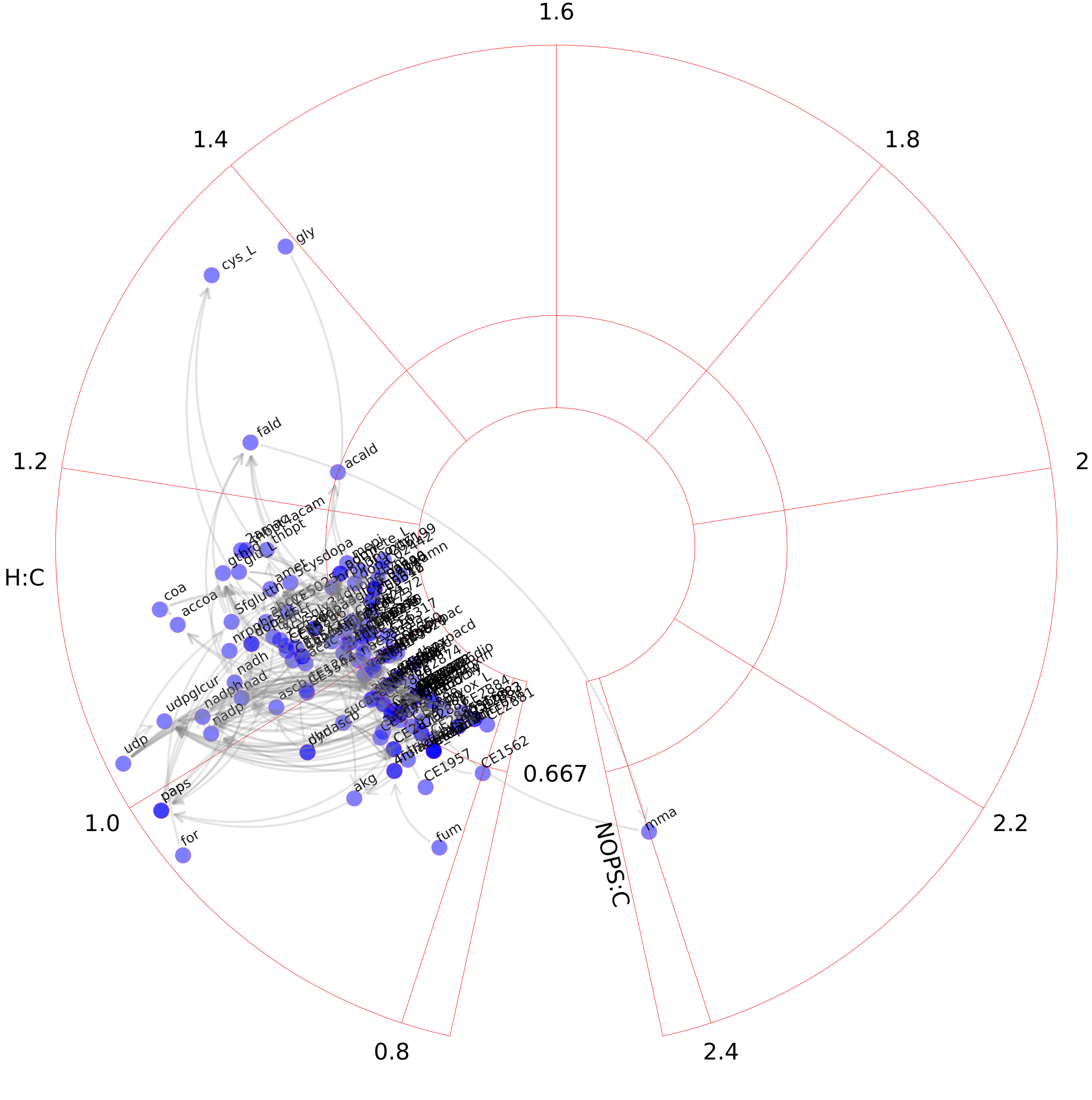

### Ubiquinone synthesis.pdf

#### Ubiquinone synthesis

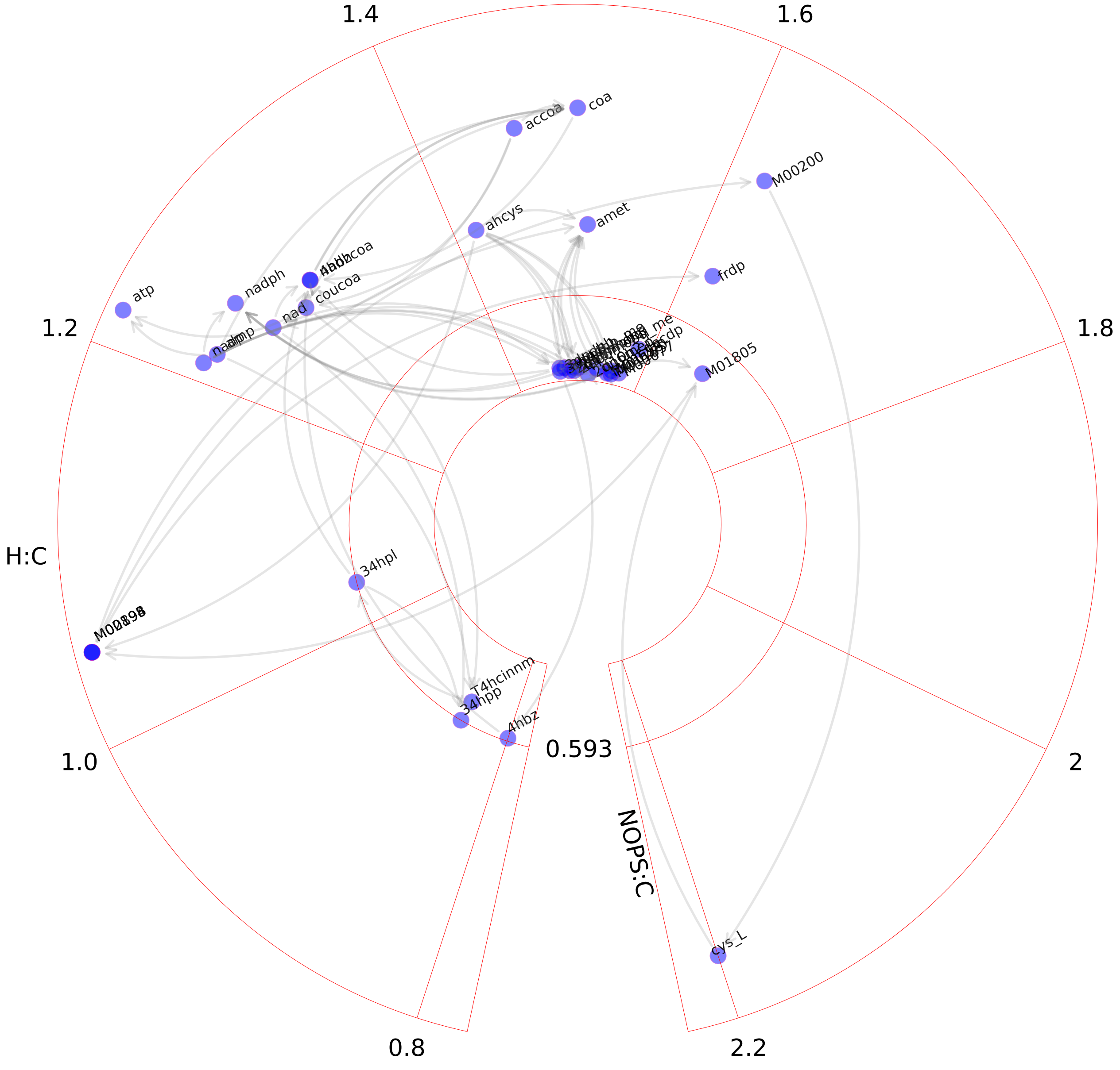

### Urea cycle.pdf

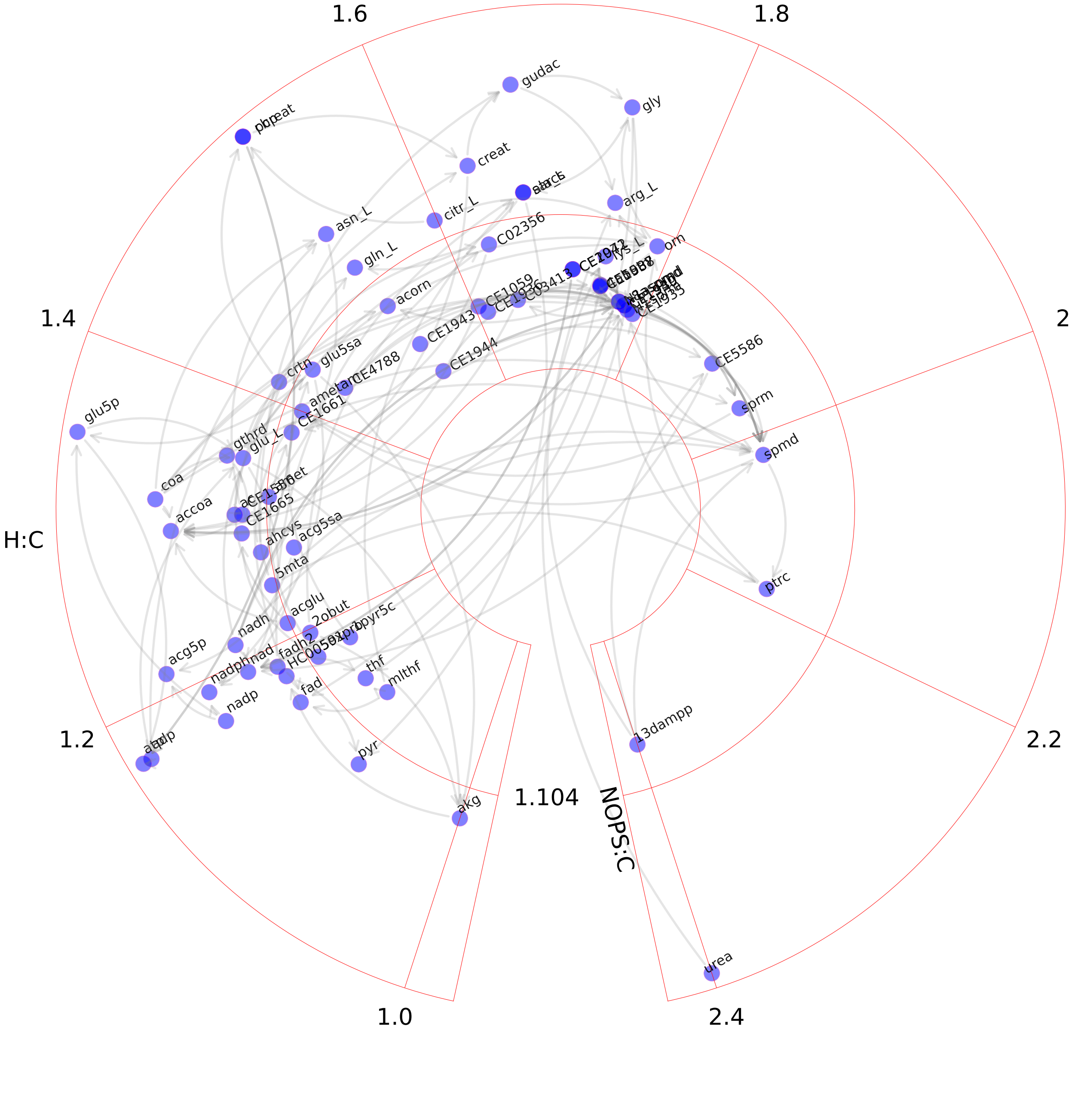

### Valine, leucine, and isoleucine metabolism.pdf

#### Valine, leucine, and isoleucine metabolism

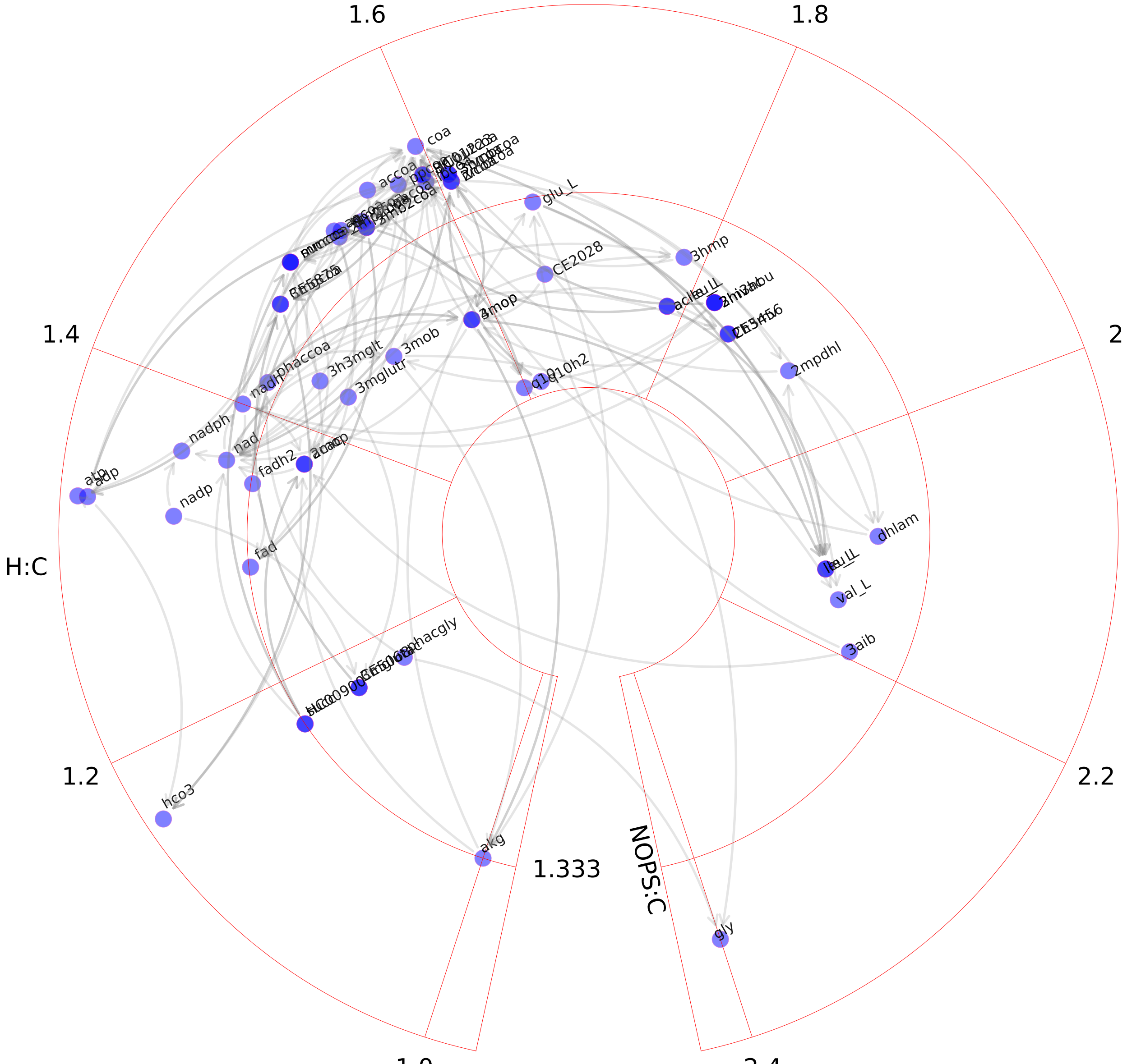

### Vitamin A metabolism.pdf

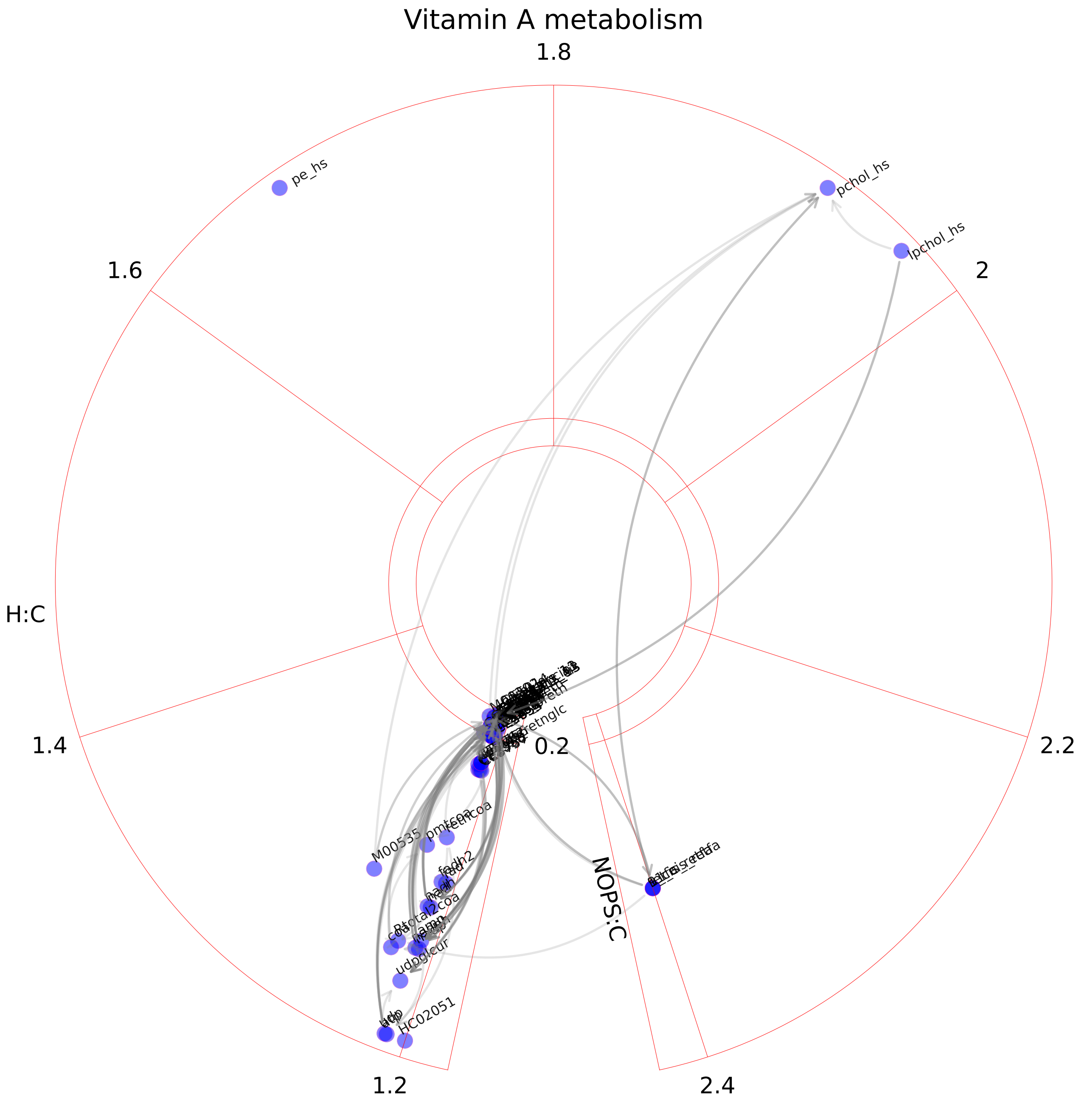

### Vitamin B2 metabolism.pdf

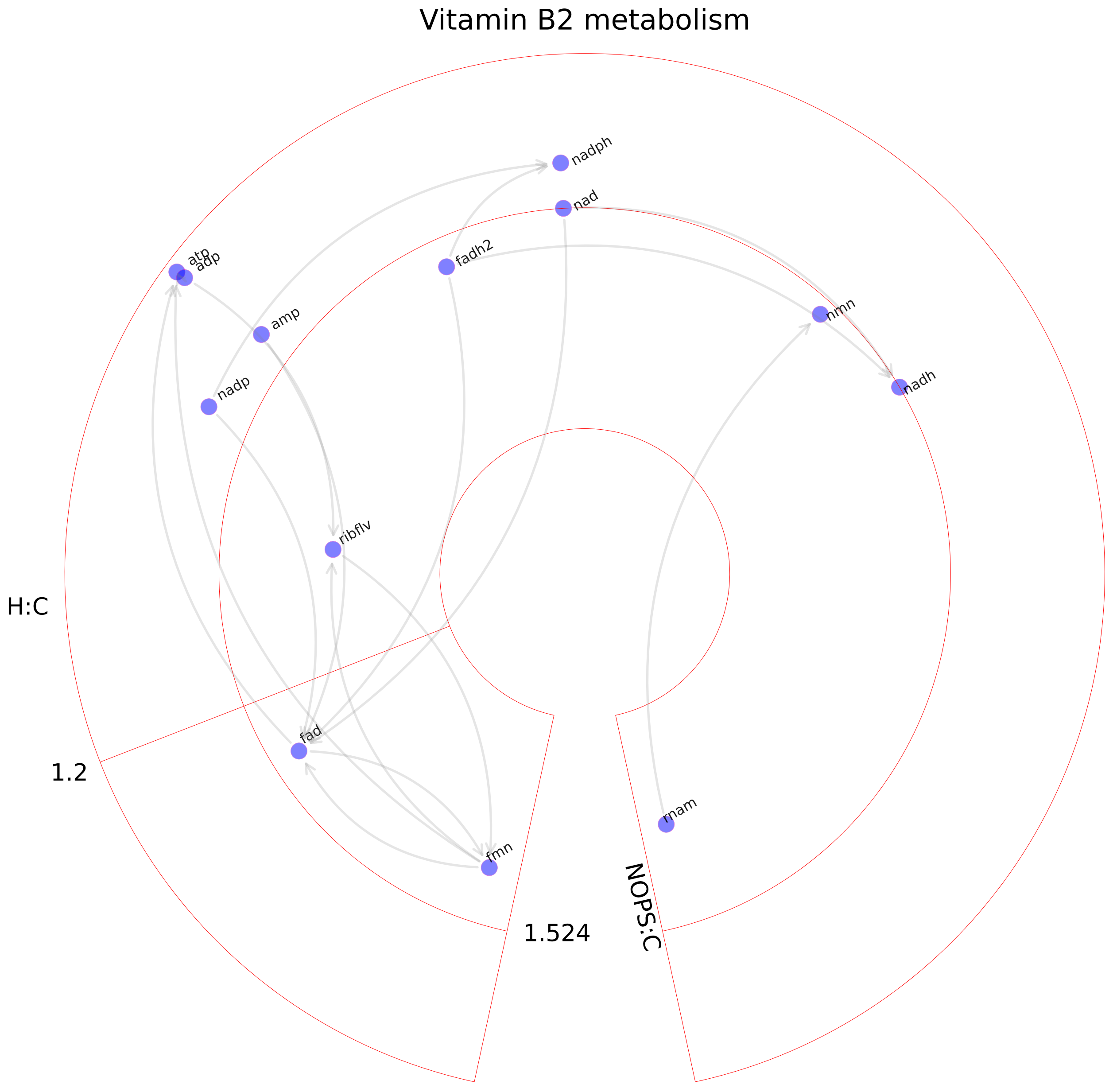

### Vitamin B6 metabolism.pdf

#### Vitamin B6 metabolism

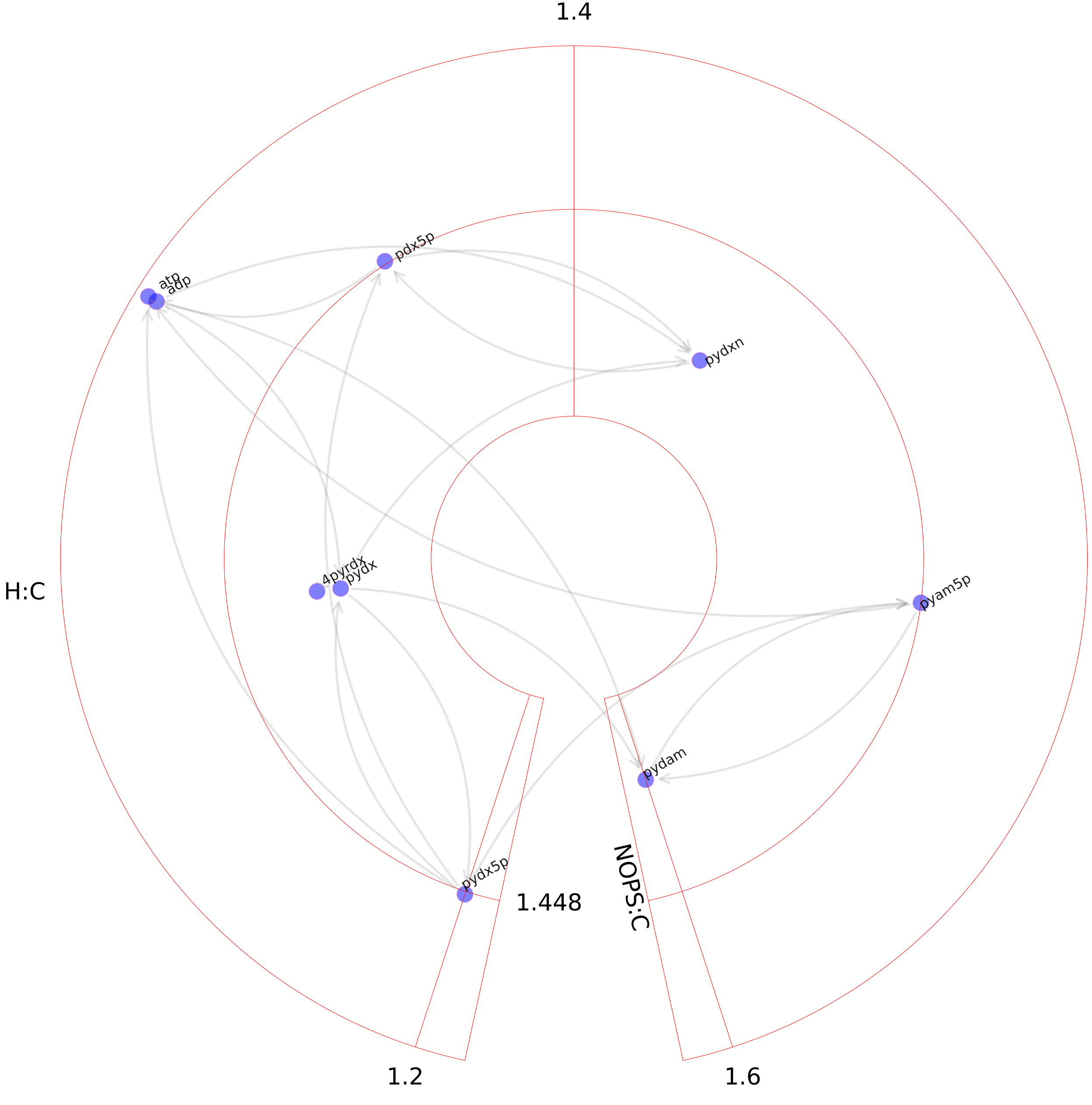

### Vitamin B12 metabolism.pdf

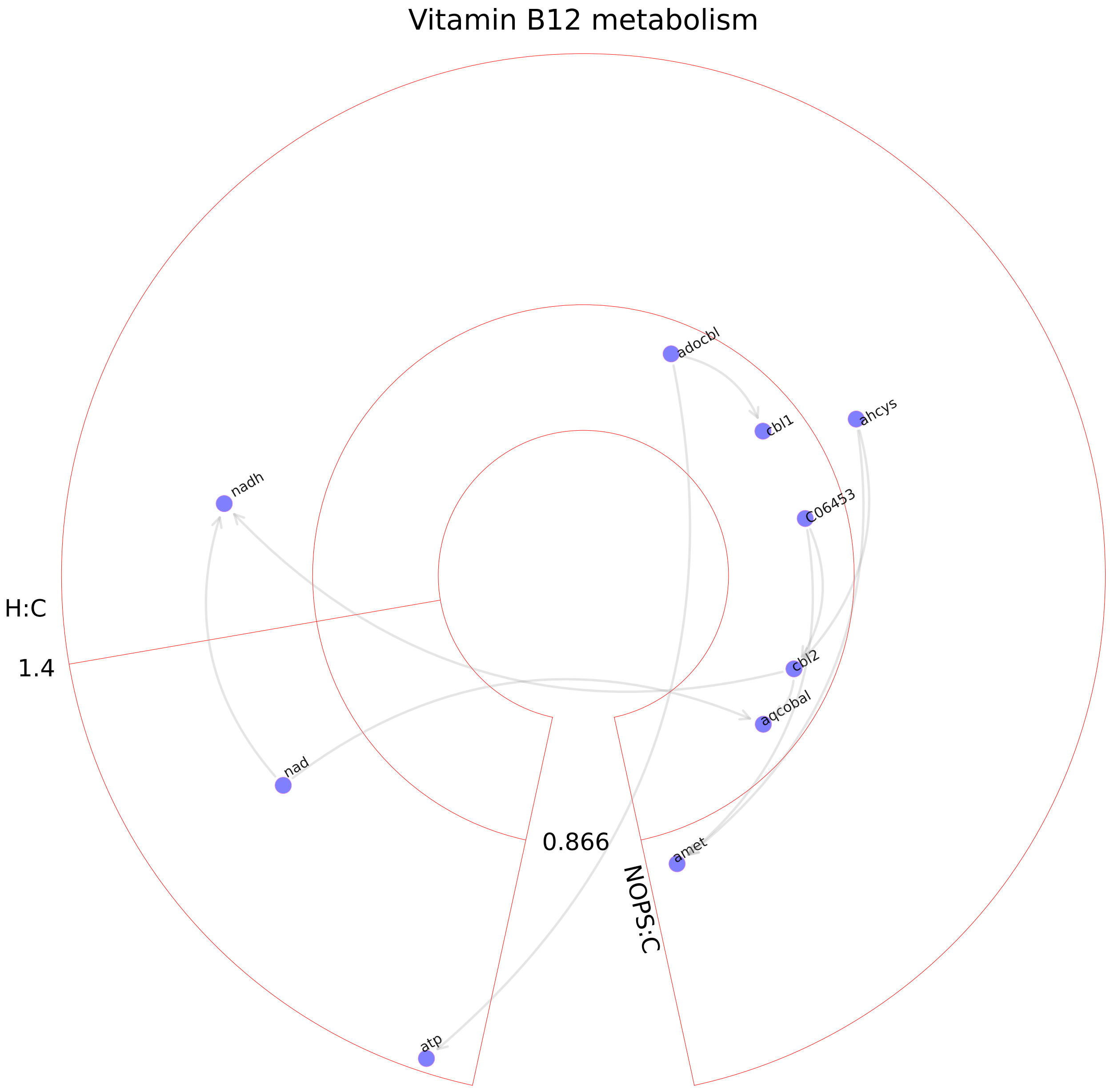

### Vitamin C metabolism.pdf

#### Vitamin C metabolism

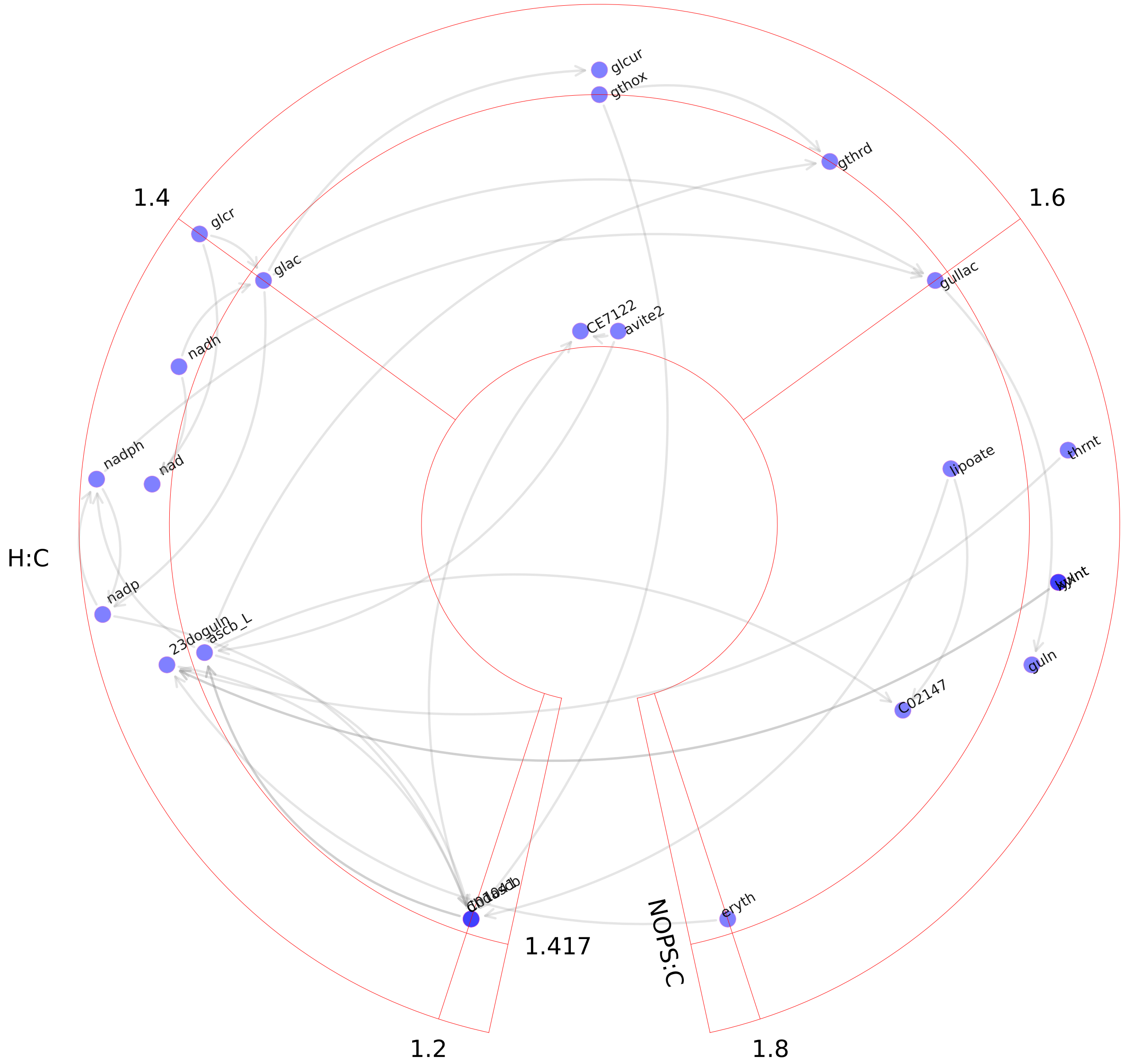

### Vitamin D metabolism.pdf

Vitamin D metabolism

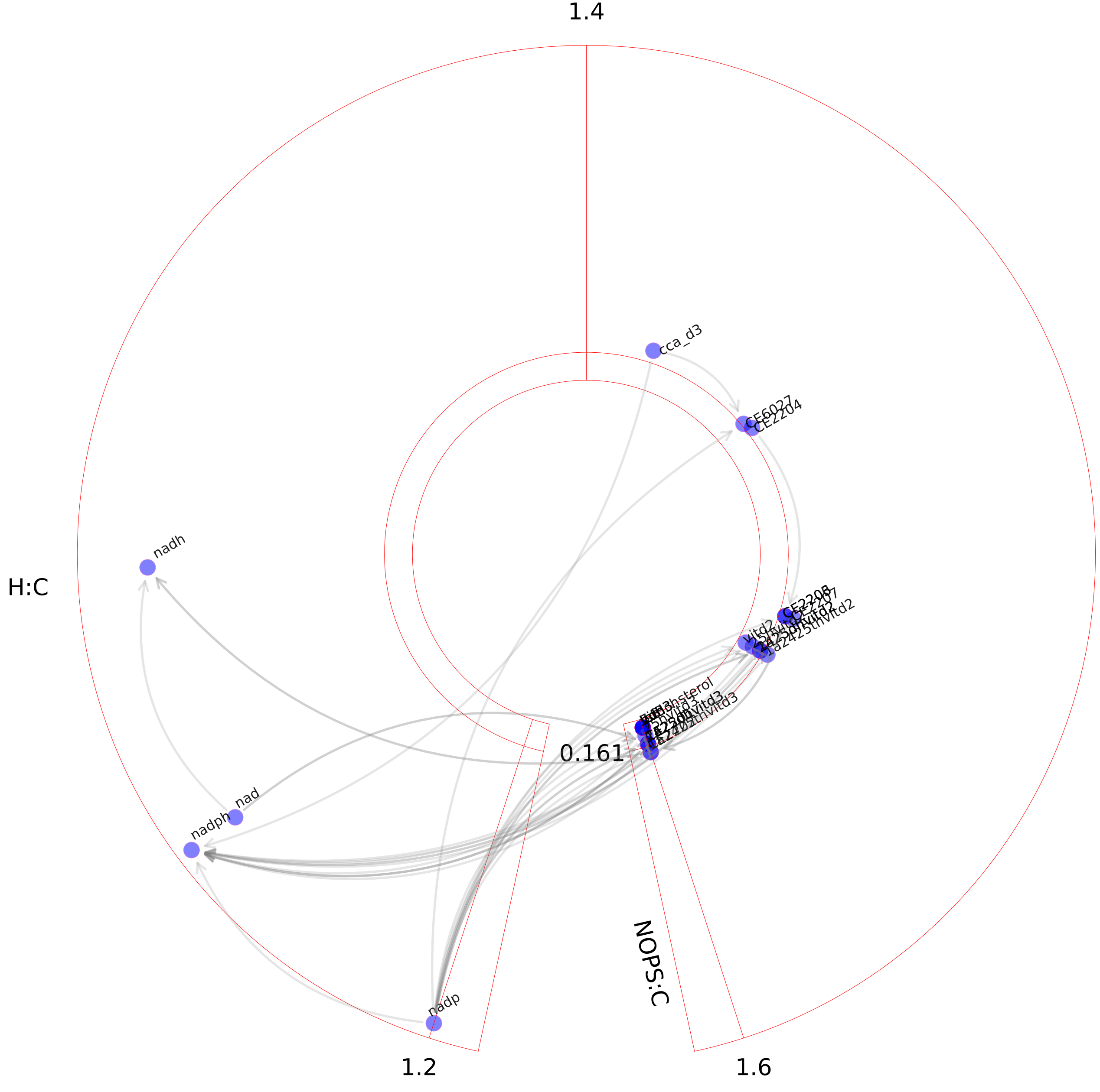

### Vitamin E metabolism.pdf

#### Vitamin E metabolism

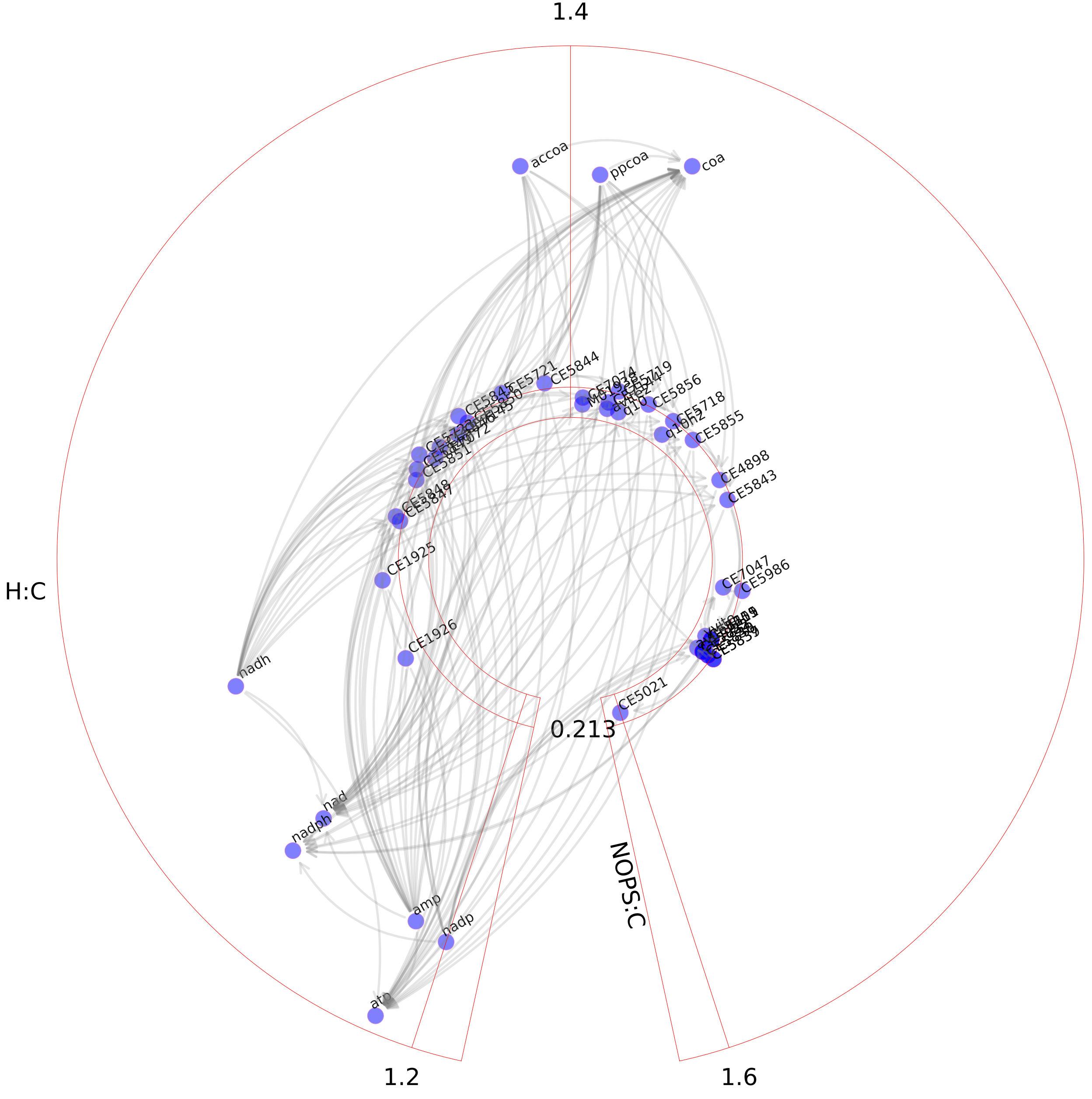

### Vitamin K metabolism.pdf

#### Vitamin K metabolism

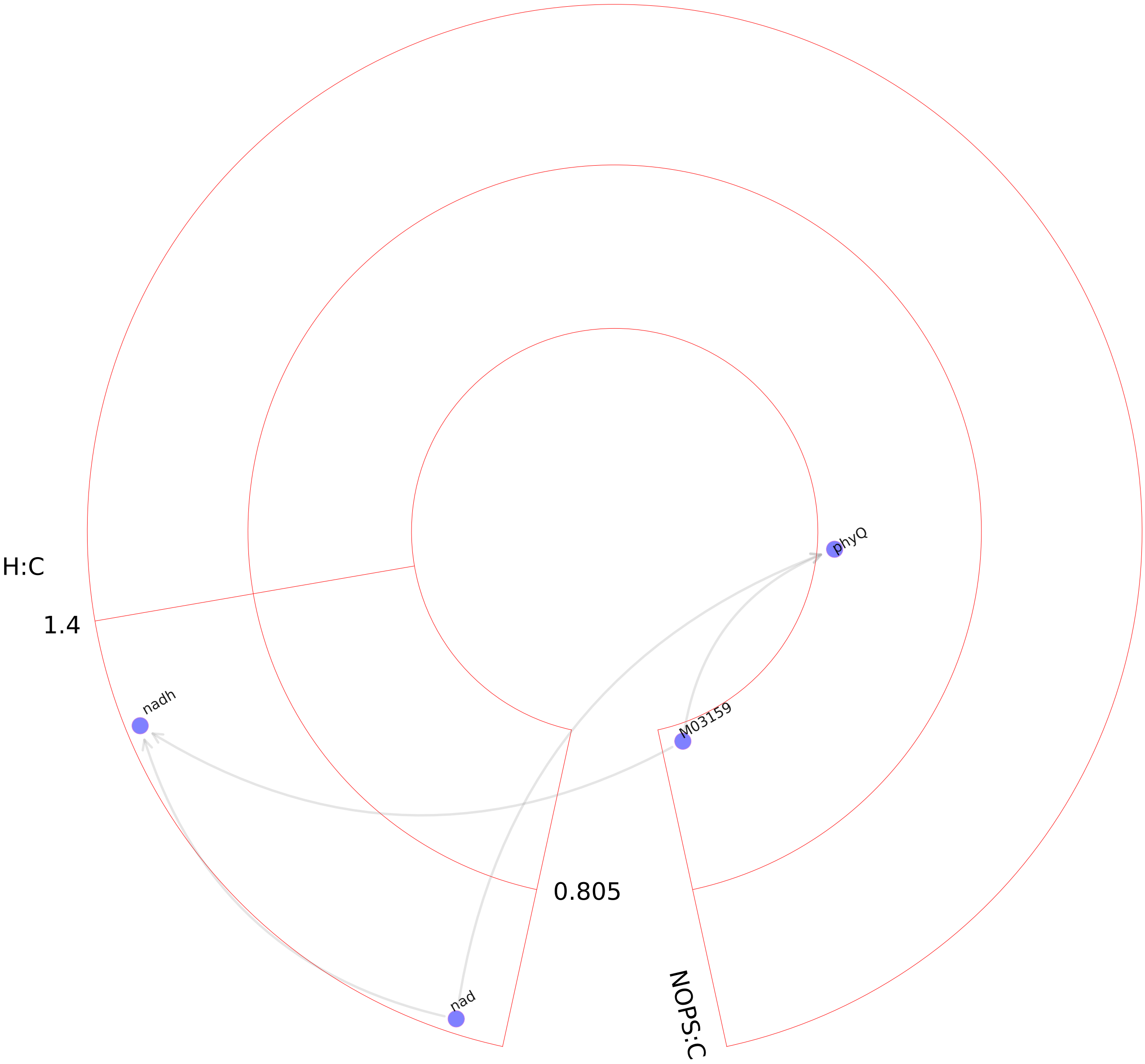

### Xenobiotics metabolism.pdf

Xenobiotics metabolism

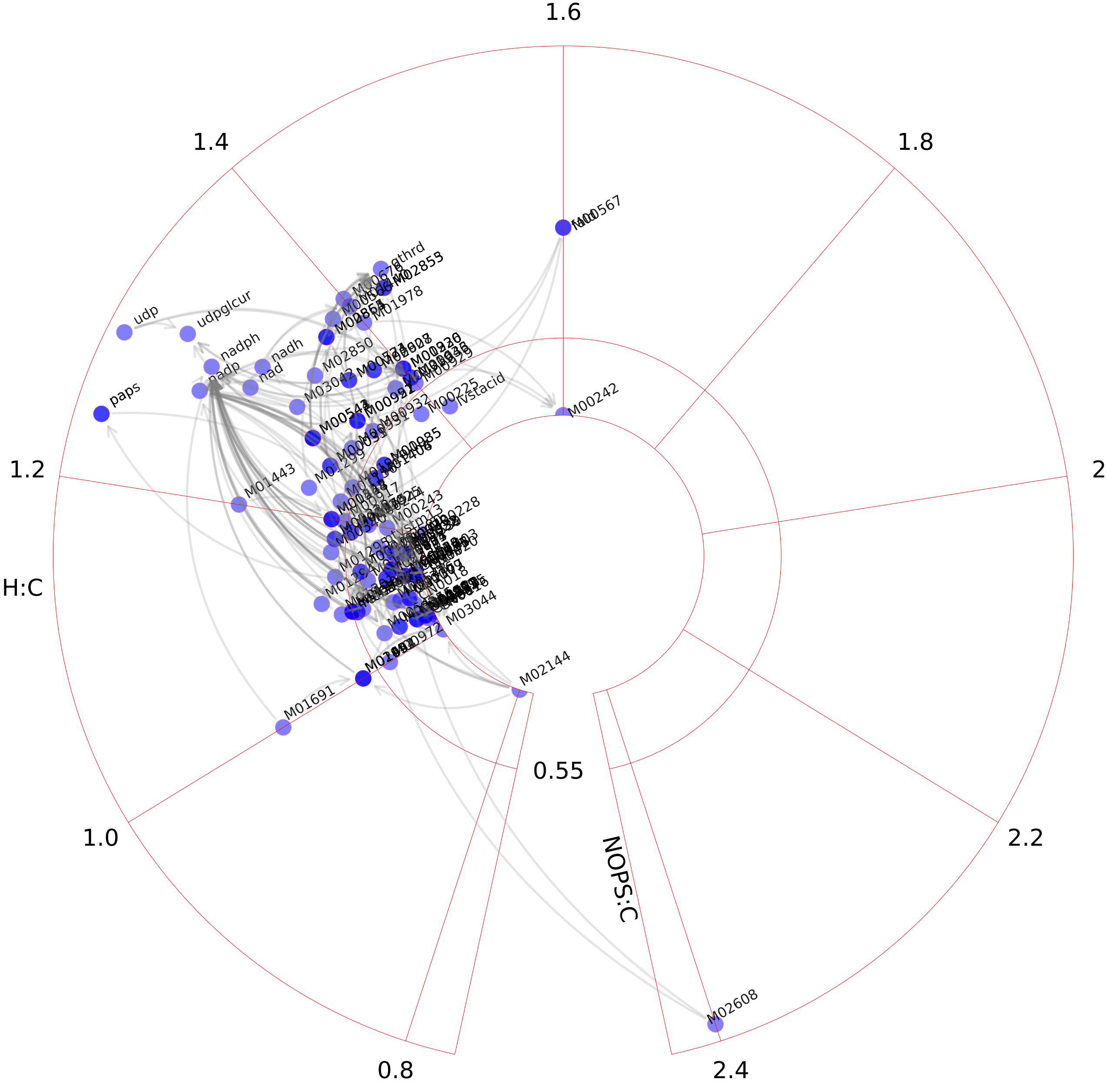
