## Supplementary figures and images for "Circular van Krevelen diagram for visualizing metabolic pathways"

### Alanine and aspartate metabolism.pdf

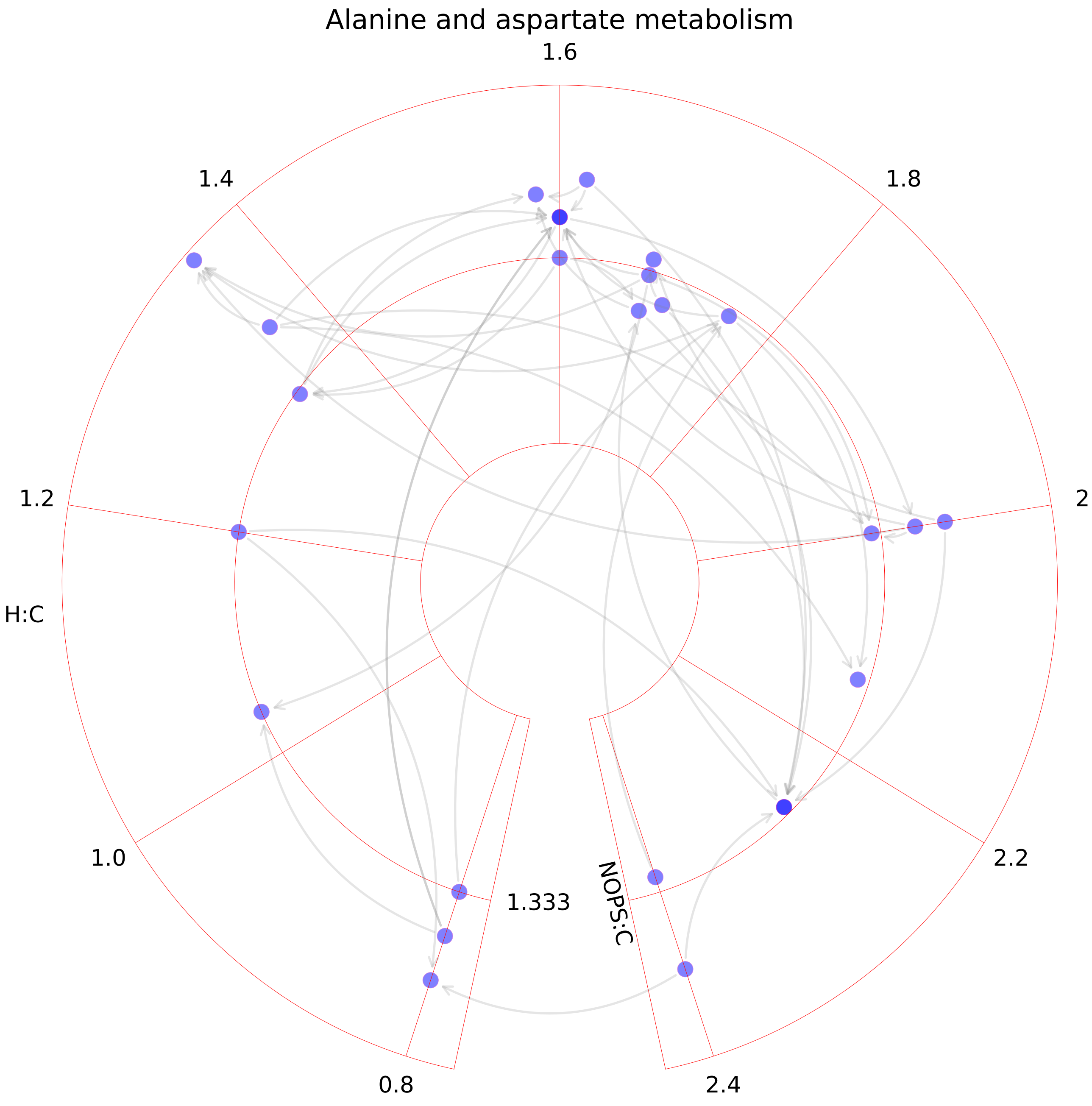

### Alkaloid synthesis.pdf

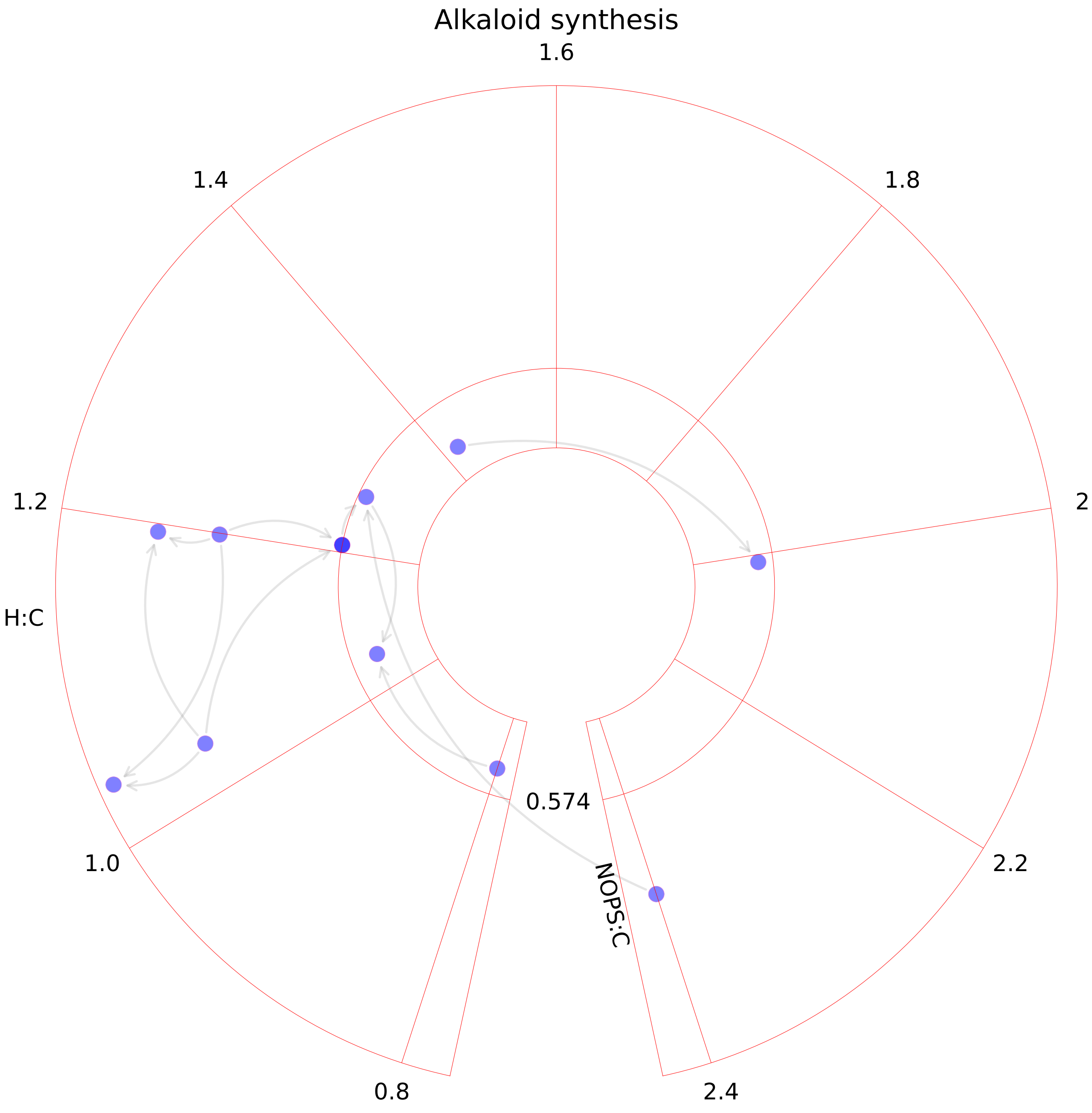

### Aminoacyl-tRNA biosynthesis.pdf

#### Aminoacyl-tRNA biosynthesis

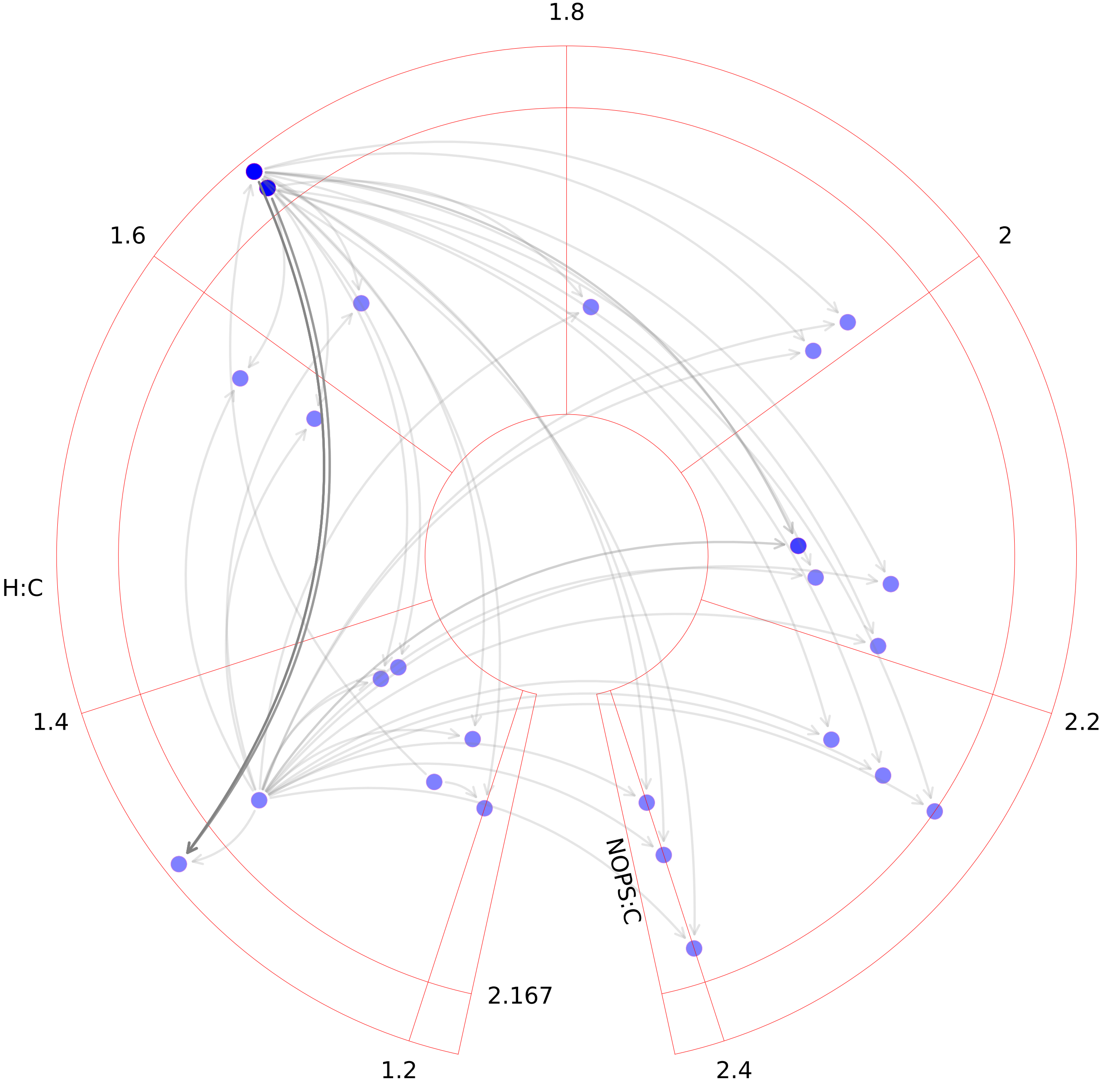

### Aminosugar metabolism.pdf

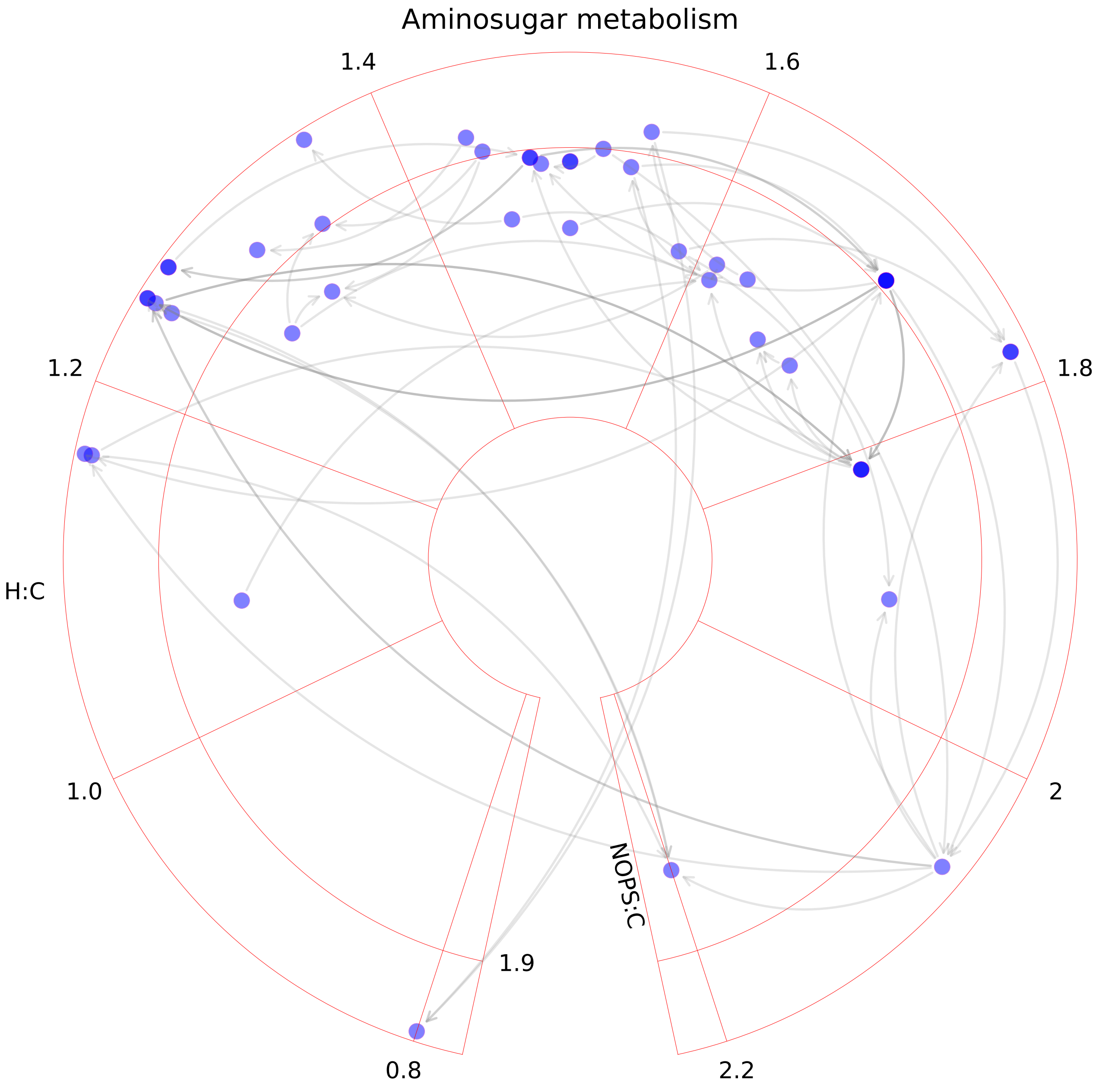

### Androgen and estrogen synthesis and metabolism.pdf

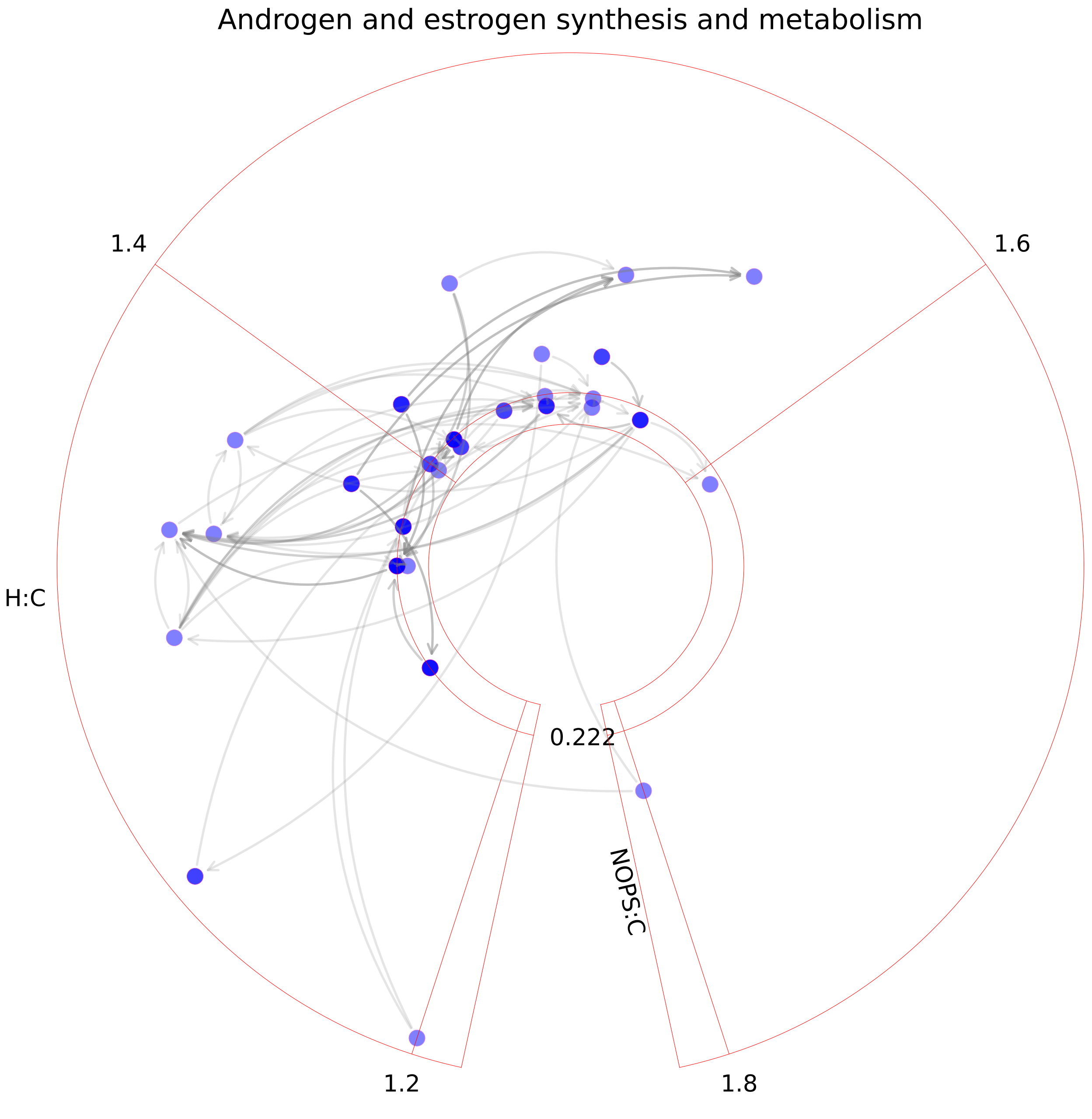

### Arachidonic acid metabolism.pdf

##### Arachidonic acid metabolism

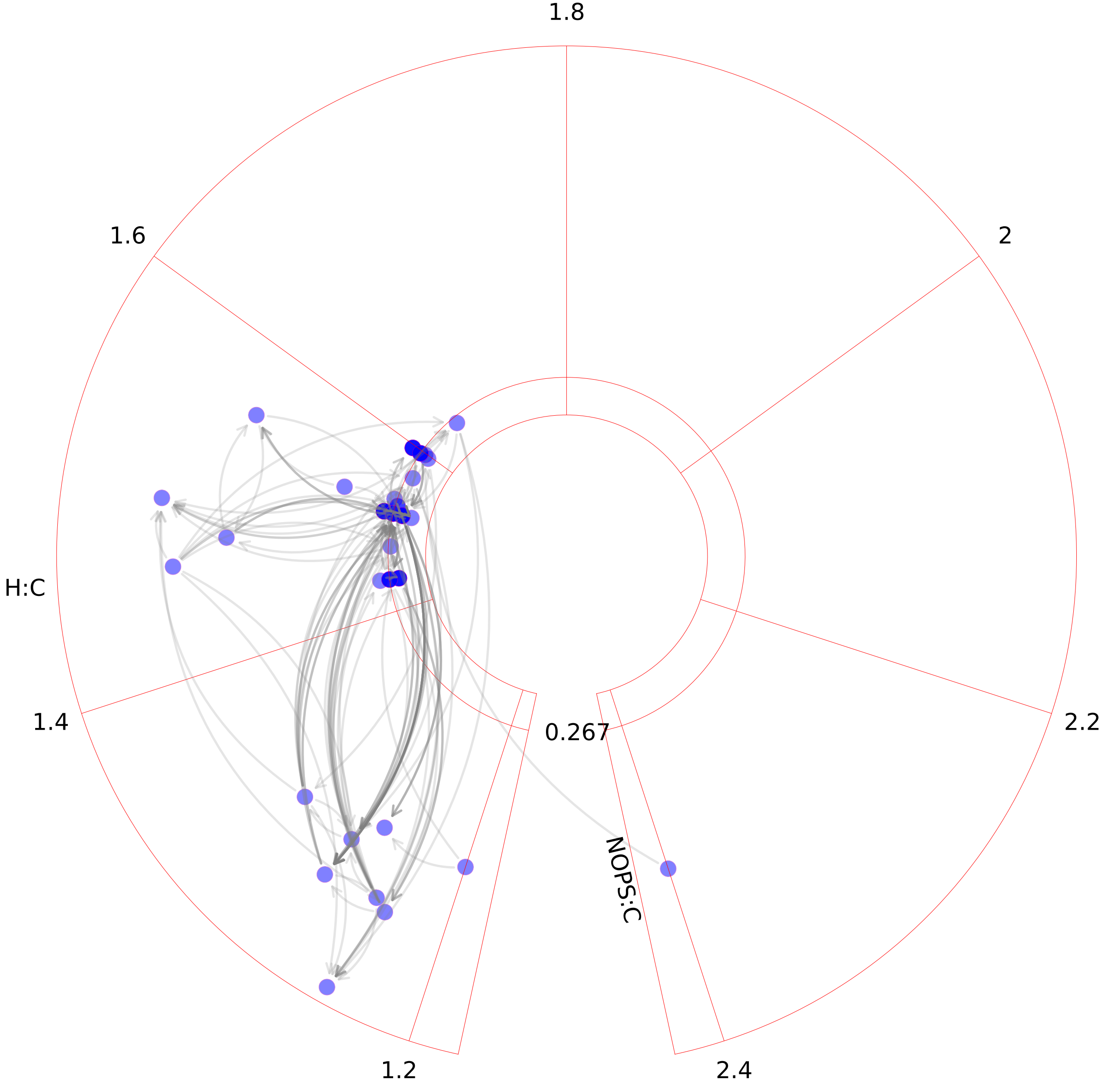

### Arginine and proline metabolism.pdf

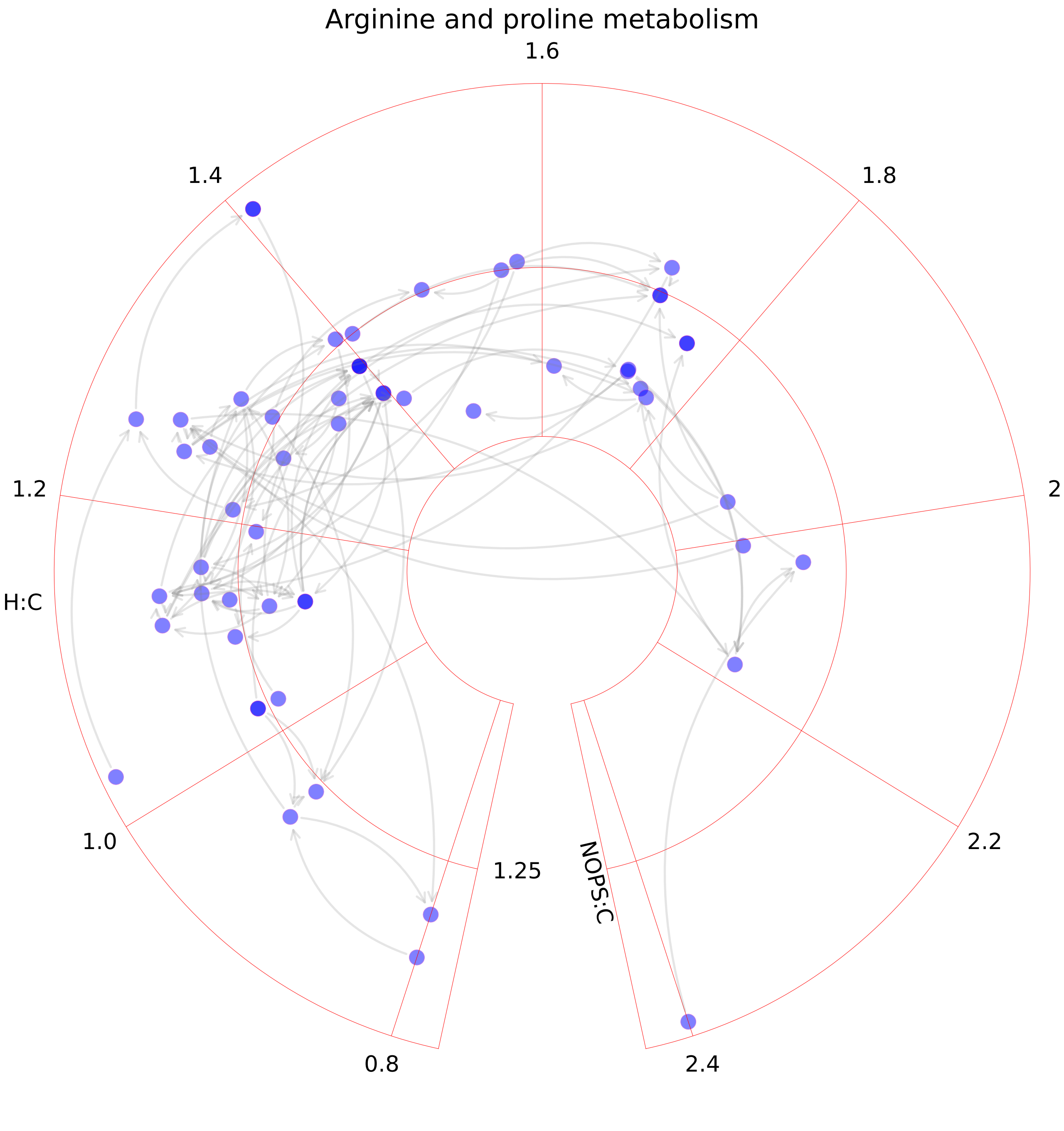

### Beta-Alanine metabolism.pdf

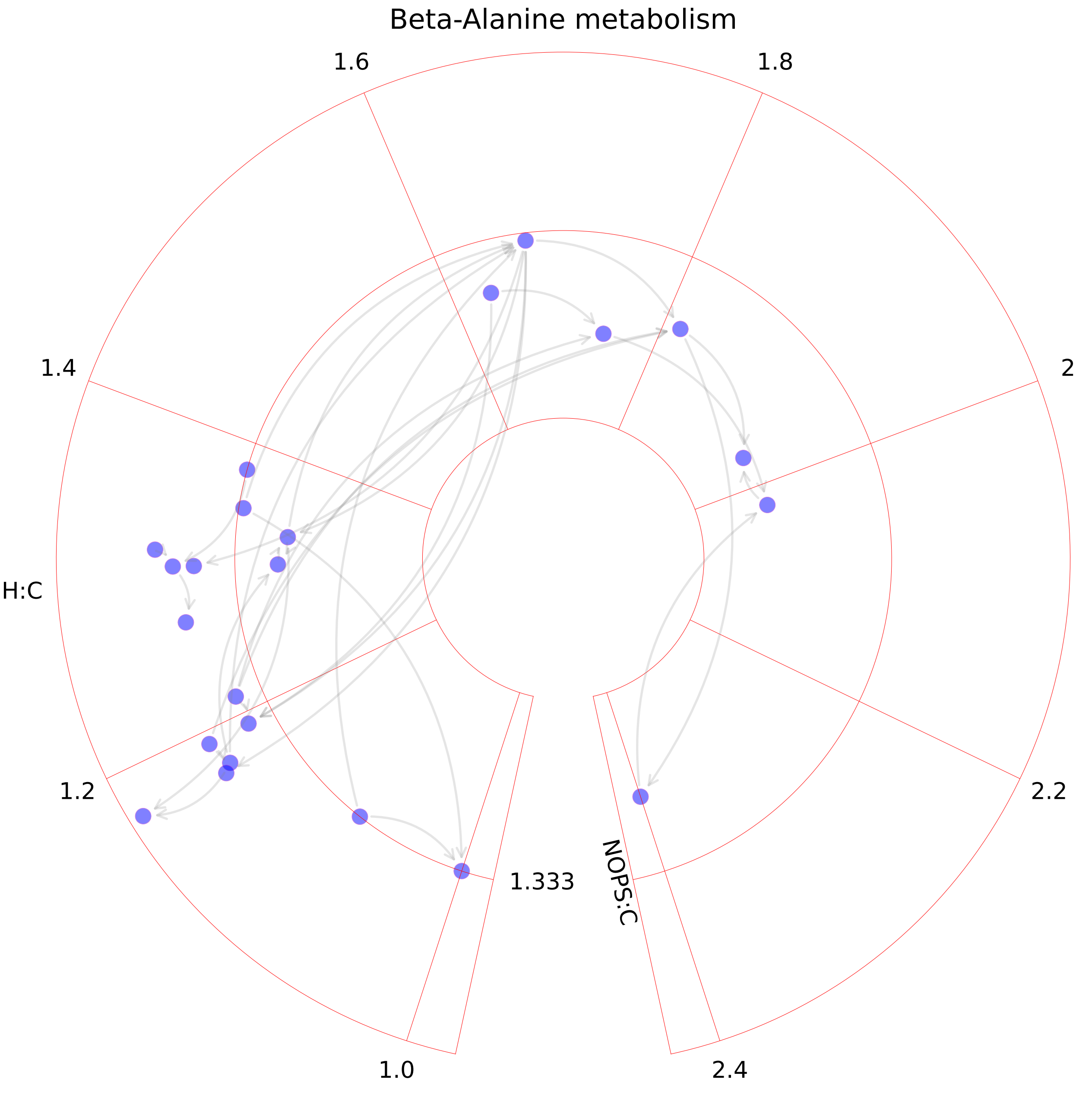

### Bile acid synthesis.pdf

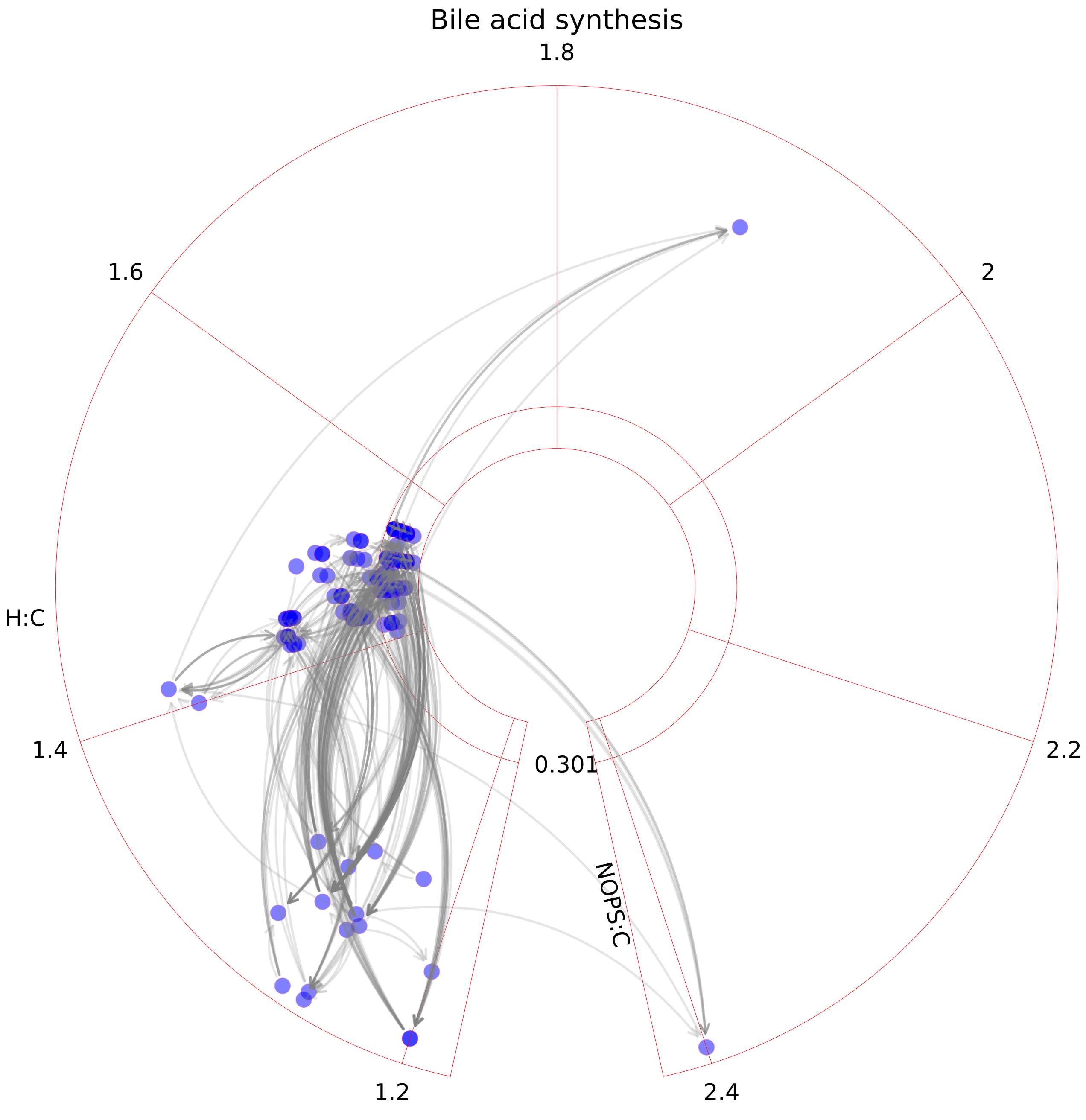

### Biotin metabolism.pdf

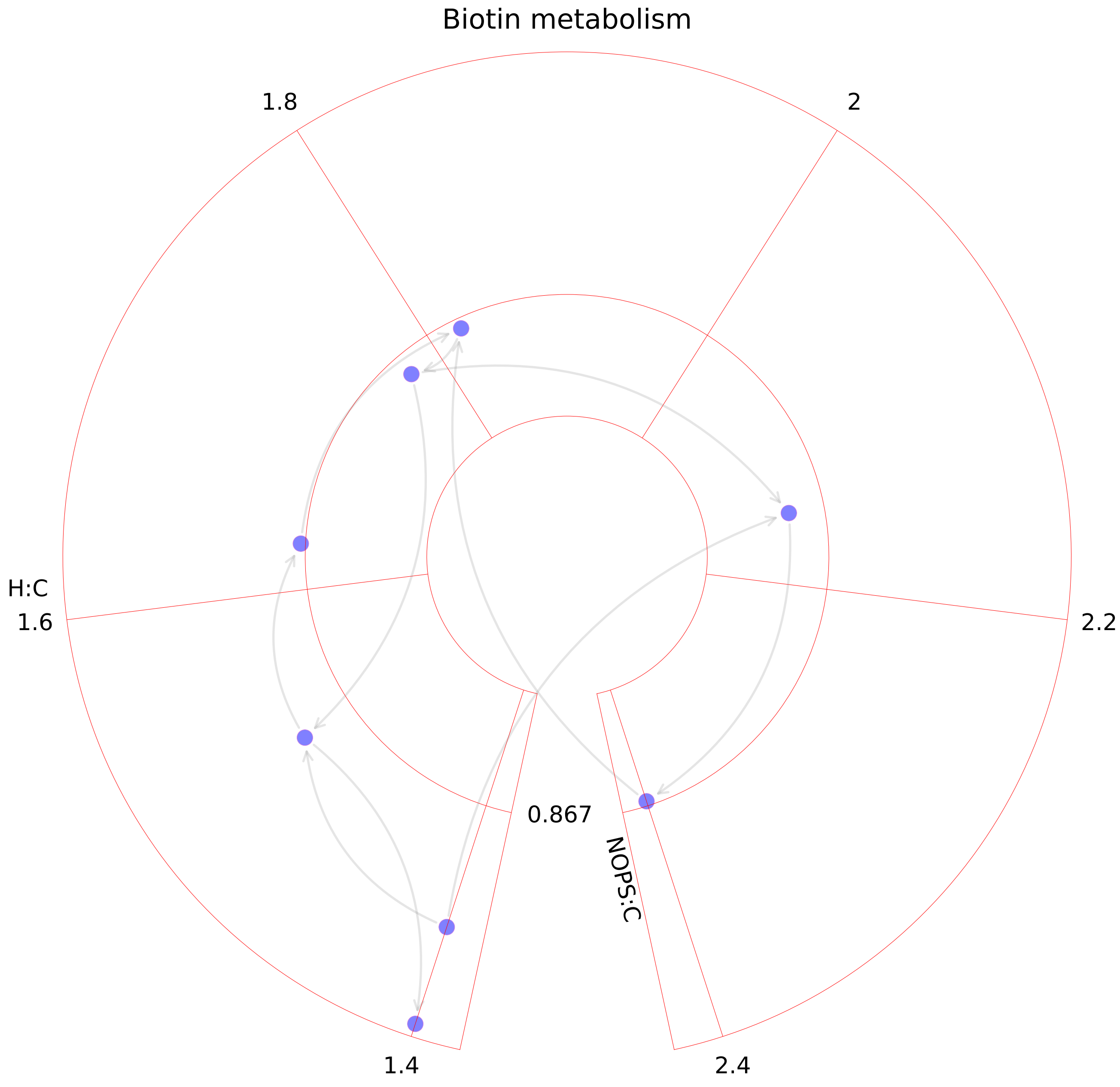

### Blood group synthesis.pdf

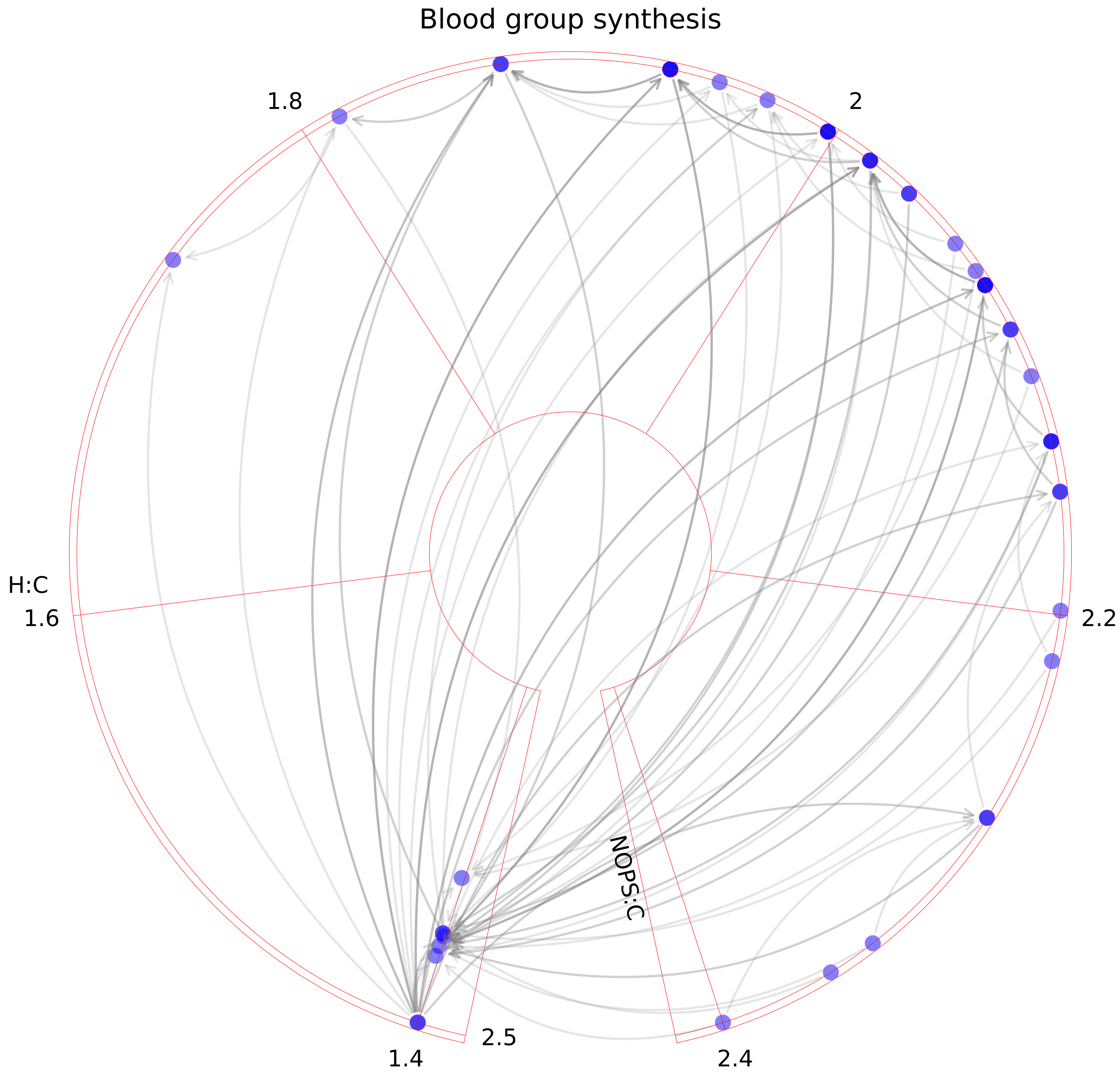

### Butanoate metabolism.pdf

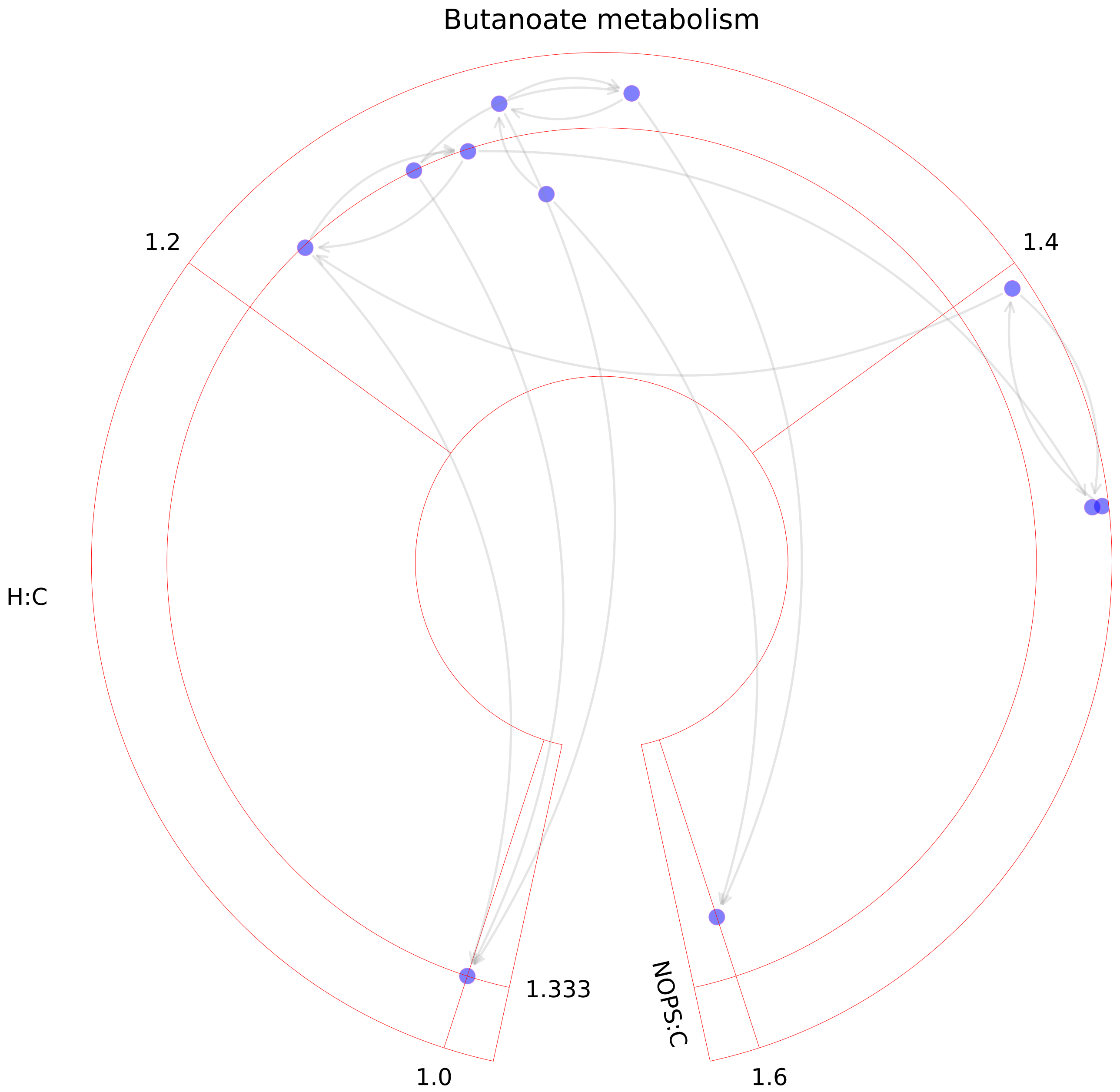

### C5-branched dibasic acid metabolism.pdf

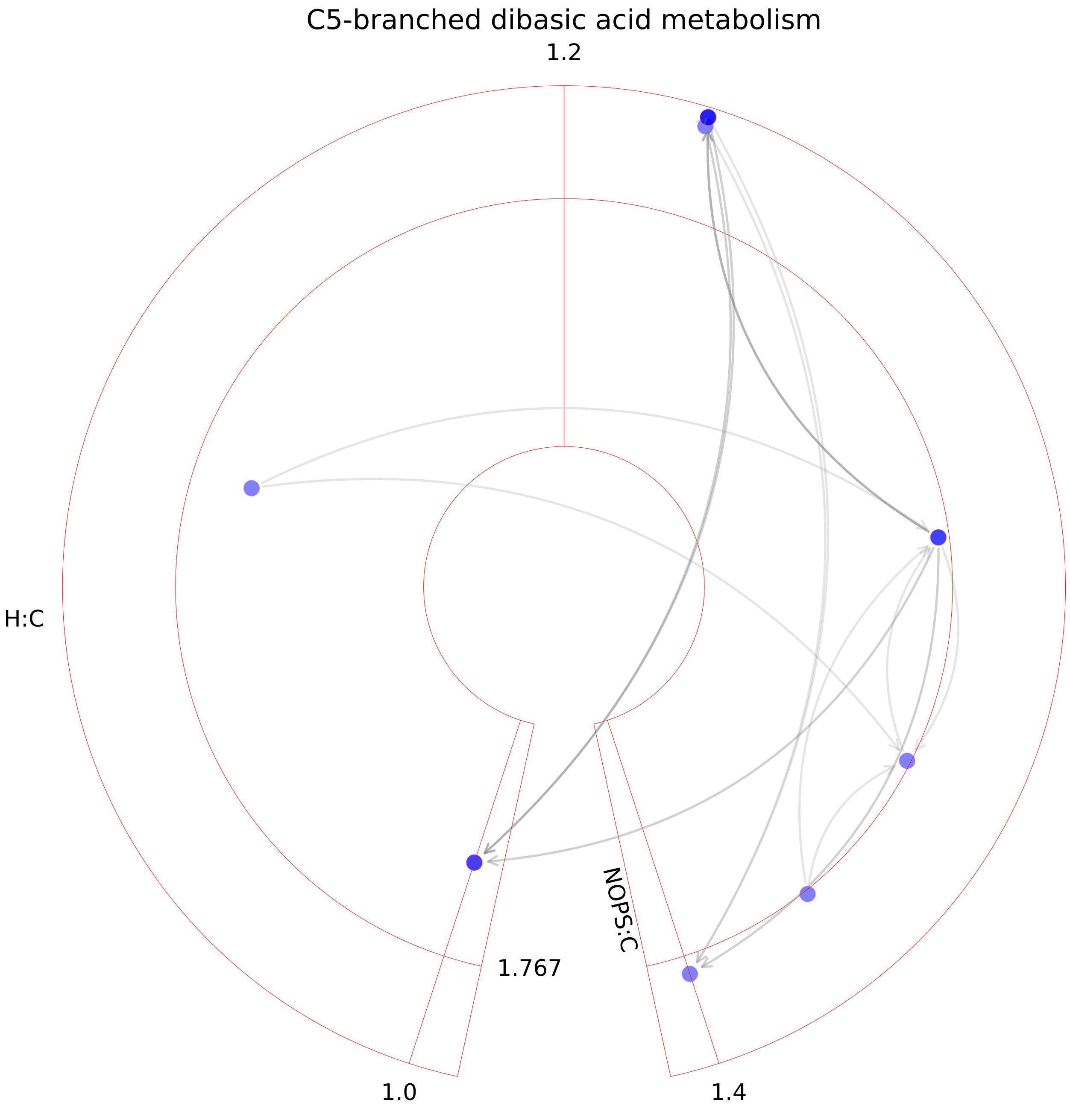

### Cholesterol metabolism.pdf

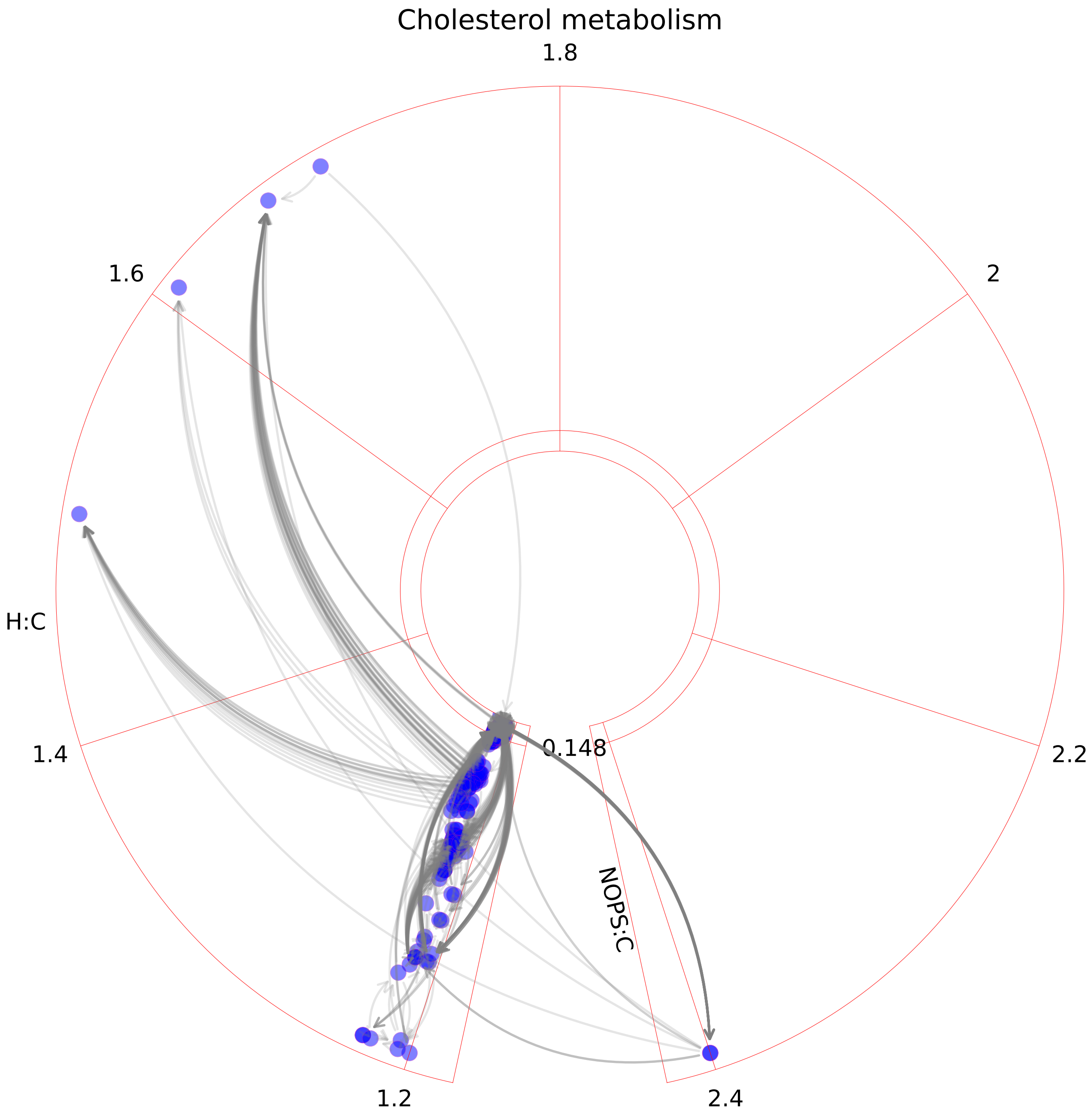

### Chondroitin sulfate degradation.pdf

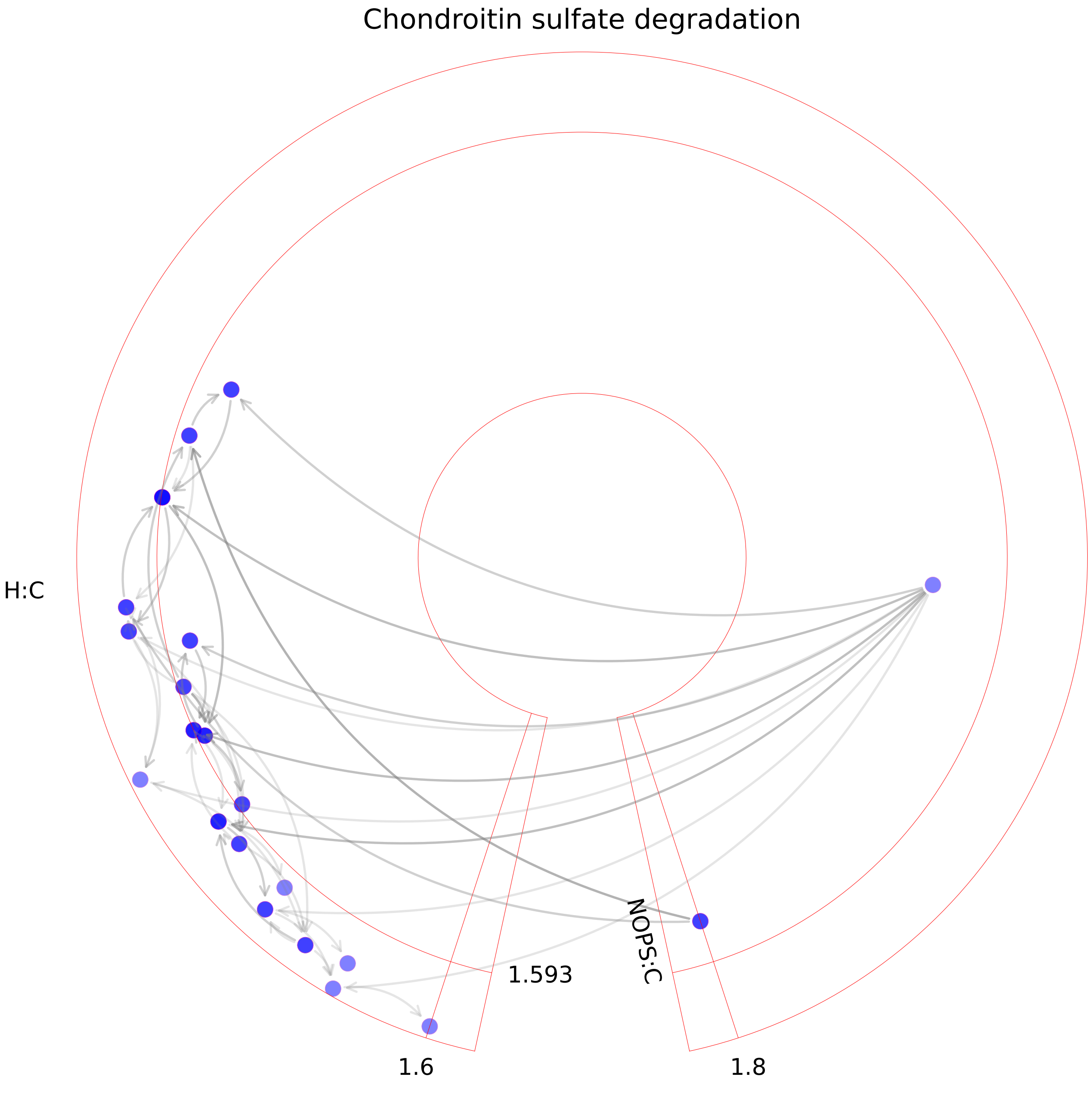

### Chondroitin synthesis.pdf

##### Chondroitin synthesis

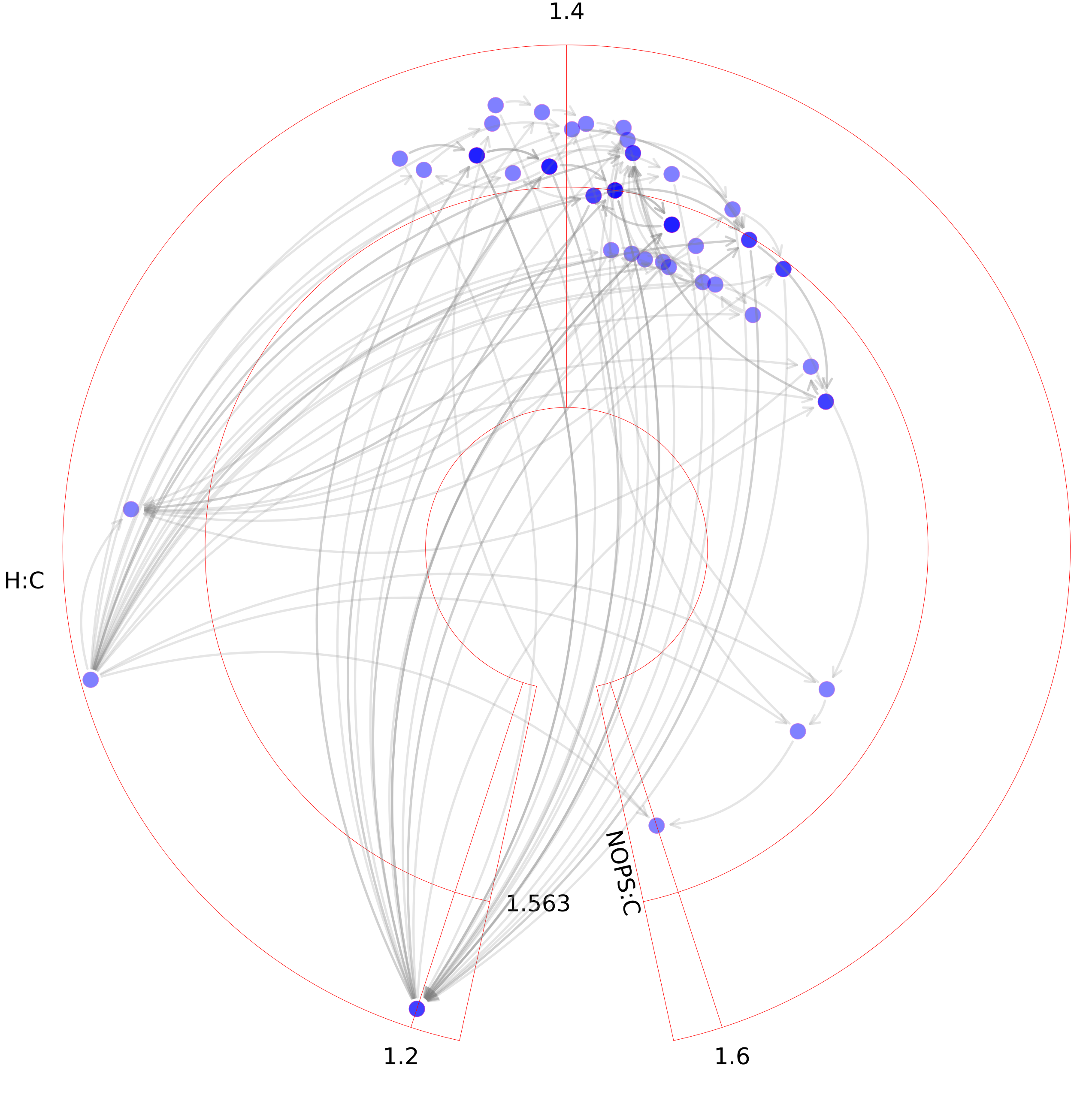

### Citric acid cycle.pdf

#### Citric acid cycle

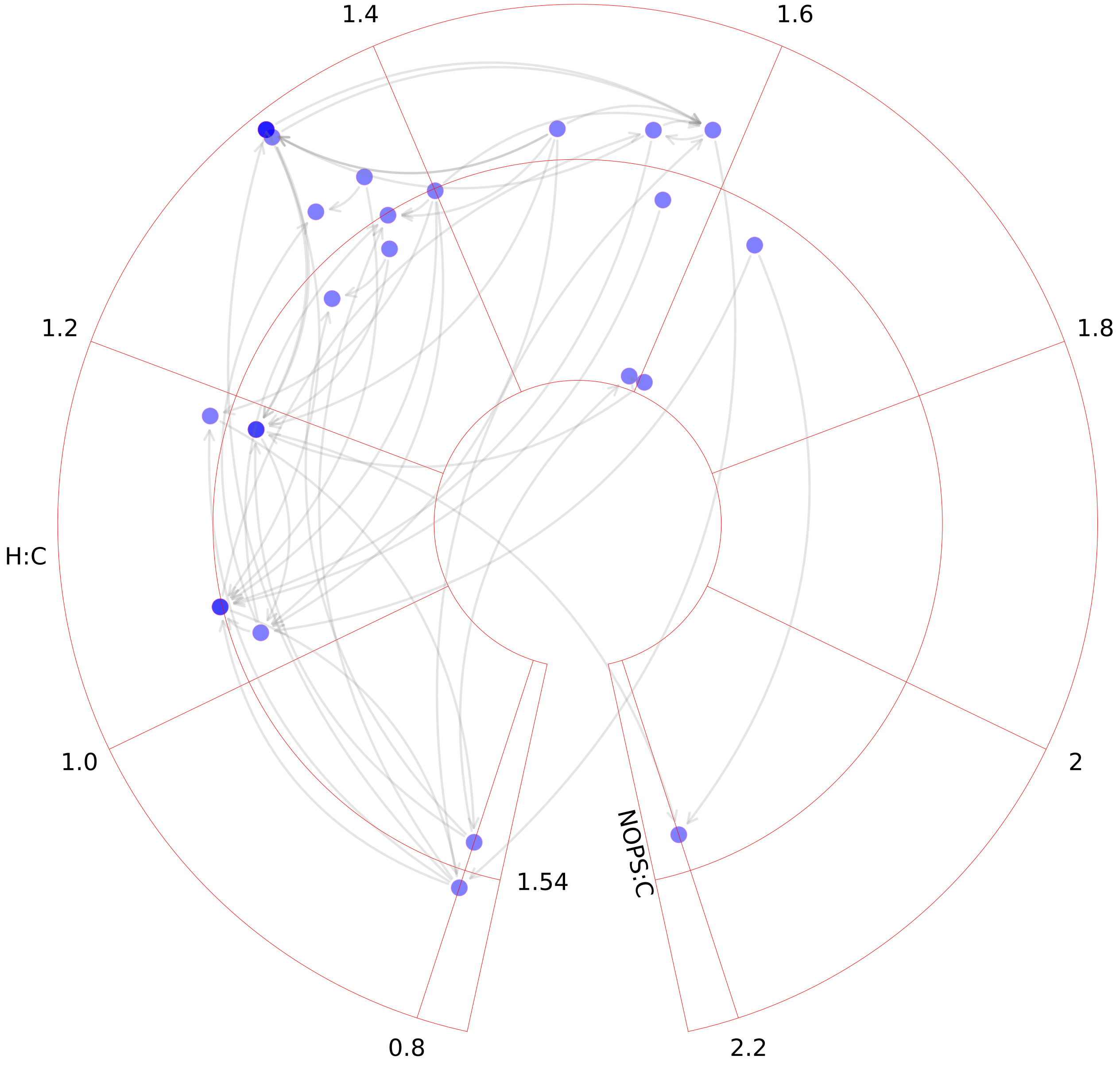

### CoA catabolism.pdf

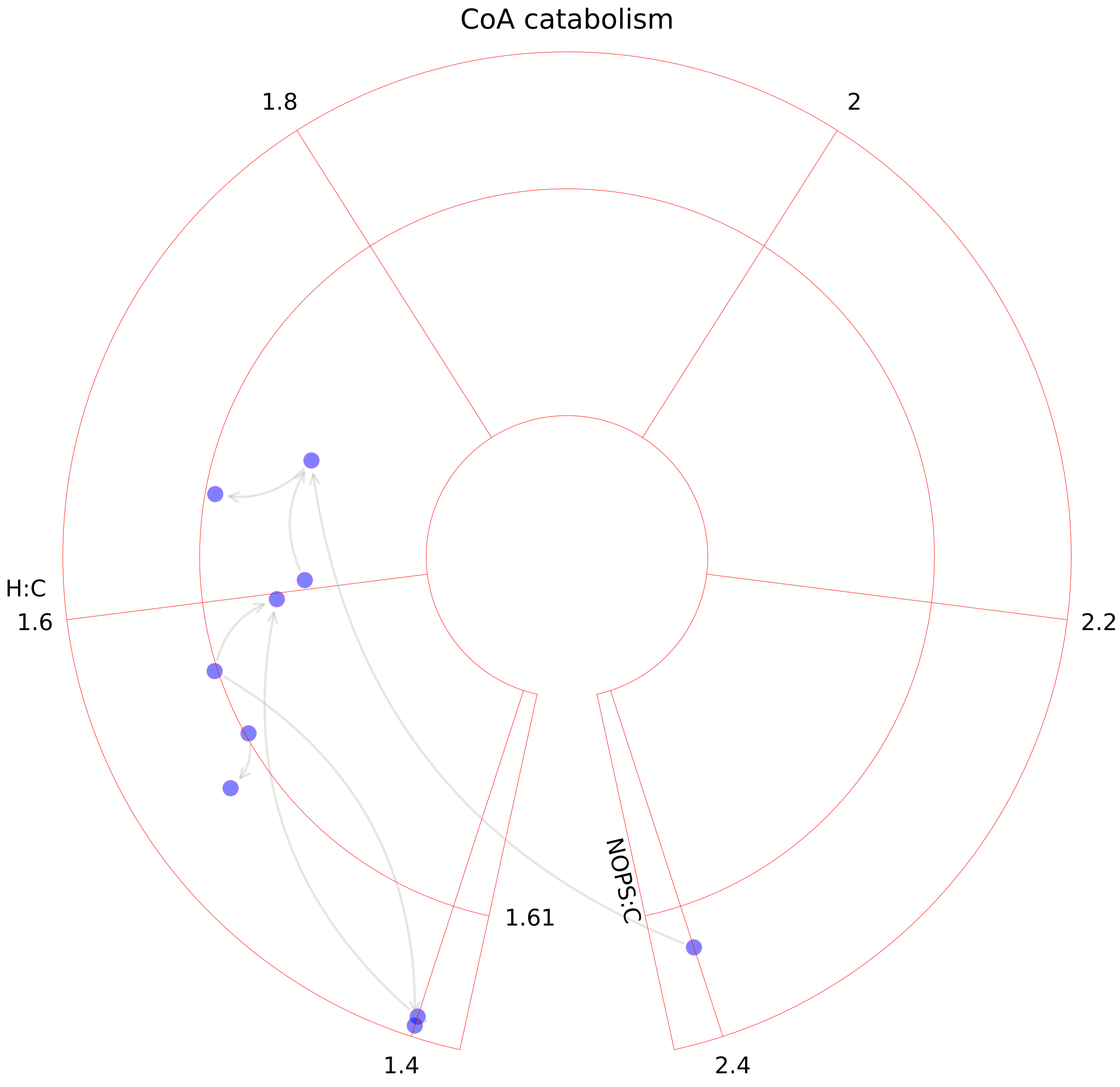

### CoA synthesis.pdf

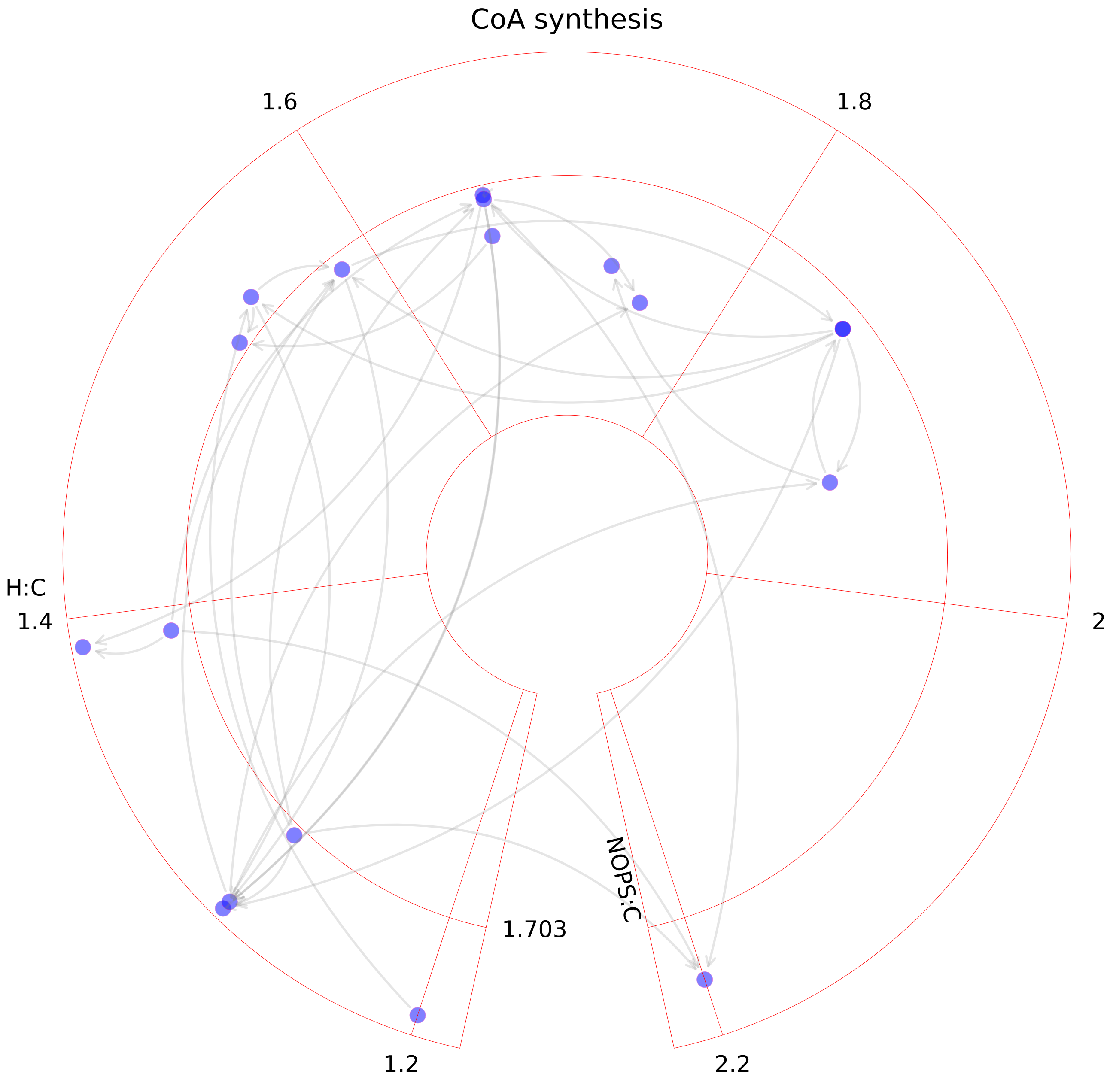

### Cytochrome metabolism.pdf

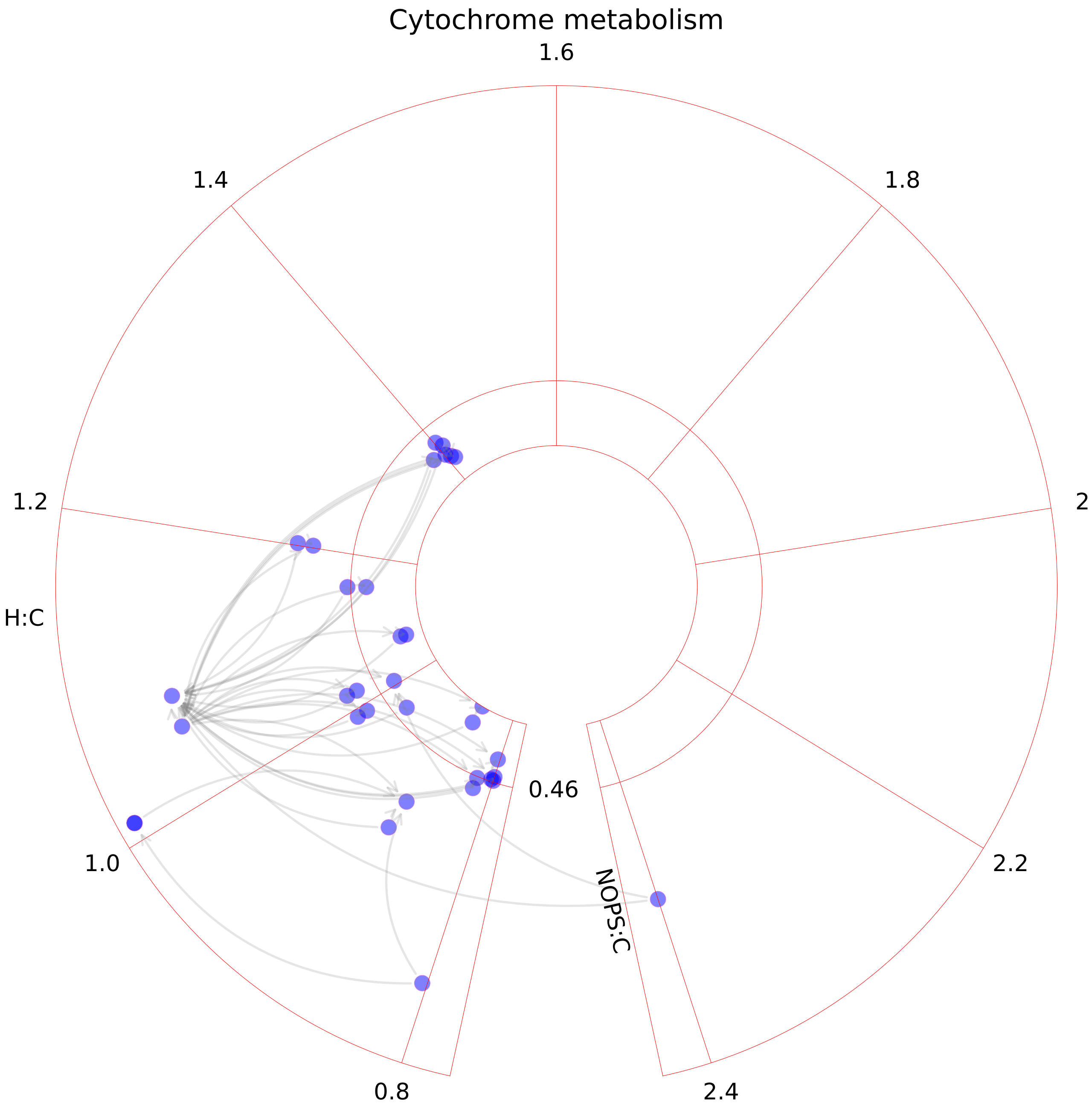

### D-alanine metabolism.pdf

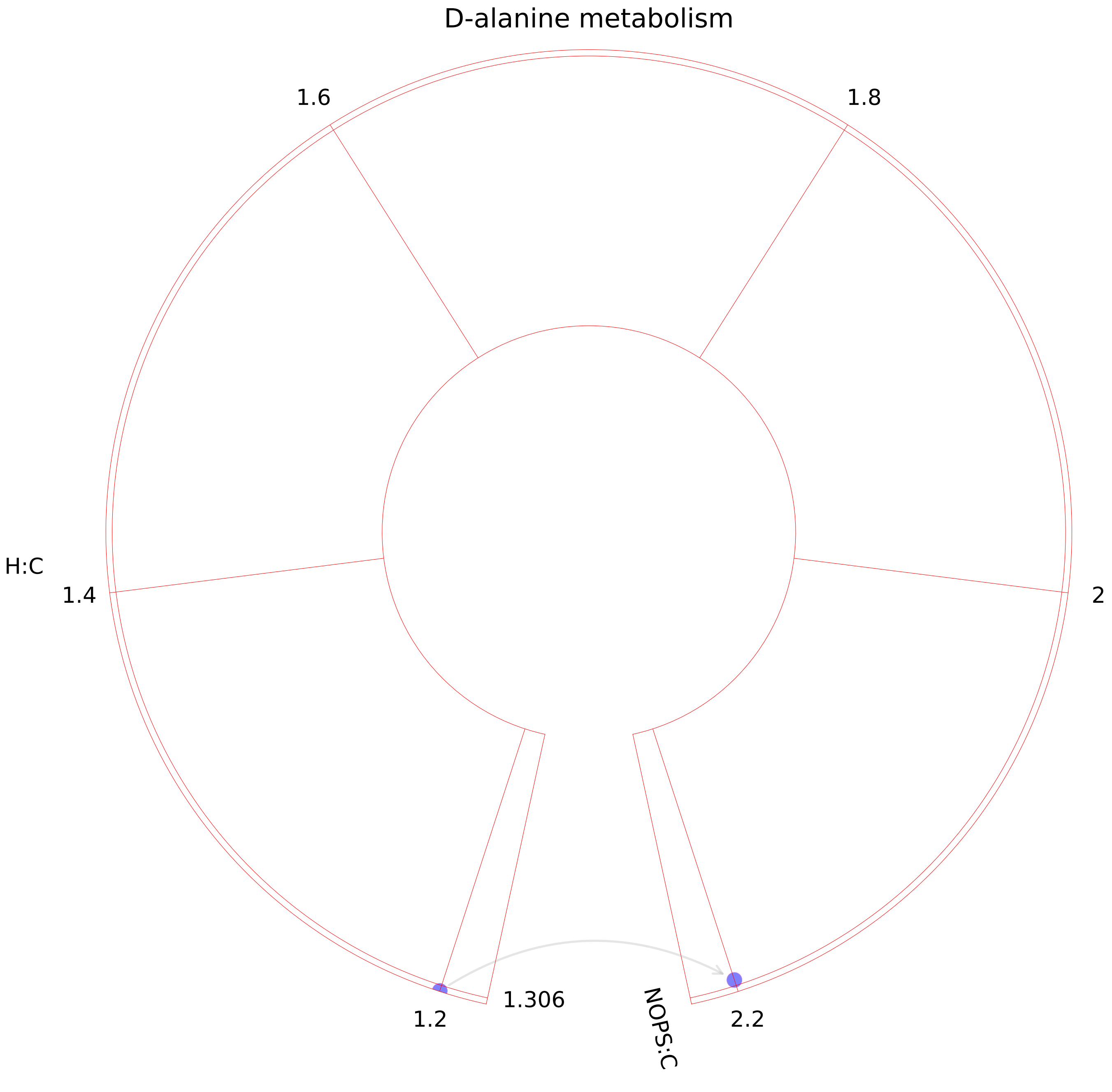

### Dietary fiber binding.pdf

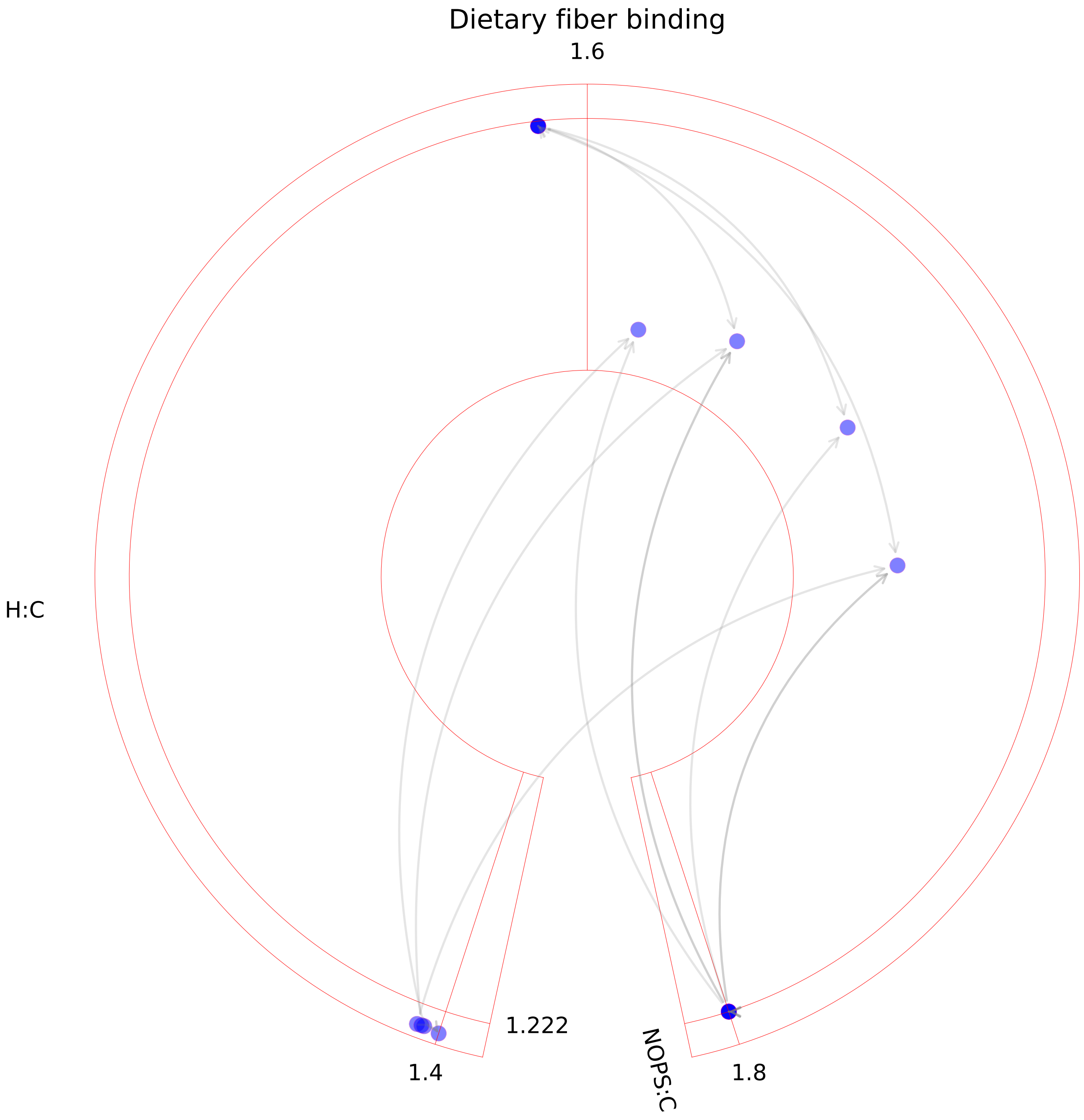

### Drug metabolism.pdf

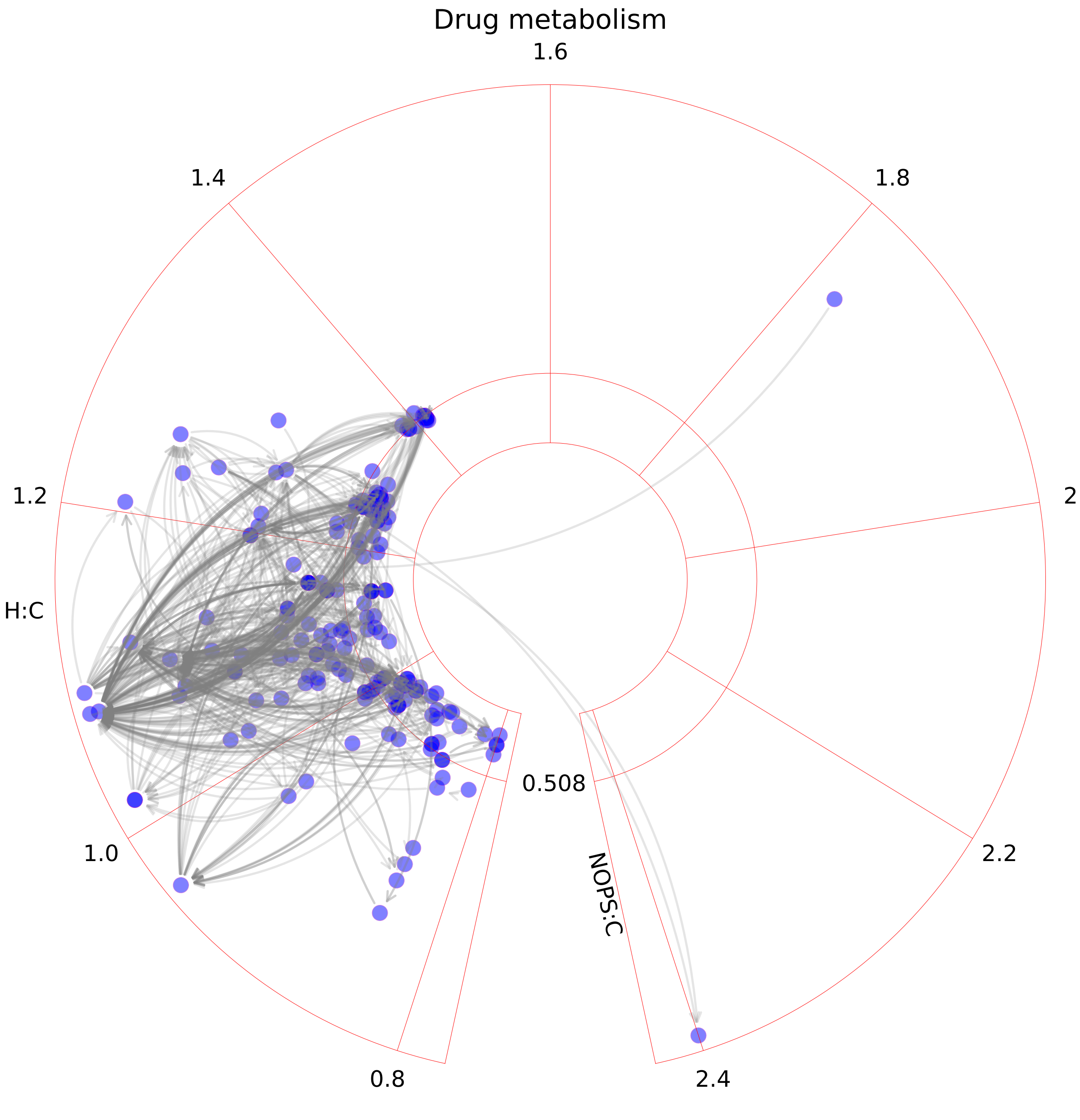

### Eicosanoid metabolism.pdf

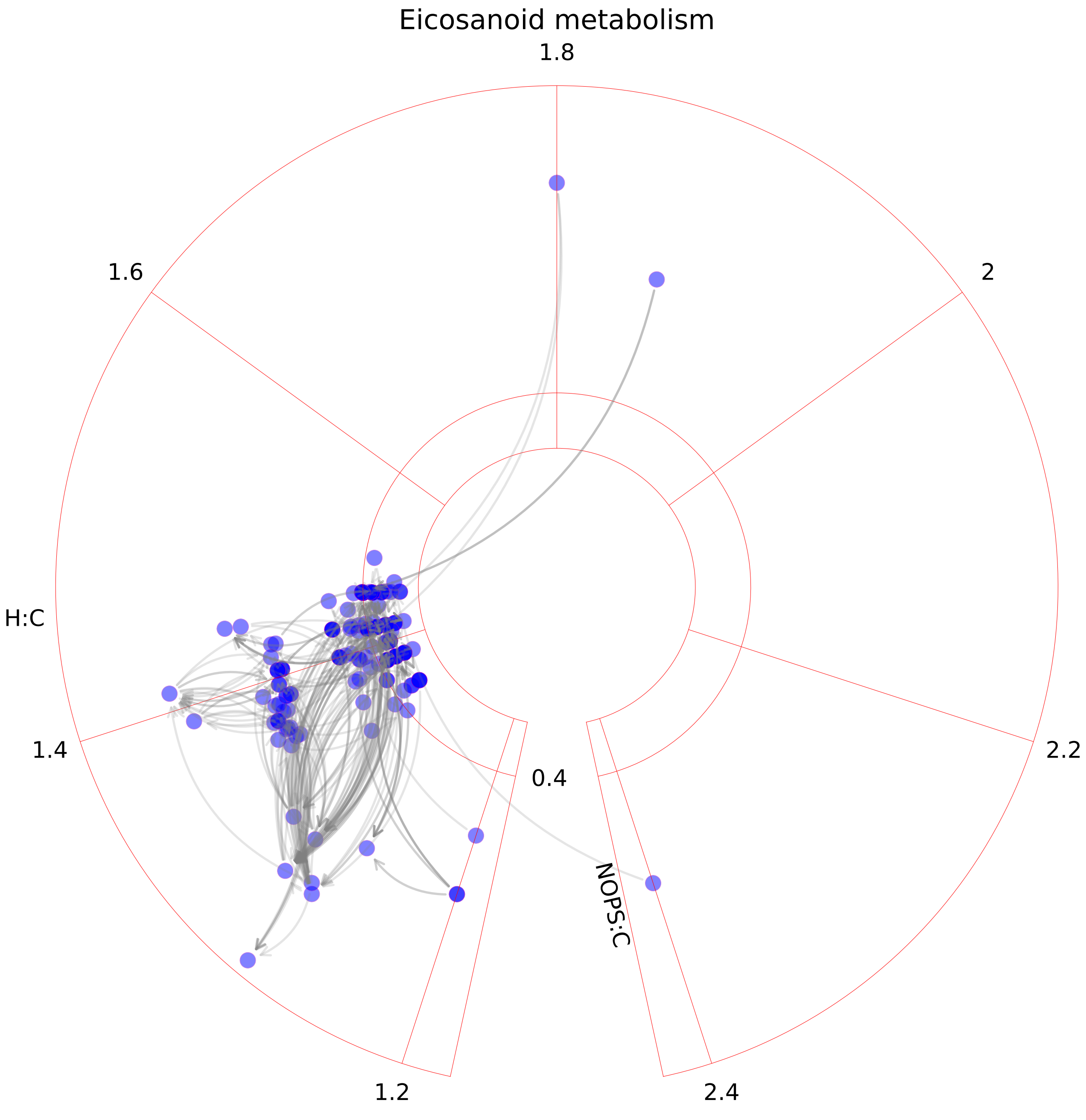

### Fatty acid oxidation.pdf

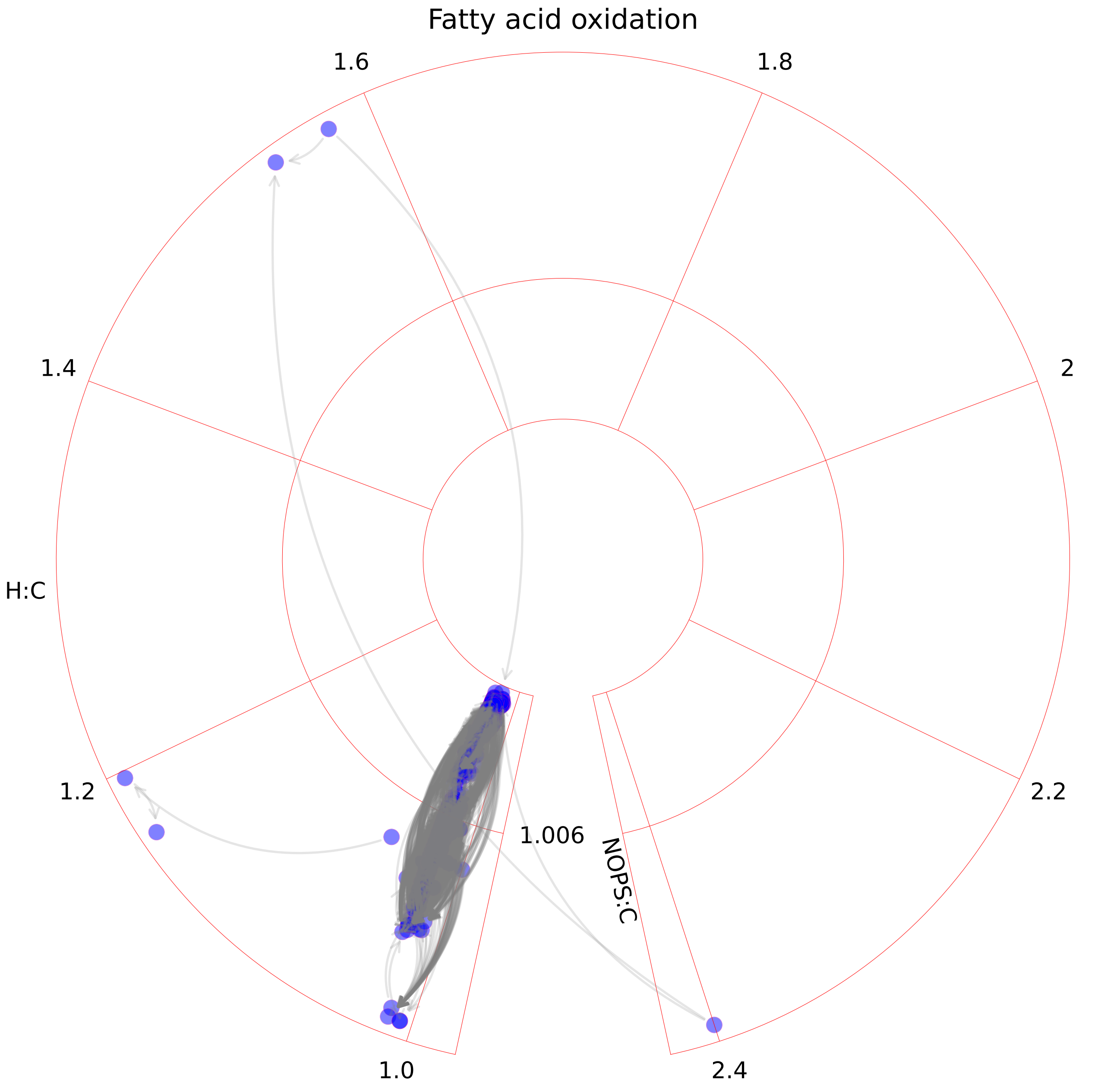

### Fatty acid synthesis.pdf

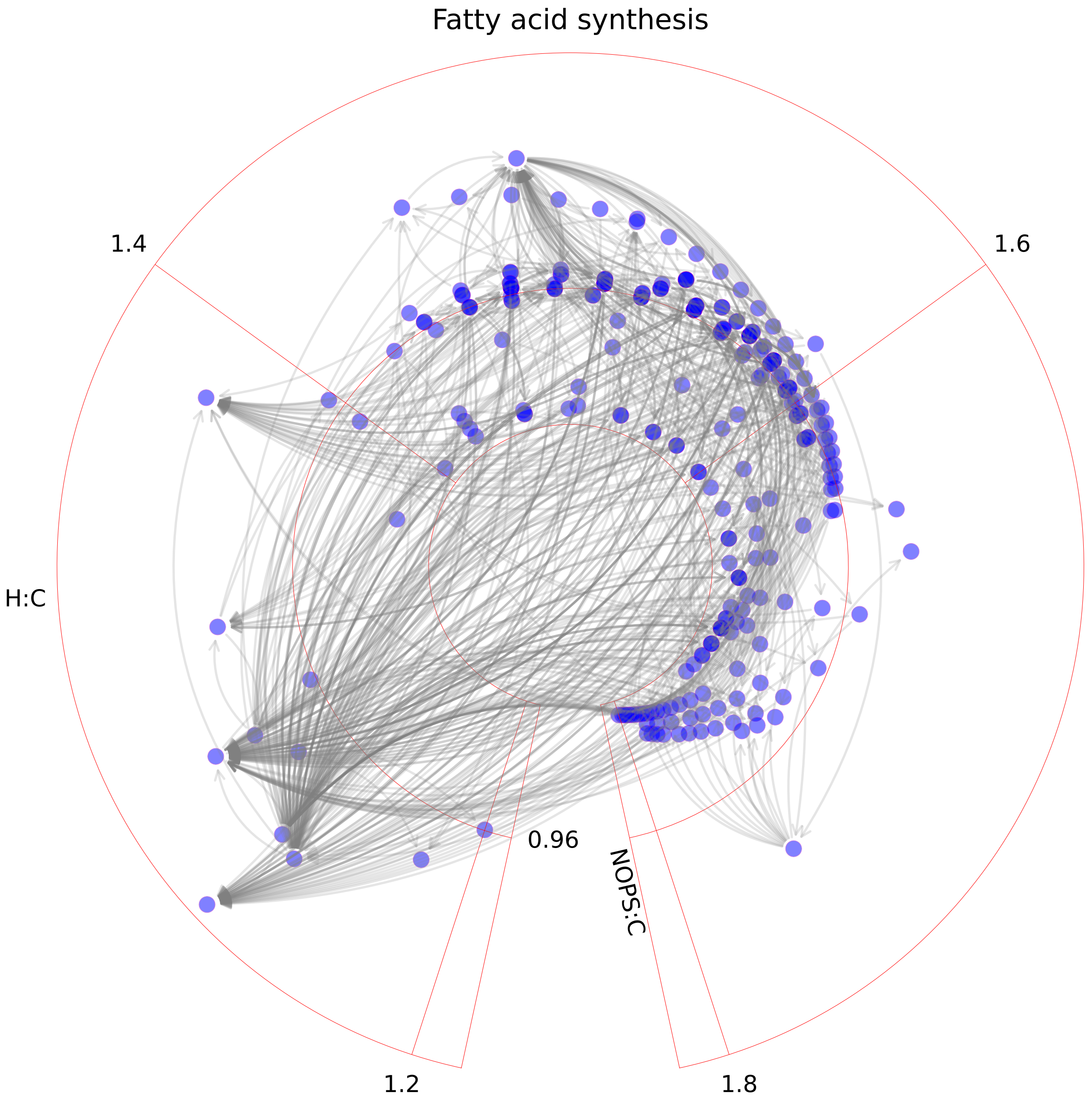

### Folate metabolism.pdf

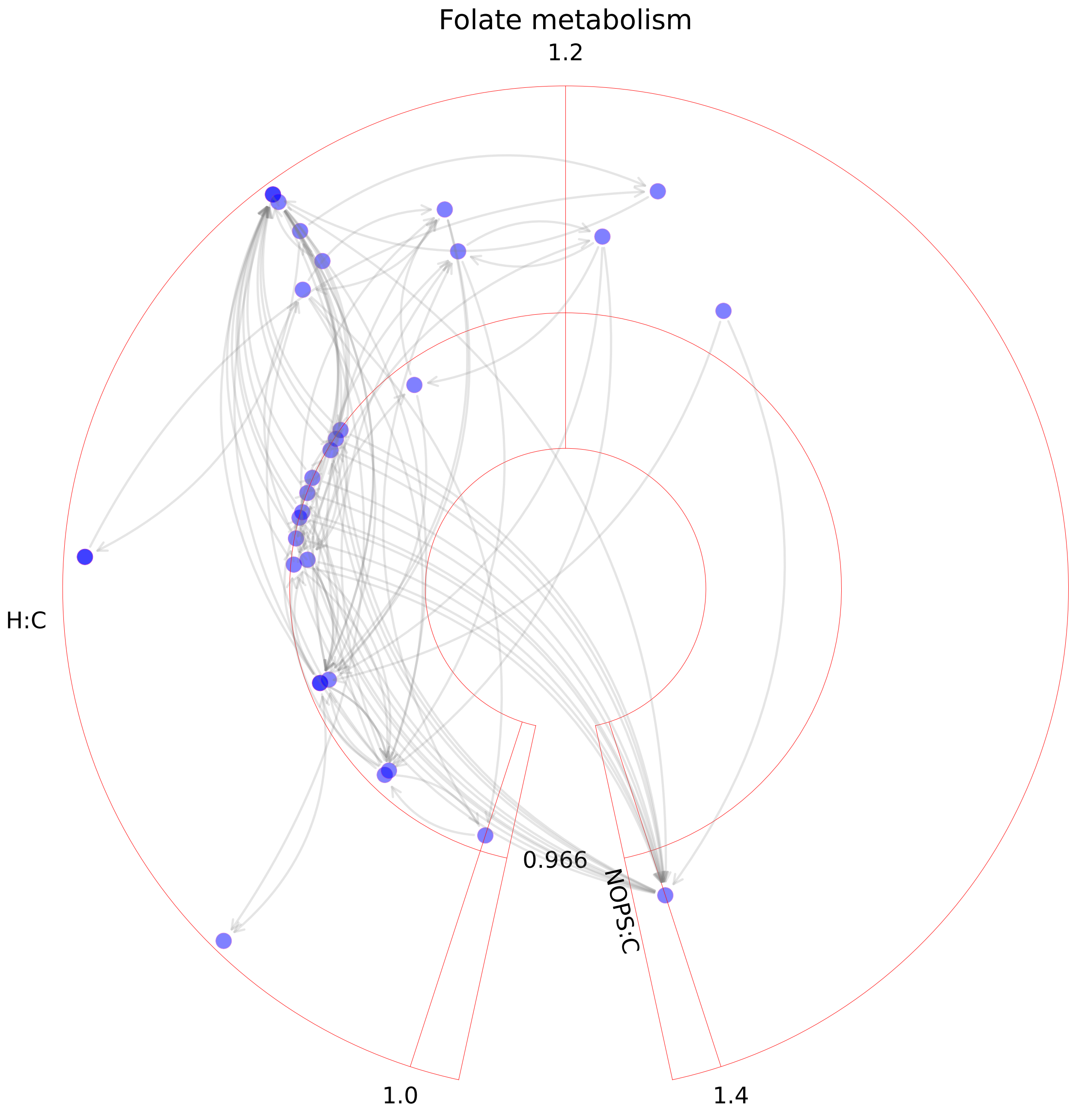

### Fructose and mannose metabolism.pdf

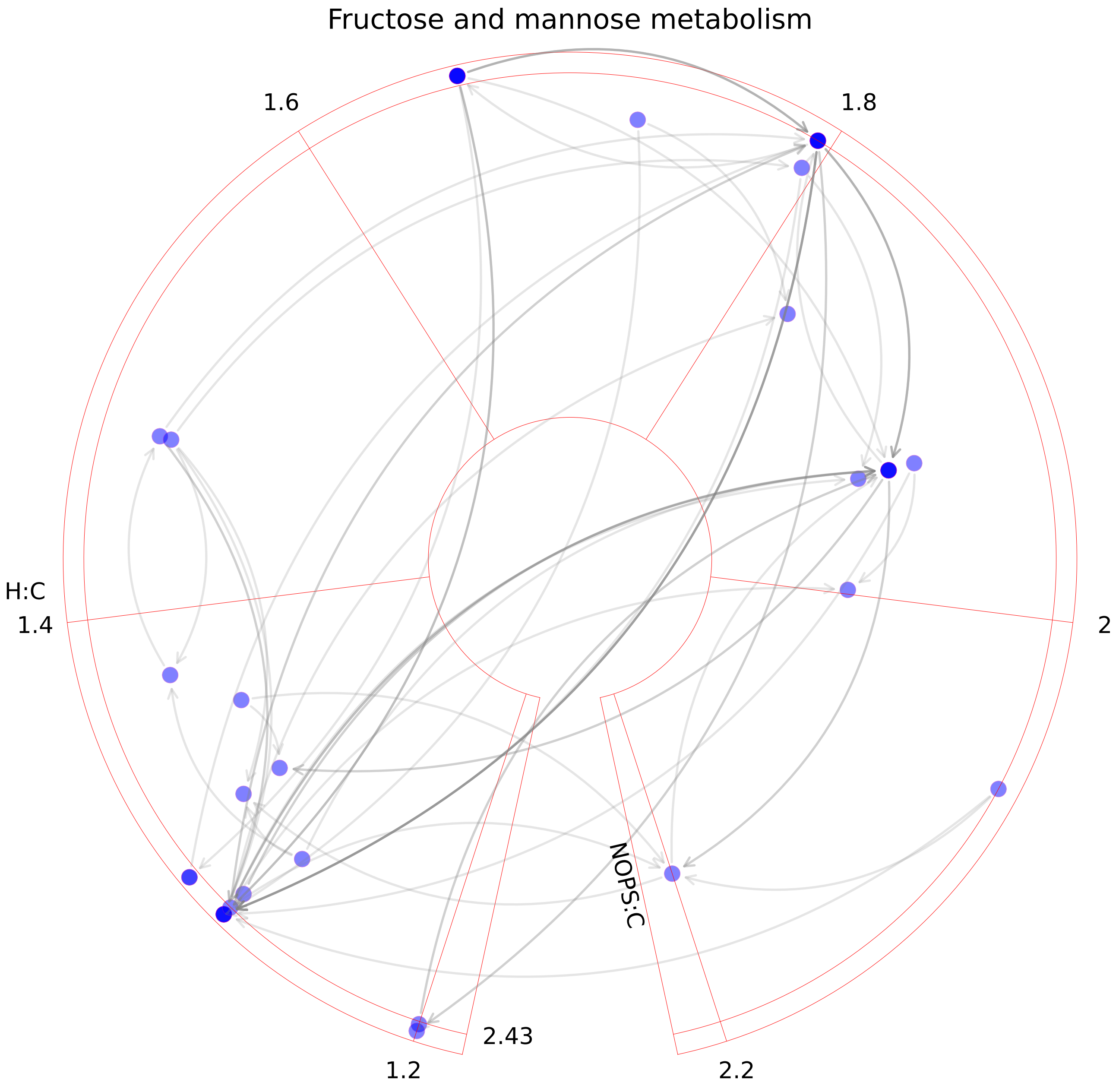

### Galactose metabolism.pdf

##### Galactose metabolism

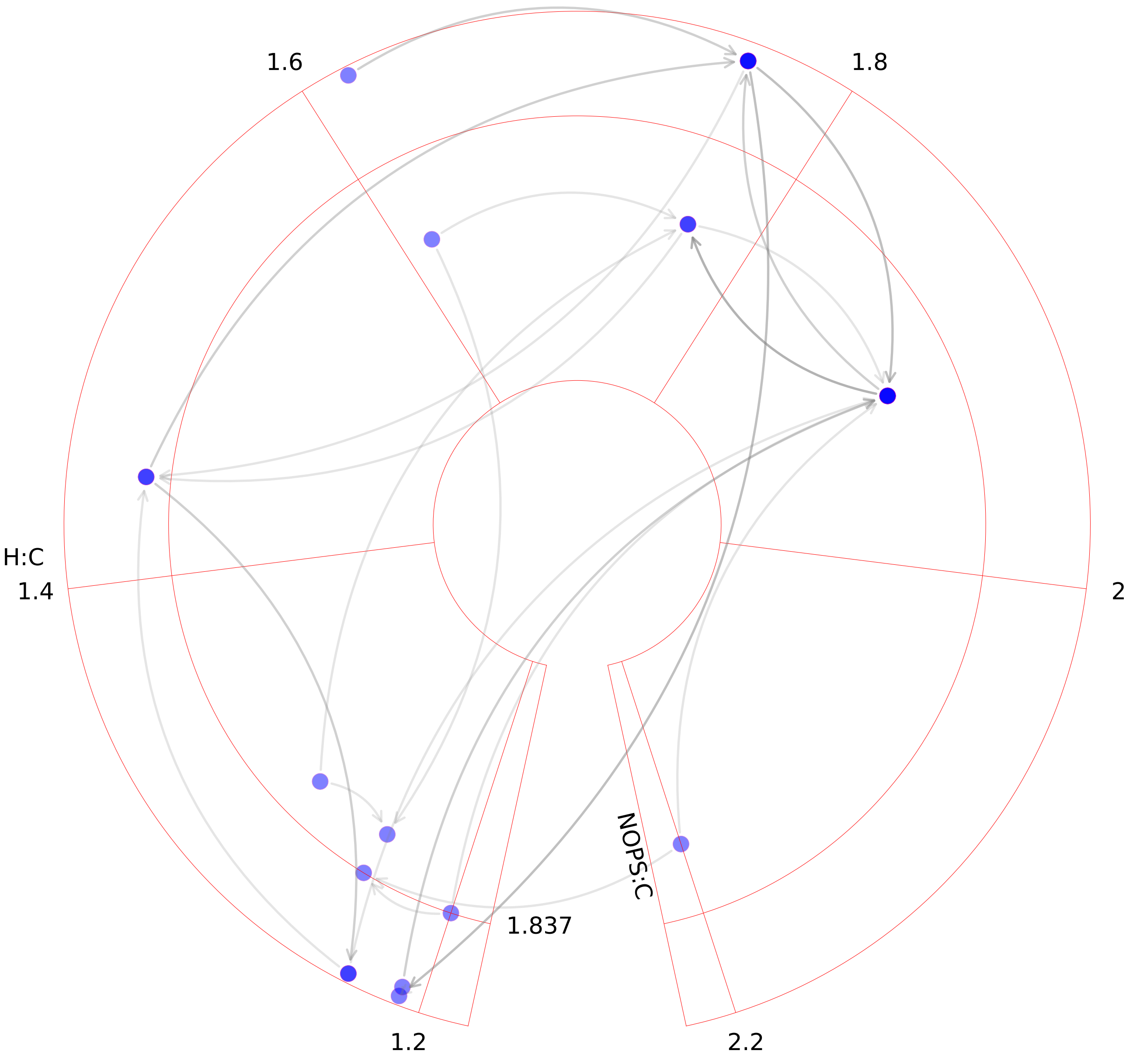

### Glutamate metabolism.pdf

#### Glutamate metabolism

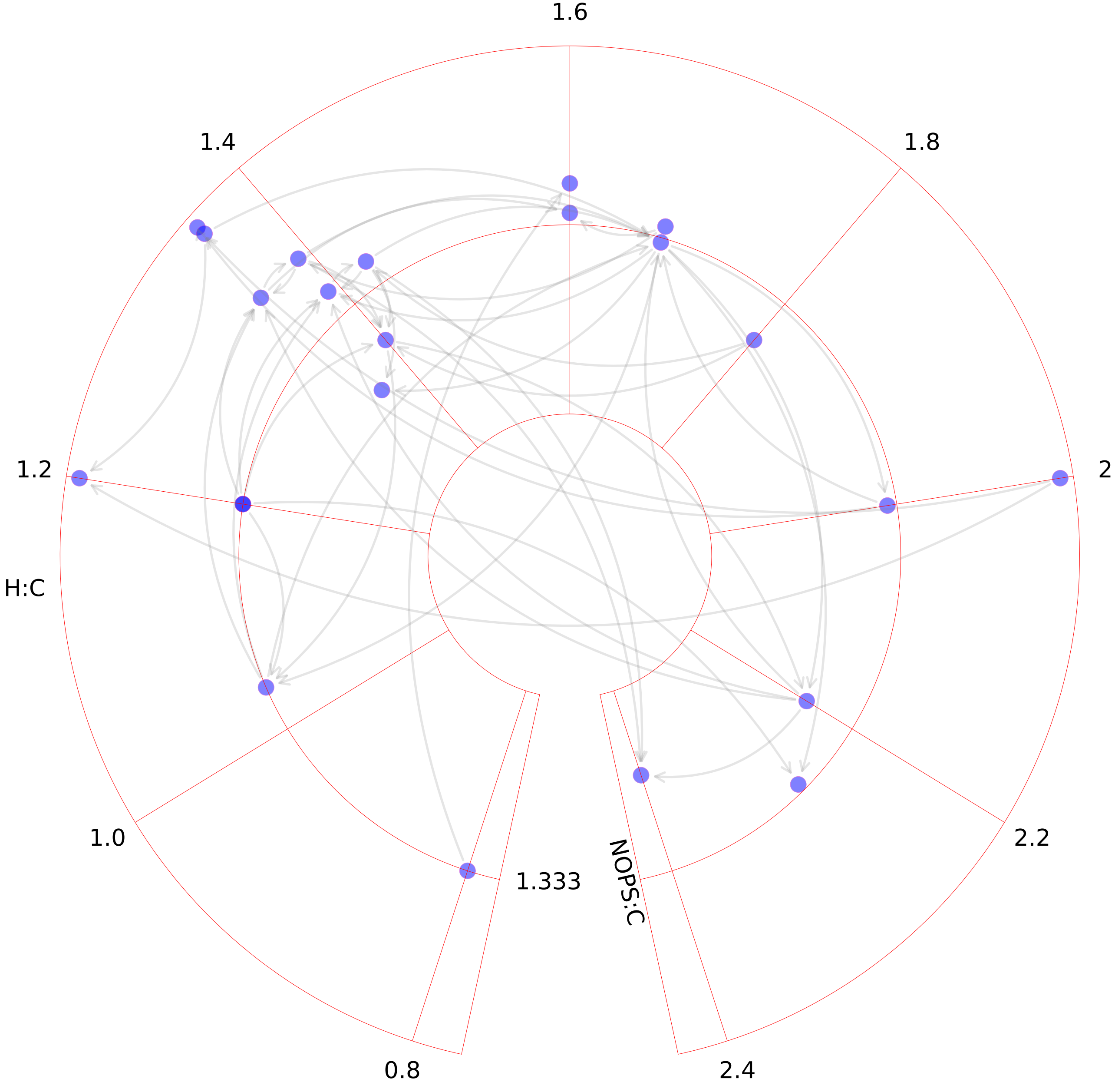

### Glutathione metabolism.pdf

#### Glutathione metabolism

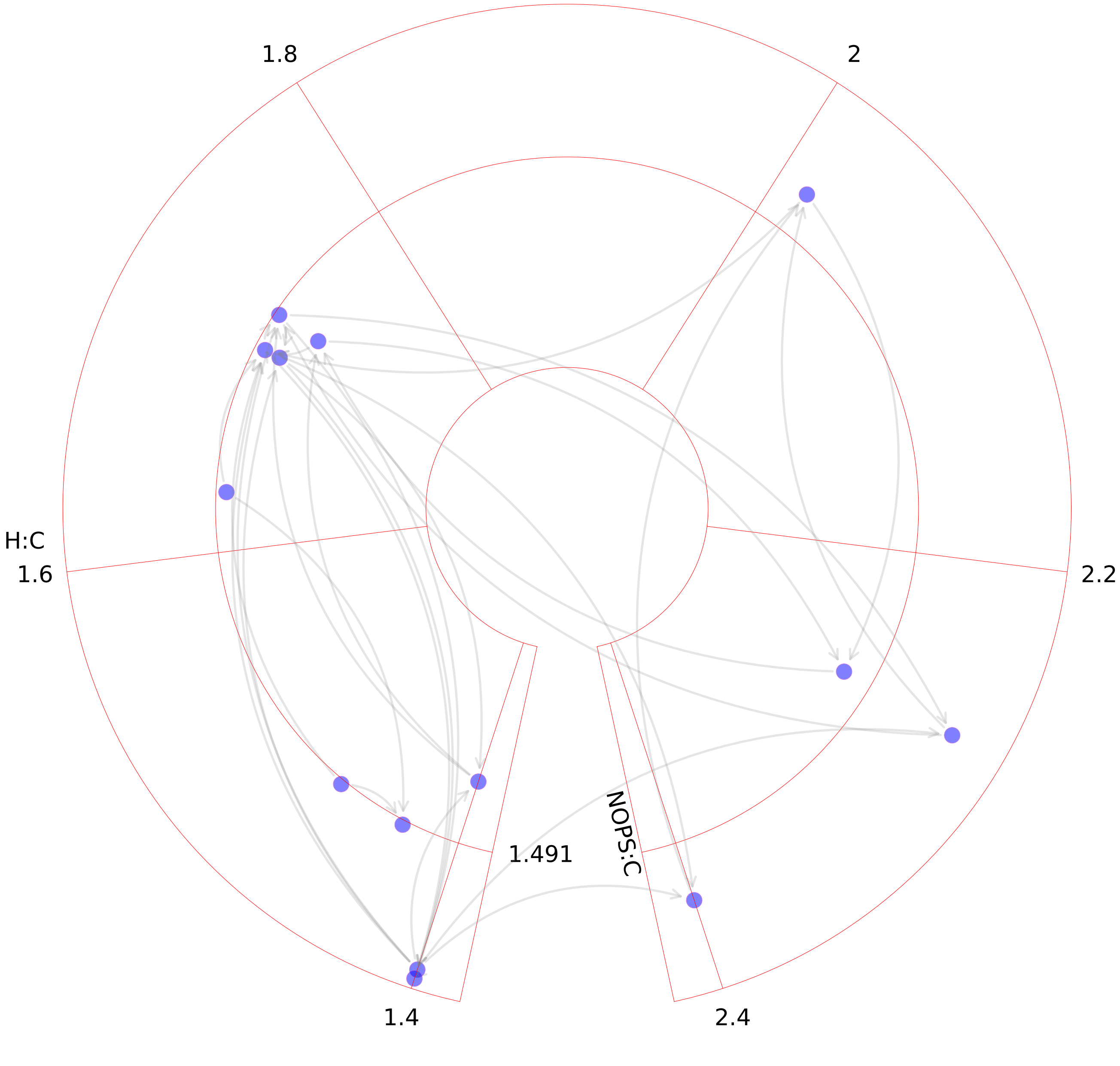

### Glycerophospholipid metabolism.pdf

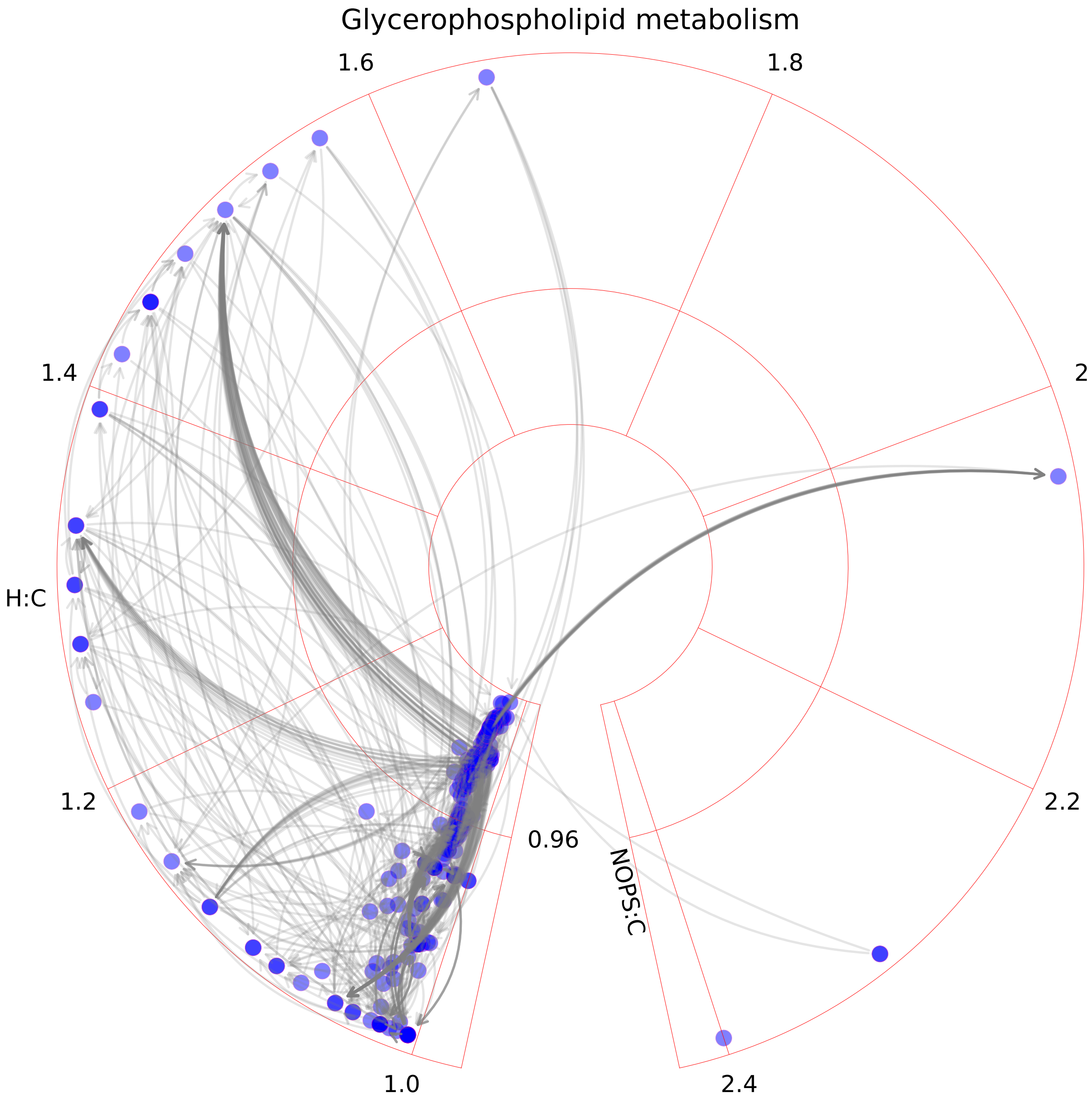

### Glycine, serine, alanine, and threonine metabolism.pdf

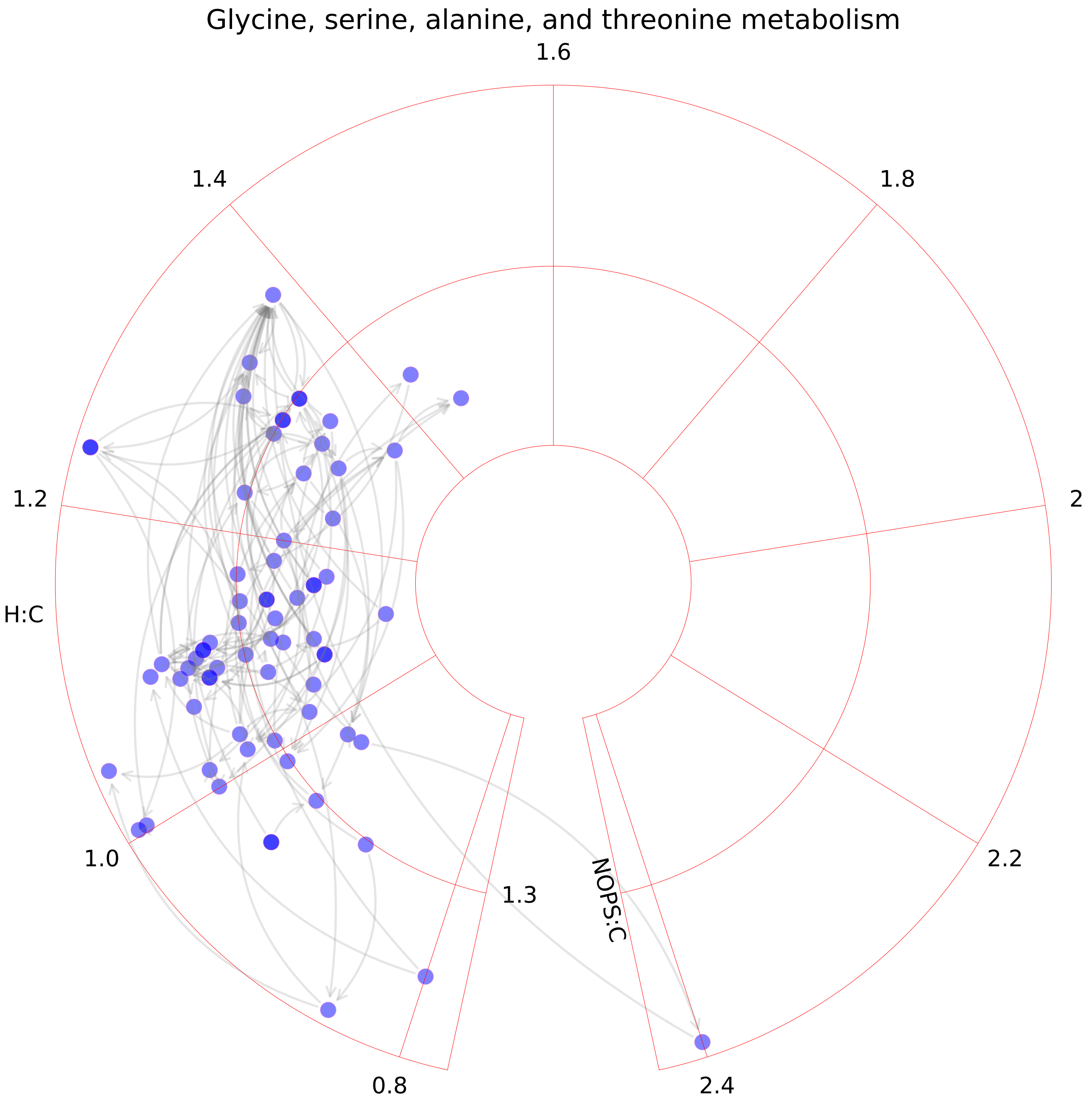

### Glycolysis_gluconeogenesis.pdf

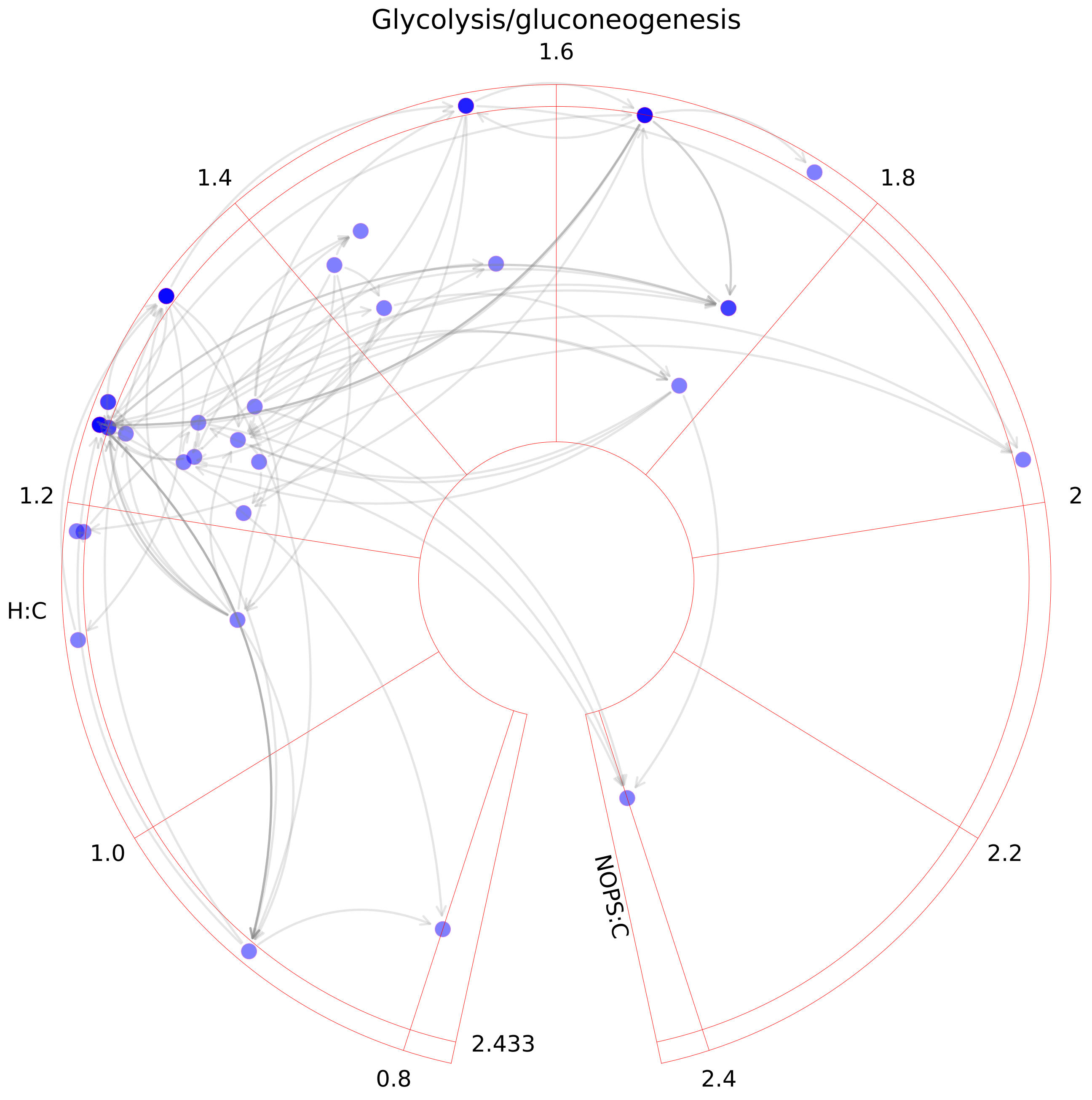

### Glycosphingolipid metabolism.pdf

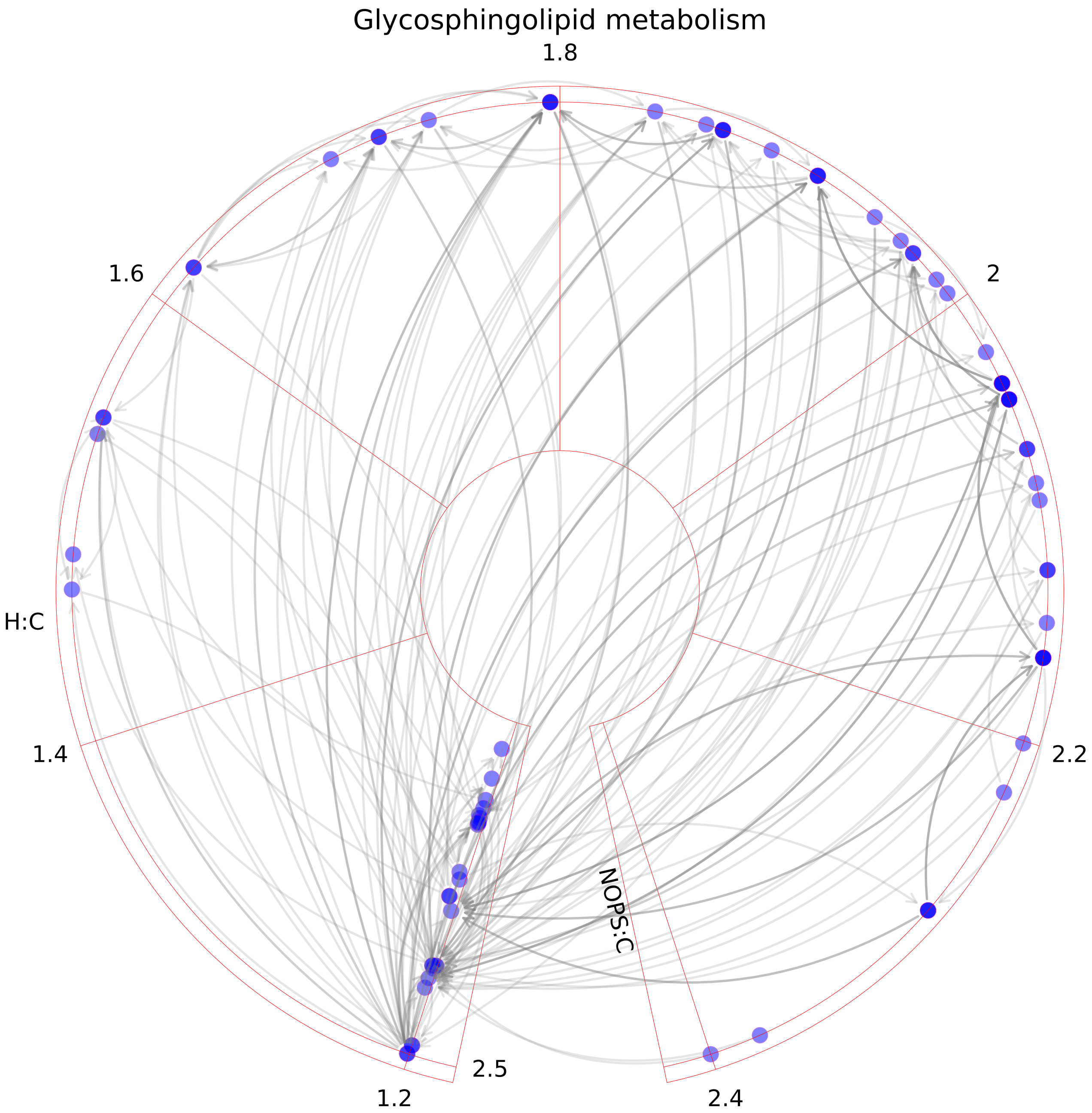

### Glyoxylate and dicarboxylate metabolism.pdf

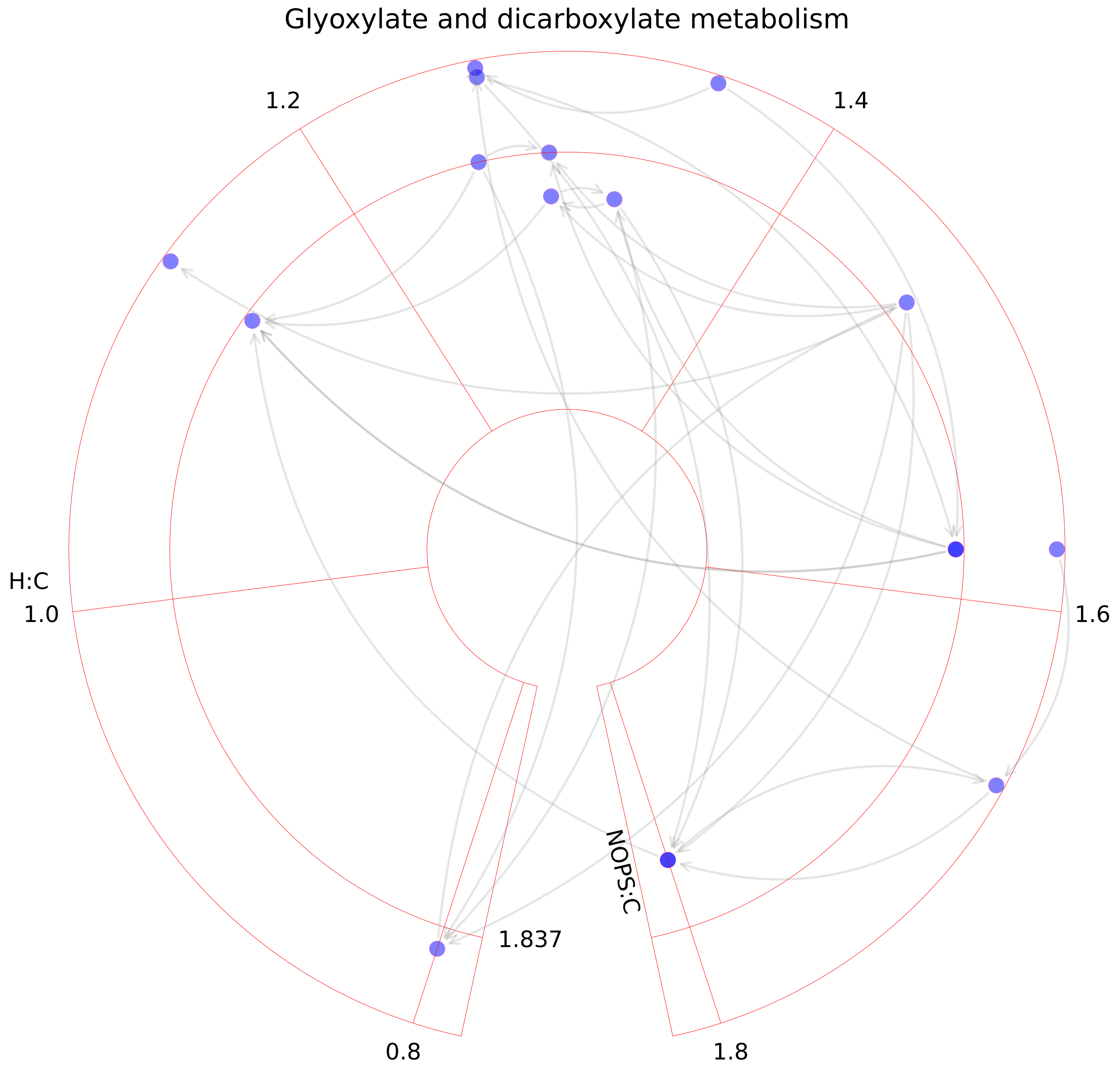

### Heme degradation.pdf

##### Heme degradation

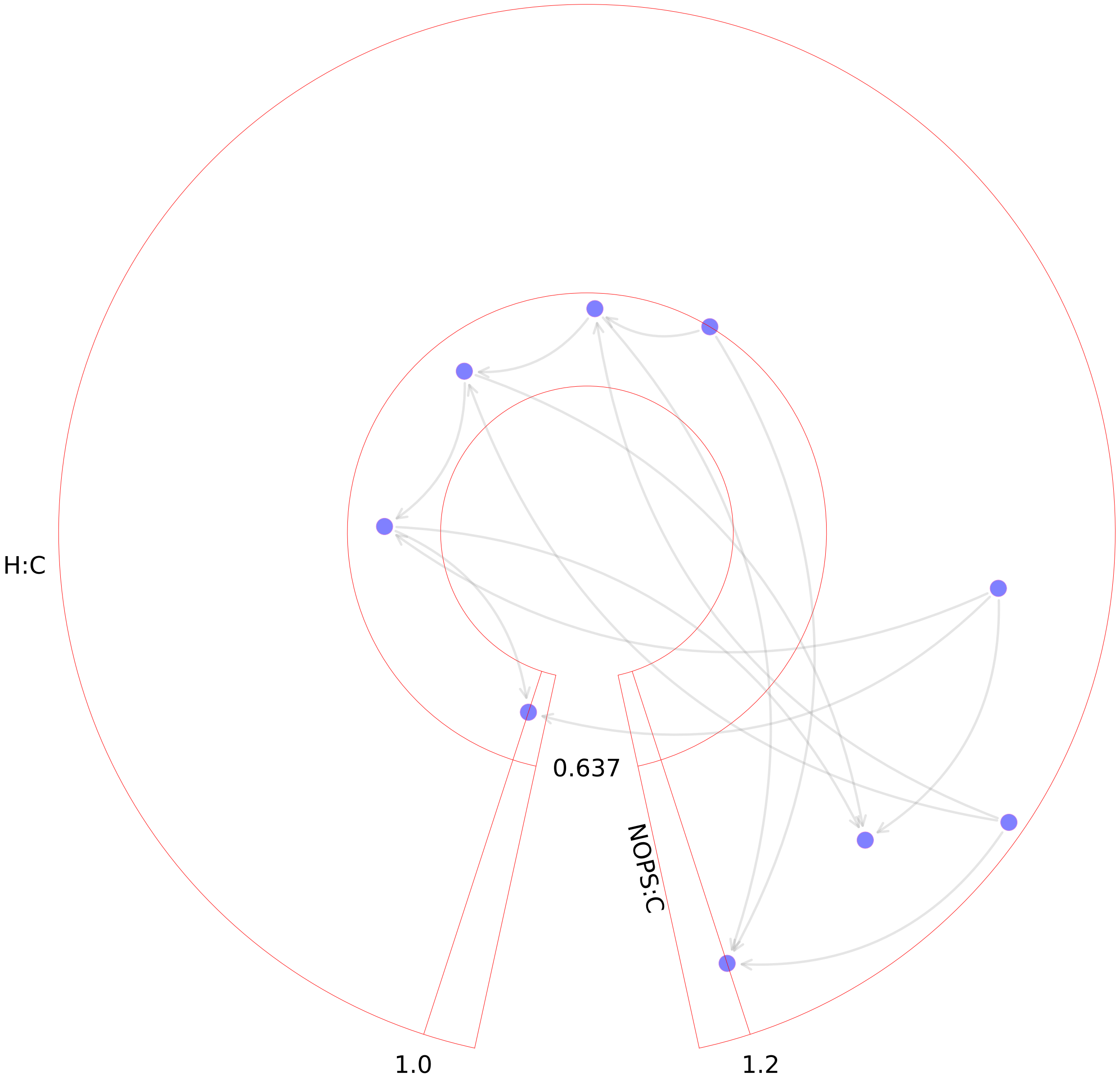

### Heme synthesis.pdf

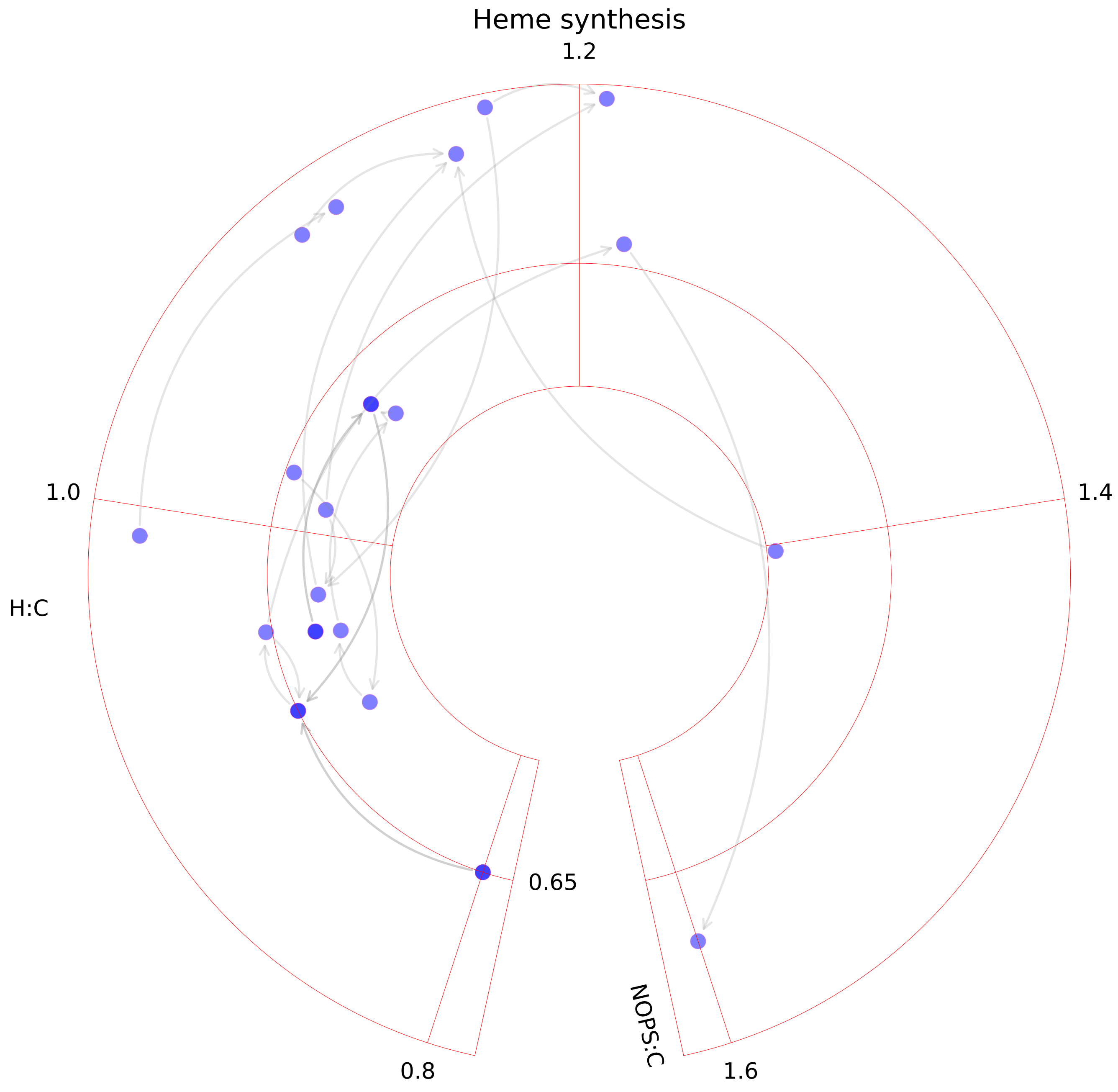

### Heparan sulfate degradation.pdf

#### Heparan sulfate degradation

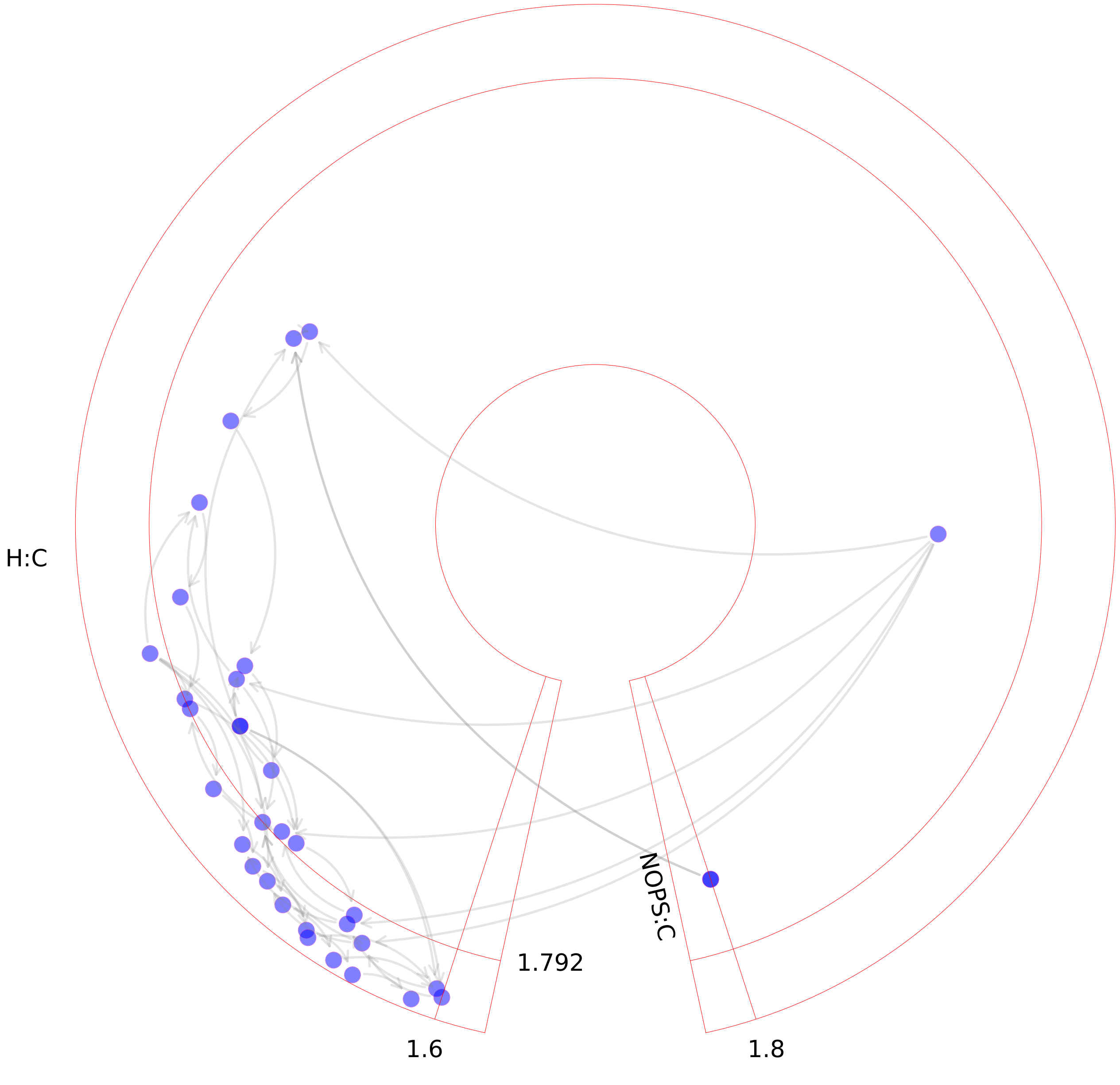

### Hippurate metabolism.pdf

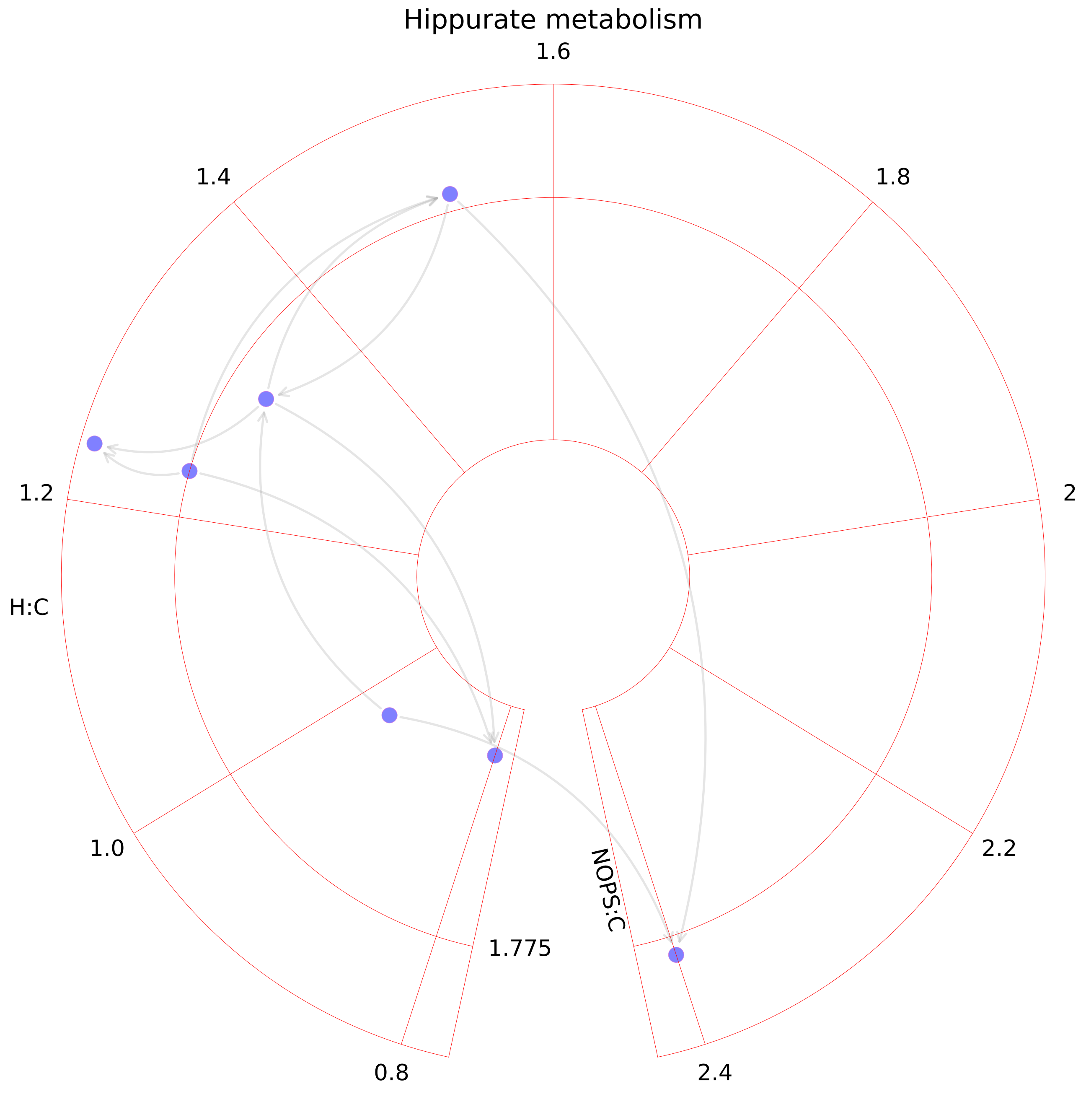

### Histidine metabolism.pdf

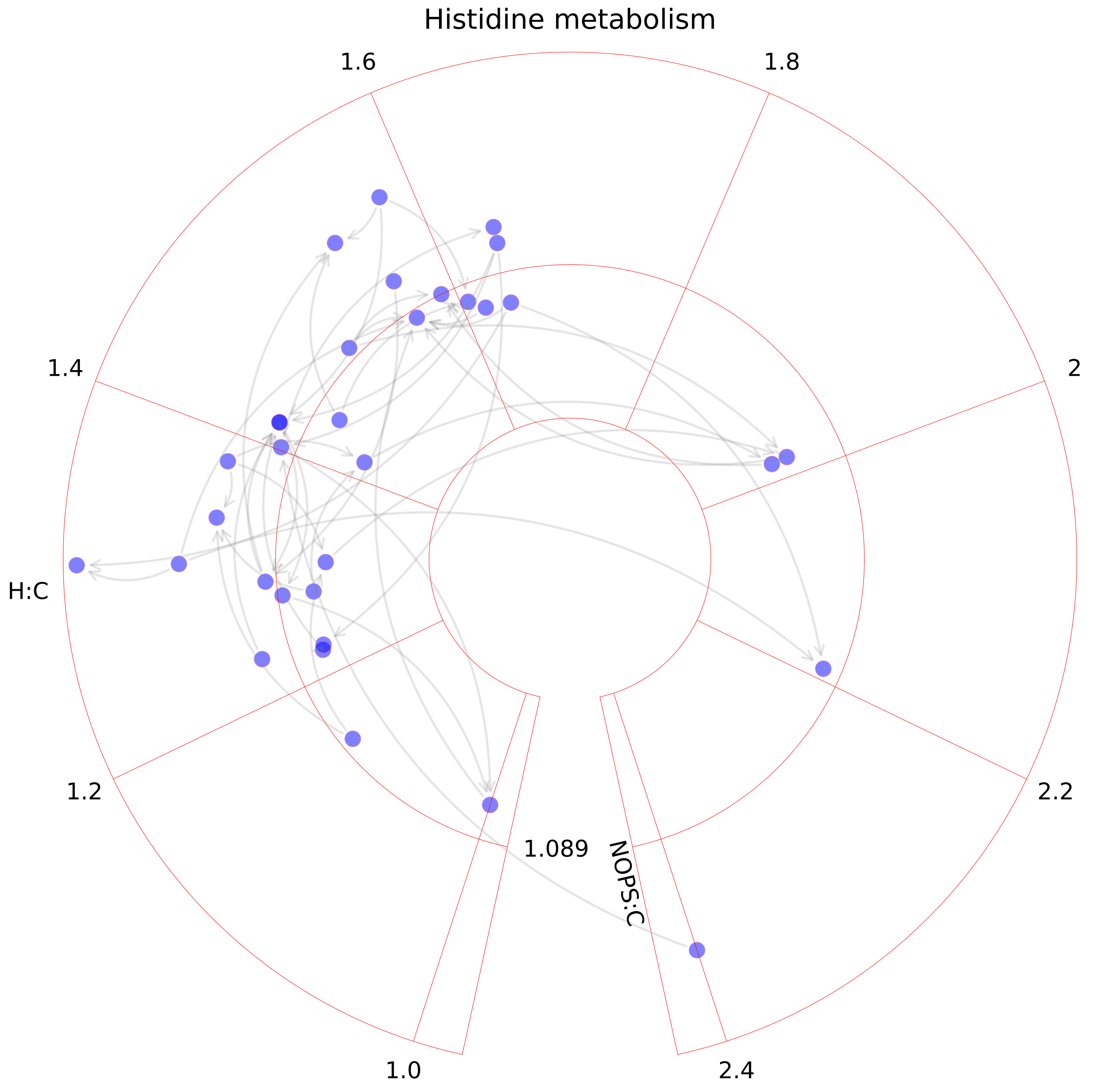

### Hyaluronan metabolism.pdf

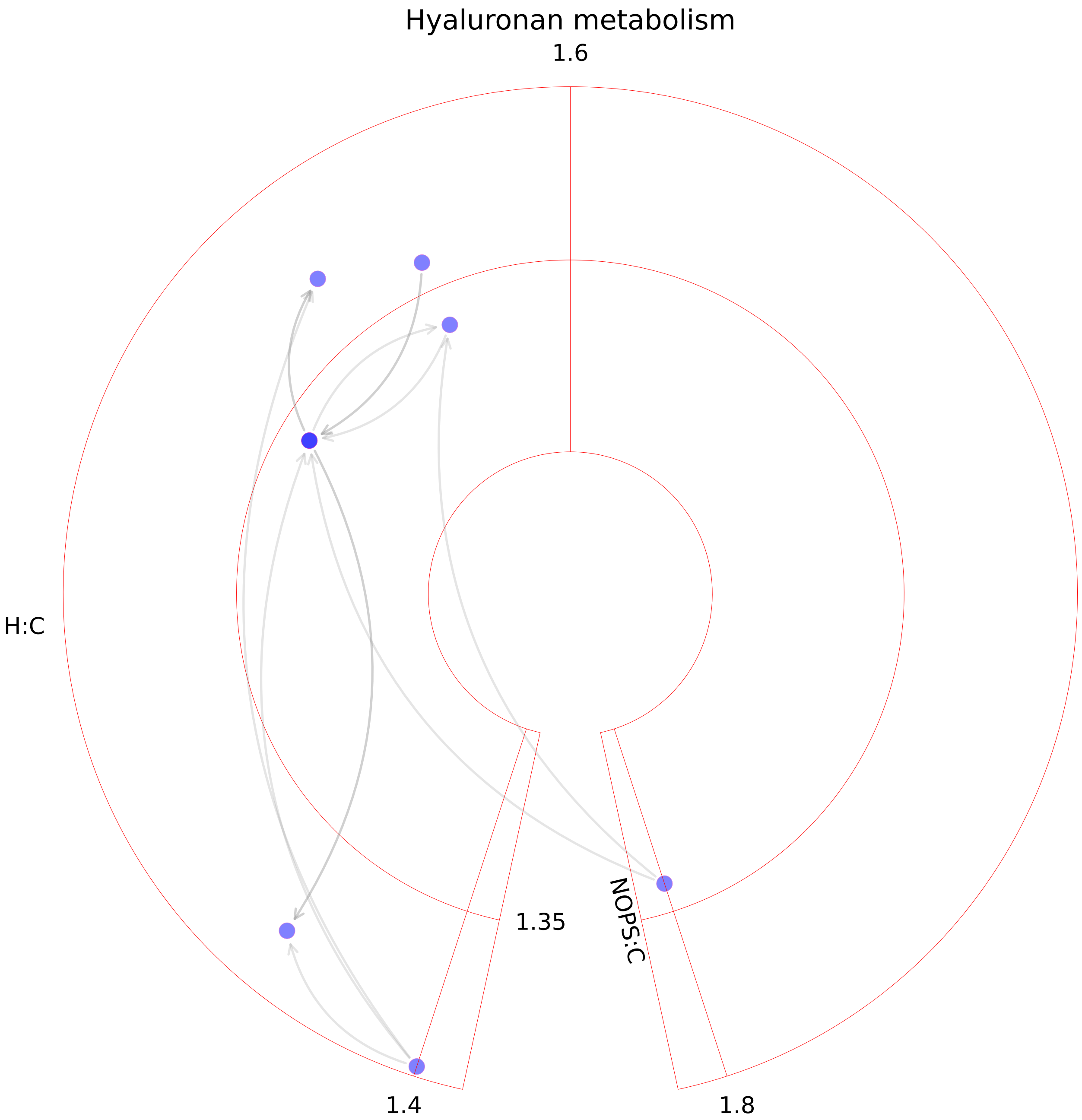

### Inositol phosphate metabolism.pdf

Inositol phosphate metabolism

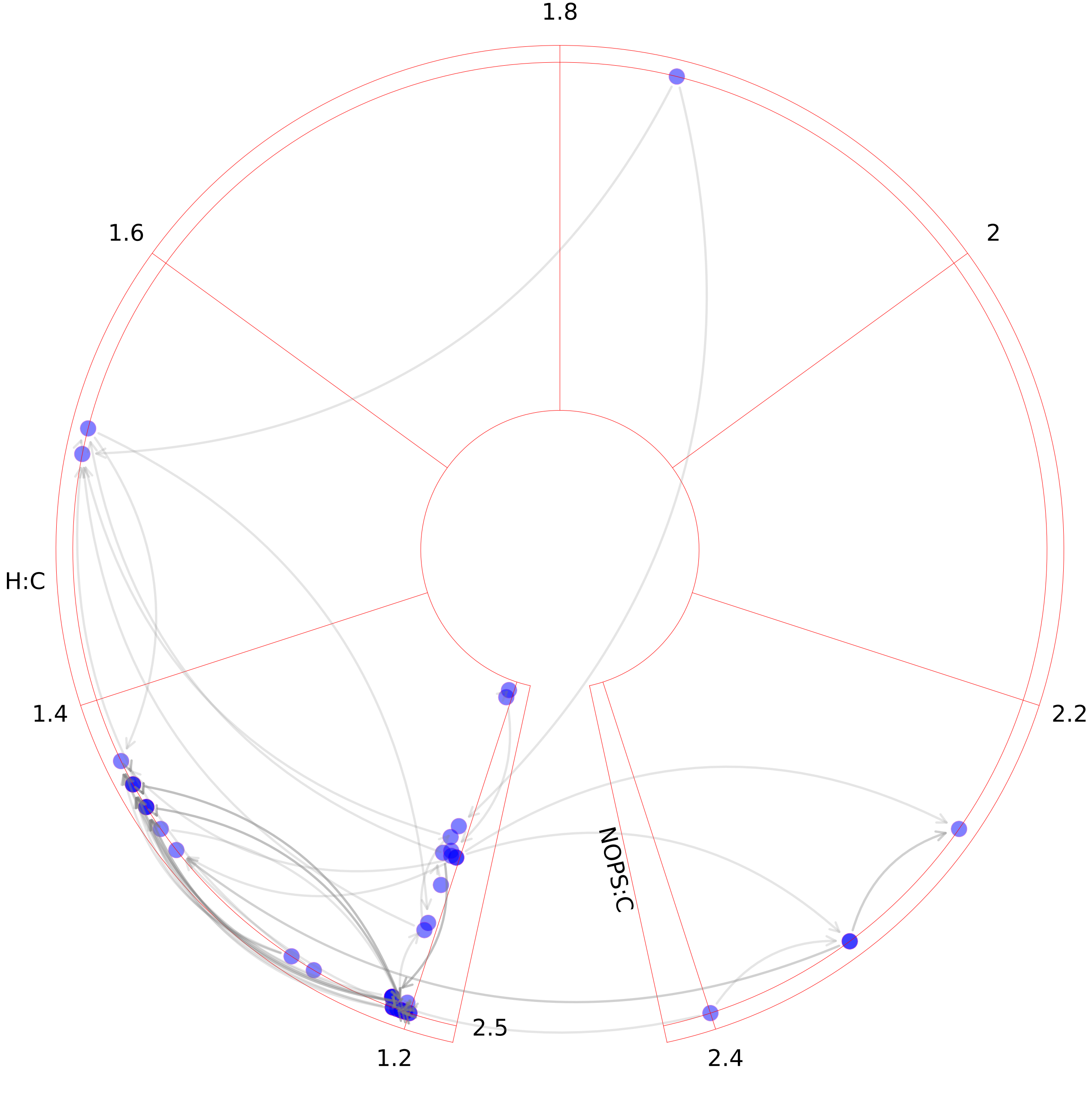

### Keratan sulfate degradation.pdf

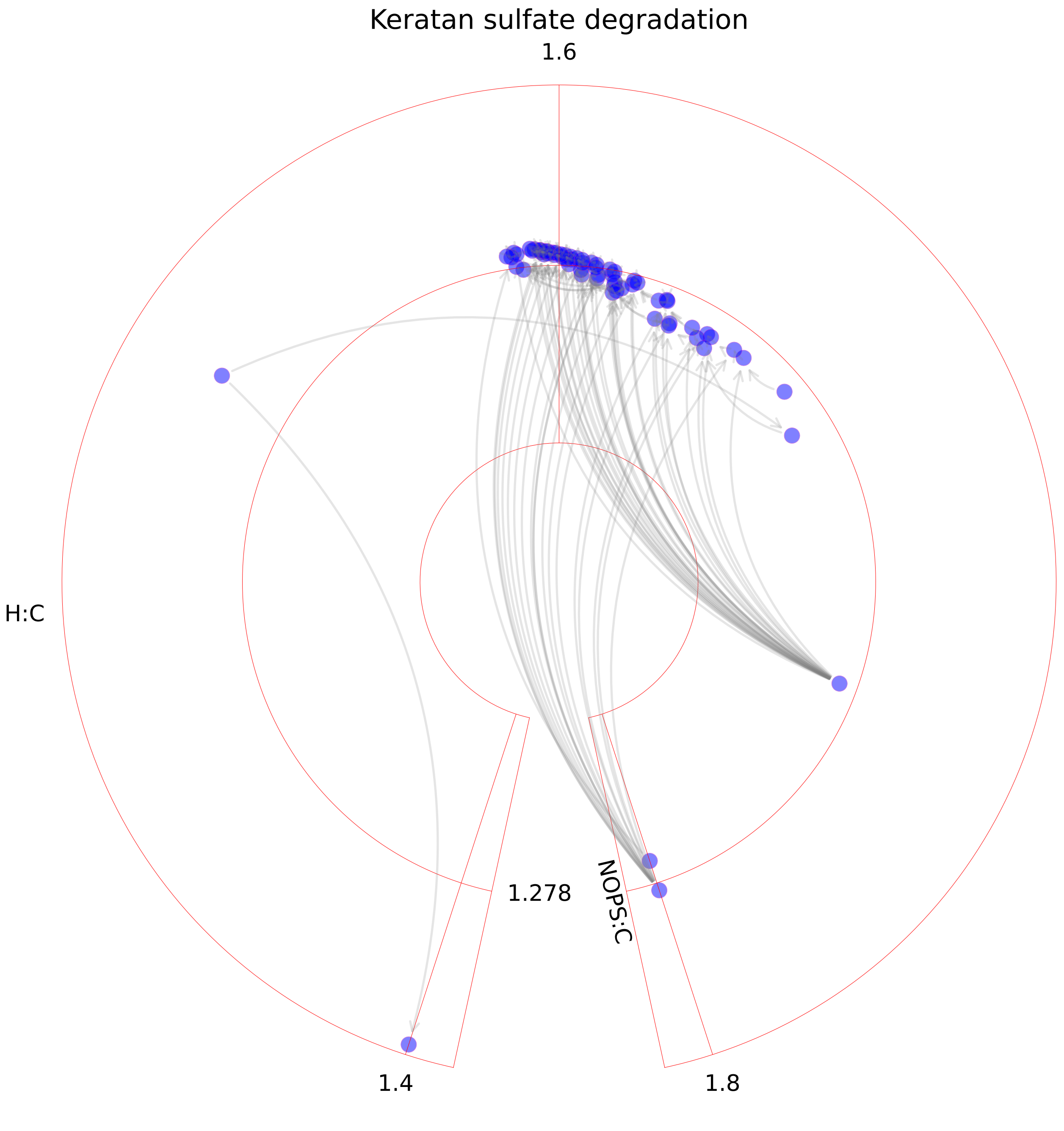

### Keratan sulfate degradation.pdf

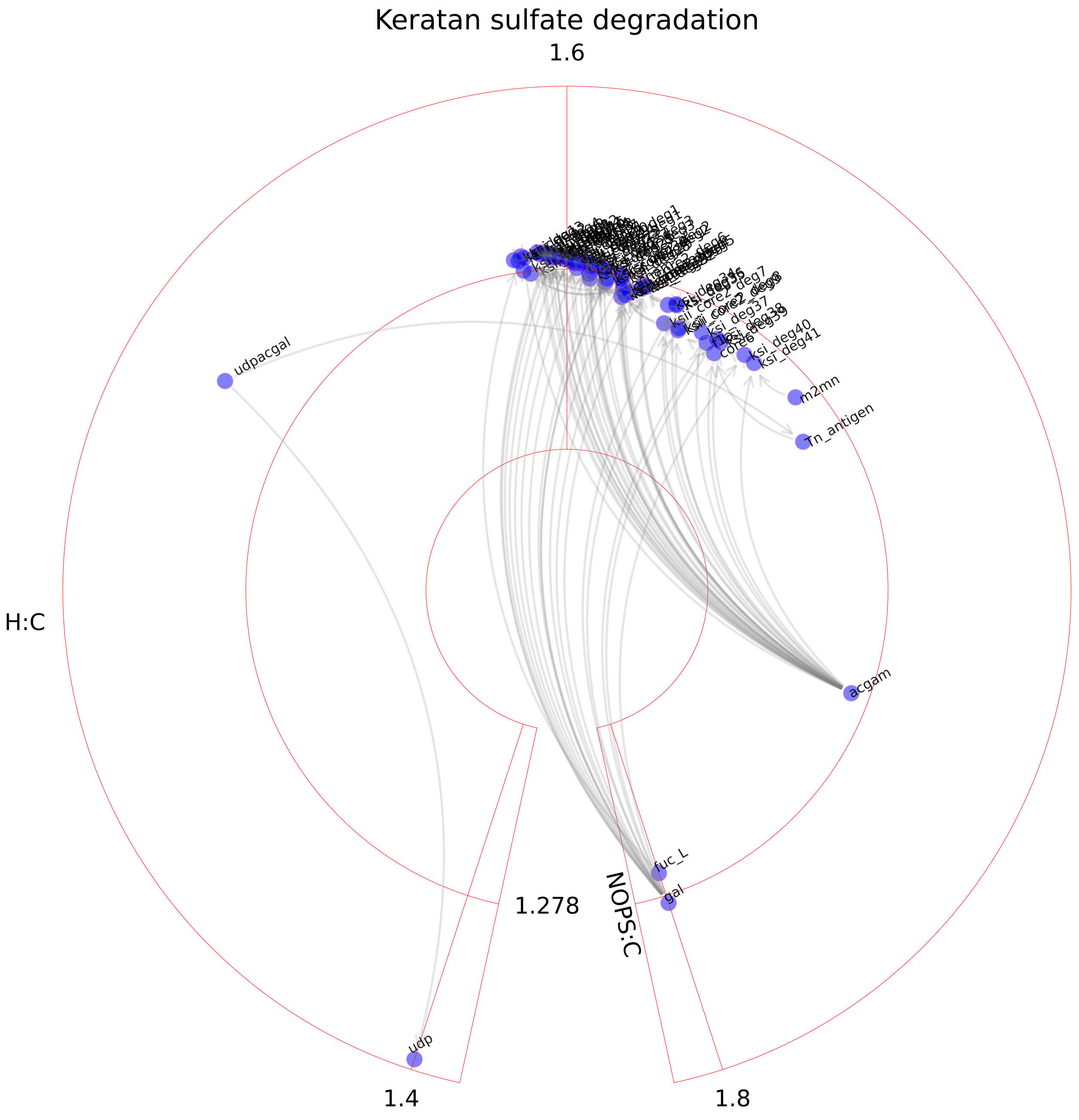

### Keratan sulfate synthesis.pdf

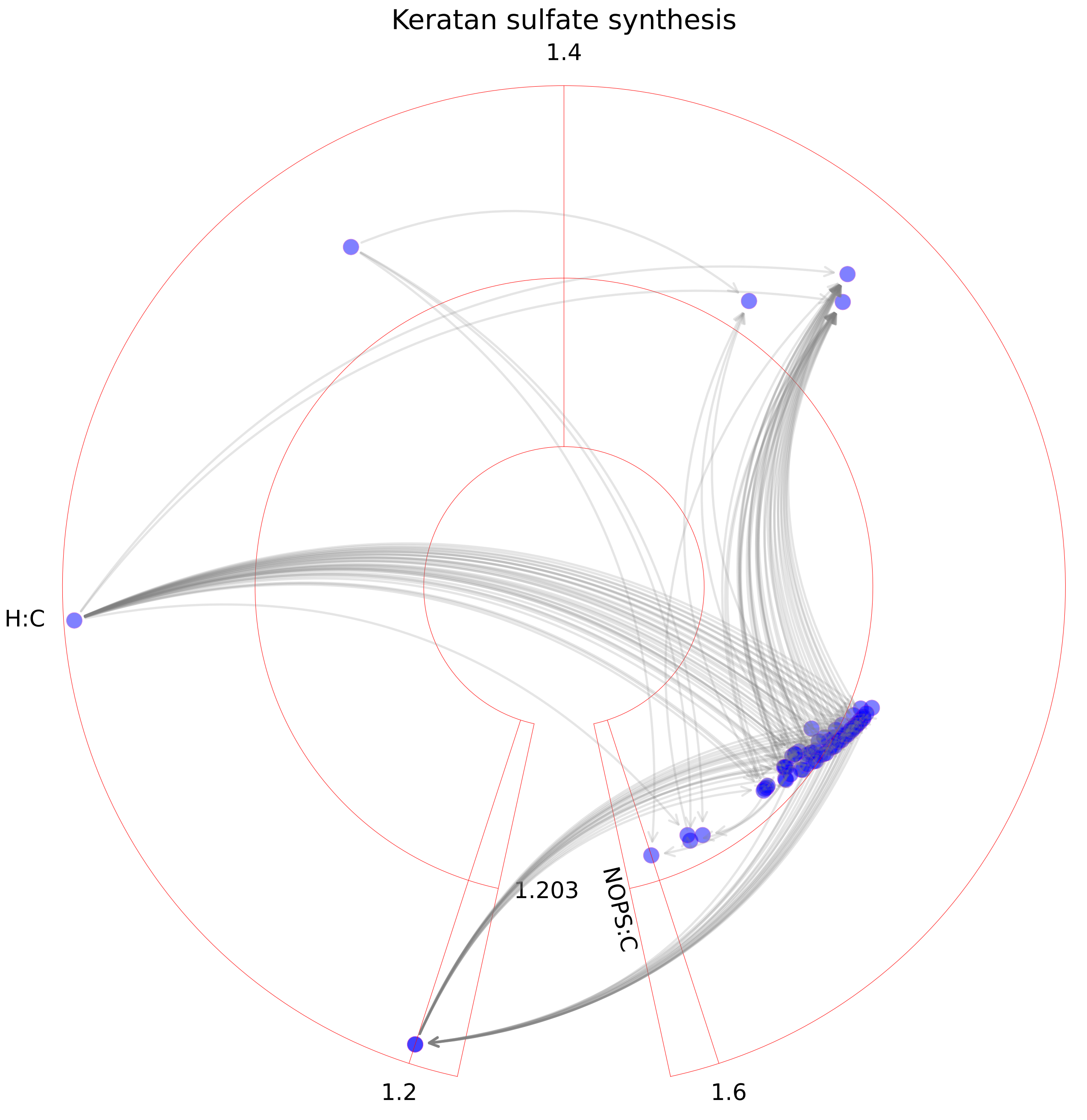

### Leukotriene metabolism.pdf

##### Leukotriene metabolism

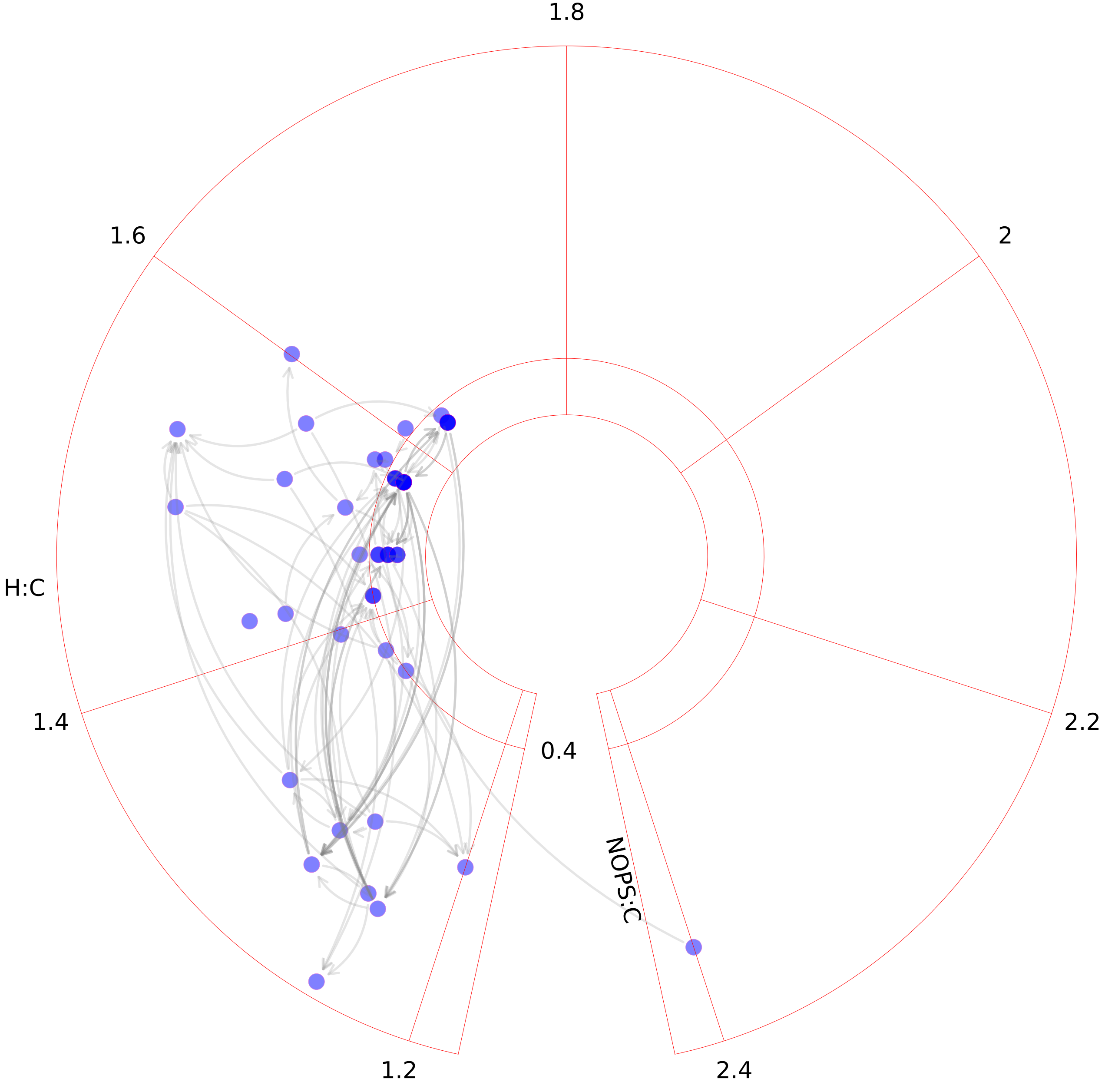

### Limonene and pinene degradation.pdf

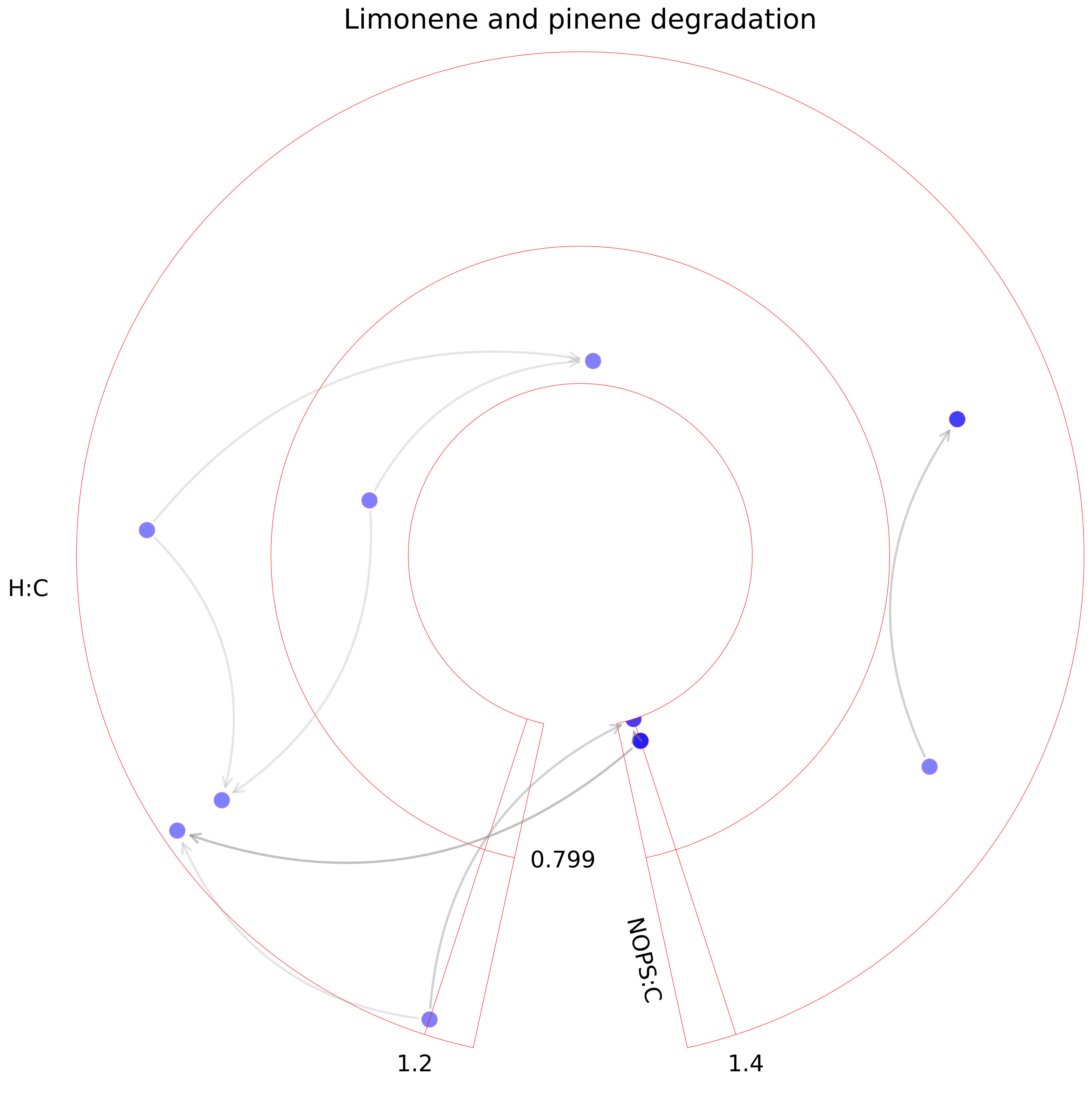

### Linoleate metabolism.pdf

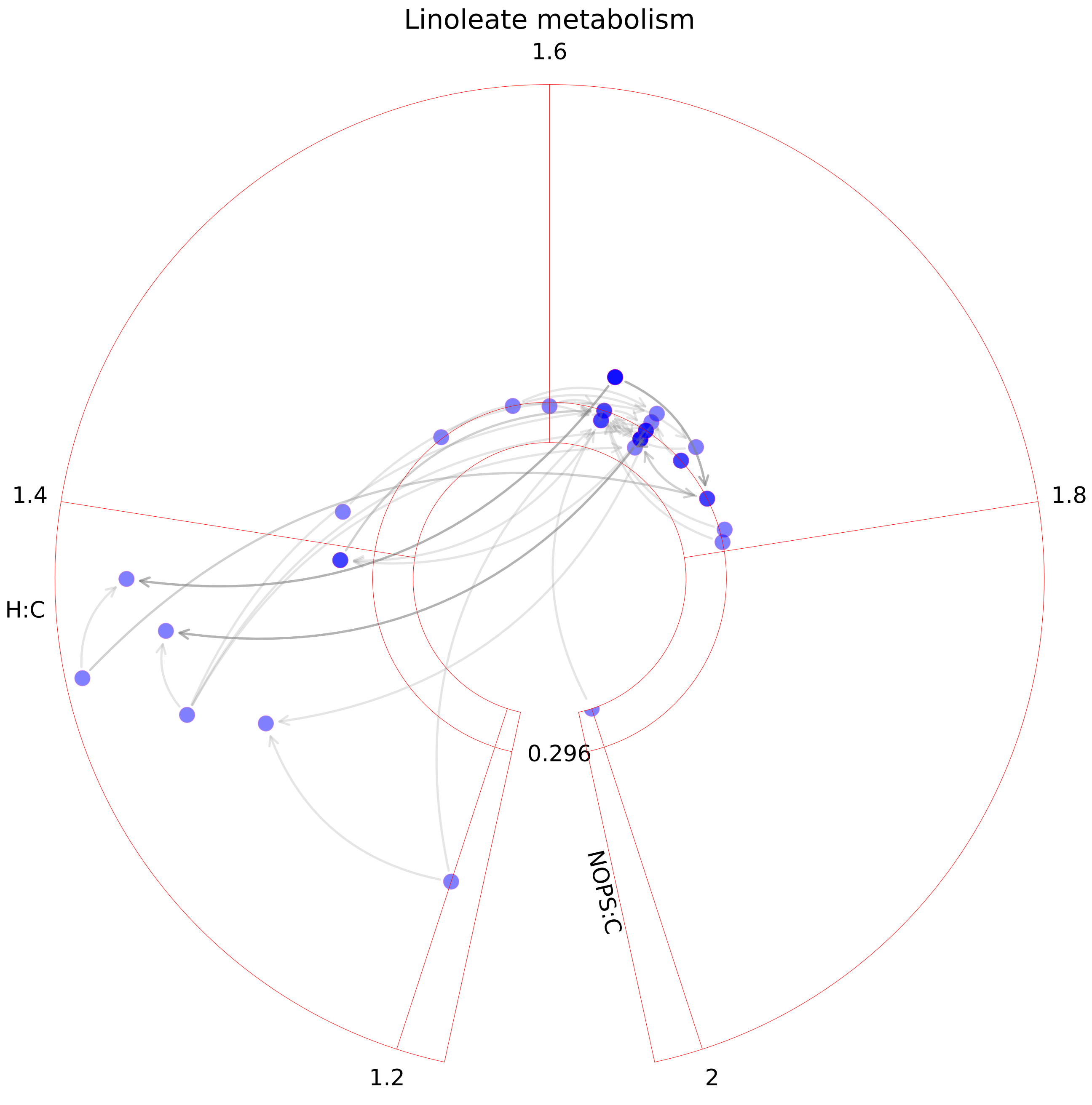

### Lipoate metabolism.pdf

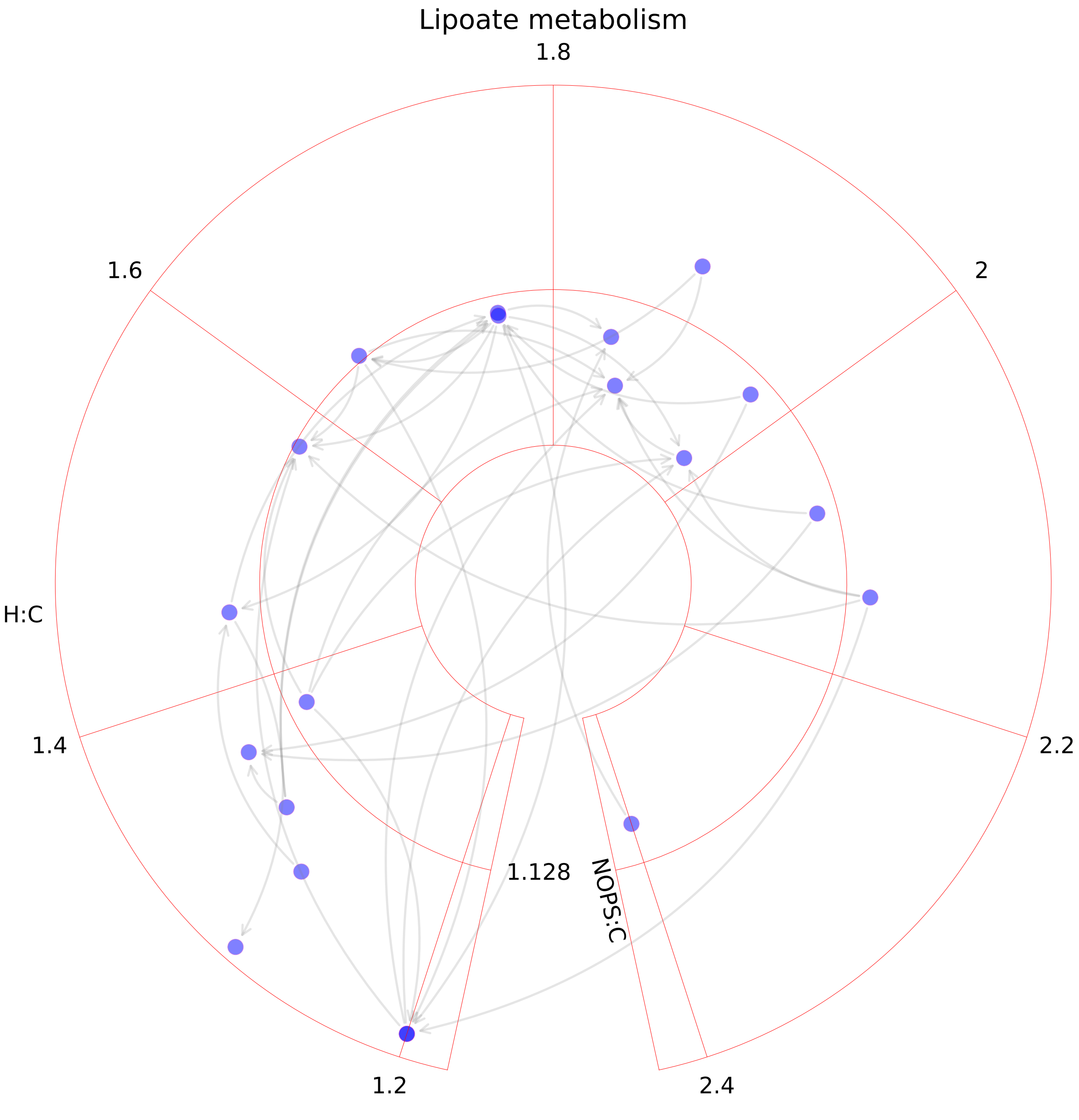

### Lysine metabolism.pdf

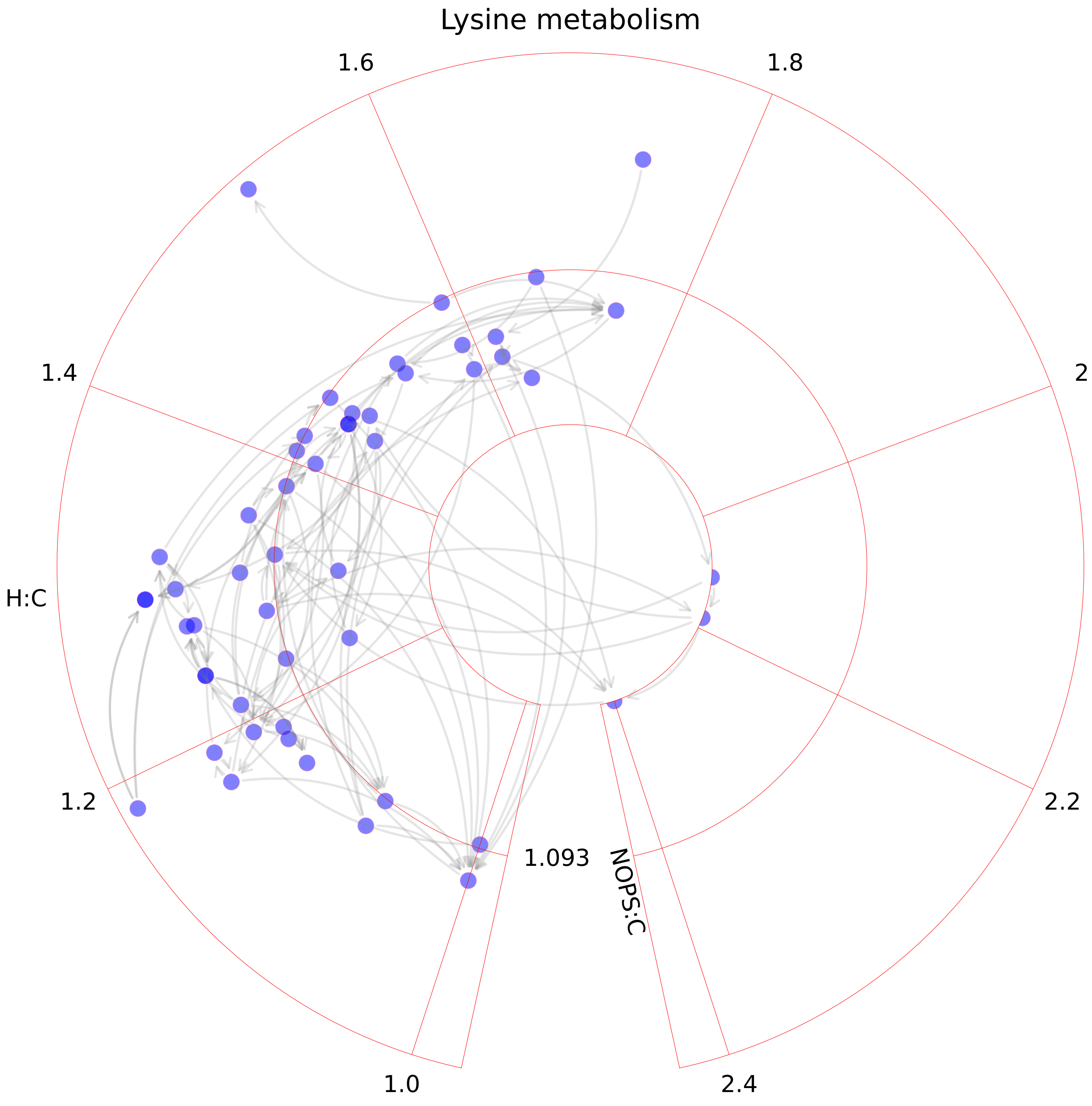

### Methionine and cysteine metabolism.pdf

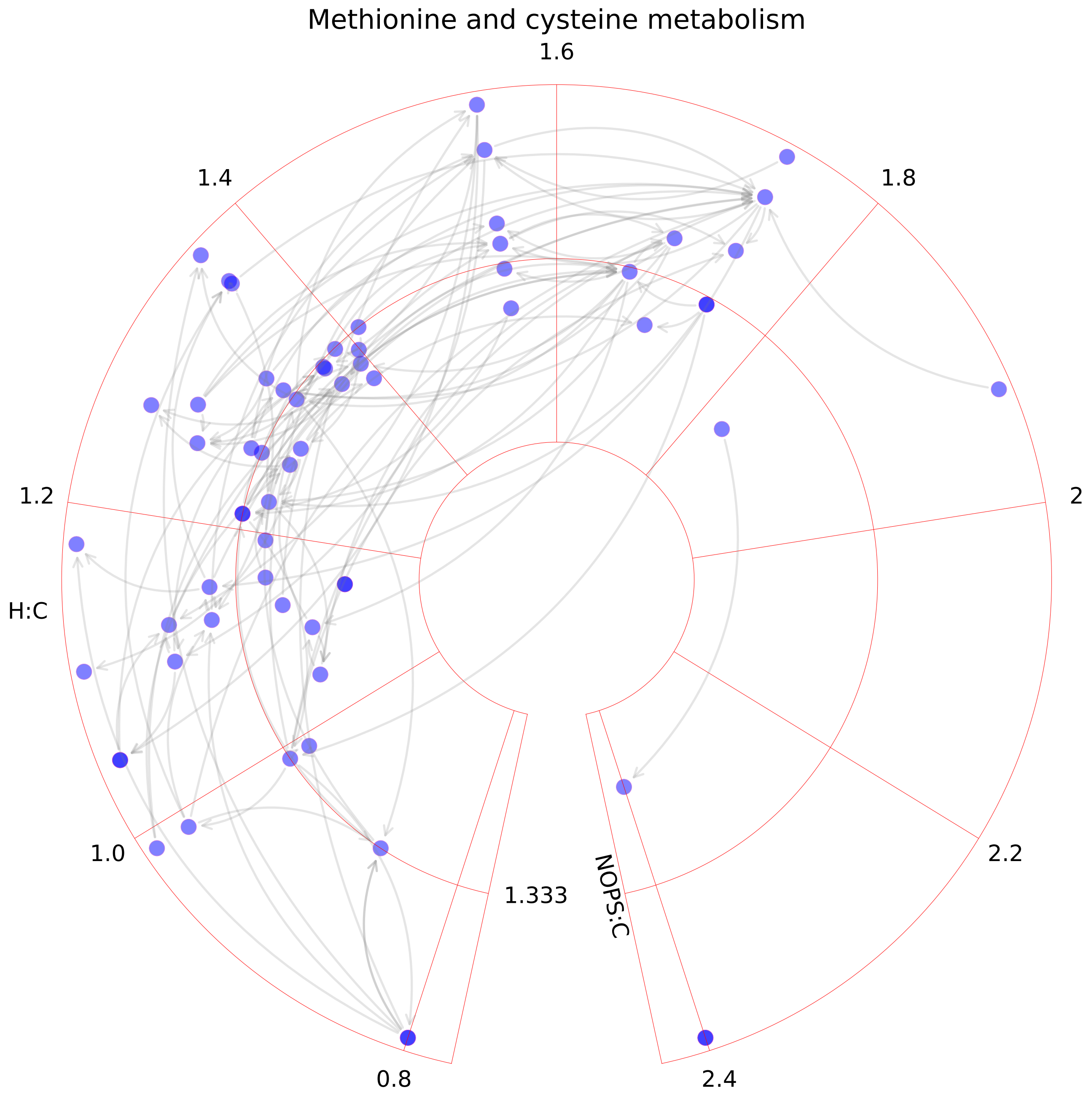

### N-glycan degradation.pdf

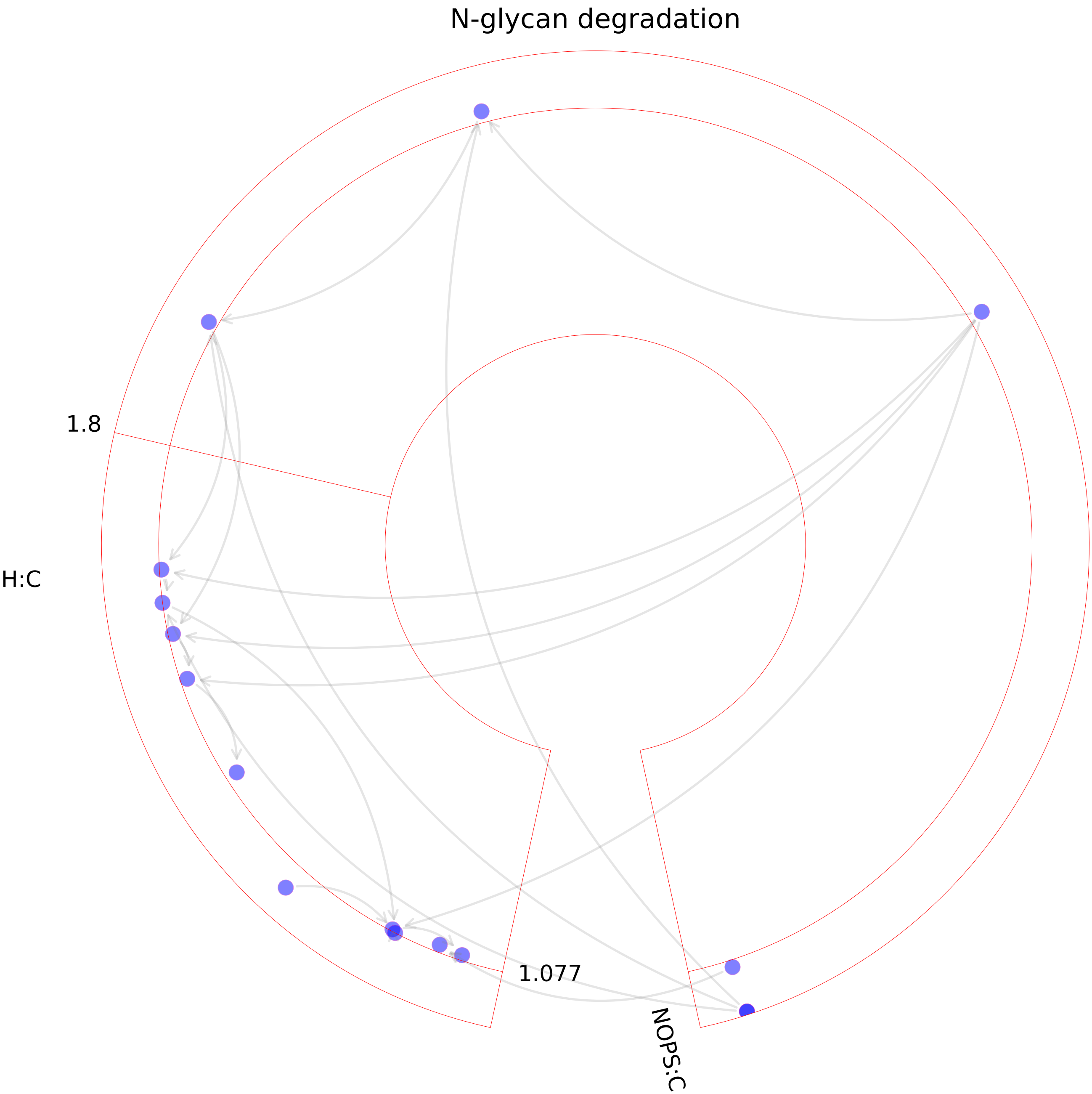

### N-glycan metabolism.pdf

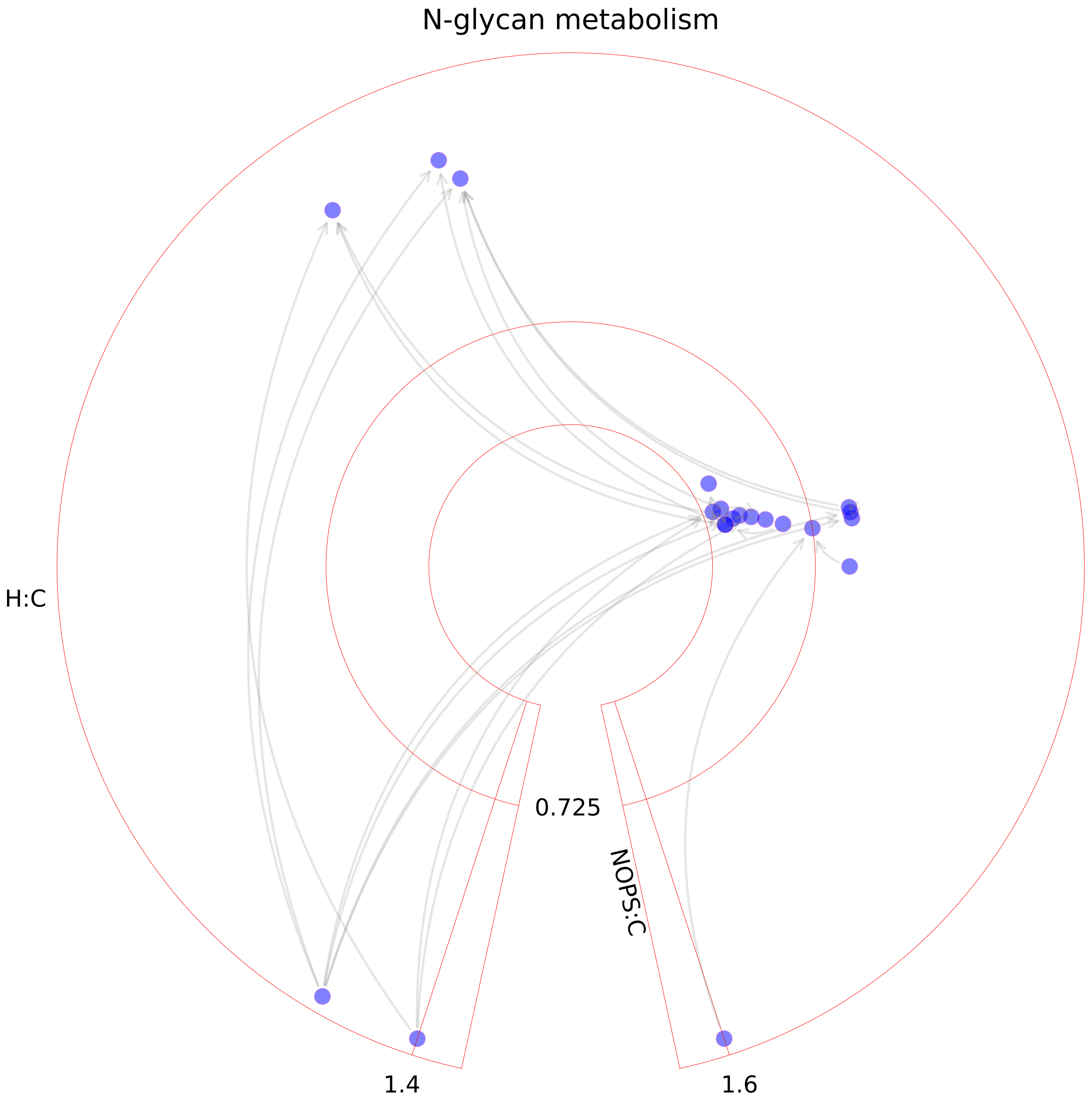

### N-glycan synthesis.pdf

#### N-glycan synthesis

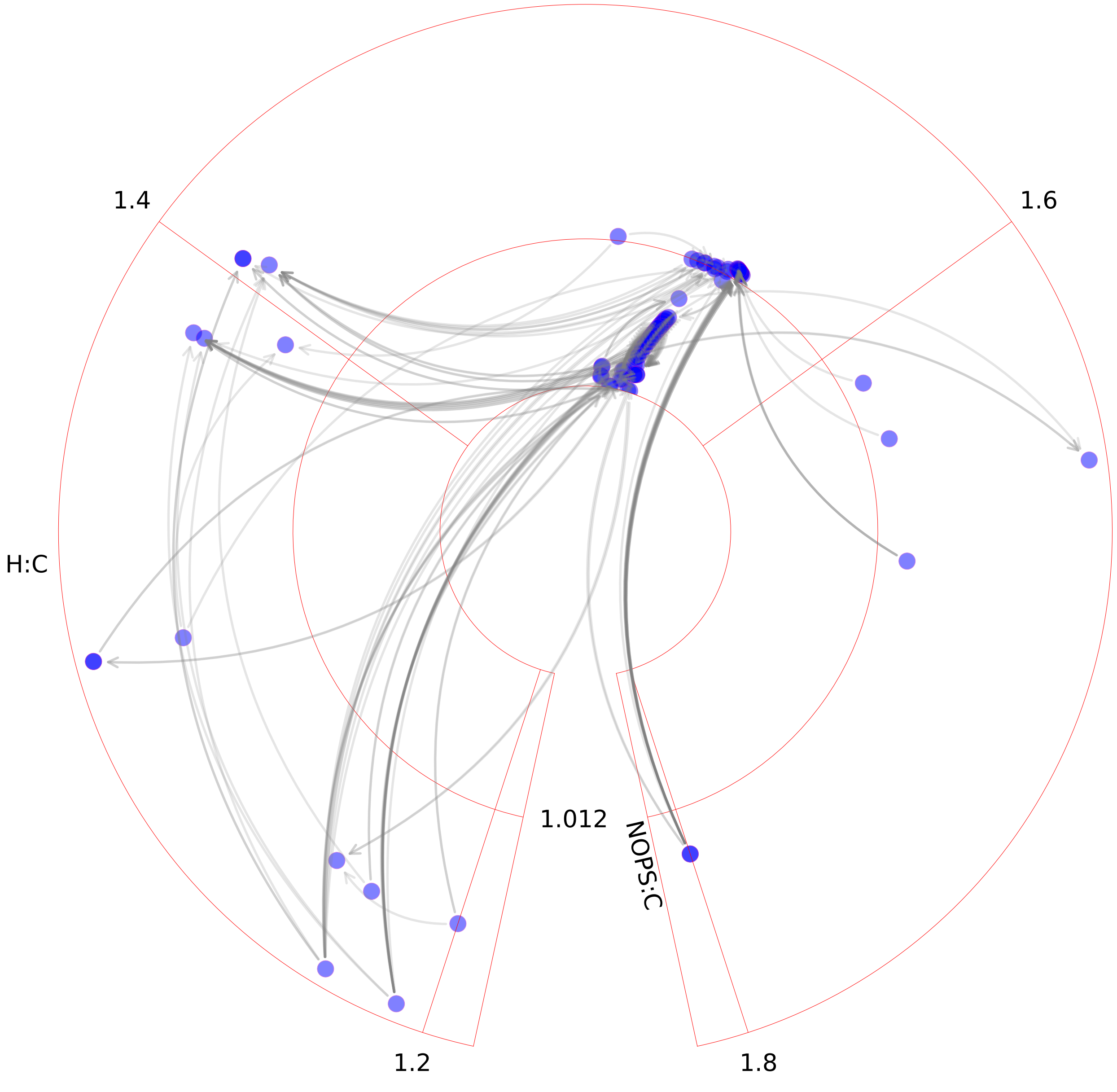

### NAD metabolism.pdf

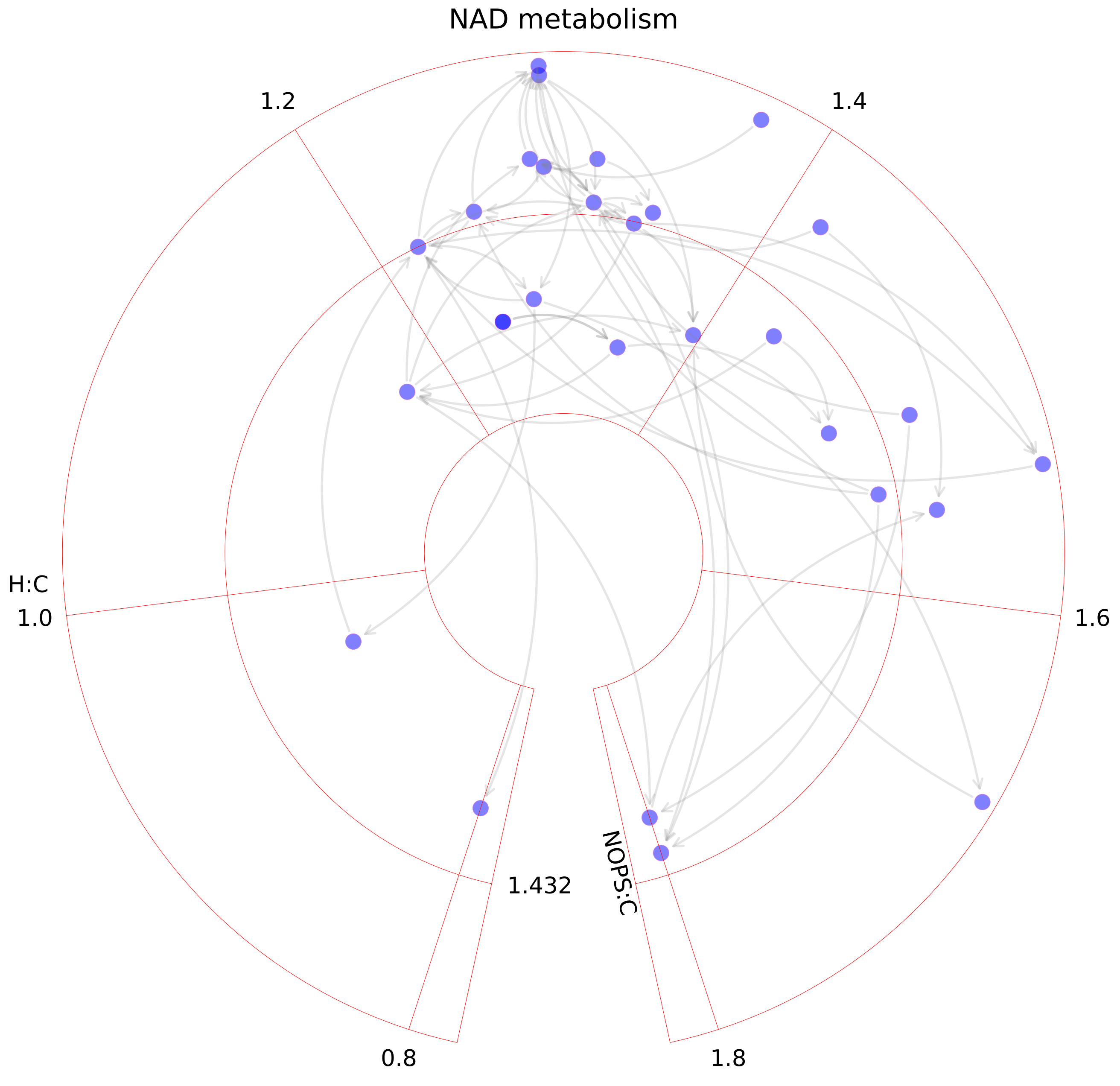

### Nucleotide interconversion.pdf

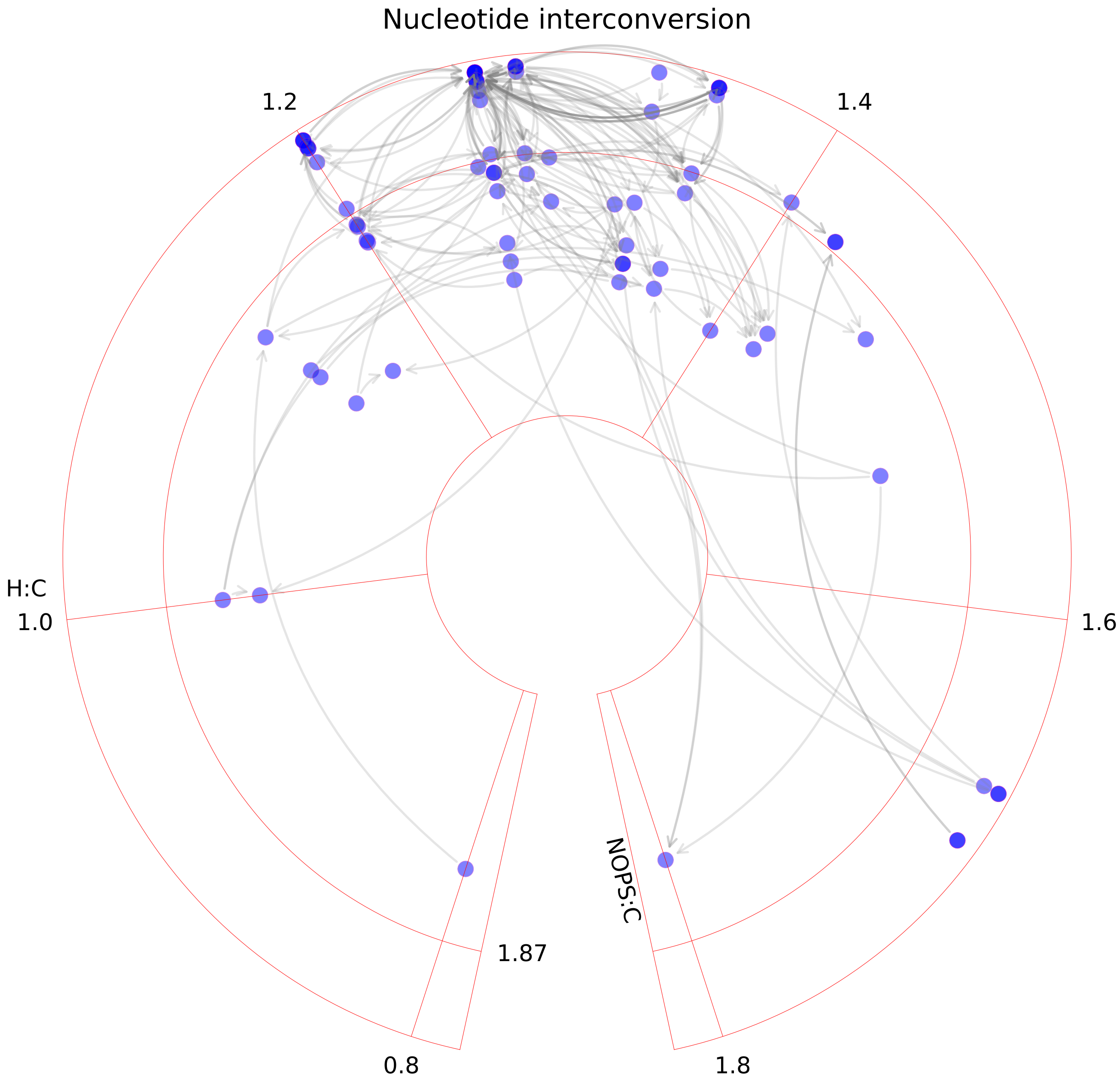

### Nucleotide metabolism.pdf

##### Nucleotide metabolism

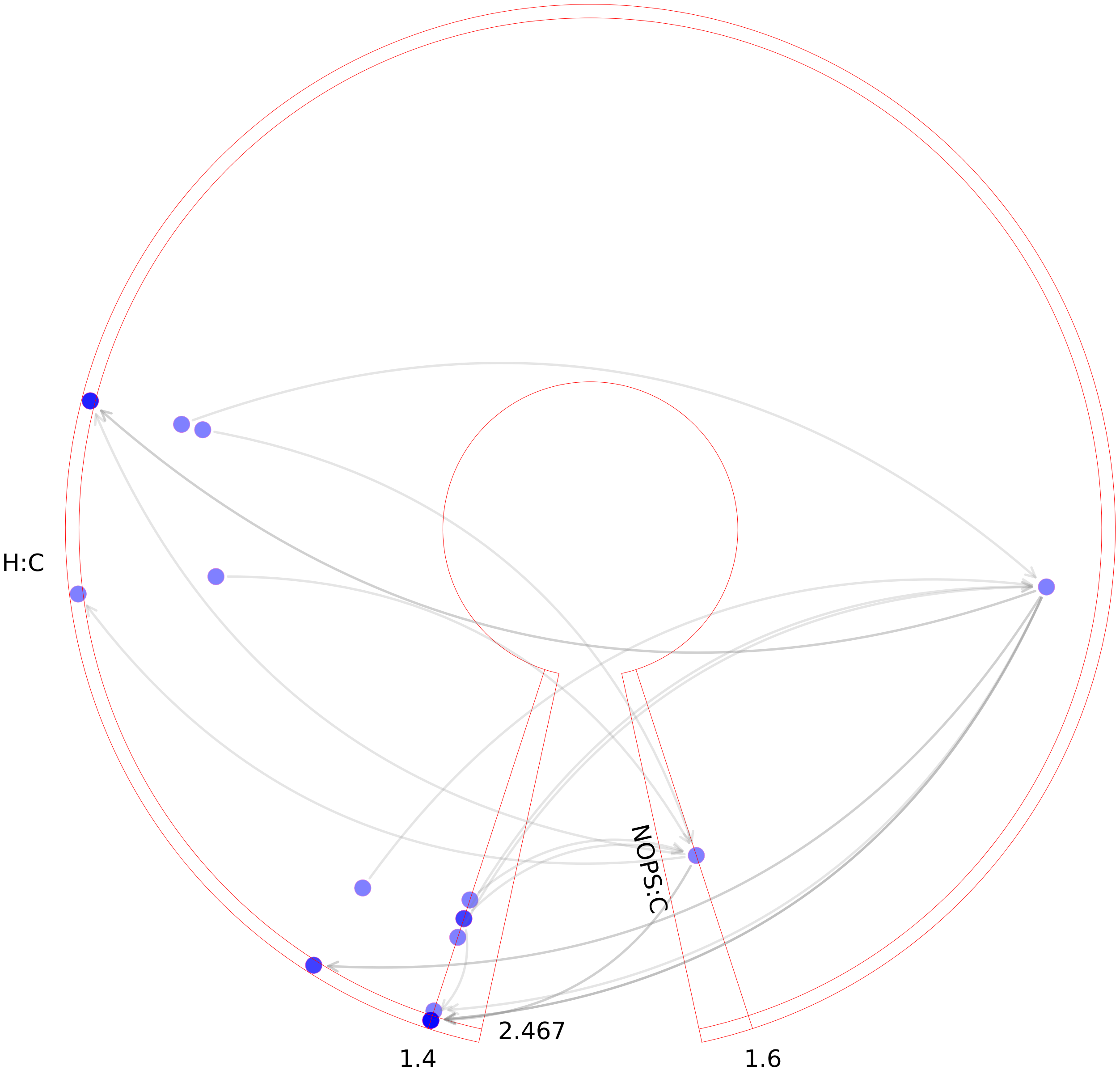

### Nucleotide metabolism.pdf

#### Nucleotide metabolism

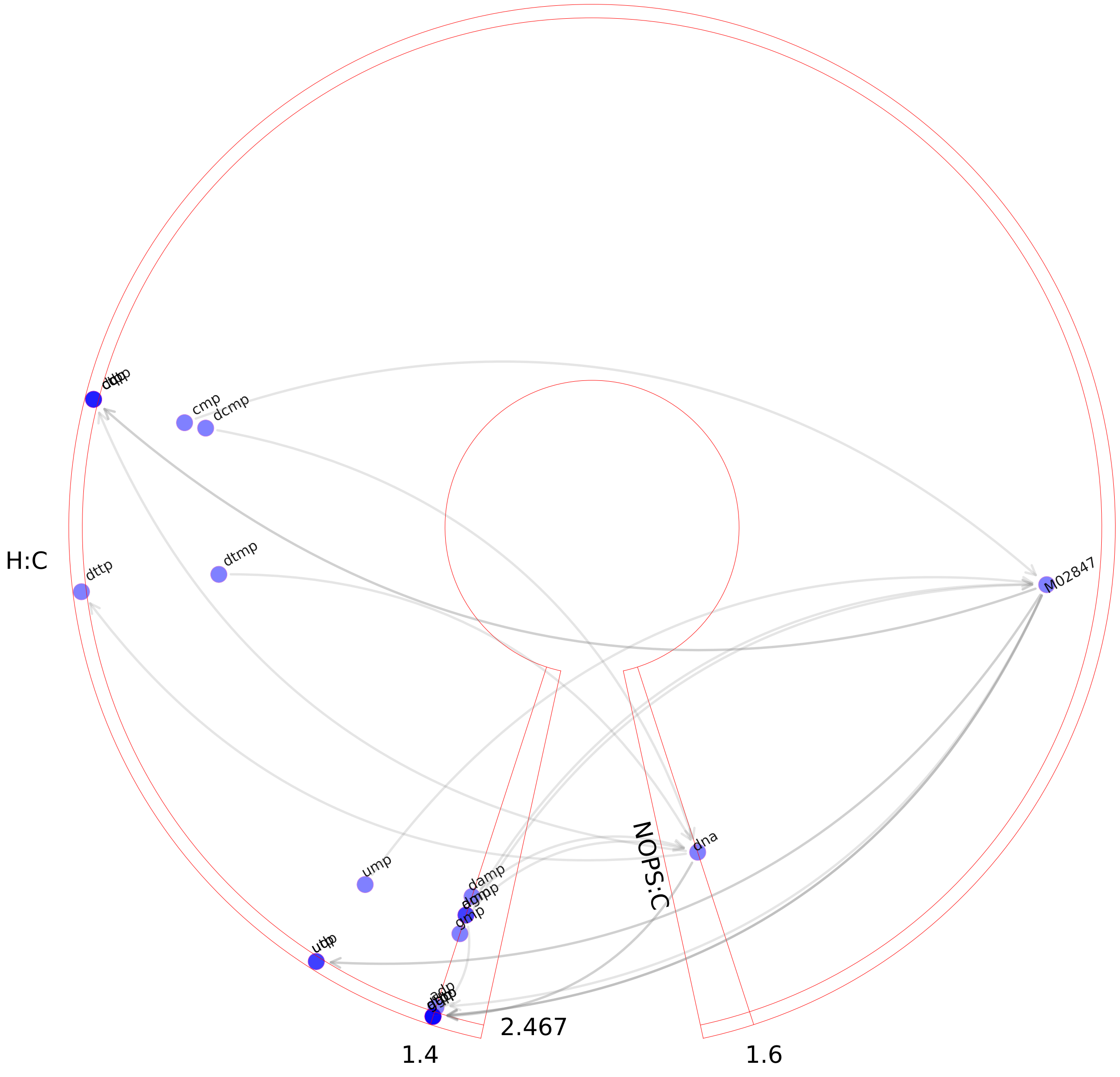

### Nucleotide salvage pathway.pdf

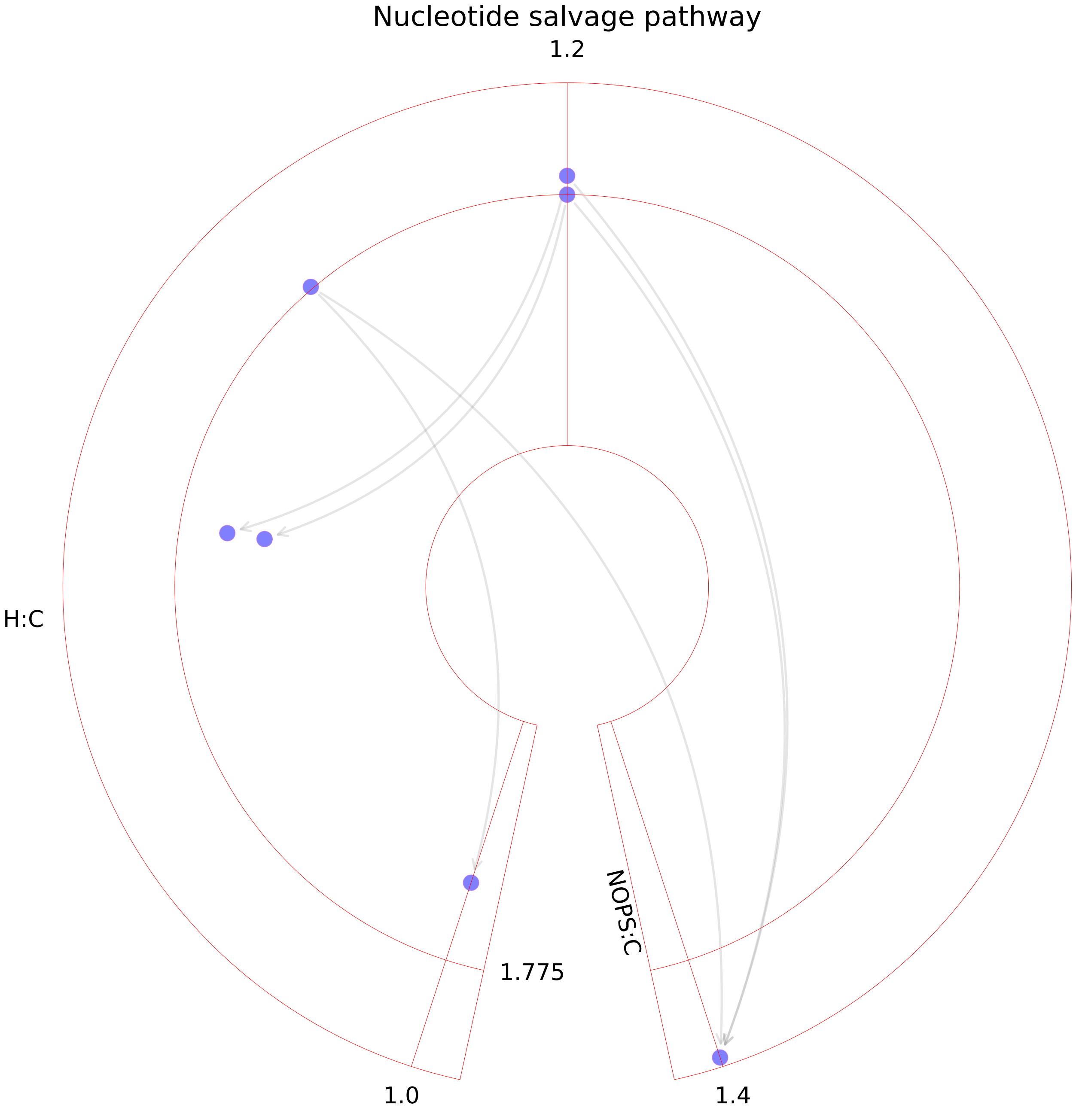

### Nucleotide sugar metabolism.pdf

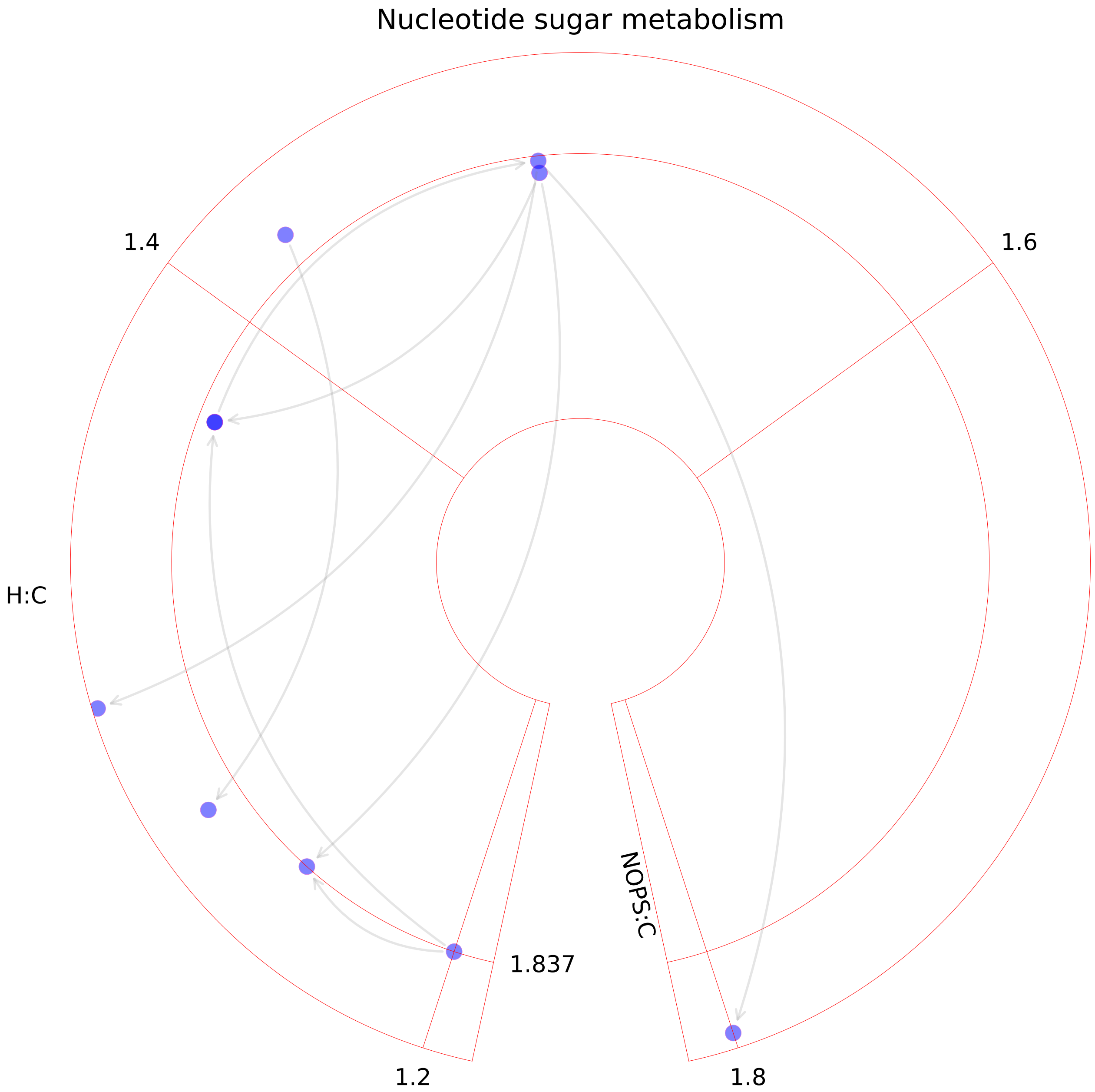

### O-glycan metabolism.pdf

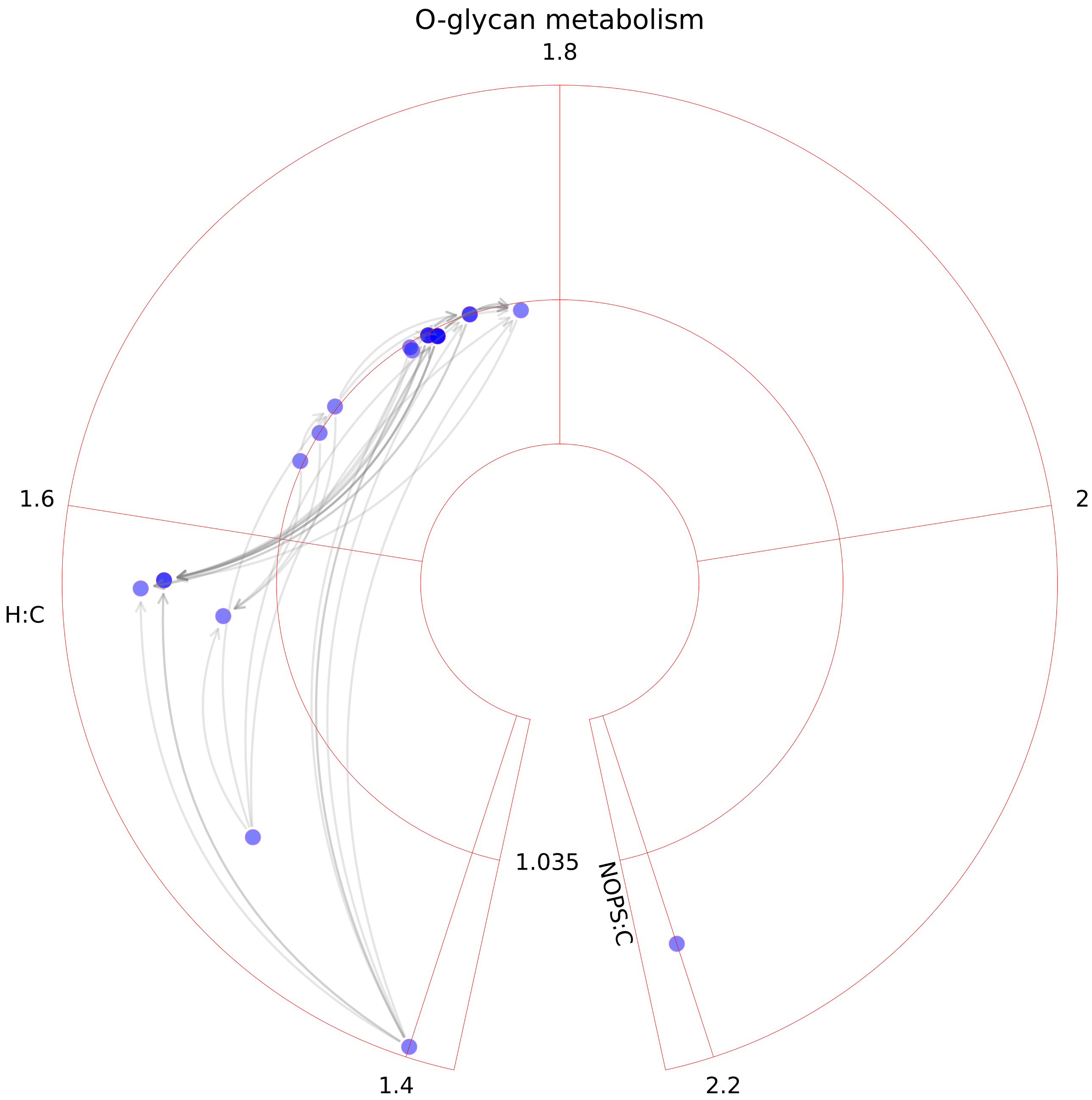

### Oxidative phosphorylation.pdf

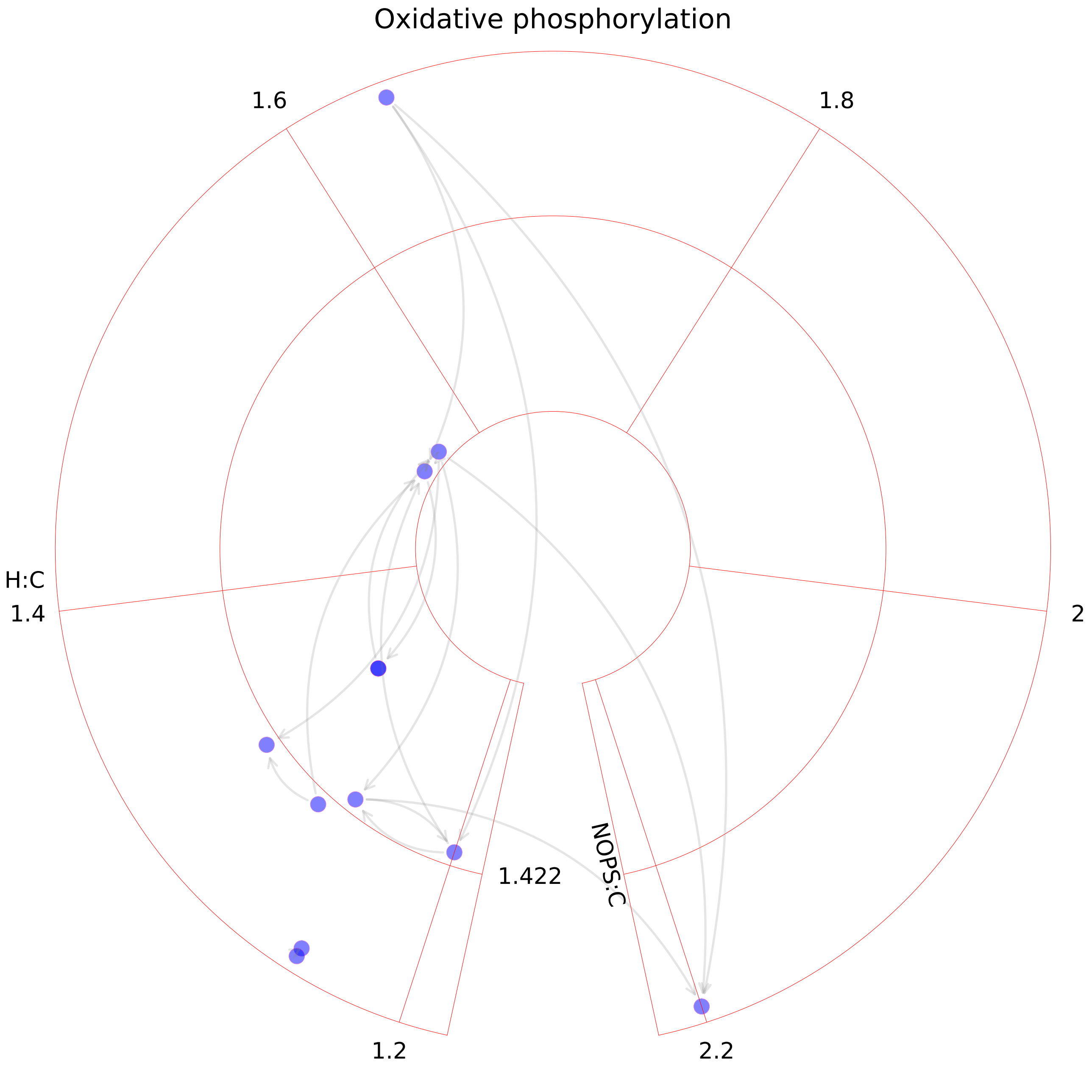

### Pentose phosphate pathway.pdf

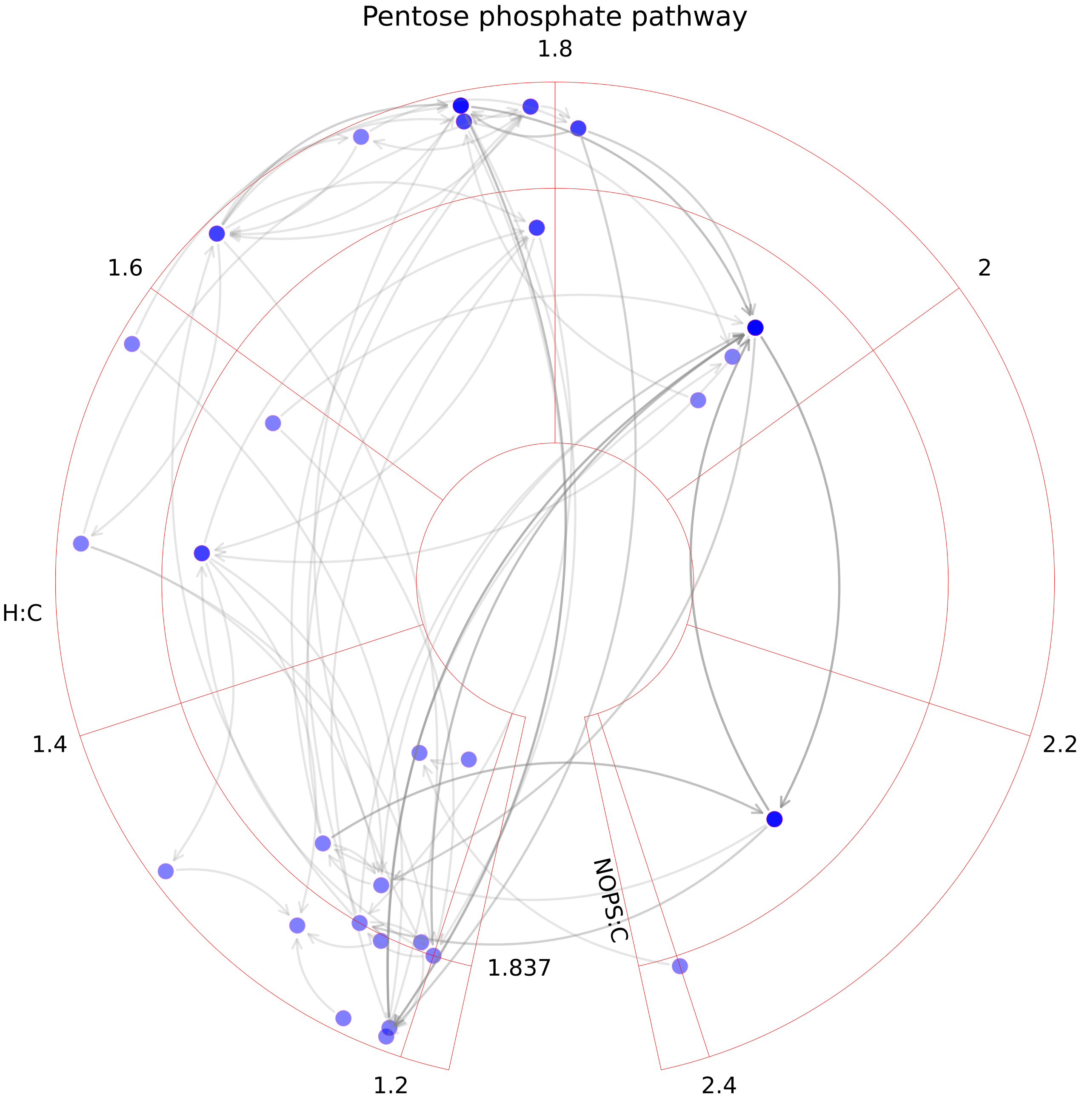

### Peptide metabolism.pdf

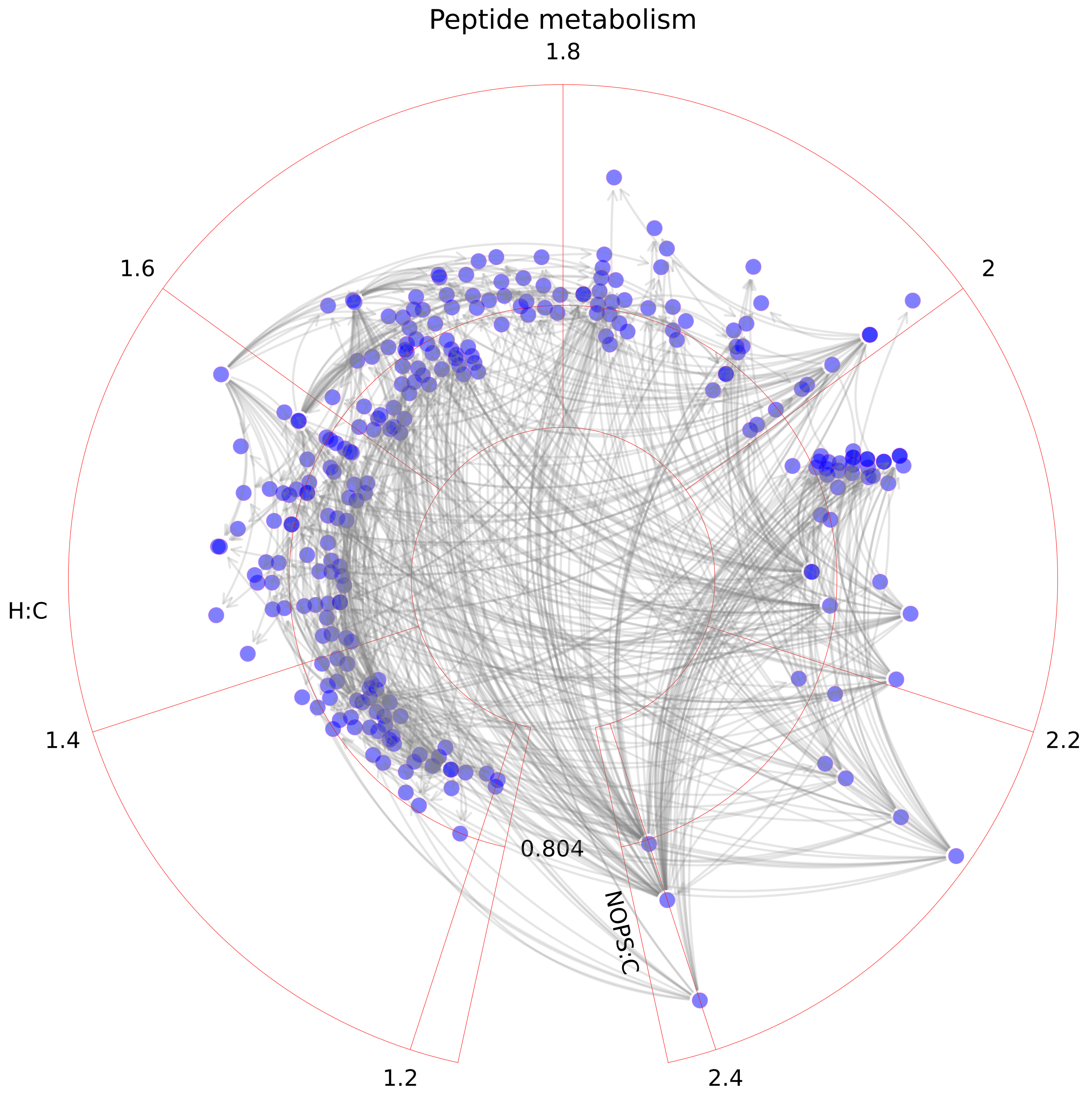

### Phenylalanine metabolism.pdf

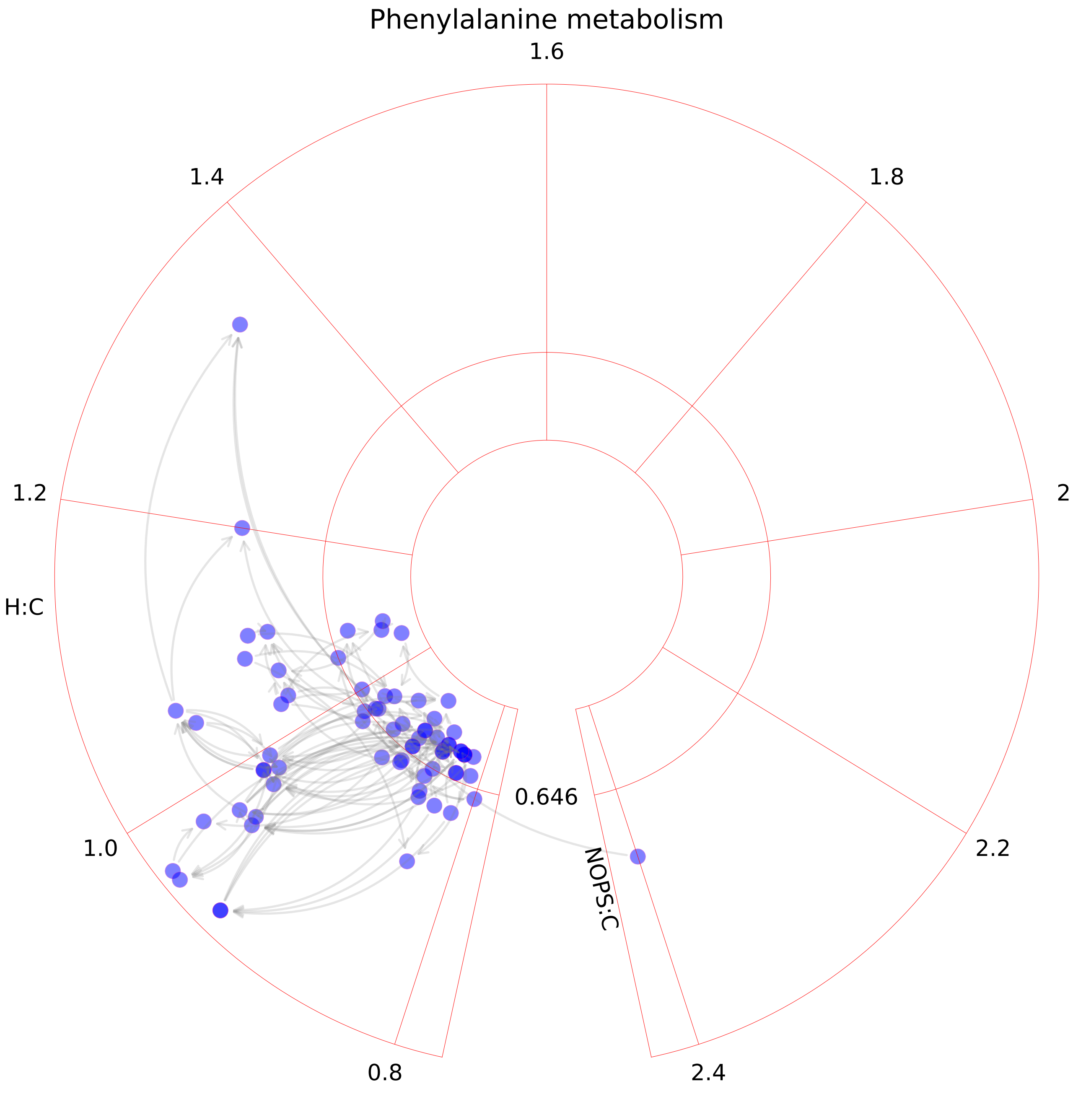

### Phosphatidylinositol phosphate metabolism.pdf

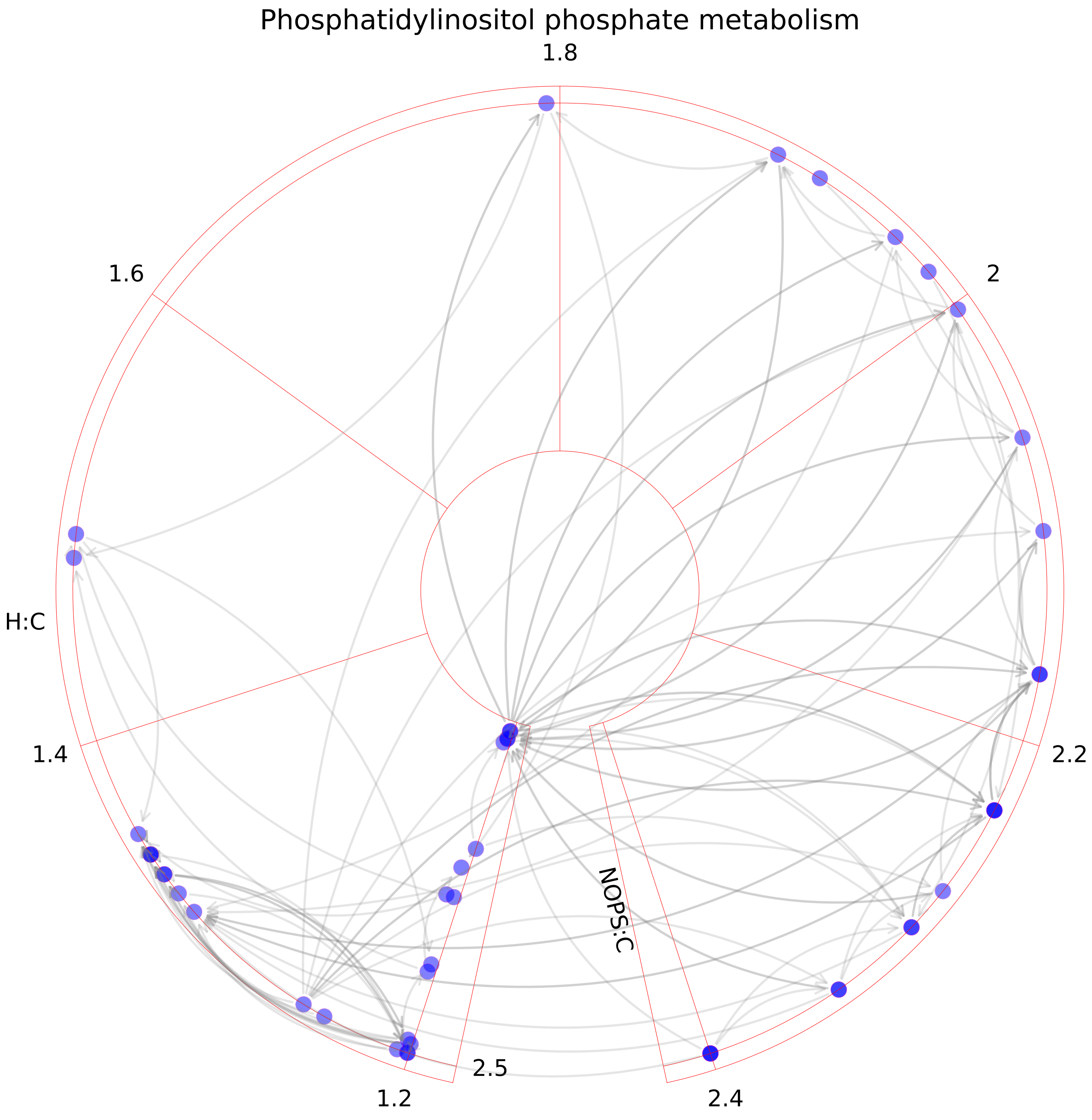

### Propanoate metabolism.pdf

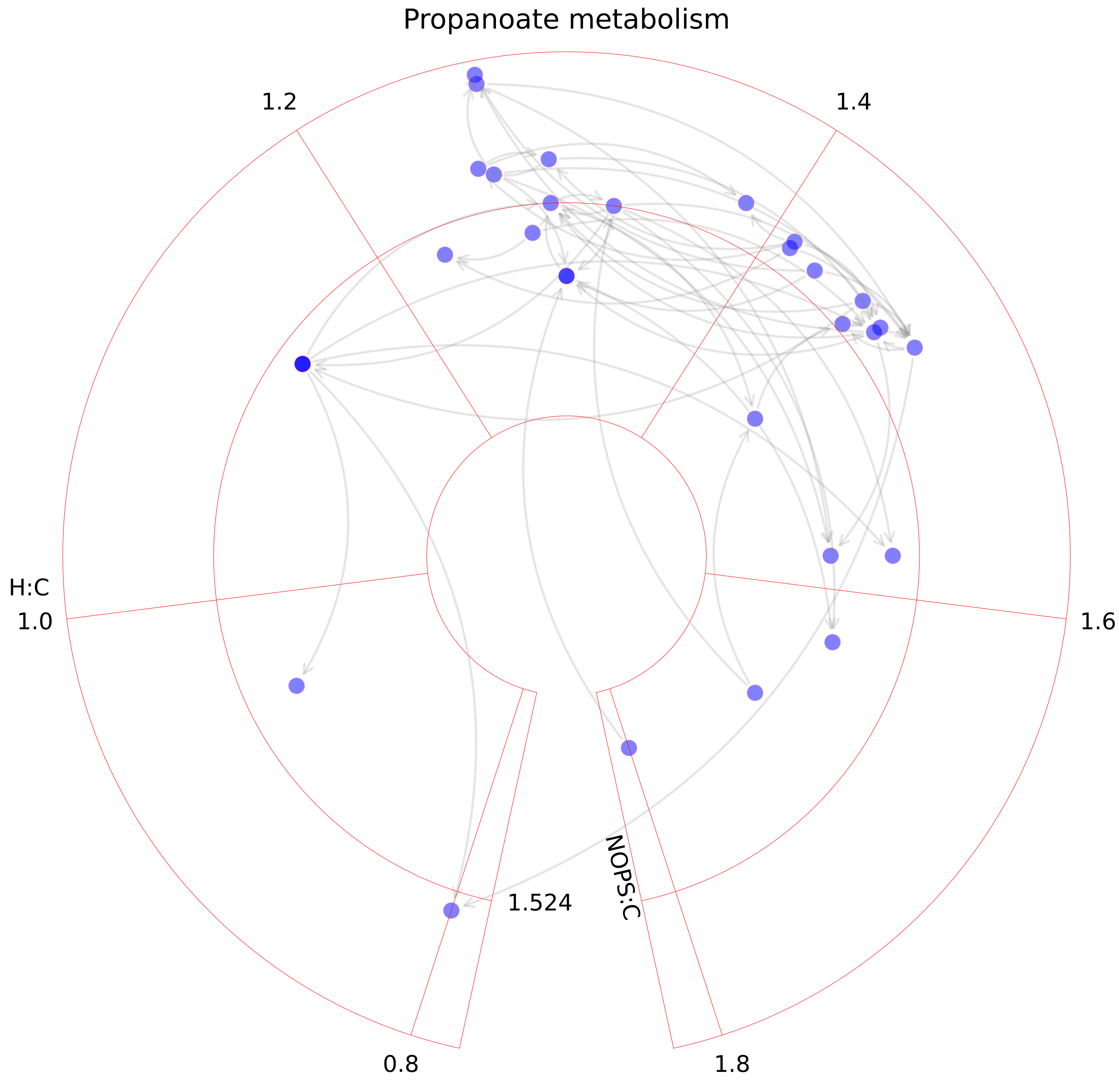

### Purine catabolism.pdf

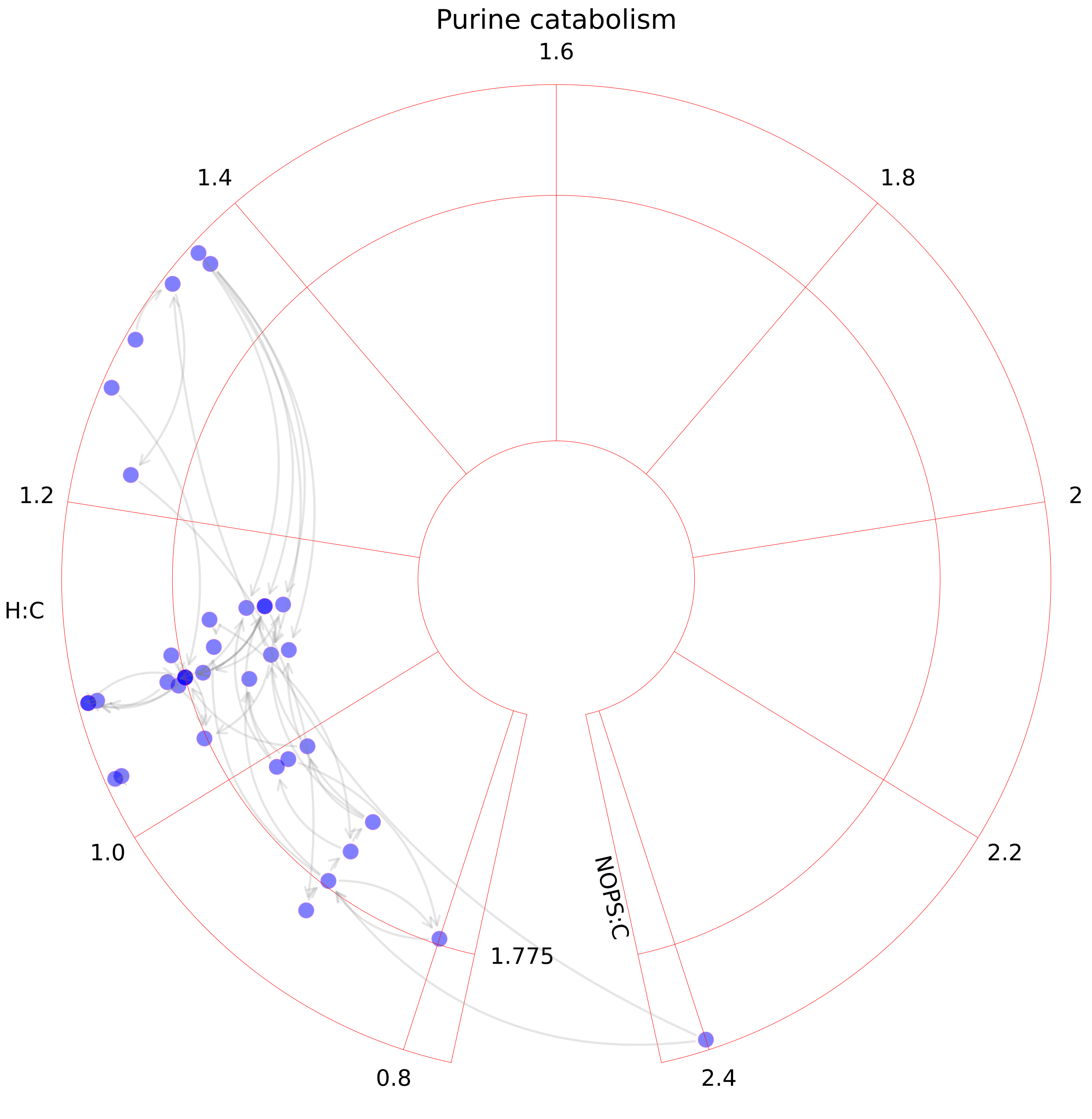

### Purine synthesis.pdf

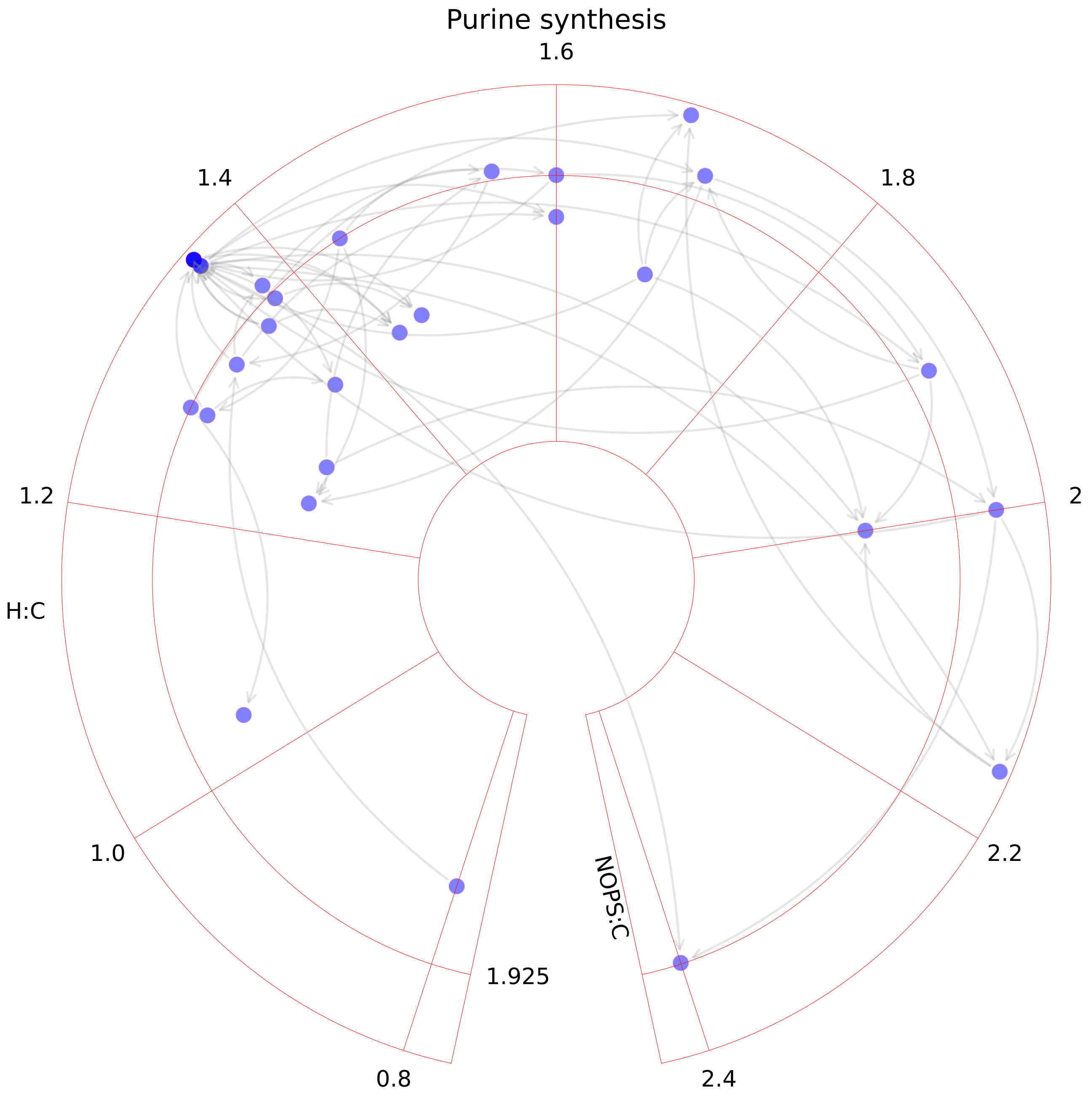

### Pyrimidine catabolism.pdf

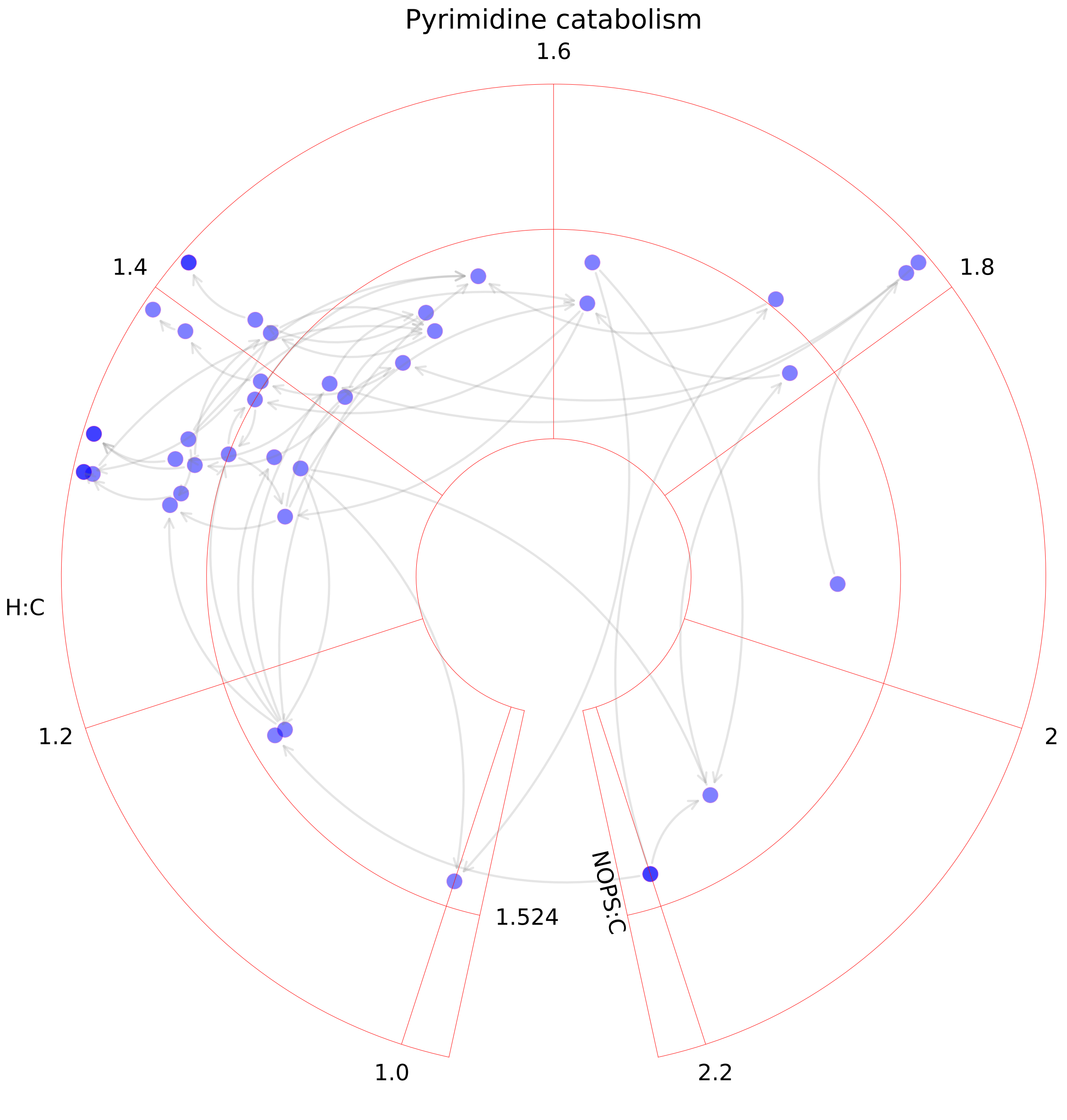

### Pyrimidine synthesis.pdf

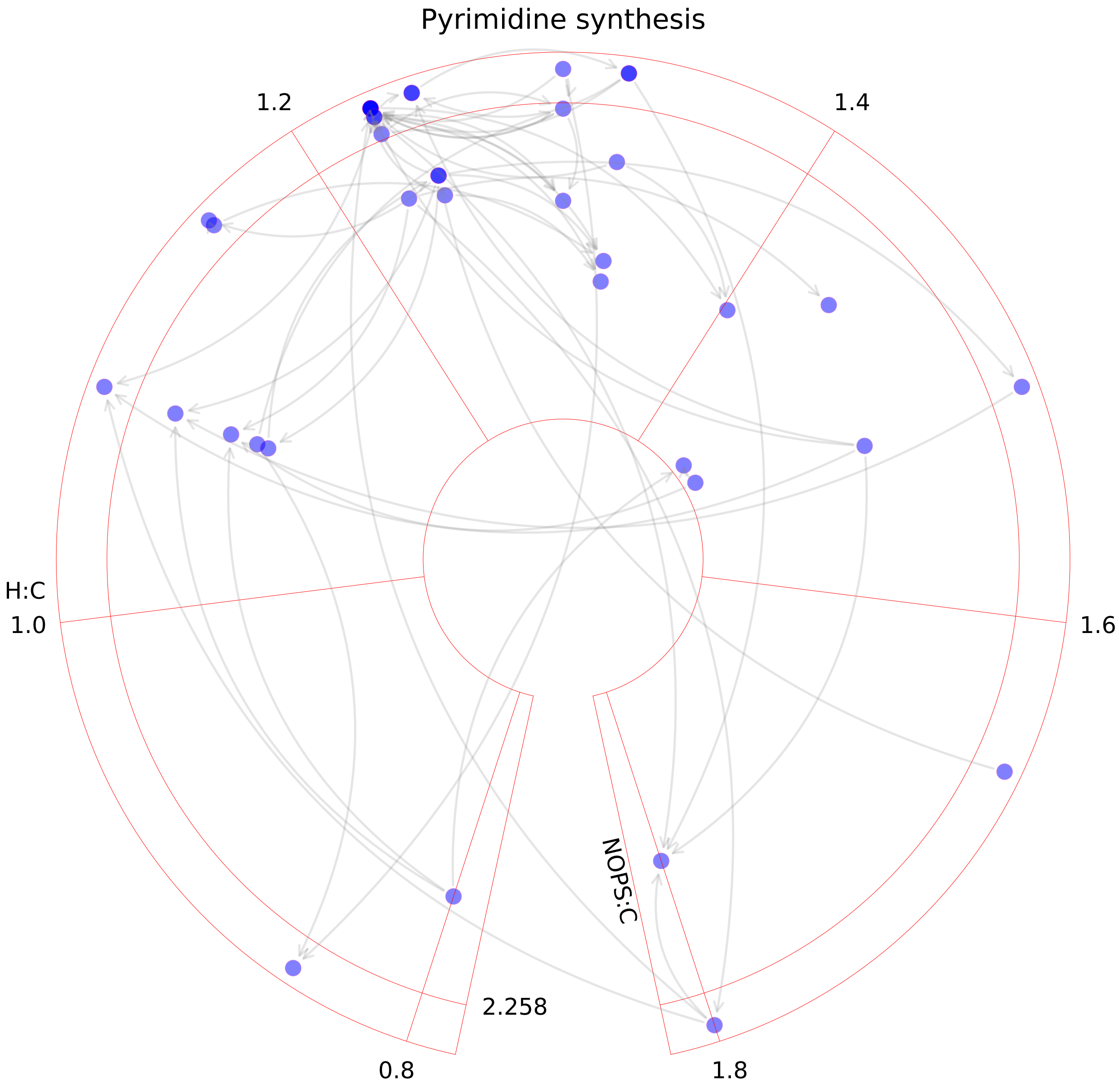

### Pyruvate metabolism.pdf

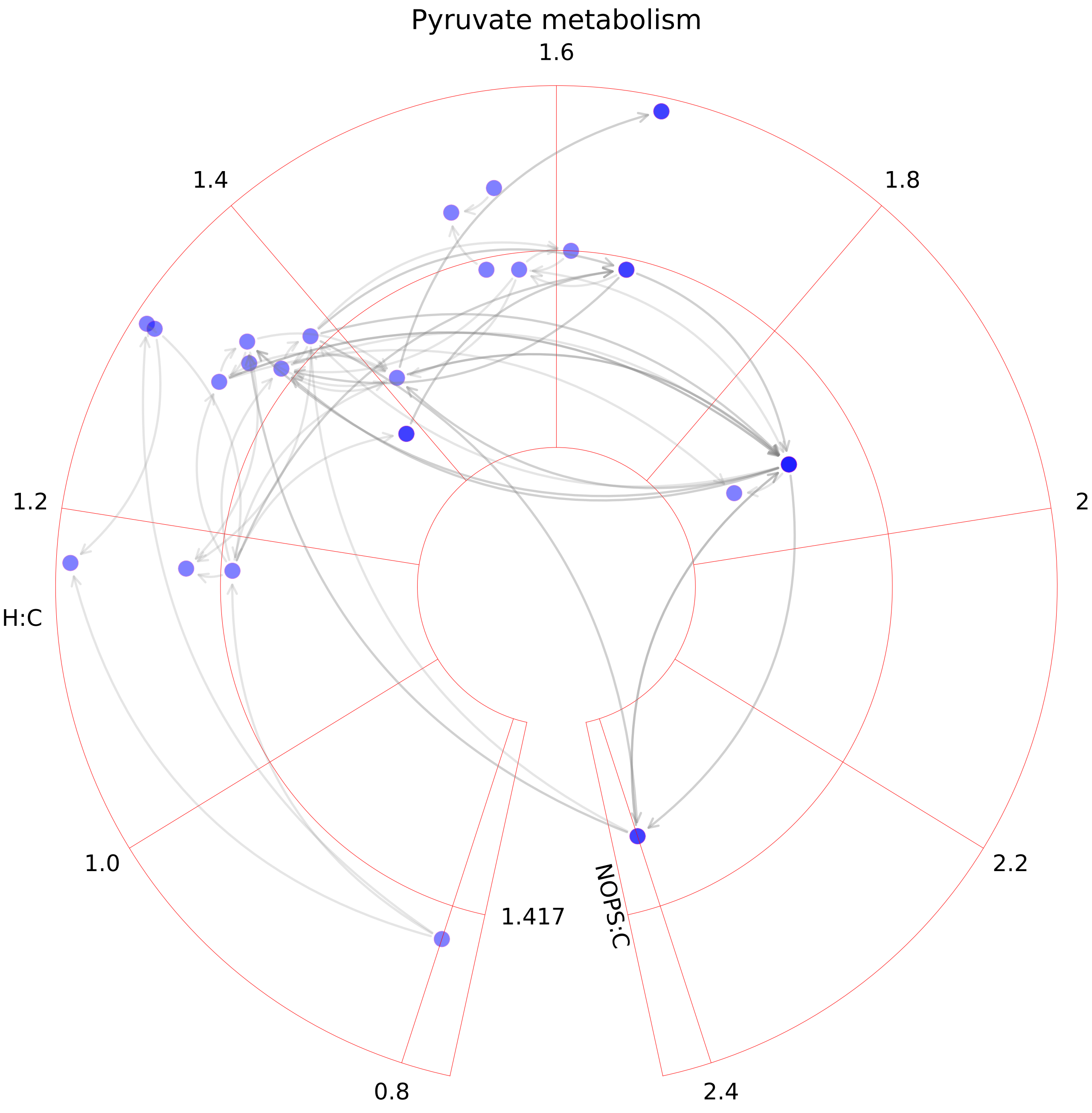

### R group synthesis.pdf

##### R group synthesis

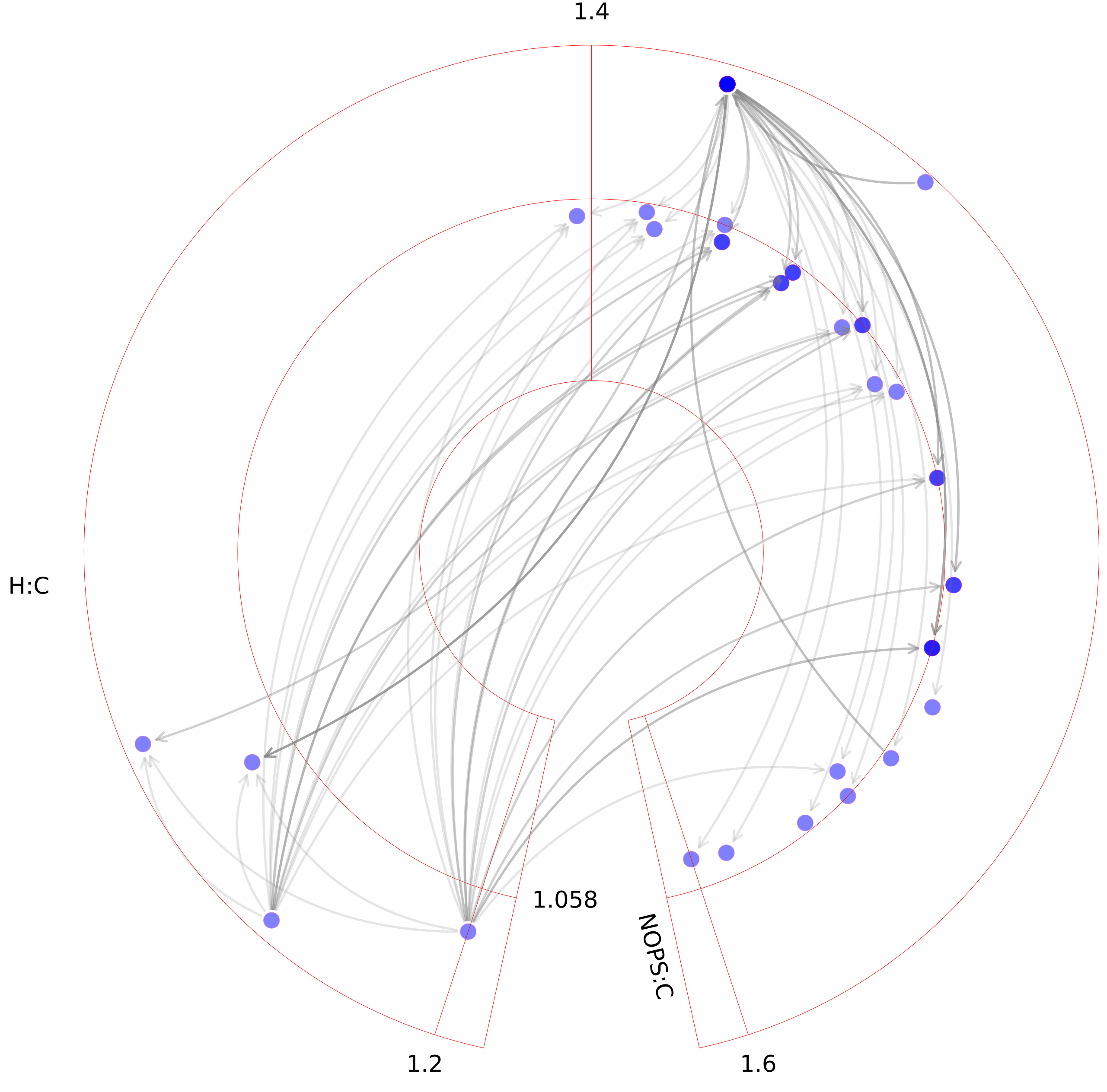

### ROS detoxification.pdf

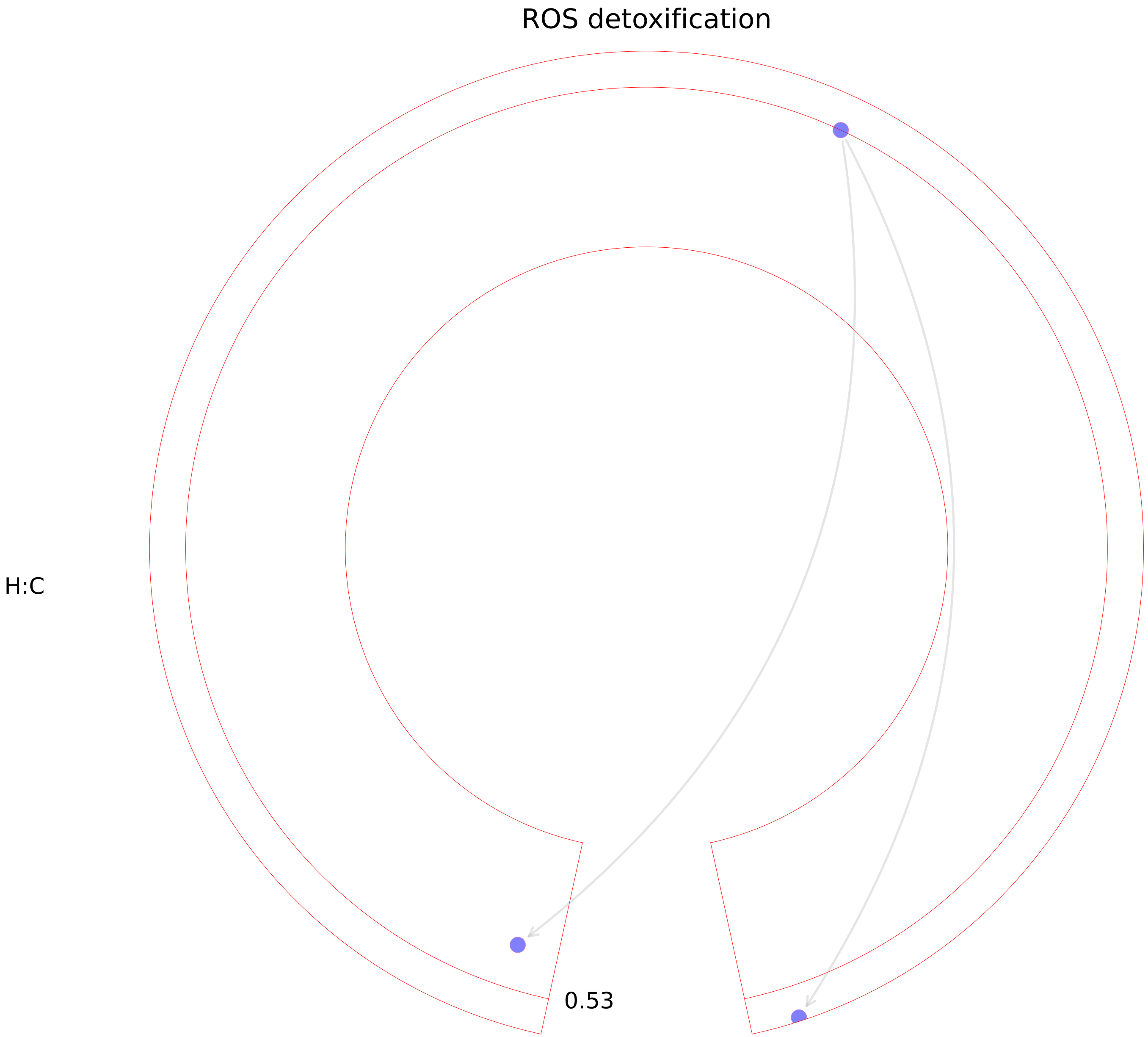

### Selenoamino acid metabolism.pdf

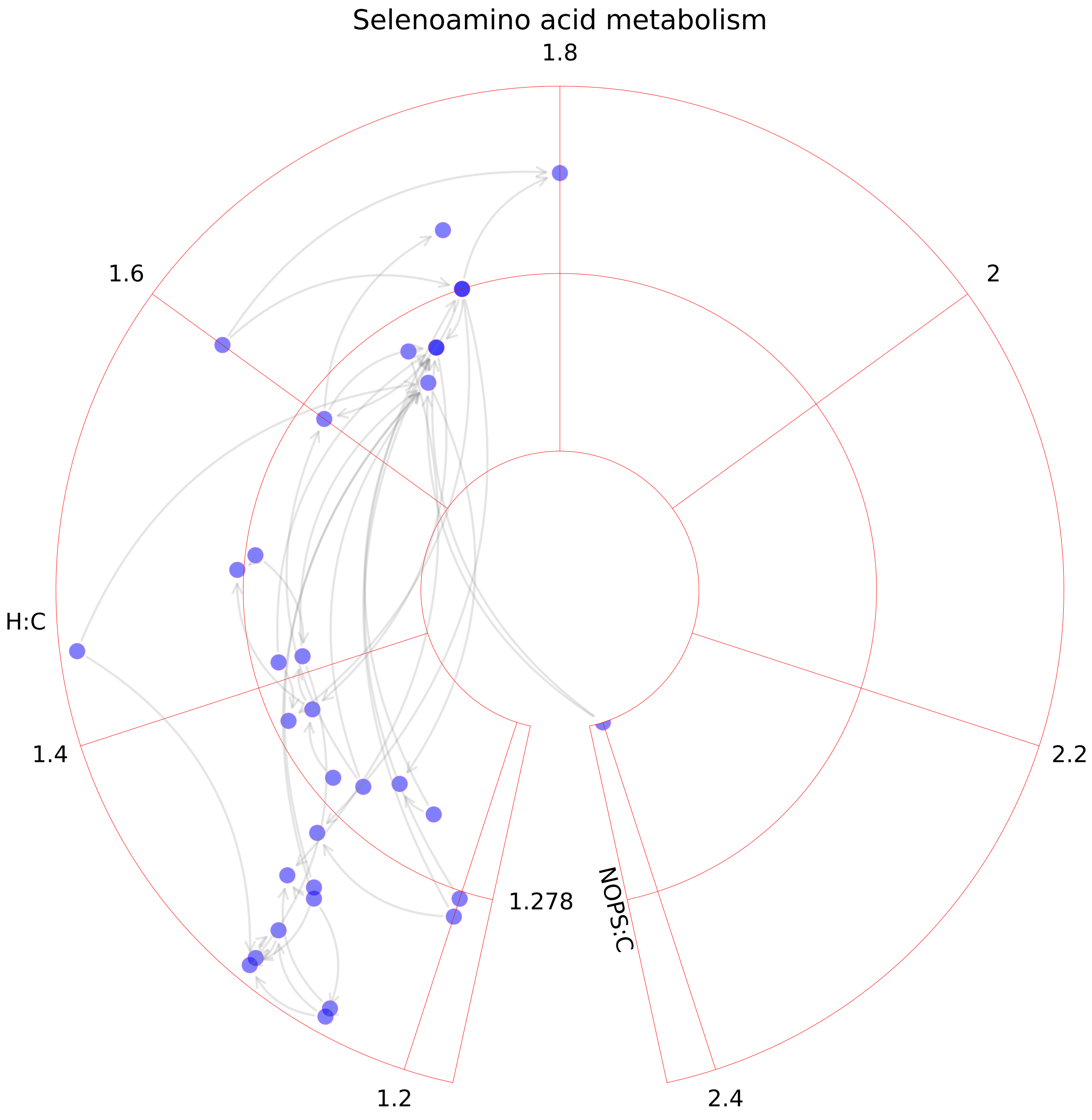

### Sphingolipid metabolism.pdf

##### Sphingolipid metabolism

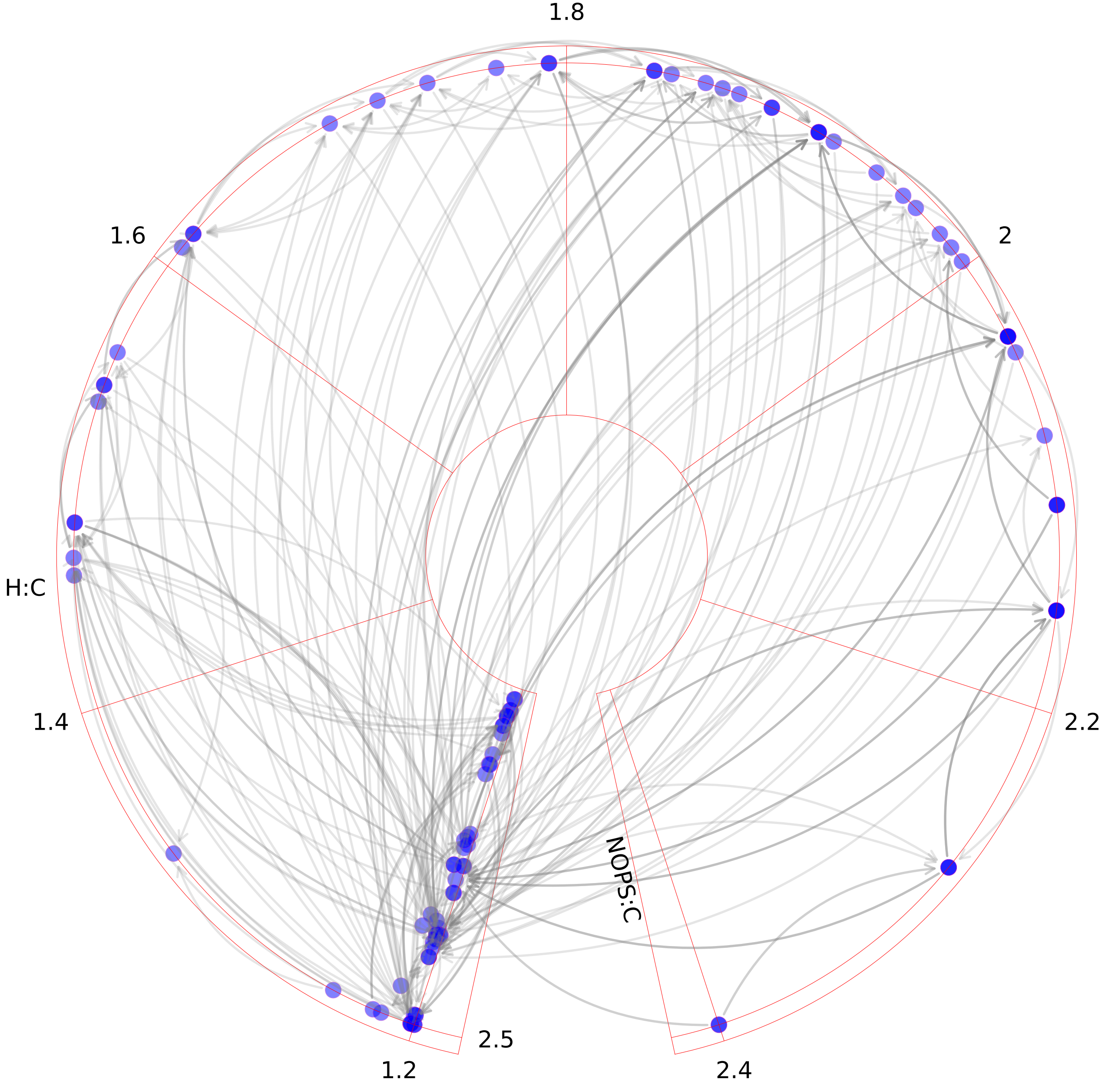

### Squalene and cholesterol synthesis.pdf

Squalene and cholesterol synthesis

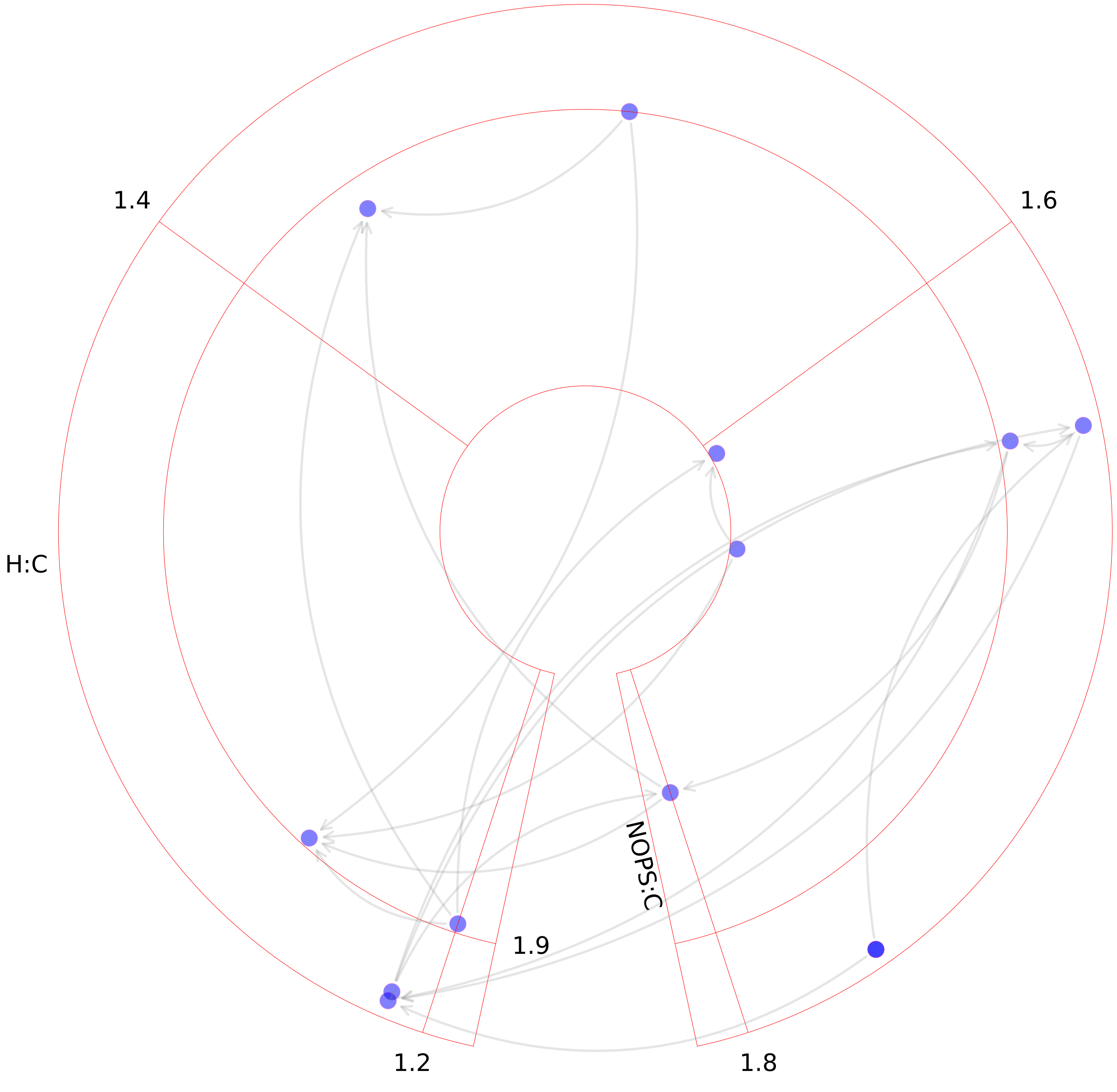

### Starch and sucrose metabolism.pdf

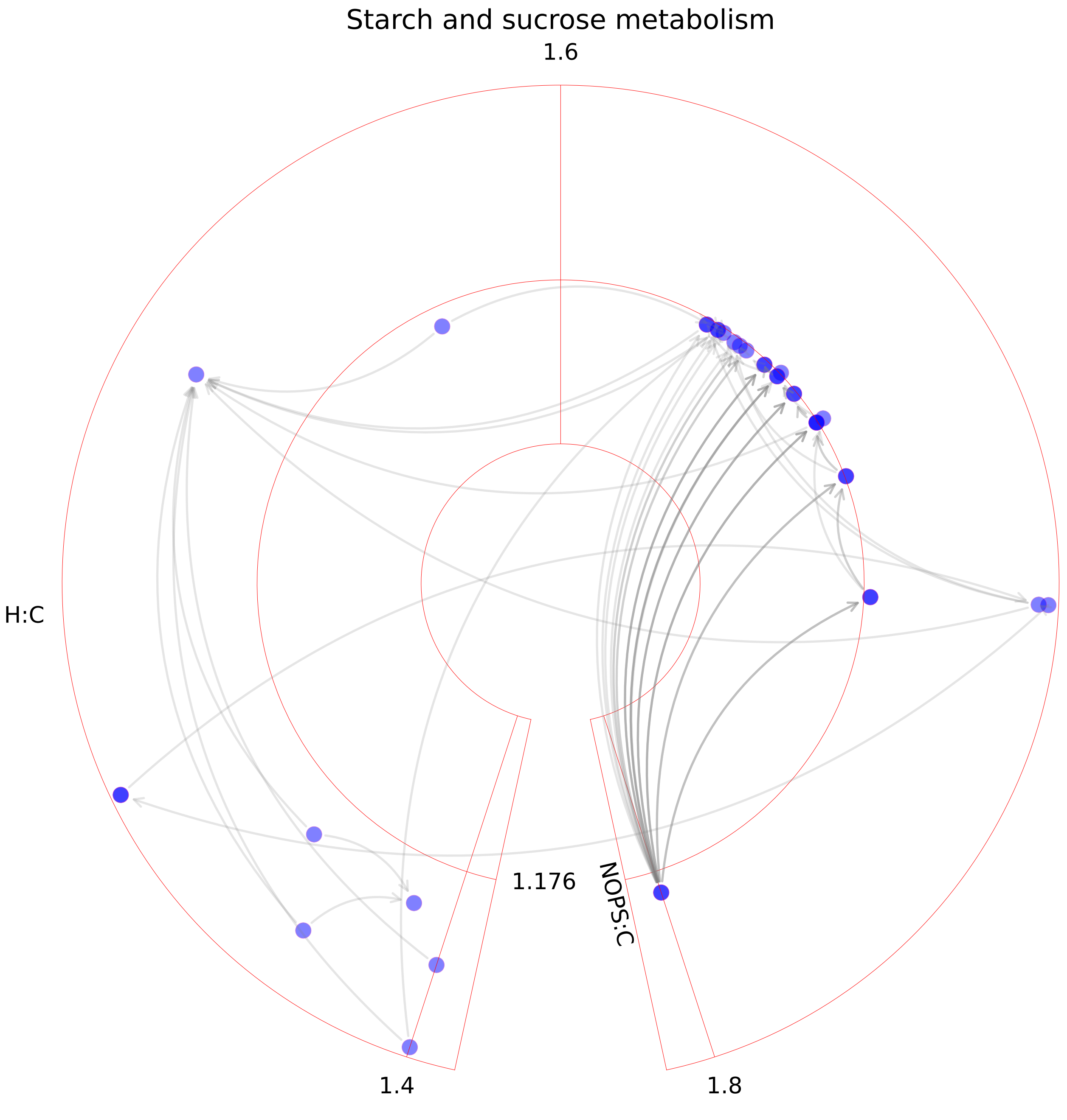

### Steroid metabolism.pdf

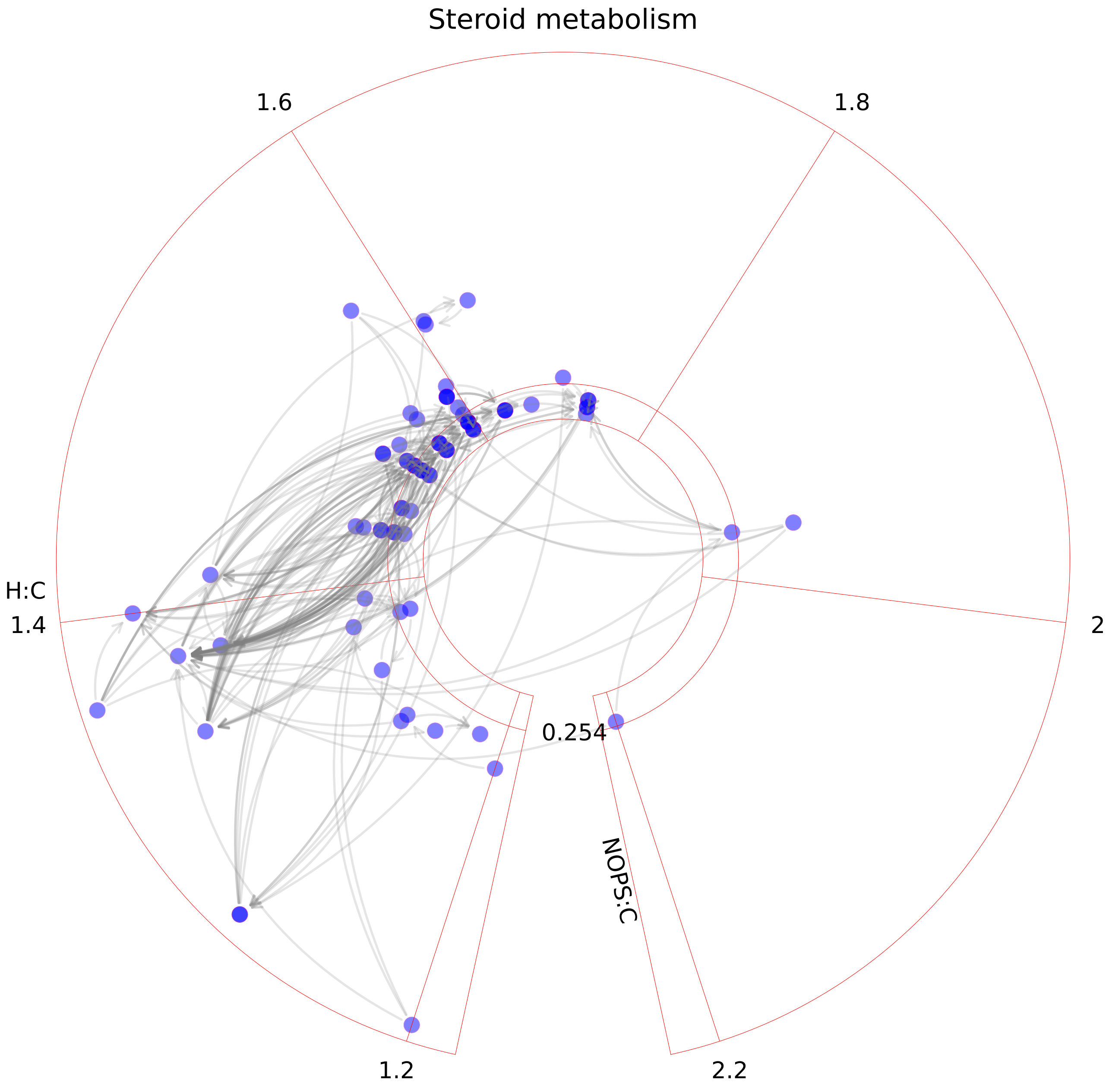

### Stilbene, coumarine and lignin synthesis.pdf

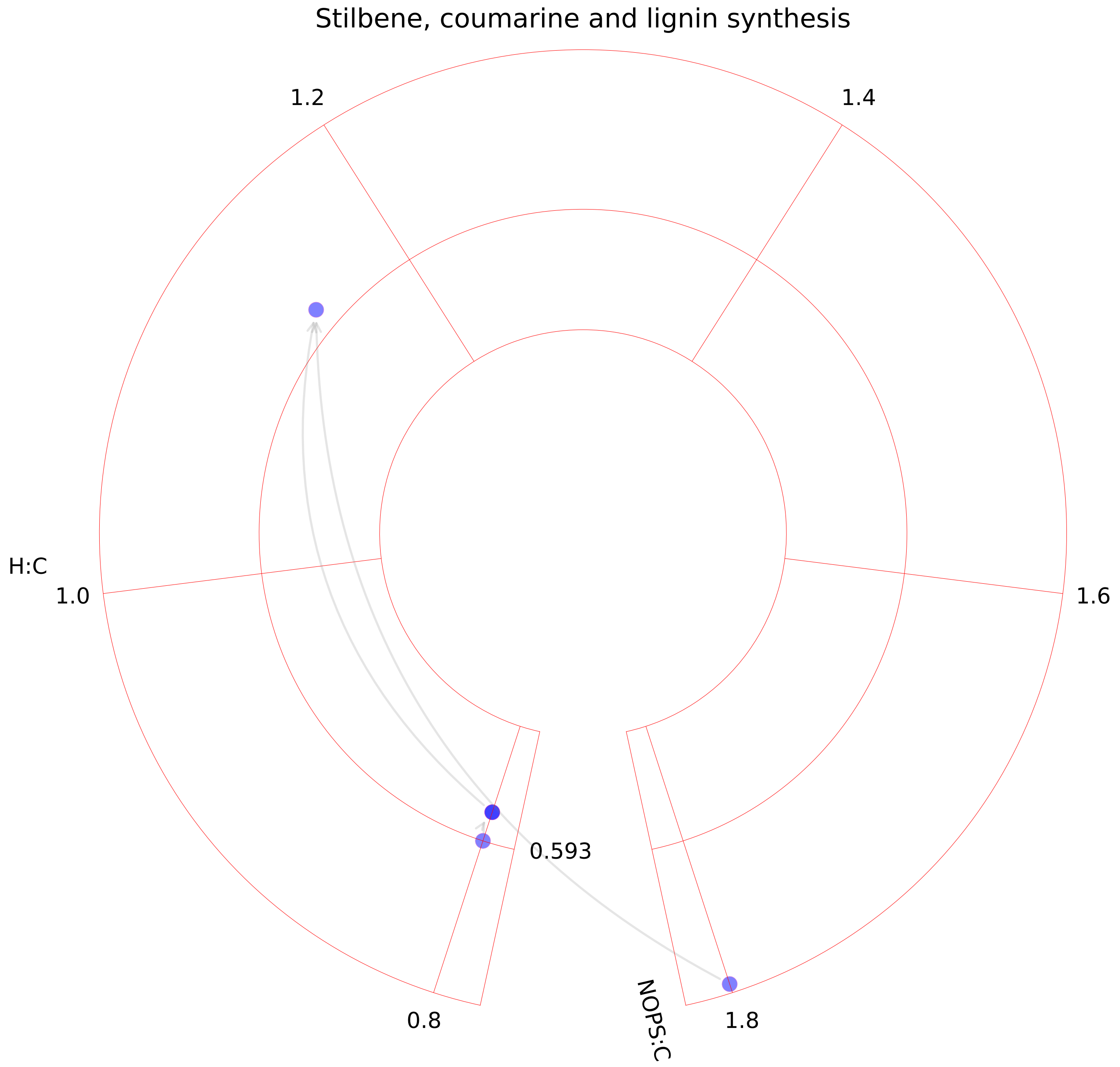

### Taurine and hypotaurine metabolism.pdf

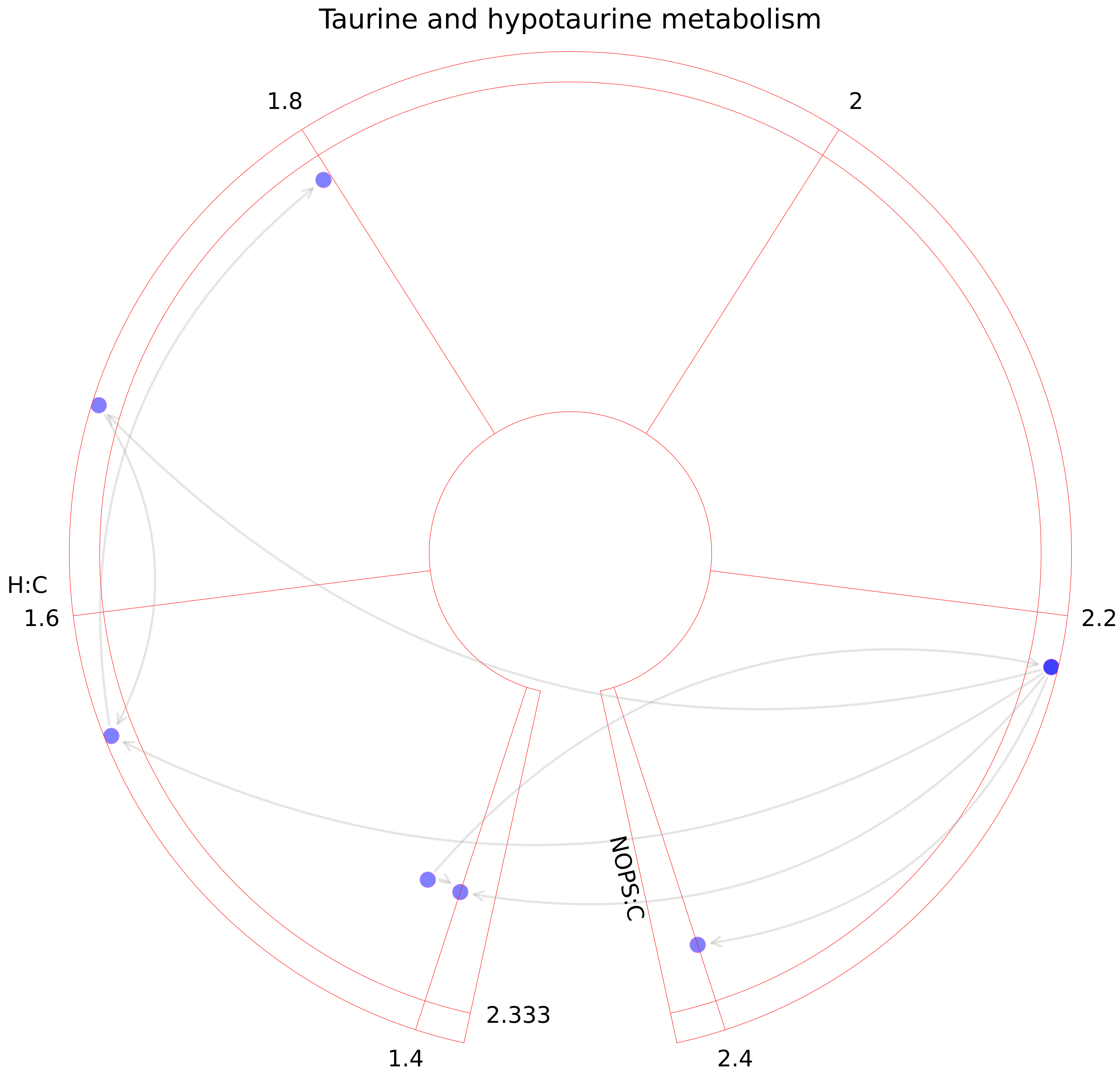

### Tetrahydrobiopterin metabolism.pdf

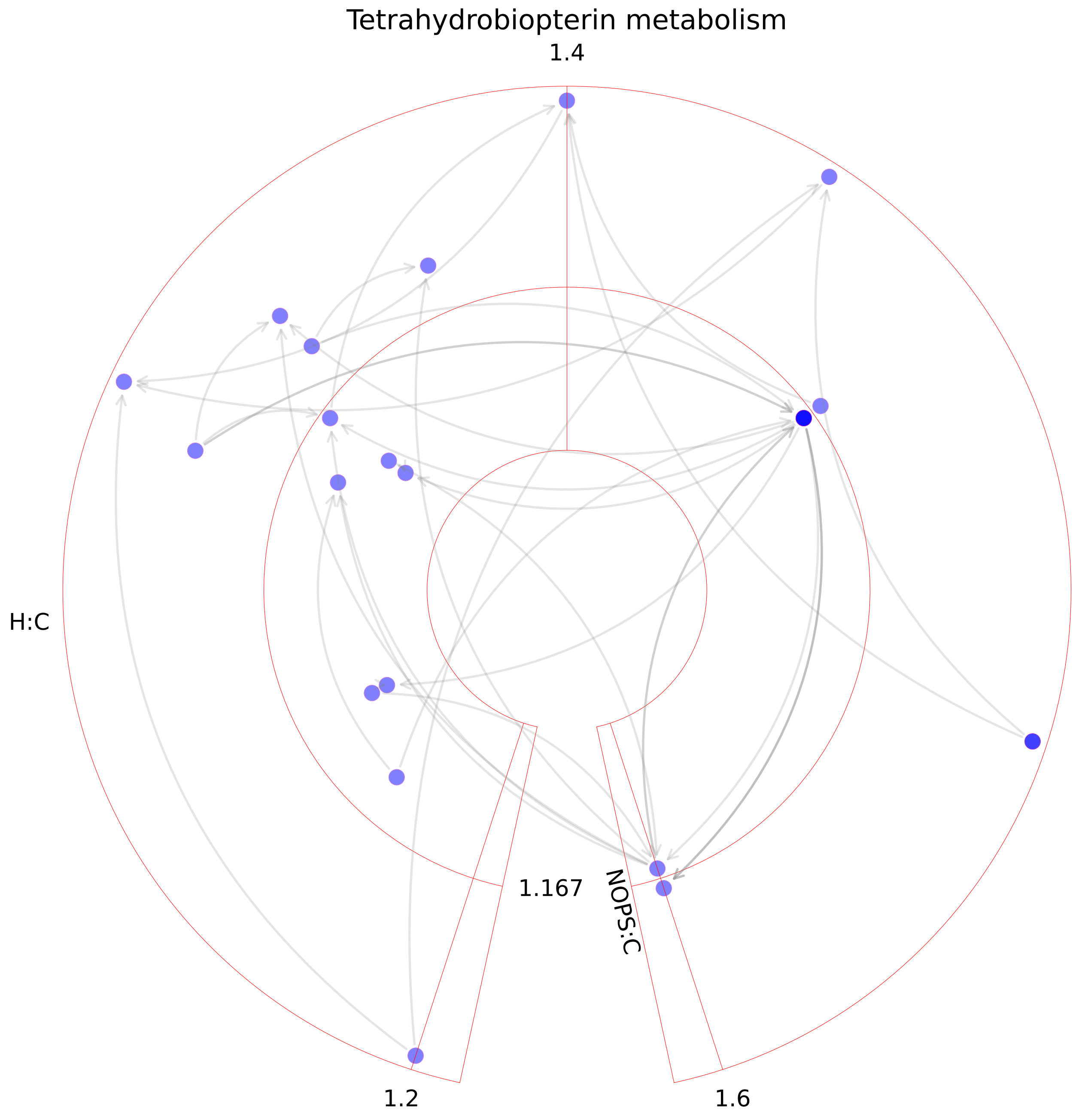

### Thiamine metabolism.pdf

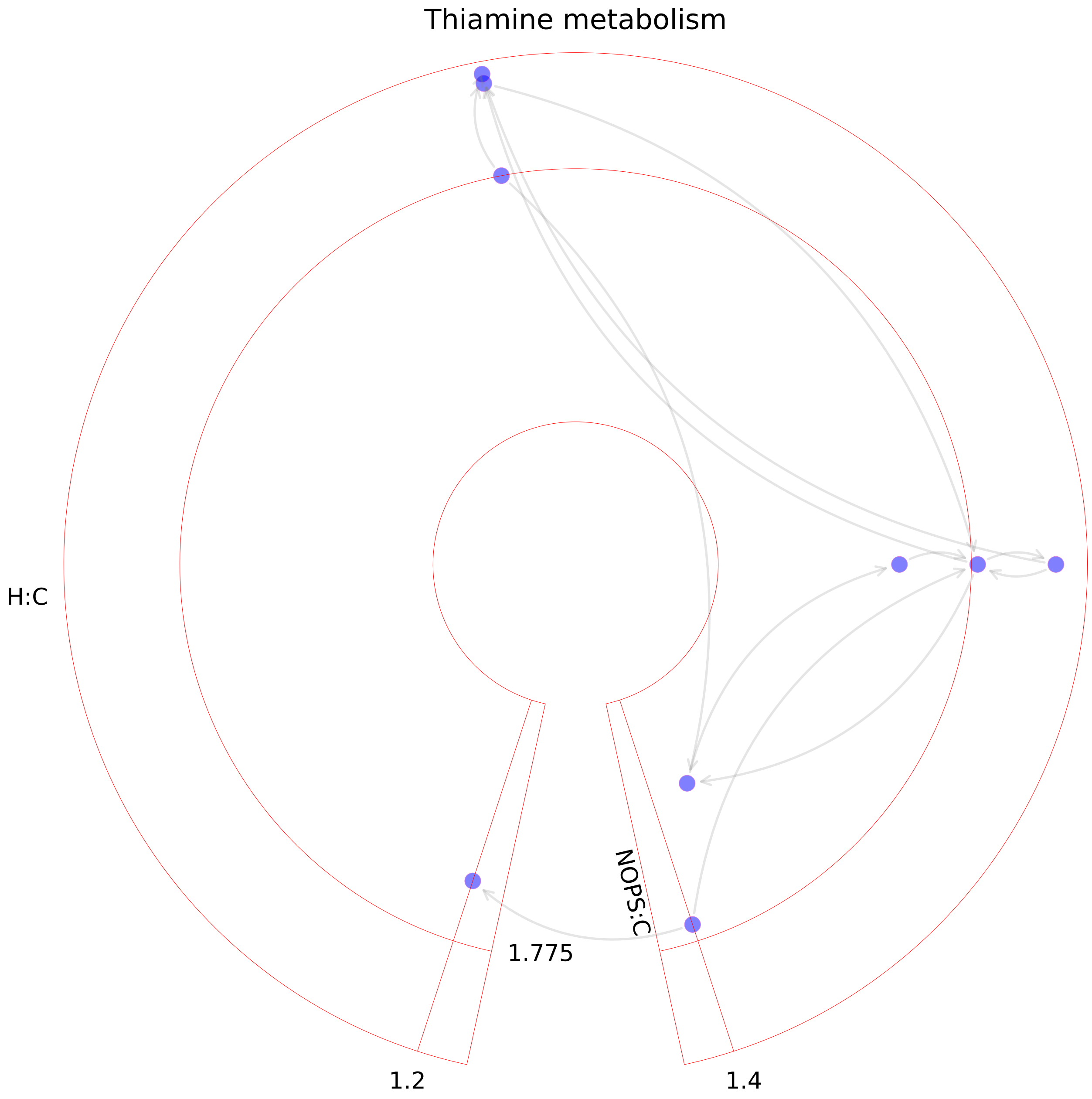

### Triacylglycerol synthesis.pdf

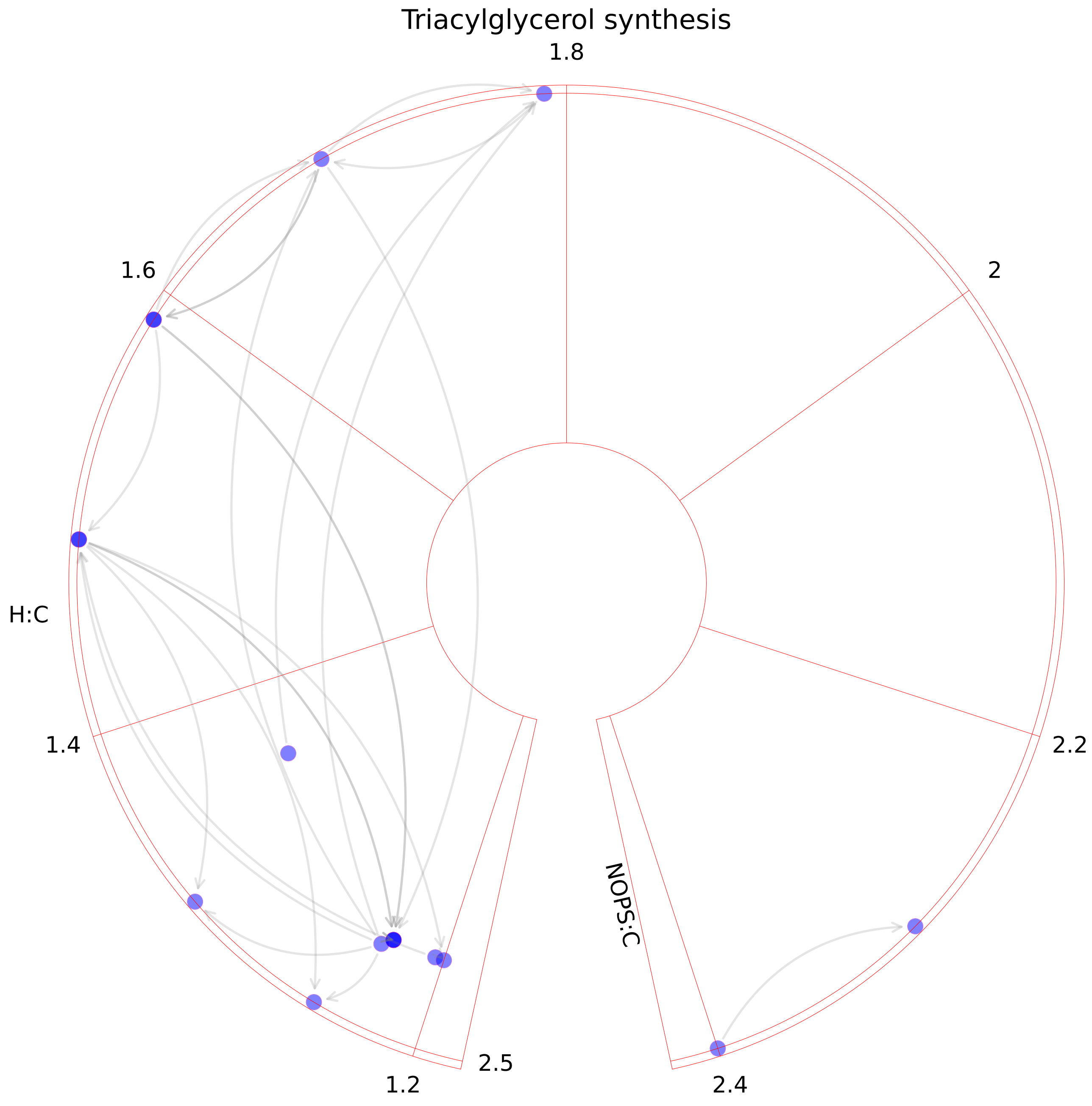

### Triglycerides formation.pdf

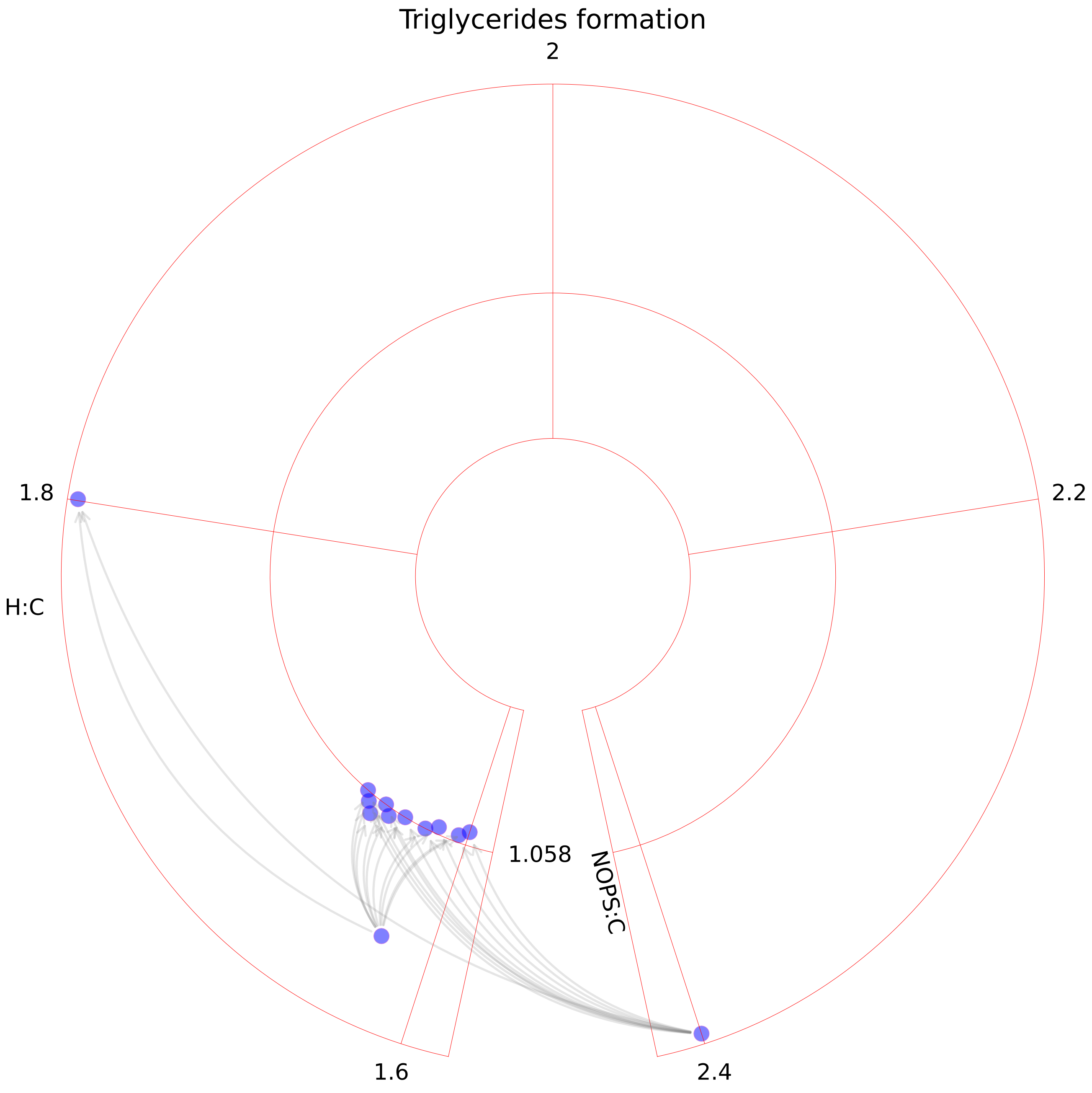

### Tryptophan metabolism.pdf

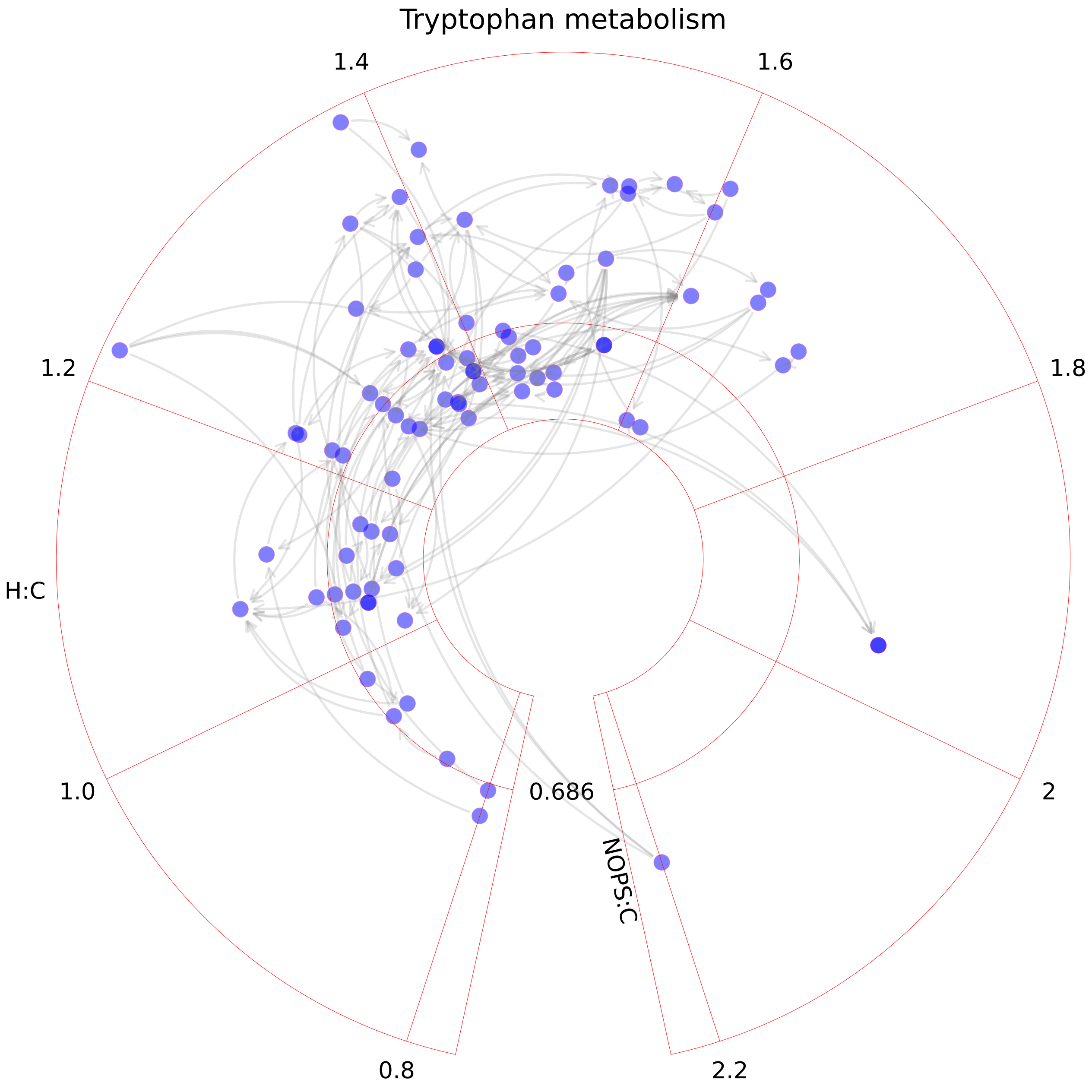

### Tyrosine metabolism.pdf

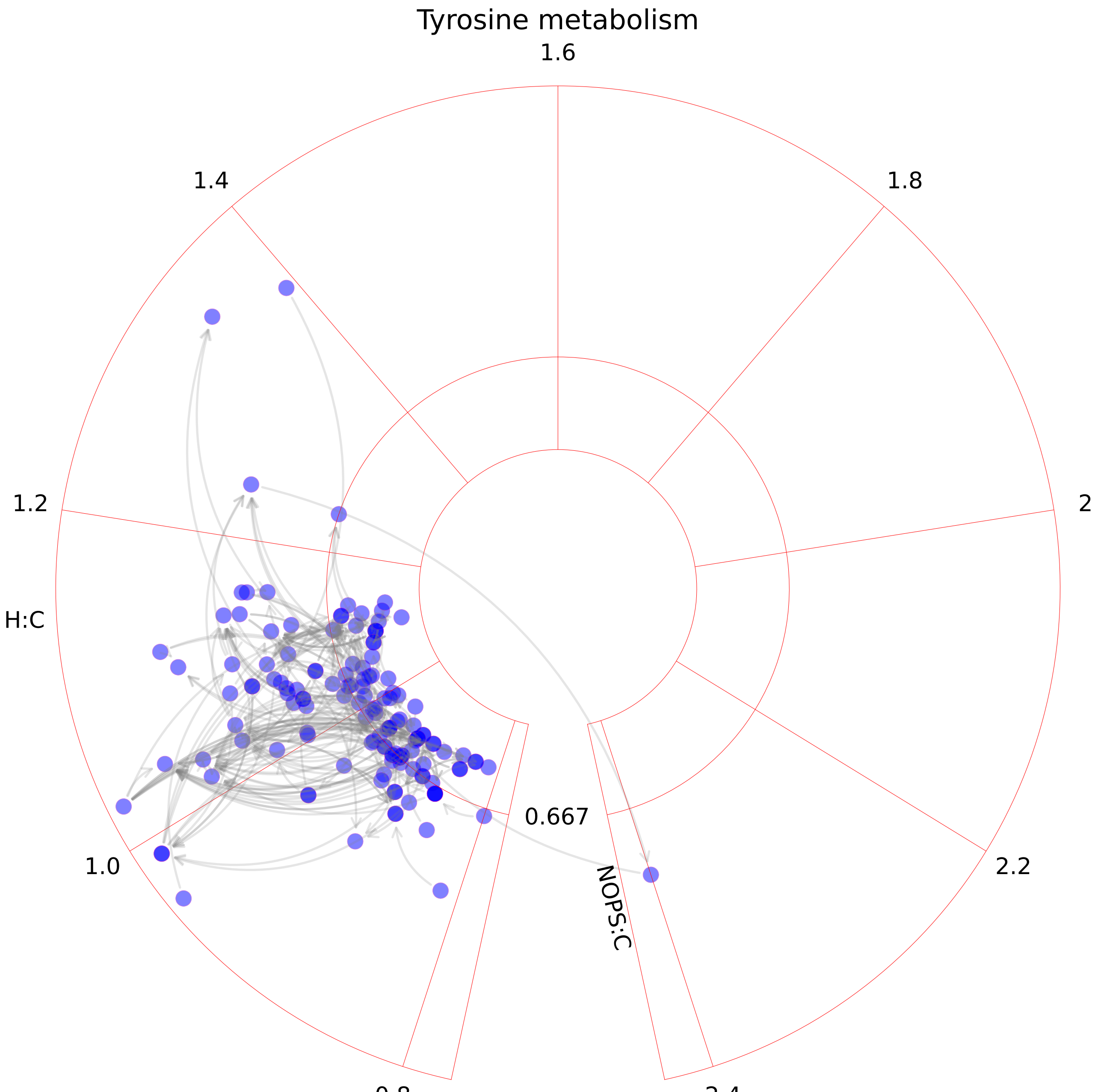

### Ubiquinone synthesis.pdf

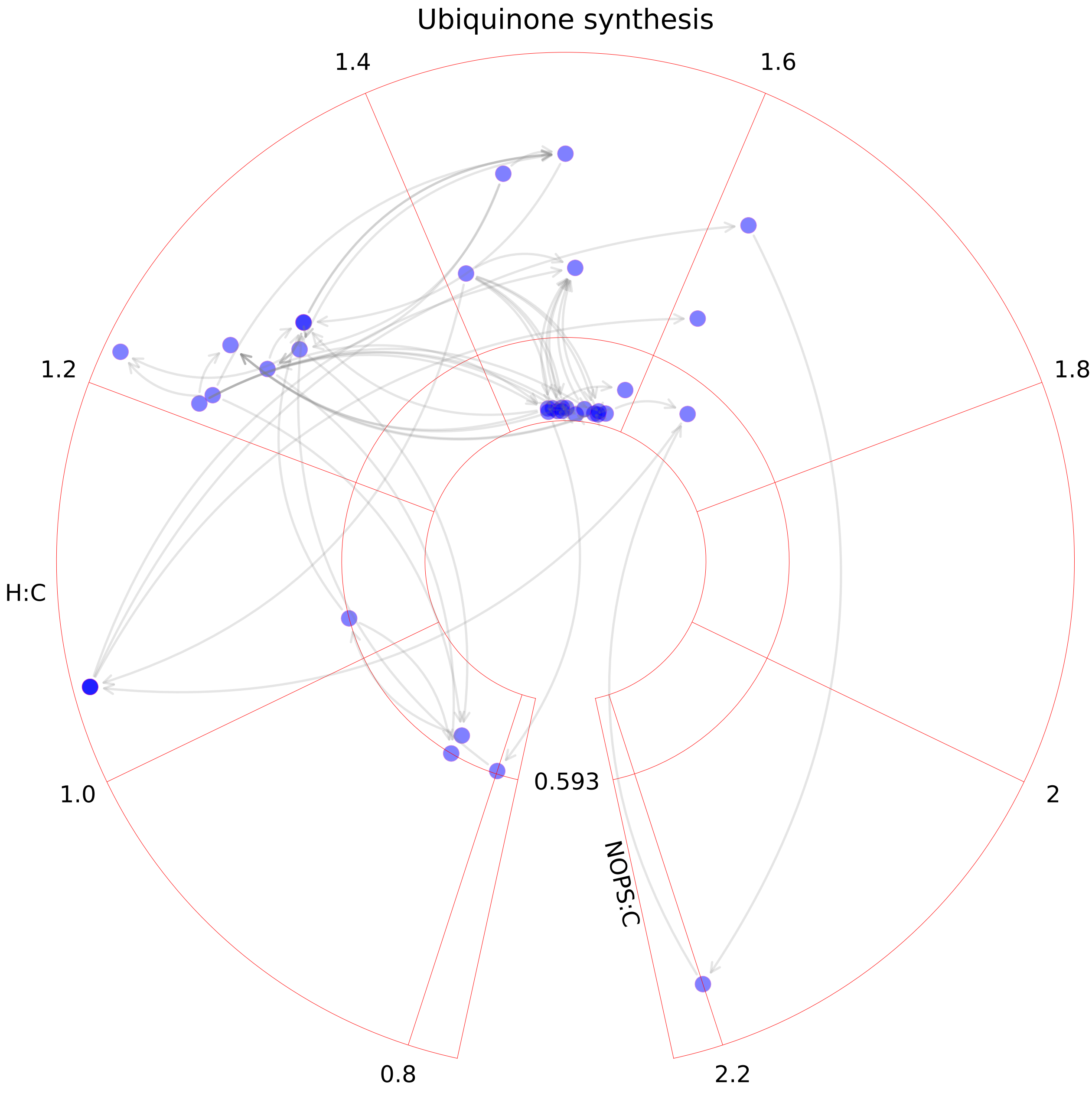

### Urea cycle.pdf

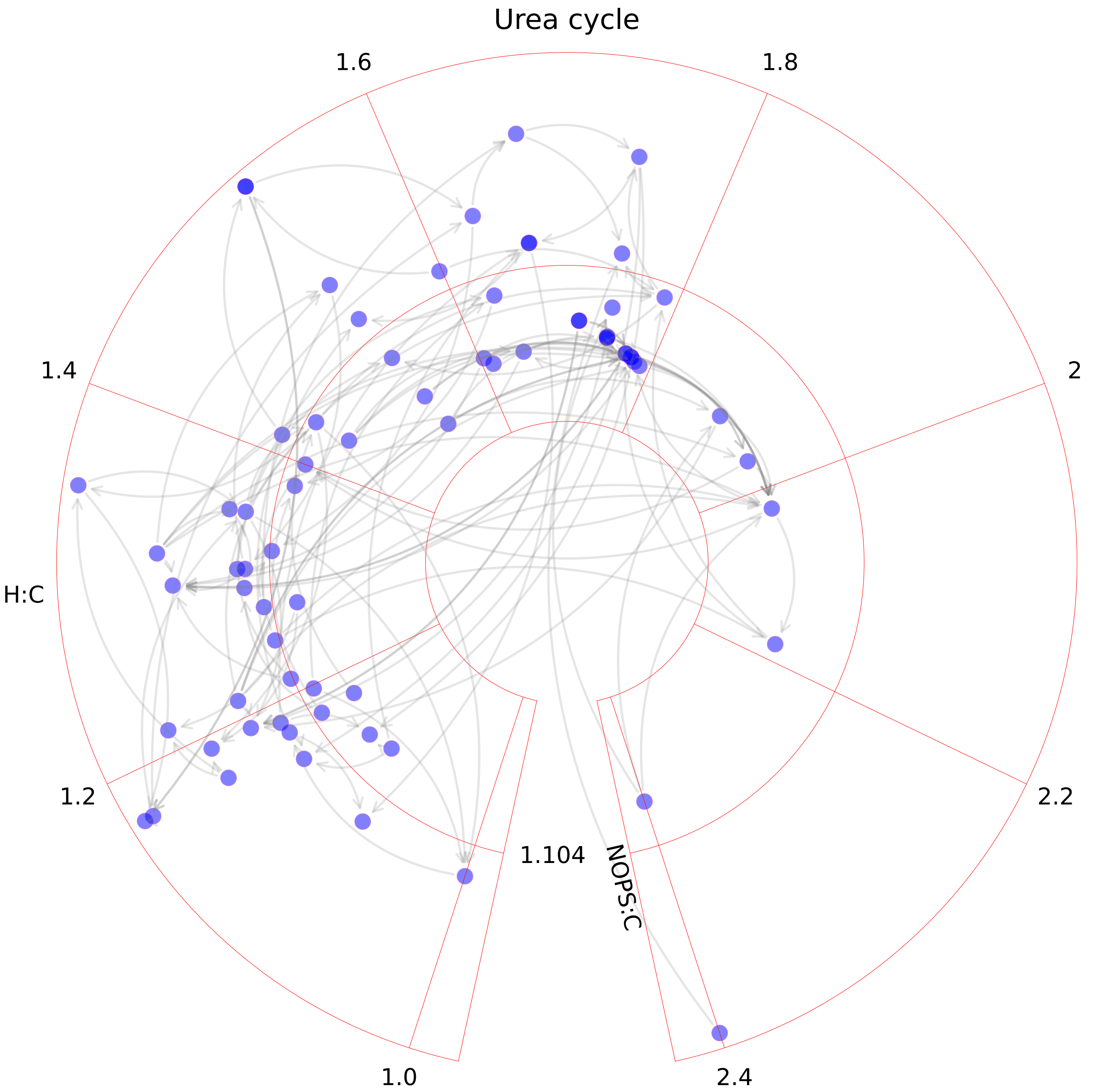

### Valine, leucine, and isoleucine metabolism.pdf

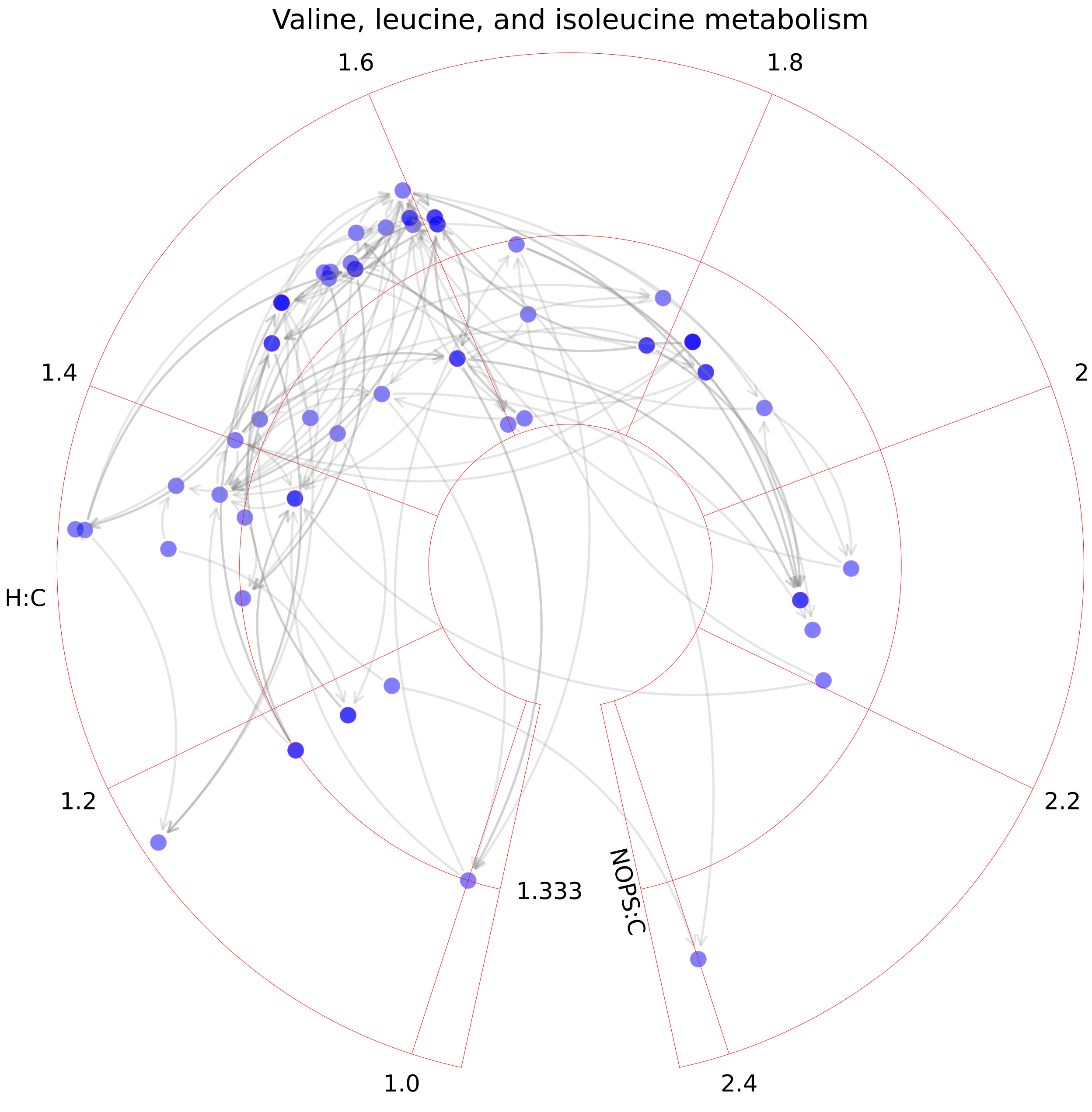

### Vitamin A metabolism.pdf

#### Vitamin A metabolism

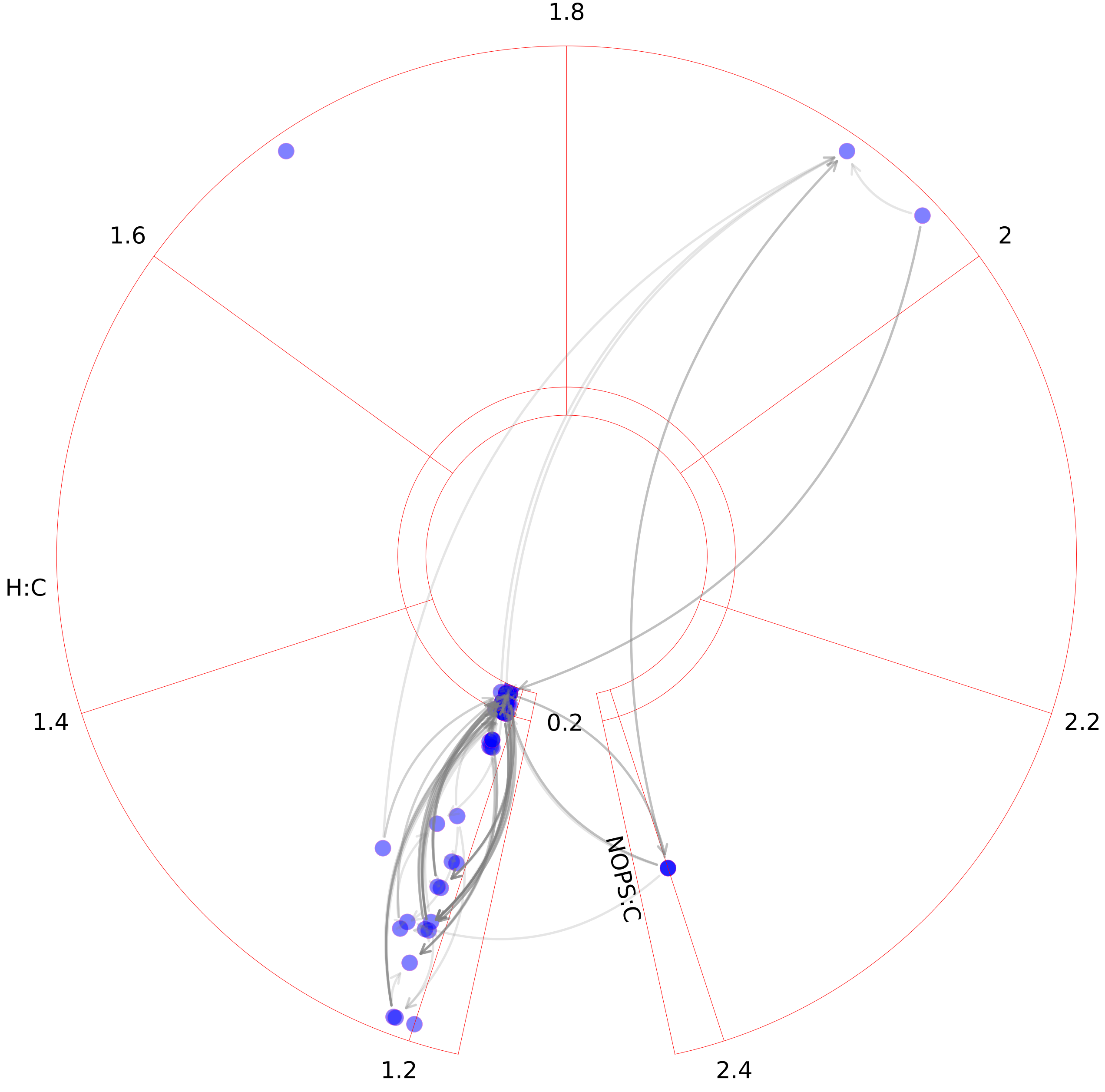

### Vitamin B2 metabolism.pdf

#### Vitamin B2 metabolism

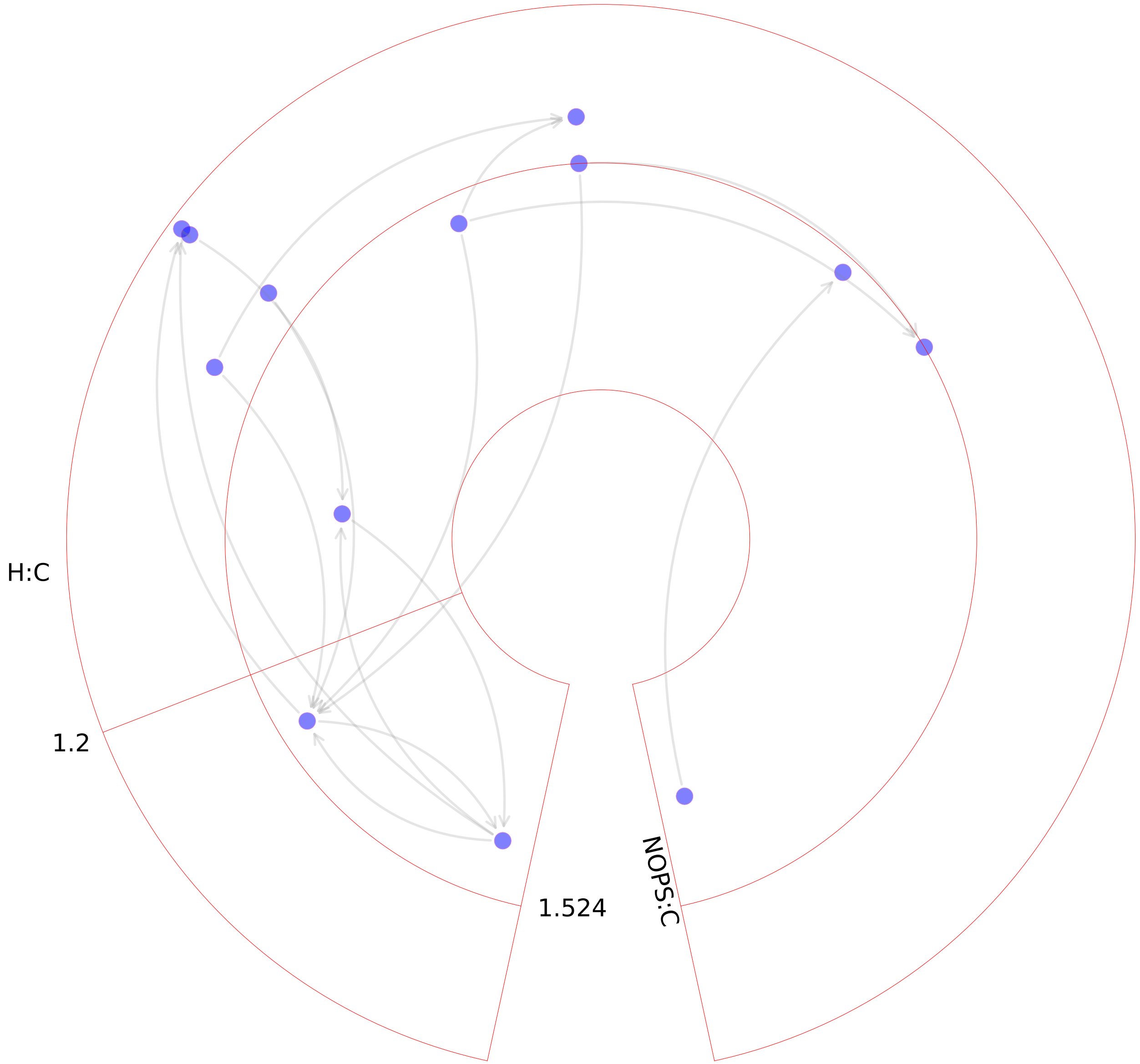

### Vitamin B6 metabolism.pdf

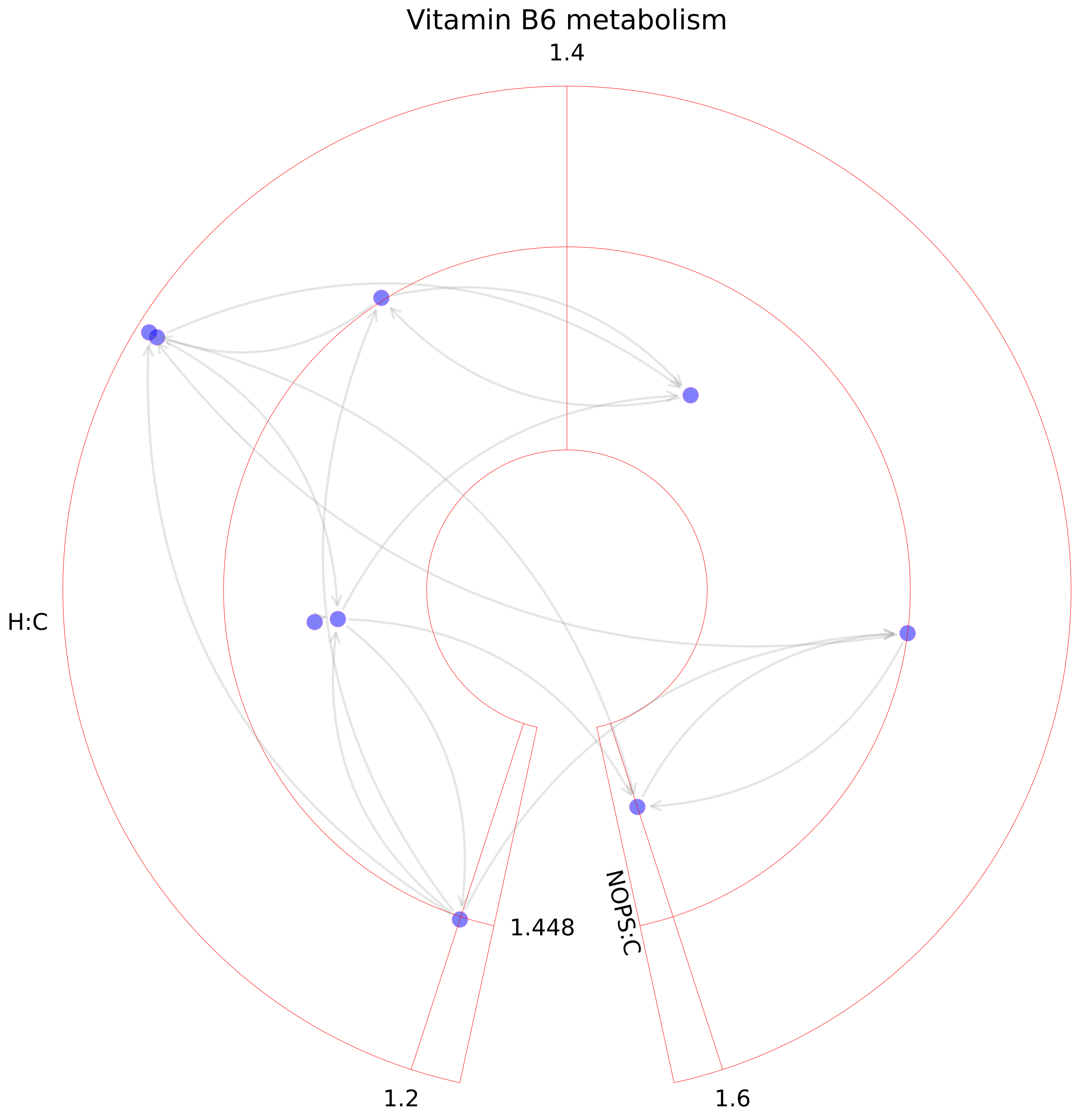

### Vitamin B12 metabolism.pdf

##### Vitamin B12 metabolism

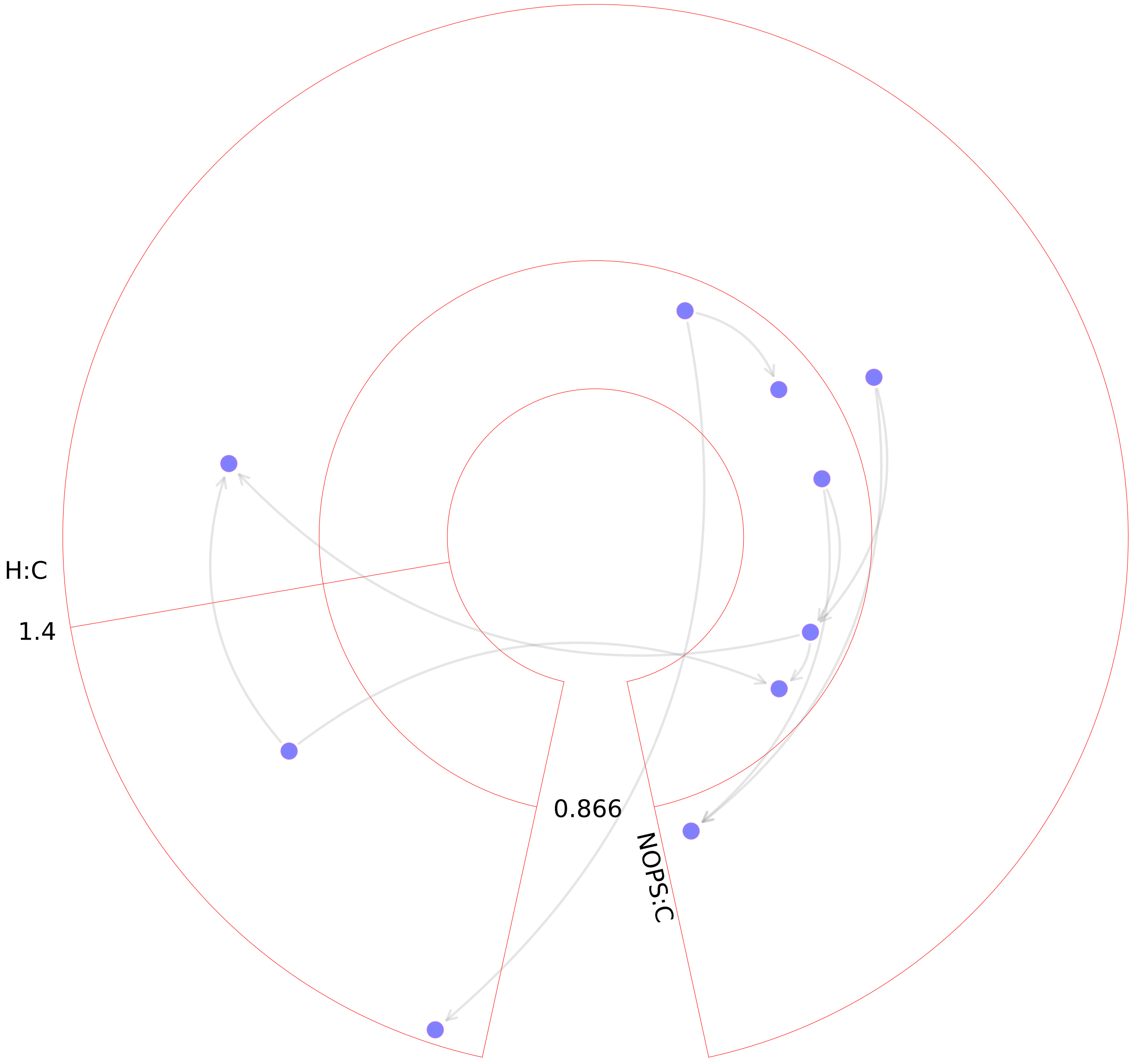

### Vitamin C metabolism.pdf

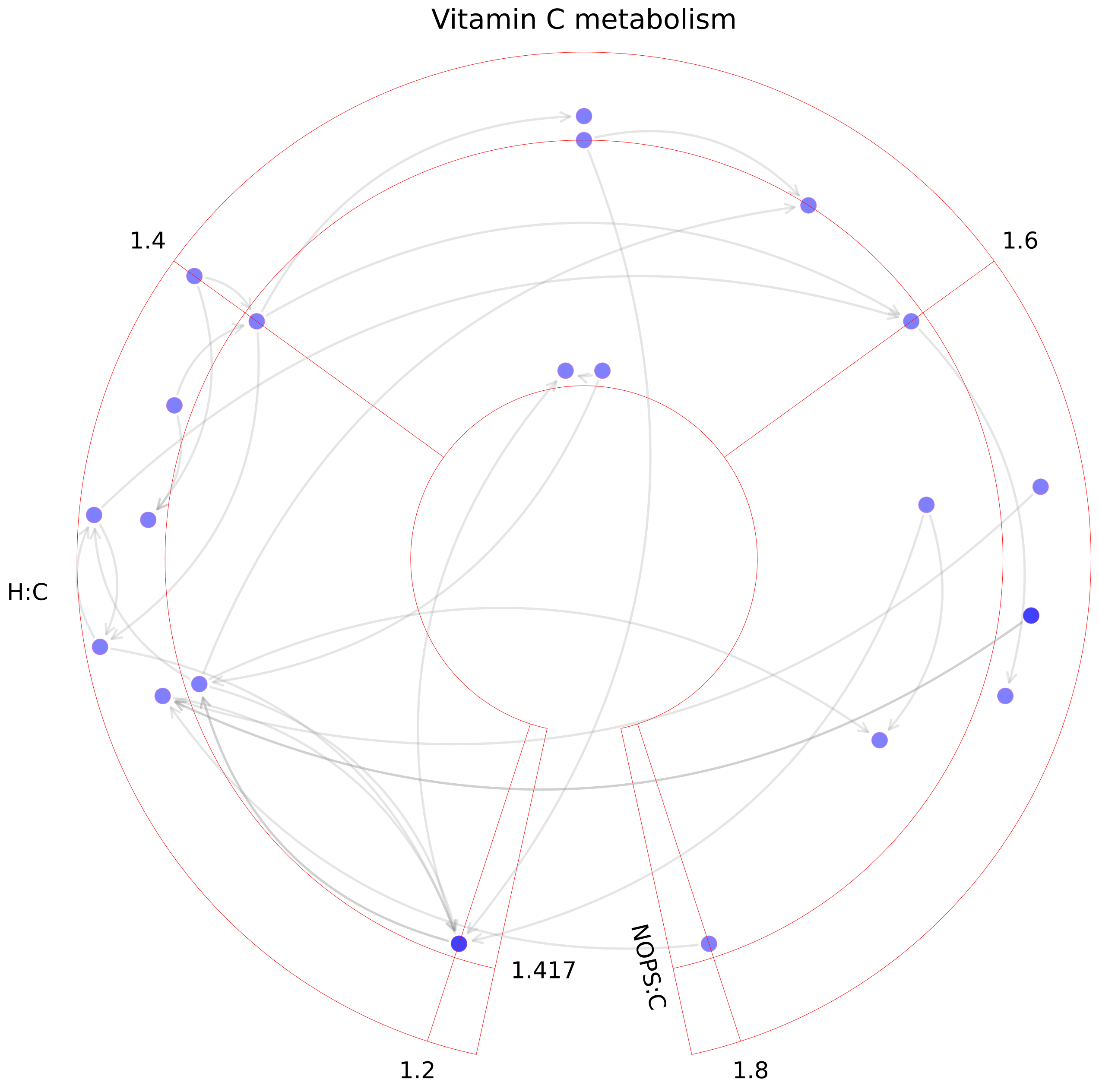

### Vitamin D metabolism.pdf

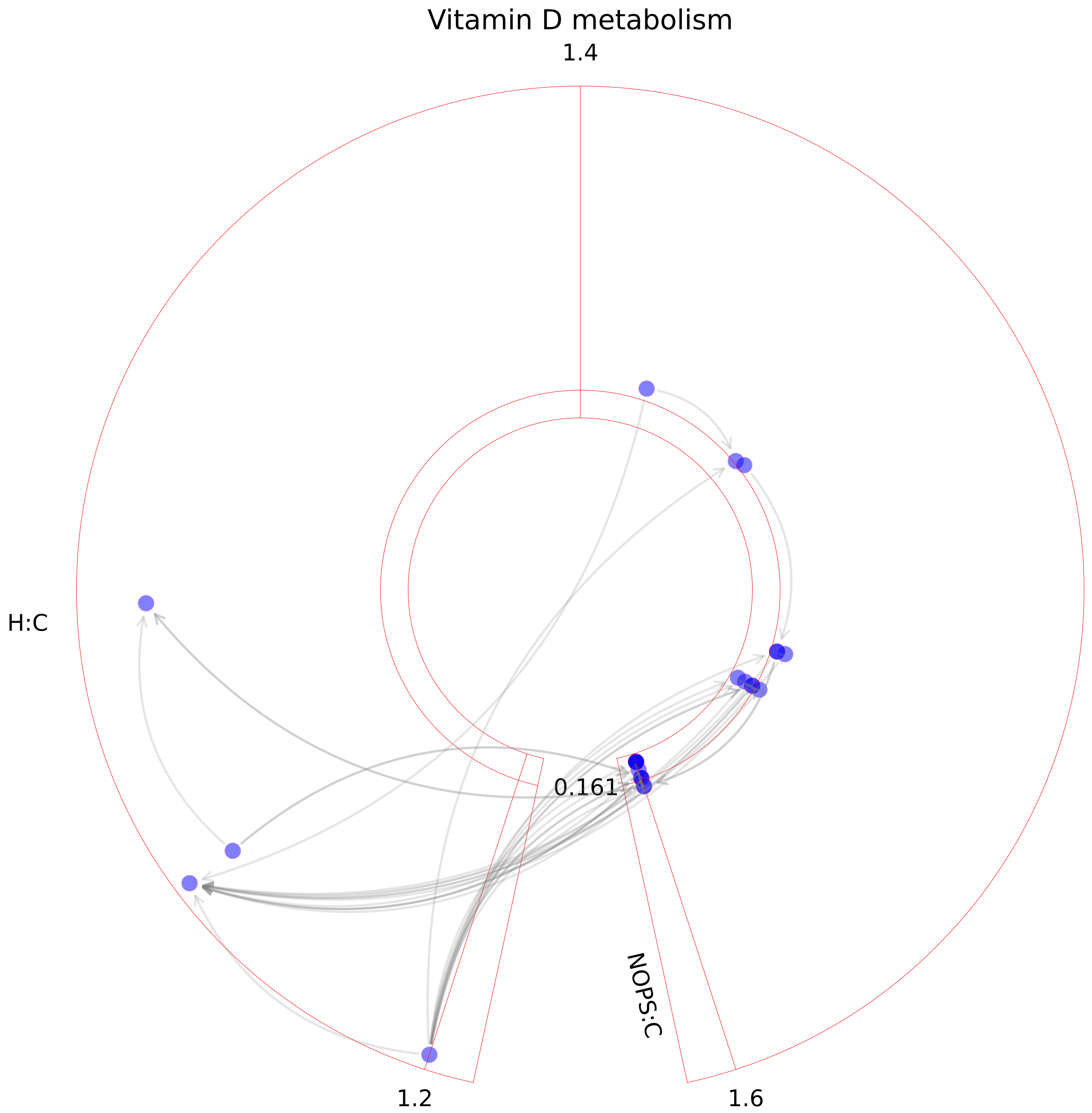

### Vitamin E metabolism.pdf

#### Vitamin E metabolism

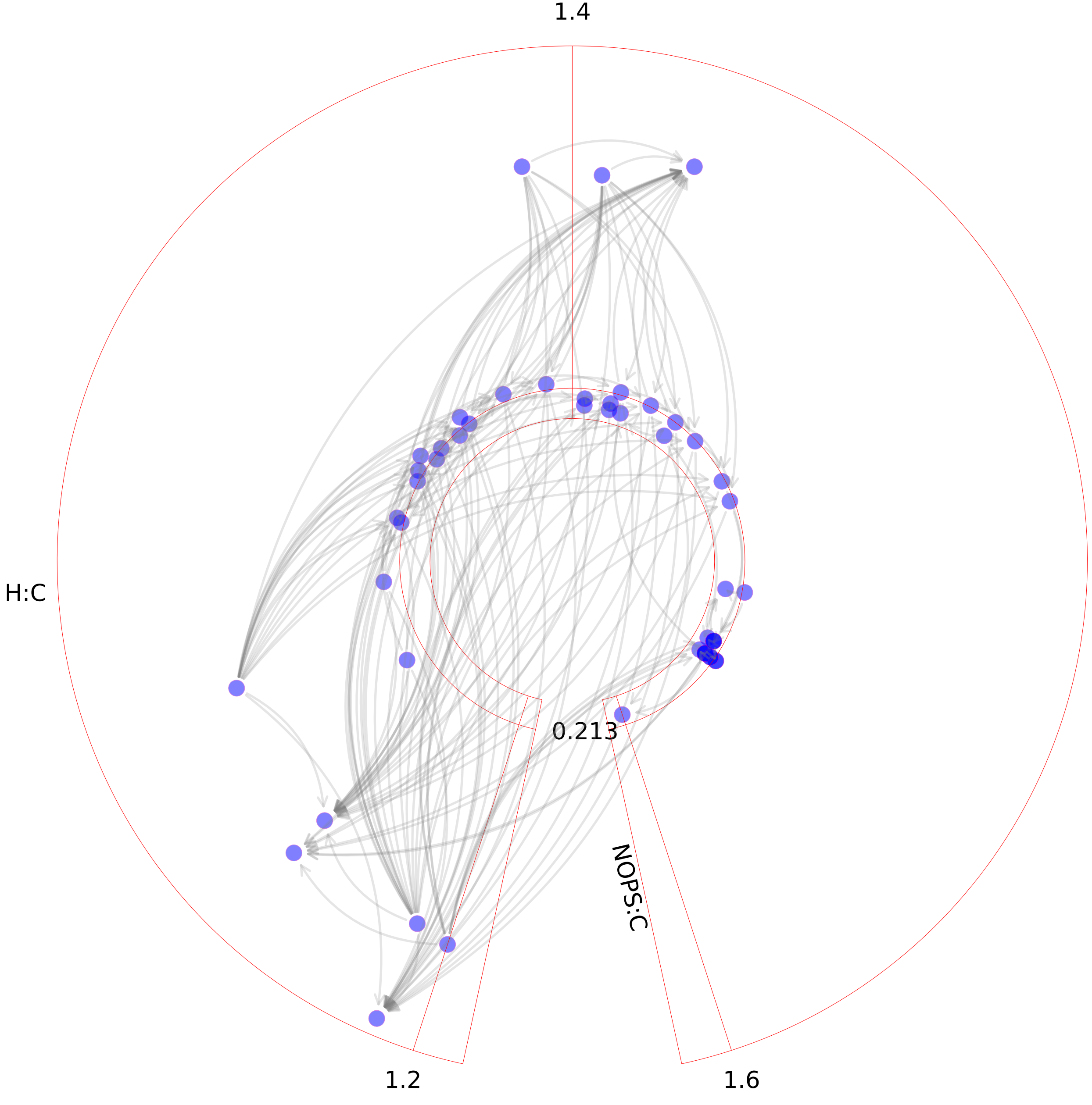

### Vitamin K metabolism.pdf

#### Vitamin K metabolism

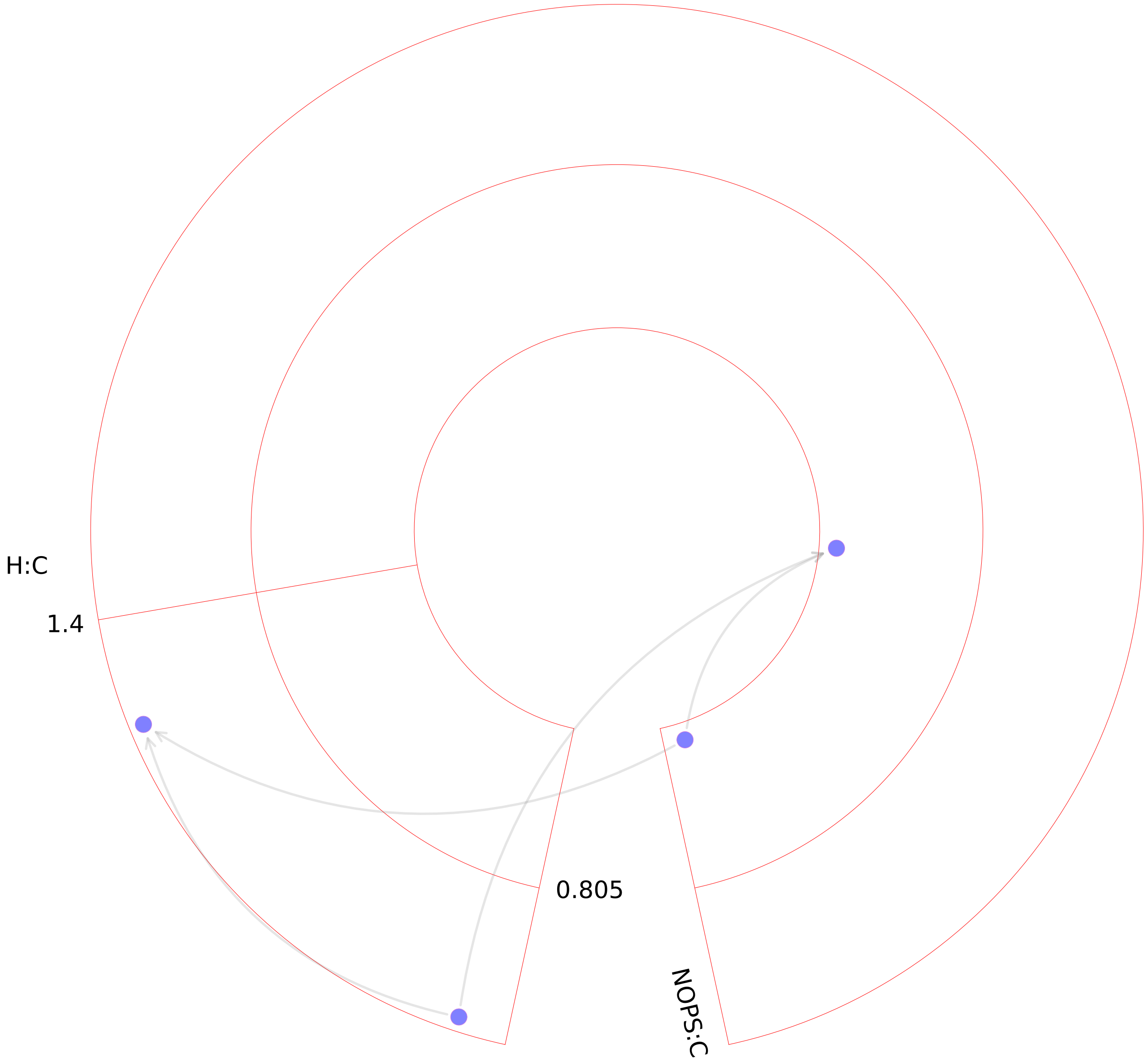

### Xenobiotics metabolism.pdf

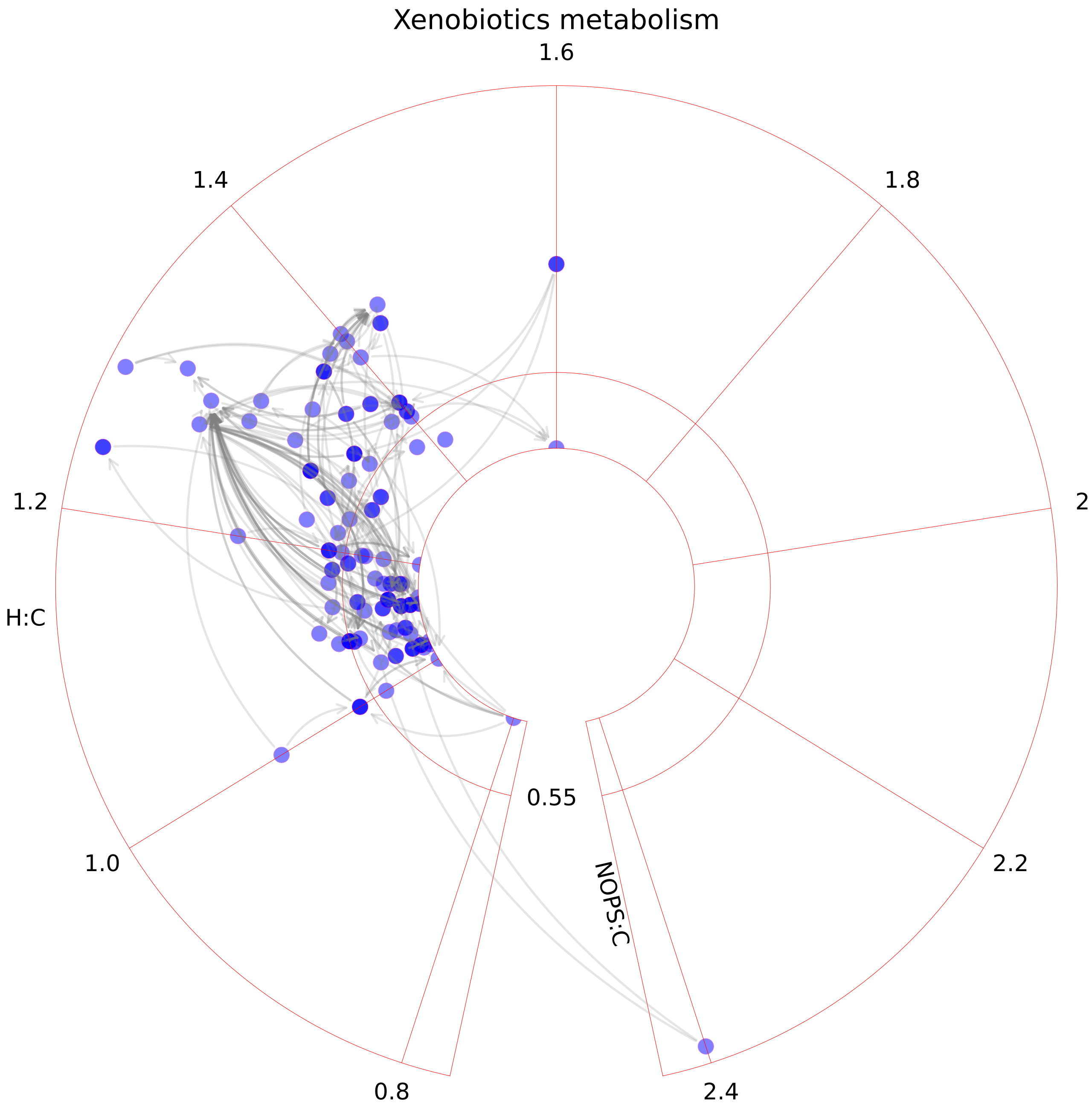
